# Pathway-based dissection of the genomic heterogeneity of cancer hallmarks’ acquisition with SLAPenrich

**DOI:** 10.1101/077701

**Authors:** Francesco Iorio, Luz Garcia-Alonso, Jonathan S. Brammeld, Iñigo Martincorena, David R. Wille, Ultan McDermott, Julio Saez-Rodriguez

## Abstract

Cancer hallmarks are evolutionary traits required by a tumour to develop. While extensively characterised, the way these traits are achieved through the accumulation of somatic mutations in key biological pathways is not fully understood. To shed light on this subject, we characterised the landscape of pathway alterations associated with somatic mutations observed in 4,415 patients across ten cancer types, using 374 orthogonal pathway gene-sets mapped onto canonical cancer hallmarks. Towards this end, we developed SLAPenrich: a computational method based on population-level statistics, freely available as an open source R package. Assembling the identified pathway alterations into sets of hallmark signatures allowed us to connect somatic mutations to clinically interpretable cancer mechanisms. Further, we explored the heterogeneity of these signatures, in terms of ratio of altered pathways associated with each individual hallmark, assuming that this is reflective of the extent of selective advantage provided to the cancer type under consideration. Our analysis revealed the predominance of certain hallmarks in specific cancer types, thus suggesting different evolutionary trajectories across cancer lineages.

Finally, although many pathway alteration enrichments are guided by somatic mutations in frequently altered high-confidence cancer genes, excluding these driver mutations preserves the hallmark heterogeneity signatures, thus the detected hallmarks’ predominance across cancer types. As a consequence, we propose the hallmark signatures as a ground truth to characterise tails of infrequent genomic alterations and identify potential novel cancer driver genes and networks.

## Introduction

The swift progression of next-generation sequencing technologies is enabling a fast and affordable production of an extraordinary amount of genome sequences. Cancer research is particularly benefiting from these advances, and comprehensive catalogues of somatic mutations involved in carcinogenesis, tumour progression and response to therapy are becoming increasingly available and ready to be exploited for the identification of new diagnostic, prognostic and therapeutic markers^1–4^. Exploration of the genomic makeup of multiple cancer types has highlighted that driver somatic mutations typically involve a few genes altered at high frequency and a long tail of more genes mutated at very low frequency^5,6^, with a tendency for both sets of genes to code for proteins involved into a limited number of biological processes^7^. As a consequence, a reasonable approach is to consider these alterations by grouping them based on a prior knowledge of the cellular mechanisms and biological pathways where the products of the mutated genes operate. This reduces the dimensionality of large genomic datasets involving thousands of altered genes into a sensibly smaller set of altered mechanisms that are more interpretable, possibly actionable in a pharmacological or experimental way^8^, and that can be used as therapeutic markers whose predictive ability is significantly improved when compared to that of genomic lesions in individual genes^9^. Additionally, this facilitates the stratification of cancer patients into informative subtypes^10^, the characterisation of rare somatic mutations^11^, and the identification of the spectrum of possible alterations underpinning a common evolutionarily successful trait acquired by a normal cell as it transforms itself into a precancerous cell and ultimately into a cancer. In two landmark papers^12,13^ these traits have been summarised into a set of 11 principles, collectively referred as the *hallmarks of cancer*.

Here we propose a computational strategy, that we call SLAPenrich (Sample-population Level Analysis of Pathway Alterations Enrichments), for characterising the set of genomically altered pathways that might contribute to the acquisition of the canonical cancer hallmarks across 10 different cancer types, via a systematic analysis of 4,415 public available cancer patients’ genomes (from the Cancer Genome Atlas). Similarly to other existing methods (such as PathScan and PathScore^14,15^), SLAPenrich aims to identify pathways that are consistently altered across the samples of a population, rather than pathways over-represented in the merged set of alterations in the population. Additionally, with respect to other existing tools, we go one step further by devising a metric to assess the predominance of alterations in pathways associated to the same canonical hallmark in each cancer type in a data-driven way. Finally, after verifying that the majority of these predominances are led by somatic mutations in established high-confidence cancer genes, we show that they are maintained when excluding these mutations from the analysis. Thus we propose to use the obtained heterogeneity signatures of cancer hallmarks as a ground truth for functionally characterising long tails of infrequent genomic alterations, across cancer types. Finally, we highlight a number of potential novel cancer driver genes and networks, identified with this approach. Our method is implemented in a publicly available R package at https://github.com/saezlab/SLAPenrich.

## Results

### Sample-population Level Analysis of Pathway Alterations Enrichments (SLAPenrich)

#### Problem definition and method overview

In the first step of our analysis we make use of SLAPenrich (Sample Level Analysis of Pathway alteration Enrichments): a computational method implementing an established statistical framework to perform pathway analyses of genomic datasets at the sample-population level. We have designed this tool as a means to characterize, in an easily interpretable way, sparse somatic mutations detected in heterogeneous cancer sample populations, which share traits of interest and are subjected to strong selective pressure, leading to combinatorial patterns.

Several computational methods have been designed to perform pathway analysis on genomic data, aiming at prioritizing sets of genomically altered genes whose products operate in the same cellular process or functional network. All the approaches proposed so far toward this aim can be categorised into two main classes^16^. The first class of approaches aims at identifying pathways whose constituent genes are significantly over-represented in the set of altered genes from all the samples of a dataset, compared with the background set of all studied genes. Many tools exist and are routinely used to perform this analysis^17–19^, sometimes incorporating additional features, such as inter-gene dependencies and signal correlations^20^, and also estimating single sample pathway deregulations based on transcriptional data^21^. To identify pathways, gene sets and gene-ontology categories that are over-represented in a selected set of genes satisfying a certain property (for example, being differentially expressed when contrasting two biological states of interest), the likelihood of their recurrence in the gene sets of interest is usually estimated. This is normally quantified through a *p-value* assignment computed through a hypergeometric (or Fisher’s exact) test, against the null hypothesis that there is no association between the pathway under consideration and the biological state yielding the selected set of genes. The test fails (producing a non-significant *p-value*) when the size of the overlap between the considered pathway and the set of genes of interests is close to that expected by random chance. The second class of approaches aims at identifying novel pathways by mapping genomic alteration patterns on large protein interaction networks. The combinatorial properties occurring among the alterations are then analyzed and used to define cost functions, for example, based on the tendency of a group of genes to be mutated in a mutually exclusive manner. On the basis of these cost functions, optimal sub-networks are identified and interpreted as novel cancer driver pathways^22–24^. However, at the moment there is no consensual method to rigorously define a mathematical metric for mutual exclusivity and compute its statistical significance, and a number of interpretations exist^22,23,25–27^.

The problem we tackle here is rather different: we want to test the hypothesis that, in a given cohort of cancer patients (or any population under evolutionary pressure), the number of samples harbouring a mutation in at least one gene belonging to a given pathway is significantly larger than its expectation (when considering the size of the measured cohort, the background mutation rate and the non-overlapping total exonic block lengths of all the genes). If this is the case, then the pathway under consideration is deemed as enriched at the population level (SLAPenriched) in relation to the whole cohort of patients. Therefore, SLAPenrich does not require somatic mutations in a pathway to be statistically enriched among those detected in an individual sample nor the merged (or aggregated) set of mutations in the population. It assumes that the mutation of a single gene of a pathway in an individual sample can be sufficient to deregulate the pathway activity. This allows pathways containing groups of genes with a tendency to be mutated in a mutually exclusive fashion (and therefore different individually mutated genes in different samples) to still be detected as enriched at the population level and further filtered based on this tendency, as additional evidence of positive selection^28^. Hence, SLAPenrich belongs roughly to the first class of computational methods described above, although it shares the mutual exclusivity consideration with the methods in the second class. More precisely, after modeling the probability of observing a genomic alteration in at least one member of a given pathway across the individual samples, SLAPenrich performs a collective statistical test against the null hypothesis that the number of samples with at least one alteration in that pathway is that expected by random chance. An additional advantage of modeling probabilities of at least an individual mutation in a given pathway (instead of, for example, the probability of the actual number of mutated genes) is that this prevents signal saturations due to hypermutated samples.

#### Statistical framework and implementation

The input to SLAPenrich is a collection of samples accounting for the mutational status of a set of genes, such as a cohort of human cancer genomes. This is modeled as a dataset where each sample consists of a somatic mutation profile indicating the status (point-mutated or wild-type) of a list of genes (Supplementary Figure S1A). For a given biological pathway *P*, each sample is considered as an individual Bernoulli trial that is successful when that sample harbours somatic mutations in at least one of the genes belonging to the pathway under consideration (Supplementary Figure S1B). The first analytical step of SLAPenrich consists in modeling the probability of such event for each individual sample. To this aim, for each sample, the likelihood of observing at least a mutation in the pathway under consideration by random chance is estimated, given the background mutation rate (for example, the number of observed mutations in the sample) and the total exonic block length of all the genes in the pathway. These individual probabilities are then aggregated in a collective test (detailed in the Methods) against the null hypothesis that the number of samples with at least one mutation in the pathway under consideration is that expected by random chance, therefore there is no association between that pathway and the disease represented by the analysed dataset.

The probability of success in each of the modeled Bernoulli trials, i.e. each sample, can be computed by either (i) a general hypergeometric model accounting for the mutation burden of the sample under consideration, the size of the gene background population and the number of genes in the pathway under consideration, or (ii) a more refined modeling of the likelihood of observing point mutations in a given pathway, accounting for the total exonic block lengths of the genes in that pathway (Supplementary Figure S1AB) and the estimated (or actual) mutation rate of the sample under consideration^29^. In addition, more sophisticated methods accounting, for example, for gene sequence compositions, trinucleotide rates, and other covariates (such as expression, chromatin state, or sequencing coverage and mappability) can be used through user-defined functions that can be easily integrated into SLAPenrich.

Once these probabilities have been computed, the expected number of samples in the population harbouring at least one somatic mutation in *P* can be estimated, and its probability distribution modeled analytically (Methods). Based on this, a pathway alteration score can be computed observing the deviance of the number of samples harbouring somatic mutations in *P* from its expectation, and its statistical significance quantified analytically (Supplementary Figure S1C). Finally, the resulting statistically enriched pathways are further filtered by looking at the tendency of their composing genes to be mutated in a mutually exclusive fashion across all the analyzed samples, as additional evidence of positive selection^22,23,30^.

A formal description of the statistical framework underlying SLAPenrich is provided in the Methods; further details are provided in the Supplementary Methods. SLAPenrich is implemented as a publicly available R package and is fully documented at https://github.com/saezlab/SLAPenrich/. It includes a visualization/report framework enabling easy exploration of outputted enriched pathways across the analyzed samples, in a way that highlights their mutual exclusivity mutation trends, and a module for the identification of core-component genes, shared by related enriched pathways. A brief description of the SLAPenrich exported functions is included in the Supplementary Note.

#### Unique features of SLAPenrich

To our knowledge, there are only two other tools enabling this type of analyses: PathScan^14^ and PathScore^15^. SLAPenrich performs comparably to both of them, showing a slightly improved ability to rank pathways containing established cancer driver genes as highly enriched. Additionally, several aspects make SLAPenrich more suitable for the analyses described in this manuscript. Particularly, PathScan does not take possible mutual exclusivity trends between patterns of mutations of genes in the same pathway into account and, in more practical terms, it requires raw sequencing data (BAM files) as input: this is quite uncomfortable for large-scale analyses where (as in our case) it is far more convenient to use available processed datasets represented through binary presence/absence matrices. PathScore uses the same mathematical framework as SLAPenrich, but the models for computing the individual pathway mutation probabilities are not fully customisable. More importantly, it is implemented as a web-application that restricts the number of individual analyses to a maximum of 10 per week. Furthermore, both PathScan and PathScore make use of fixed pathway collections from public repositories (KEGG^31^ for PathScan, and MsigDB^32^ for PathScore). In contrast, the SLAPenrich R package allows users to define and use any collection of gene sets and, by default, it employs a large pathway collection from Pathway Commons^33^ (including 2,794 pathways, covering 15,281 genes, 15 times the pathways and 3 times the genes considered by Pathscan, and twice the pathways and 1.72 times the genes of PathScore). Additionally, the SLAPenrich R package includes routines to update, on the fly, gene attributes and exonic lengths, to check and update gene nomenclatures in datasets and reference pathway gene sets, to perform mutual exclusivity sorting of binary matrices, and to identify core-components (i.e. subsets of genes leading the enrichment of different pathways).

These and other aspects are discussed in the Supplementary Methods, together with results from applying SLAPenrich to a case study Lung Adenocarcinoma Dataset to identify pathways that are differentially enriched across subpopulations of Smokers/non-smokers and mucinous/non-mucinous bronchioalveolar types, and from a systematic comparison of SLAPenrich, PathScan and PathScore (Supplementary Tables S1, S2, S3, S4, S5 and Supplementary Figure S2, S3, and S4).

### SLAPenrich analyses across different cancer types

Leveraging the capacities of SLAPEnrich, we set out to perform a systematic large-scale analysis of pathway deregulation in cancer. To this aim we used a collection of pathways from the Pathway Commons data portal (v8, 2016/04)^33^ (post-processed as detailed in the Methods), and we performed individual SLAPenrich analyses of 10 different genomic datasets containing somatic point mutations, preprocessed as described in^34^, from 4,415 patients across 10 different cancer types, from publicly available studies, in particular The Cancer Genome Atlas (TCGA) and the International Cancer Genome Consortium (ICGC). In these analyses we used a Bernoulli model to define individual pathway alteration probabilities across the single samples (equation 5). With respect to the hypergeometric models (equations 3 and 4), this formulation upon full expansion sums the individual gene mutation probabilities, each accounting for the individual gene lengths.

The analysed samples (see Methods) comprise breast invasive carcinoma (BRCA, 1,132 samples), colon and rectum adenocarcinoma (COREAD, 489), glioblastoma multiforme (GBM, 365), head and neck squamous cell carcinoma (HNSC, 375), kidney renal clear cell carcinoma (KIRC, 417), lung adenocarcinoma (LUAD, 388), ovarian serous cystadenocarcinoma (OV, 316), prostate adenocarcinoma (PRAD, 242), skin cutaneous melanoma (SKCM, 369), and thyroid carcinoma (THCA, 322). Results from all these individual SLAPenrich analyses are contained in Supplementary Table S6.

We tested the stability of SLAPenrich with respect to variations in mutation calling reliability, evaluating the effect of random noise and errors at the level of the SLAPenrich input matrices. To this aim, we increasingly introduced (respectively removed) uniformly distributed false-positives (respectively true-positives) mutations, in each of the 10 analysed genomic datasets. This was performed simulating a reduction of mutation call sensitivity (respectively, specificity) to 95, 80, 70, and 50%, producing 10 noise-inflated versions of the considered dataset for each reduction level. Subsequently, we executed a SLAPenrich analysis on each of these datasets and compared the sets of outputted pathways with those obtained when running SLAPenrich on the corresponding original datasets. Results from these analyses (detailed in the Supplementary Methods) are shown in Supplementary Figure S5. They highlight that the output of SLAPenrich is highly stable with respect to the introduced noise (median area under the Receiver Operating Characteristic (ROC) curves stably over 0.995 for all the tested ratios of Variants False Positives, and over 0.99 for all the tested ratios of Variants False Negatives).

### Mapping pathway enrichments onto canonical cancer hallmarks

Subsequently, we reasoned that since the main role of cancer driver alterations is to enable cells to achieve a series of phenotypic traits termed the ‘cancer hallmarks’^12,13^, that can be linked to gene mutations^35^, it would be informative to group the pathways according to the hallmark with which they are associated. Towards this end, through a computer-aided manual curation (see Methods and Supplementary Table S7) we were able to map 374 gene-sets (from the most recent release of pathway commons^33^) to 10 cancer hallmarks^12,13^ (Figure 1AB), for a total number of 3,915 genes (included in at least one gene set associated to at least one hallmark; Supplementary Table S8). The vast majority (99%, 369 sets) of the considered pathway gene-sets were mapped on two hallmarks at most, and 298 of them (80%) was mapped onto one single hallmark (Figure 1C). Regarding the individual genes contained in at least one pathway gene-set, about half (49%) were associated with a single hallmark, 22% with two, 12% with three, and 7% with four (Figure 1D). Finally, as shown in Figure 1E, the overlaps between the considered pathway gene-sets was minimal (74% of all the possible pair-wise Jaccard indexes was equal to 0 and 99% < 0.2). In summary, our manual curation produced a non-redundant matching in terms of both pathways- and genes-hallmarks associations. Mapping pathway enrichments into canonical cancer hallmarks through this curation covered 46% of significant results on average across cancer types (Supplementary Figure S6A).

**Figure 1.**
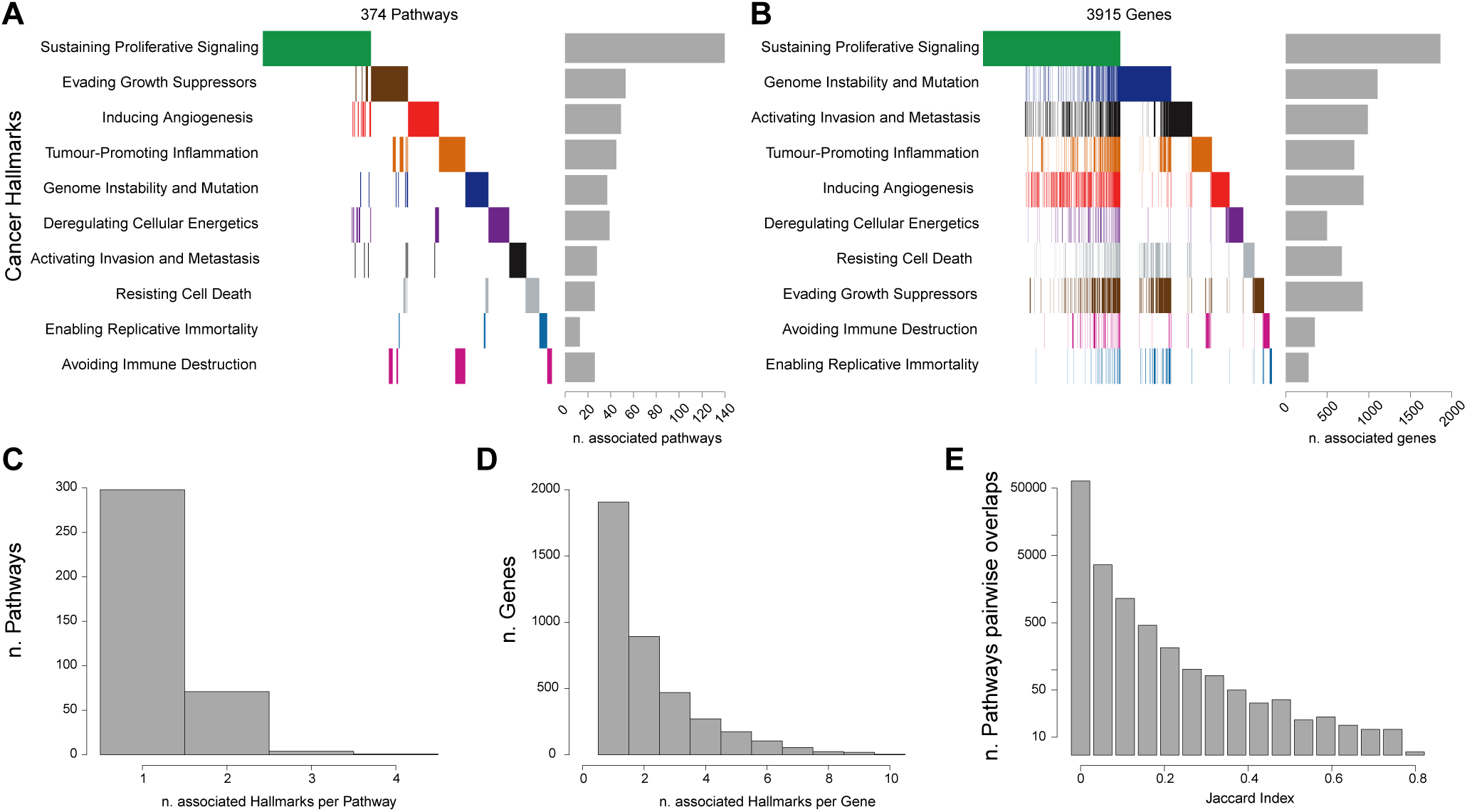
Manually curated mapping between genes, pathways and hallmarks (properties): (A) Heatmap with cancer hallmarks on the rows, pathways gene sets on the columns. A coloured bar in position (*i*, *j*) indicates that the *j*-th pathway is associated with the *i*-th hallmark; bar diagram on the right shows the number of pathways associated with each hallmark. (B) Heatmap with cancer hallmarks on the rows and genes on the columns. A coloured bar in position (*i*, *j*) indicates that the *j*-th gene is contained in at least one pathway associated with the *i*-th hallmark (thus associated with the *i*-th hallmark); bar diagram on the right shows the number of genes associated with each hallmark. (C) Number of associated hallmarks per pathways: the majority of the pathways is associated with 1 hallmark. (D) Number of associated hallmarks per gene: the majority of the genes is associated with less than 3 hallmarks. (E) Distribution of Jaccard similarity scores (quantifying the extent of pair-wise overlaps) computed between pairs of pathway gene sets.

We observed a weak correlation (*R* = 0.53, *p* = 0.11) between the number of hallmark-associated (HMA) enriched pathways across the different analyses and the number of available samples in the analysed dataset (Supplementary Figure S6B), but a down-sampled analysis showed that our results are not broadly confounded by the sample sizes (see Methods and Supplementary Figure S6C).

We investigated how our HMA-pathway enrichments capture known tissue-specific cancer driver genes. To this aim, we used a list of high-confidence and tissue-specific cancer driver genes^34,36^ (from now high-confidence Cancer Genes, HCGs, assembled as described in the Methods). We observed that the majority of the HCGs was contained in at least one SLAPenriched HMA-pathway, across the 10 different tissues analyses (median percentage = 63.5, range = 88.5%, for BRCA, to 28.7% for SKCM) (Supplementary Figure S6D).

Interestingly, we found that the number of HMA-SLAPenriched pathways per cancer type (median = 130, range = 55 for PRAD, to 200 for BRCA and COREAD) was independent of the average number of mutated genes per sample across cancer types (median = 46, range from 15 for THCA to 388 for SKCM) with a Pearson correlation *R* = 0.16 (*p* = 0.65), Figure 2A, as well as from the number of high confidence cancer driver genes (as predicted in^36^, median = 100, range from 33 for THCA to 251 for SKCM, Figure 2B). Particularly, THCA has the lowest average number of mutations per sample (15.03), but there are 4 tissues with a lower number of HMA-pathways enriched. In contrast, SKCM has the highest average number of point mutations per sample (387.63), but the number of affected pathways is less than half of those of BRCA and GBM (82 enrichments against an average of 191), which have on average less than 100 mutations per sample (Figure 2A). GBM, OV, KIRC, PRAD and BRCA are relatively homogeneous with respect to the average number of somatic mutations per sample (mean = 41.03, from 34.76 for KIRC to 45.95 for PRAD) but when looking at the number of enriched HMA-pathways for this set of cancer types we can clearly distinguish two separate groups (Figure 2A). The first group includes BRCA and GBM that seem to have a more heterogeneous set of processes impacted by somatic mutations (average number of SLAPenriched pathways = 191) with respect to the second group (63 SLAPenriched pathways on average). These results suggest that there is a large heterogeneity in the number of processes deregulated in different cancer types that is independent of the mutational burden. This might also be indicative of different subtypes with dependencies on different pathways (and at least for BRCA this is expected) but could also be biased by the composition of the analysed cohorts being representative of selected subtypes only.

**Figure 2.**
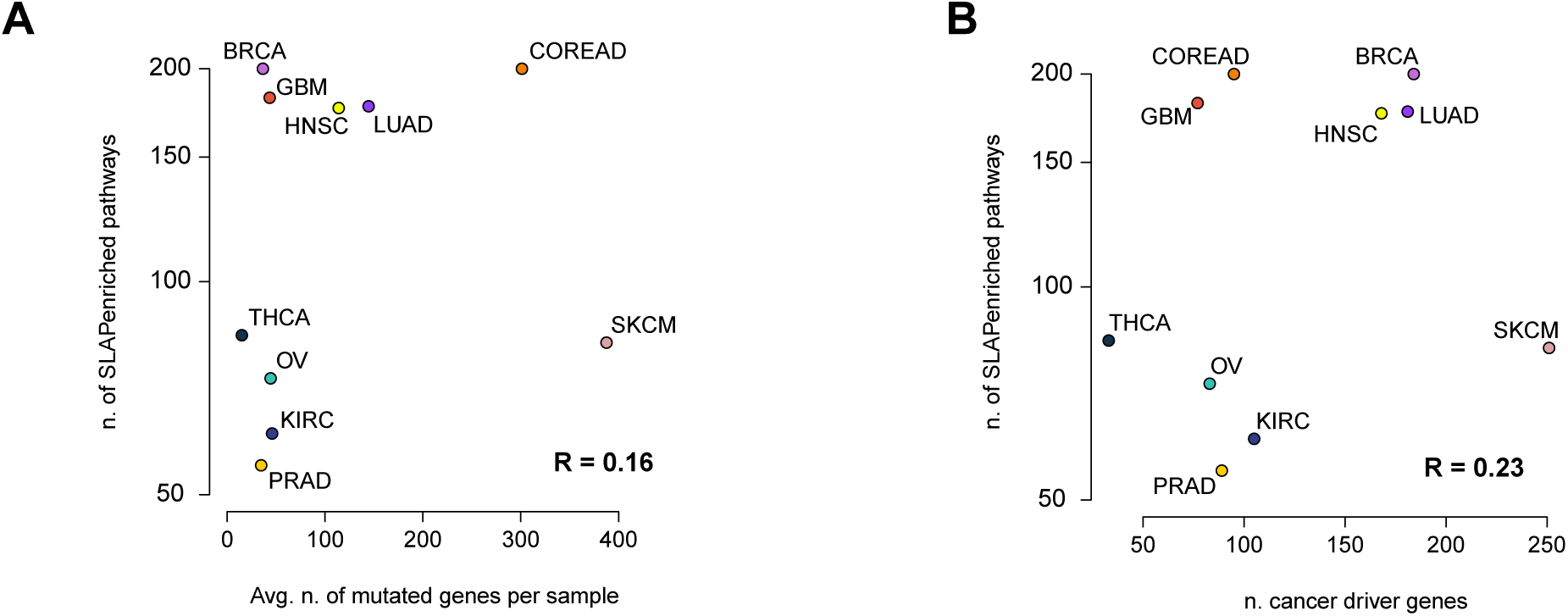
Number of SLAPenrichments versus mutation burdens and number of established cancer genes: (A) Number of pathways enriched at the population level across cancer types compared with the average number of mutated genes and (B) the average number of high confidence cancer driver genes.

### Genomic heterogeneity of cancer hallmarks’ acquisition across cancer types

Inspecting the sets of enriched HMA-pathways across the performed analyses allowed us to explore how different cancer types might acquire the same hallmark by selectively altering different pathways. Heatmaps in Figure 3, and Supplementary Figure S7 (one per each hallmark) show a different level of enrichment of pathways associated with the same hallmark across different tissues, with clearly distinguishable patterns and well-defined clusters.

**Figure 3.**
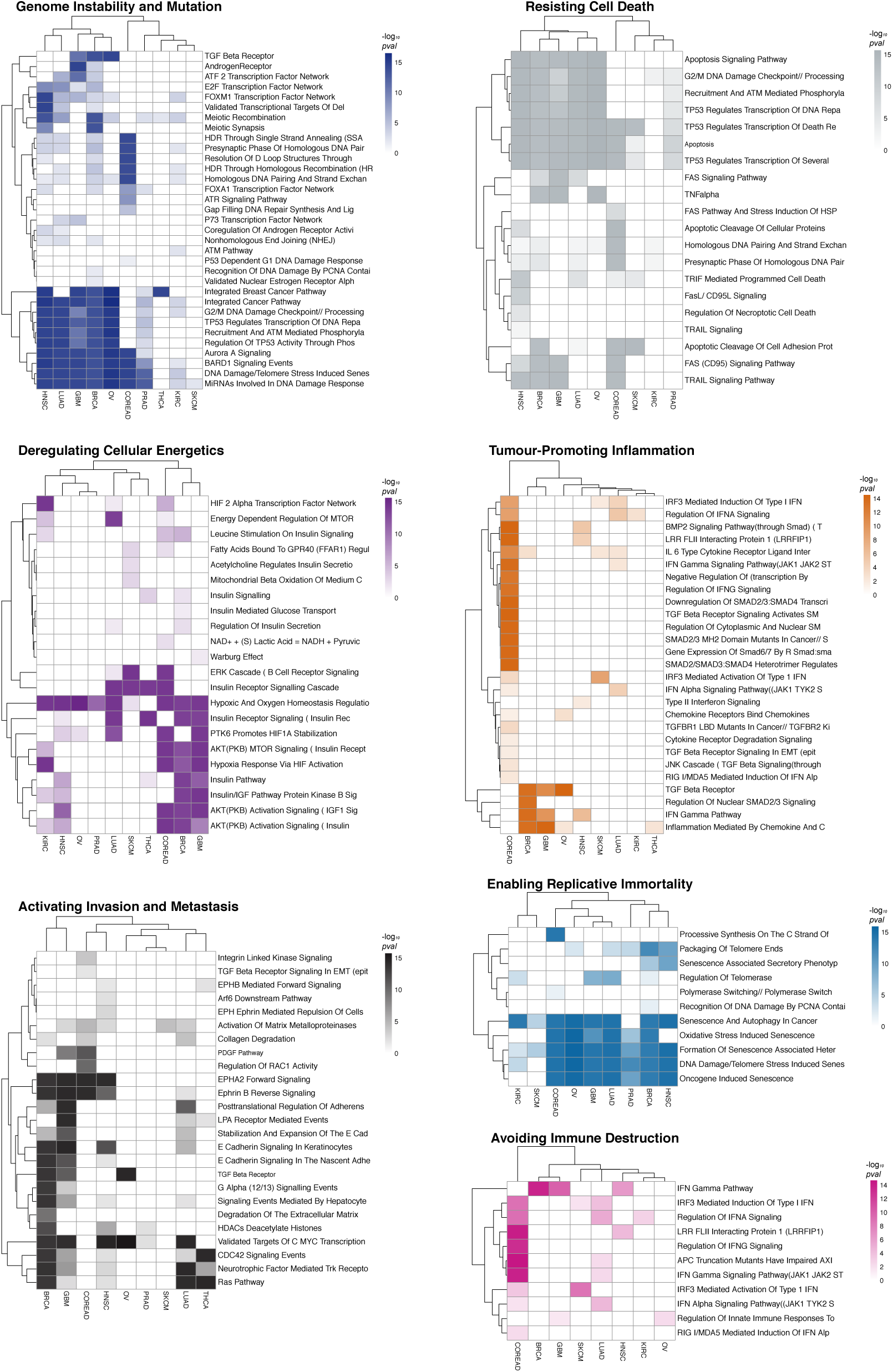
Heterogeneity of hallmark acquisition across cancer types: Heatmaps showing pathways enrichments at the population level across cancer types for individual hallmarks (representative cases). Color intensities correspond to the enrichment significance. Cancer types and pathways are clustered using a correlation metric. See also Supplementary Figure 7.

As an example, the heatmap related to the *Genome Instability and mutation* hallmark shows that BRCA, OV, GBM, LUAD and HNSC might achieve this hallmark by selectively altering a group of pathways related to homologous recombination deficiency, whose prevalence in BRCA and OV is established^37^. This deficiency has been therapeutically exploited recently and translated into a clinical success thanks to the introduction of PARP inhibition as a very selective therapeutic option for these two cancer types^38^.

Pathways preferentially altered in BRCA, OV, GBM, LUAD and HNSC include *G2/M DNA Damage Checkpoint // Processing Of DNA Double Strand Break Ends*, *TP53 Regulates Transcription Of DNA Repair Genes* and other signaling networks related to BRCA1/2 and its associated RING Domain 1 (BARD1). Conversely, the *Androgen receptor* pathway, known to regulate the growth of glioblastoma multiforme (GBM) in men^39^ is exclusively and preferentially altered in this cancer type.

The acquisition of the *Genome Instability and mutation* hallmark seems to be dominated in COREAD by alterations in the *HDR Through Single Strand Annealing (SSA)*, *Resolution Of D Loop Structures Through Synthesis Dependent Strand Annealing (SDSA)*, *Homologous DNA Pairing And Strand Exchange* and other pathways more specifically linked to a microsatellite instability led hypermutator phenotype, known to be prevalent in this cancer type^40^.

Finally, the heatmap for *Genome Instability and Mutation* shows nearly no enriched pathways associated to the acquisition of this hallmark in SKCM. This is consistent with the high burden of mutations observed in melanoma being the effect of this hallmark rather than leading its acquisition. In fact, genomic instability in SKCM originates from cell extrinsic processes such as UV light exposure^41^.

The maintenance of genomic integrity is guarded by a network of damage sensors, signal transducers, and mediators, and is regulated through changes in gene expression. Recent studies show that miRNAs play a crucial role in the response to UV radiation in skin cells^42^. Our analysis strikingly detects *MiRNAs Involved In DNA Damage Response* as the unique pathway associated to *Genome instability and mutation* in SKCM. This suggests that mutations in this pathway, involving ATM (as the most recurrently mutated gene, and known to induce miRNA biogenesis following DNA damage^43^), impair the ability of melanocytes to properly respond to insults from UV light and may have a significant role in the tumourigenesis of melanoma.

The *Avoiding Immune destruction* heatmap (Figure 3) highlights a large number of pathways selectively enriched in COREAD, whereas very few pathways associated to this hallmark are enriched in the other analysed cancer types. This could explain why immunotherapies, such as PD-1 inhibition, have a relatively low response rate in COREAD when compared to, for example, non-small cell lung cancer^44^, melanoma^45^ or renal-cell carcinoma^46^. In fact, response to PD-1 inhibition in COREAD is limited to tumours with mismatch-repair deficiency, perhaps due to their high rate of neoantigen creation^47^.

Moreover, in the context of COREAD, the *Tumor-promoting inflammation* heatmap (Figure 3) also highlights several pathways predominantly and very specifically altered in this cancer type. Chronic inflammation is a proven risk factor for COREAD and studies in animal models have shown a dependency between inflammation, tumor progression and chemotherapy resistance^48^. Indeed, a number of clinical trials evaluating the utility of inflammatory and cytokine-modulatory therapies are currently underway in colorectal cancer^49,50^. Interestingly, according to our analysis this hallmark is acquired by SKCM by exclusively preferentially altering IRF3 related pathways.

Several other examples would be worthy of mention. For example, the detection of the *Warburg effect* pathway contributing to the acquisition of the *Deregulating cellular energetics* hallmark in GBM only (Figure 3). The Warburg effect is a unique bioenergetic state of aerobic glycolysis, whose reversion has been recently proposed as an effective way to decrease GBM cell proliferation^51^. Additionally, the pathway *Formation of senescence-associated heterochromatin*, associated to the *Enabling replicative immortality* hallmark is enriched in multiple cancer types. Genomic alterations in this pathway have not been linked to cancer so far. More interestingly the enrichment of this pathway, across cancer types, is not driven by any established cancer gene.

Finally, we quantified the diversity of altered pathways mapped to each cancer hallmark in a given tumor type, via a cumulative heterogeneity score (CHS). The CHS of a hallmark is computed as the proportion of the pathways associated to that hallmark that are significantly enriched. We hypothesize that a large CHS points to the exploitation of many evolutionary trajectories pursued to acquire a defined hallmark. This might suggest that the hallmark with a higher CHS is more advantageous evolutionary than others for the cancer type under consideration.

The pattern of CHSs per cancer hallmark in a cancer type gives its *hallmark heterogeneity signature* (Figure 4). Results show consistency with the established predominance of certain hallmarks in determined cancer types such as, for example, a high CHS for *Genome instability and mutation* in BRCA and OV^52^, for *Tumour-promoting inflammation* and *Avoiding immune-destruction* in COREAD^53^. Lastly, and as expected for *Sustaining proliferative-signaling* and *Enabling replicative immortality*, the key hallmarks in cancer initiation^12^, high CHSs are observed across the majority of the analysed cancer types.

**Figure 4.**
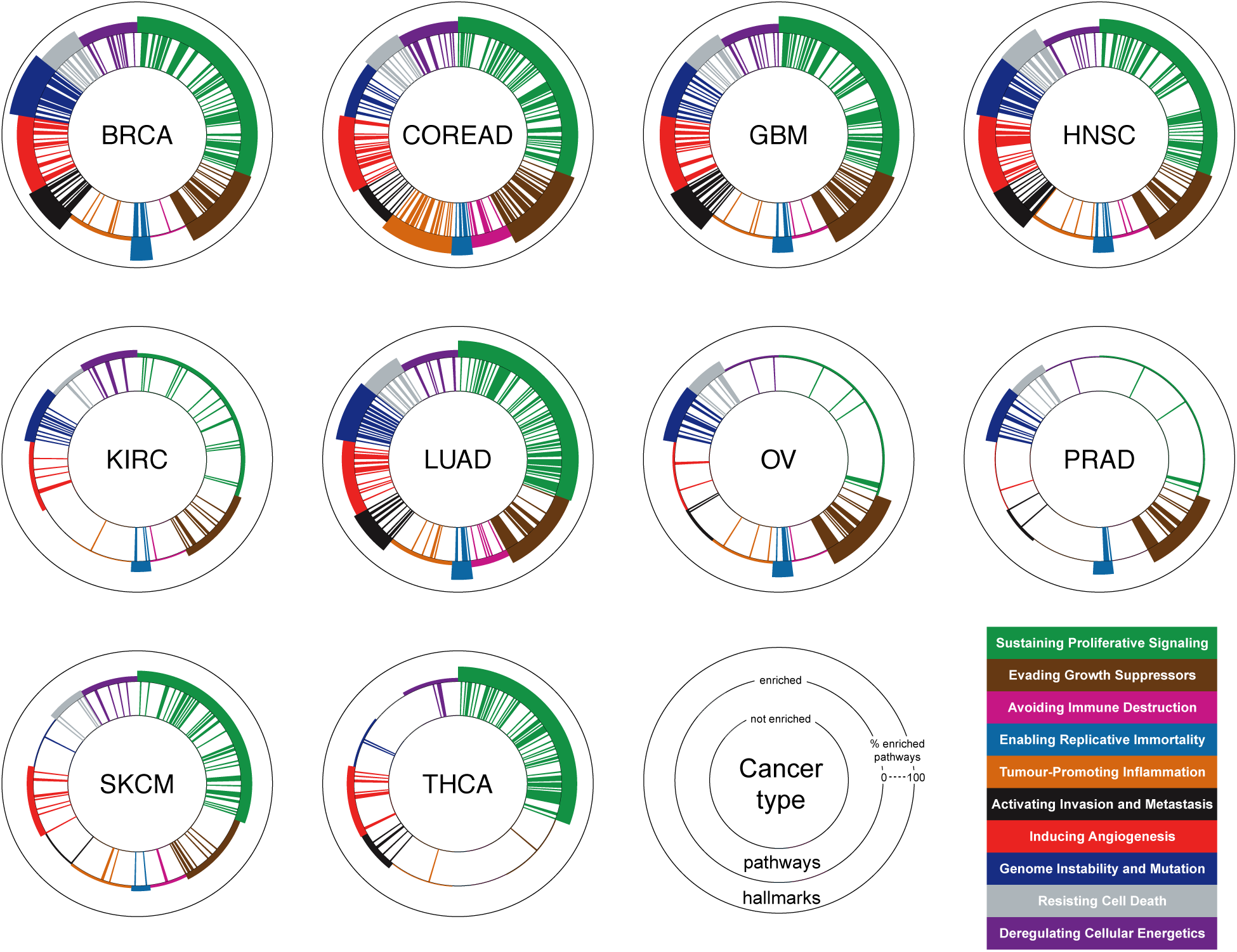
Cancer hallmark heterogeneity signatures: Each cancer hallmark signature plot is composed of three concentric circles. Bars between the inner and middle circles indicate pathways, bars between the middle and external circle indicate cancer hallmarks. Different colors indicate different cancer hallmarks. Pathway bars are coloured based on their hallmark association. The presence of a pathway bar indicates that the corresponding pathway is enriched at the population level (FDR < 5%, EC = 50%) in the cancer type under consideration. The thickness of the hallmark bars are proportional to the ratio of enriched pathways over those associated with that hallmark.

Taken together, these results show the potential of our pipeline to perform systematic landscape analyses of large cohorts of cancer genomes. In this case, this is very effective in highlighting commonalities and differences in the acquisition of the cancer hallmarks across tissue types, confirming several known relations between cancer types, and pinpointing preferentially altered pathways and hallmark acquisitions.

### Hallmark heterogeneity analysis points at novel cancer driver genes and networks

To investigate the potential of our computational method in identifying novel cancer driver genes and networks, we evaluated first to what extent the identified enriched HMA-pathways were dominated by somatic mutations in established high-confidence cancer genes (HCGs)^36^ across cancer types. To this aim, for each pathway *P* enriched in a given cancer type *T*, we computed an HCG-dominance score as the ratio between the number of samples with mutations in HCGs in *P* and the number of samples with mutations in any gene in *P*. Results of this analysis are shown in Supplementary Figures S7 and S8. We observed a median of 15% of pathway enrichments, across hallmarks, with a HCG-dominance score ¡ 50%, thus not led by somatic mutations in HCGs (range from 9% for *Deregulating Cellular Energetics* to 21% for *Genome Instability and Mutation*). Additionally, a median of 3% of pathway enrichments had a null HCG-dominance, thus did not involve somatic mutations in HCGs (range from 0.25% for *Evading Growth Suppression* to 15% for *Avoiding Immune Destruction*). Across all the hallmarks, the cancer type with the lowest median HCG-dominance was KIRC (33%), whereas that with the highest was THCA (91%).

Subsequently, we re-analysed the TCGA data excluding all the variants involving HCGs from each cancer type (from now the *filtered analysis*). Results from this exercise (Figure 5, Supplementary Table S9 and Supplementary Figure S10), showed that the majority of the enrichments identified in the original analyses (on the unfiltered genomic datasets) were actually led by alterations in the HCGs (consistent with their condition of high reliable cancer genes). The average ratio of retained enrichments in the filtered analyses across cancer types (maintained enrichments (MA) in Figure 5 and Supplementary Figure S10) was 21%, (range from 2.1% for GBM to 56.2% for COREAD). However, several HMA-pathway enrichments (some of which did not include any HCGs) were still detected in the filtered analysis and, most importantly, the corresponding hallmark heterogeneity signatures were largely conserved across the filtered and unfiltered analyses for most of the cancer types, with coincident top fitting hallmarks and significantly high over-all correlations (Figure 5, Supplementary Figure S10).

**Figure 5.**
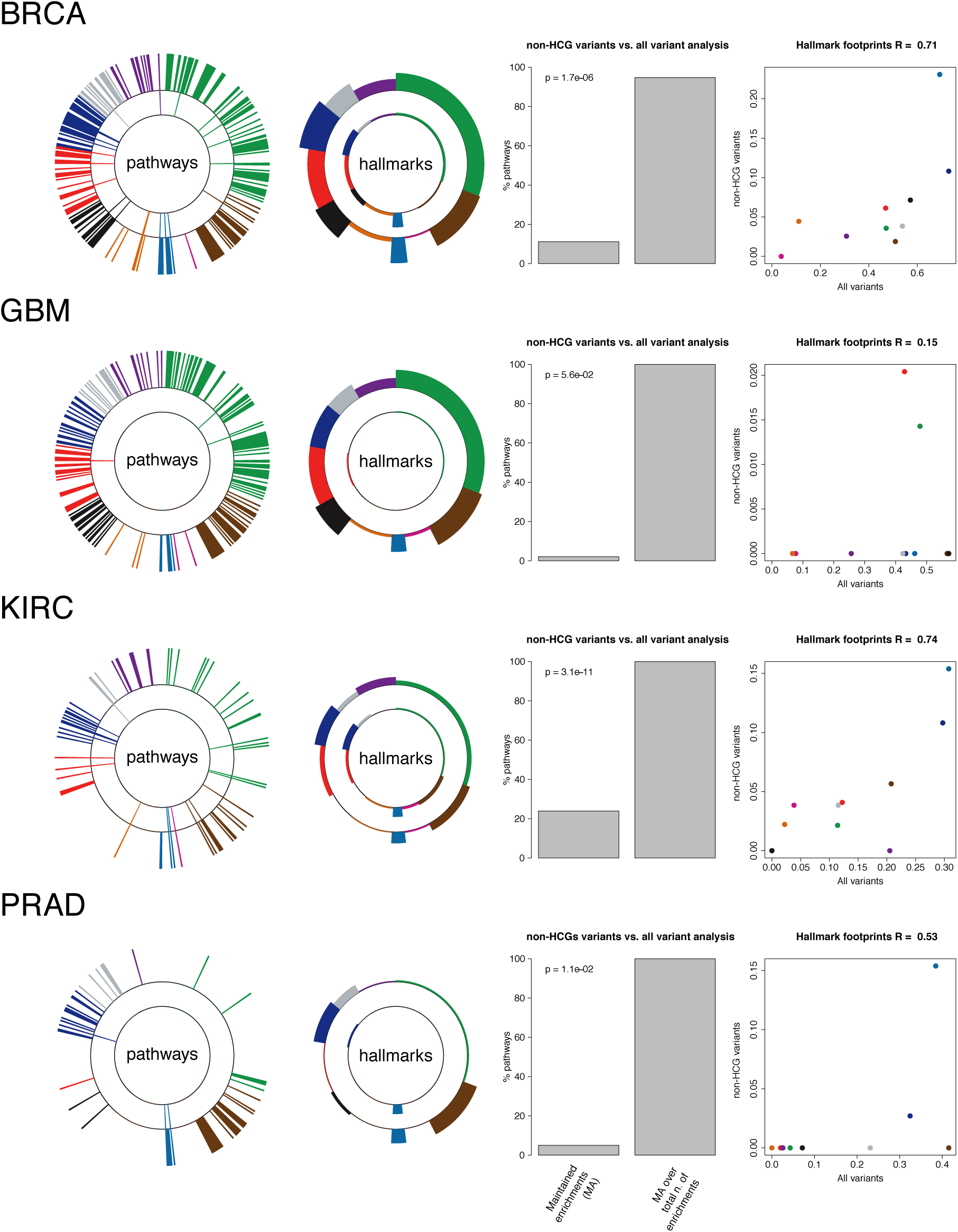
Hallmark heterogeneity signature analysis including and not including known cancer driver genes: In each row, the first circle plot show pathway enrichments at the population level when considering all the somatic variants (bars on the external circle) and when considering only variants not involving known high-confidence cancer driver genes (internal circle); the second circle plot compares the hallmark signatures resulting from SLAPenrich analysis including (bars on the external circle) or excluding (bars on the internal circle) the variants involving known high-confidence cancer genes. The bar plot shows a comparison, in terms of true-positive-rate (TPR) and positive-predictive-value (PPV), of the SLAPenriched pathways recovered in the filtered analysis vs. the complete analysis., The scatter plots on the right show a comparison between the resulting hallmark signatures.

If the hallmark signatures from the original unfiltered analyses are faithful representations of the mutational landscape of the analysed cancer types and the filtered analyses still detect this landscape despite the removal of known drivers, then the filtered analyses might have uncovered novel cancer driver networks composed by infrequently mutated genes. In fact, these new gene modules are typically composed by groups of functionally interconnected and very lowly frequently mutated genes (examples are shown in Figure 6 and the whole bulk of identified network is included in the Supplementary Results).

**Figure 6.**
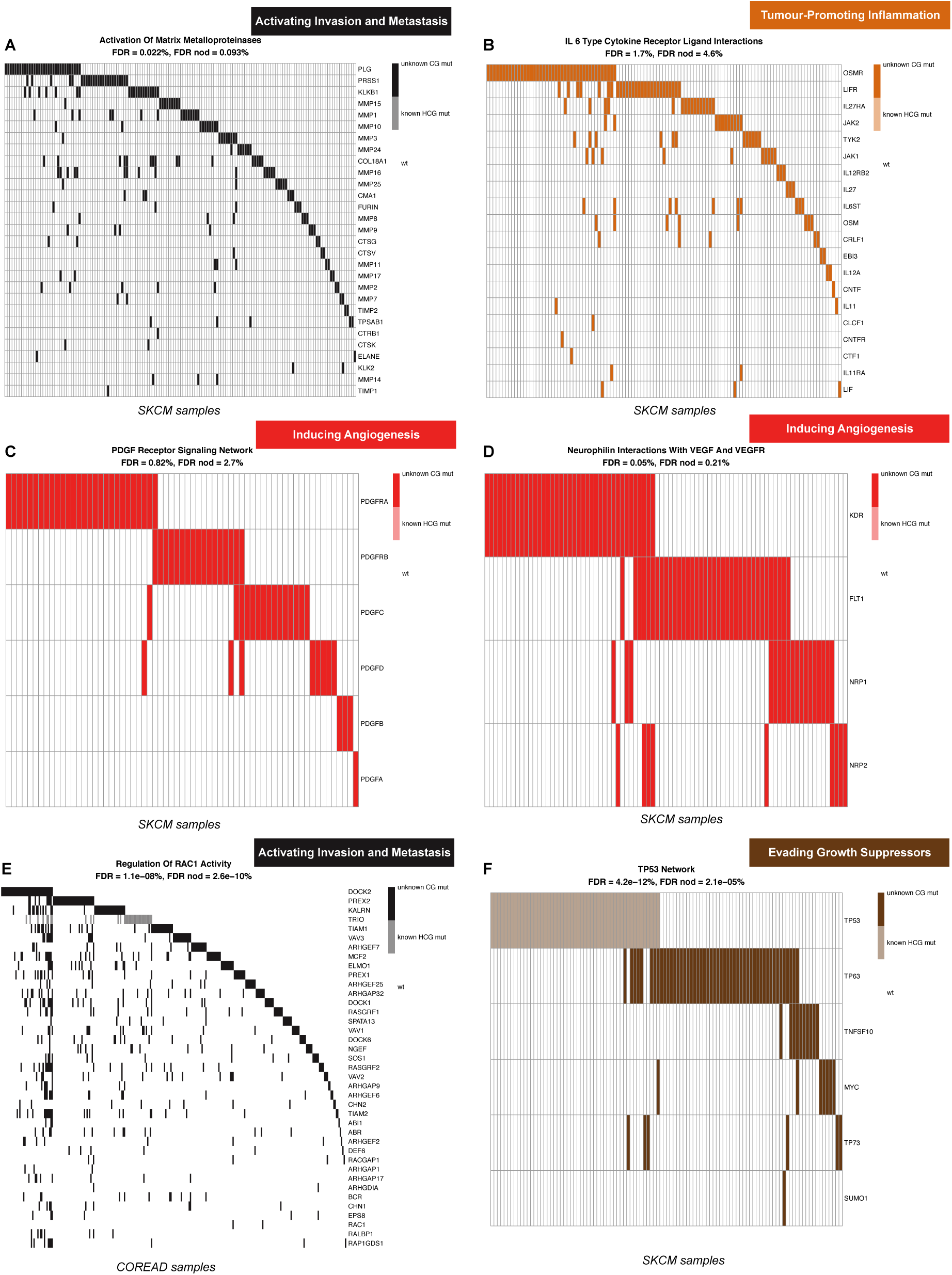
Example of potential novel cancer genes and networks: Picked examples of novel putative cancer driver genes and networks. The first FDR value refers to the unfiltered analysis, whereas the second FDR refers to the filtered one (in which variants involving high confidence and highly frequently mutated cancer driver genes have been removed).

An example is given by the pathway *Activation Of Matrix Metalloproteinases* associated with the *Invasion and metastasis* hallmark and highly enriched in the filtered analyses of COREAD (FDR = 0.002%), SKCM (0.09%) (Figure 6A), LUAD (0.93%), and HNSC (3.1%). The activation of the matrix metalloproteases is an essential event to enable the migration of malignant cells and metastasis in solid tumors^54^. Although this is a hallmark acquired late in the evolution of cancer, according to our analysis this pathway is still detectable as significantly enriched. As a consequence, looking at the somatic mutations of its composing genes (of which only Matrix Metallopeptidase 2 - MMP2 - has been reported as harbouring cancer-driving alterations in LUAD^36^) might reveal novel key components of this pathway leading to metastatic transitions. Interestingly, among these, one of the top frequently mutated genes (across all the 4 mentioned cancer types) is Plasminogen (PLG), whose role in the evolution of migratory and invasive cell phenotype is established^55^. Furthermore, blockade of PLG with monoclonal antibodies, DNA-based vaccination or silencing through small interfering RNAs has been recently proposed to counteract cancer invasion and metastasis^56^. The remaining altered component of this pathway is mostly made of a network of very lowly frequently mutated (and in a highly mutually exclusive manner) other metalloproteinases.

Another similar example is given by the *IL 6 Type Cytokine Receptor Ligand Interactions* pathway significantly enriched in the filtered analysis of SKCM (FDR = 4.6%) and associated with the *Tumour-promoting inflammation hallmark* (Figure 6B). IL-6-type cytokines have been observed to modulate cell growth of several cell types, including melanoma^57^. Increased IL-6 blood levels in melanoma patients correlate with disease progression and lower response to chemotherapy^58^. Importantly, studies proposed OSMR, an IL-6-type cytokine receptor, to play a role in the prevention of melanoma progression^59^, and as a novel potential target in other cancer types^60^. Consistent with these findings, OSMR is the member of this pathway with the largest number of mutations in the SKCM cohort (Figure 6B), complemented by a large number of other lowly frequently mutated genes (most of which are interleukins).

In the context of melanoma, we observed two other highly enriched pathways in the filtered analysis: *PDGF receptor signaling network* (FDR = 2.7%) (Figure 6C) and *Neurophilin Interactions with VEGF And VEGFR* (0.21%)(Figure 6D), both associated with the *Inducing angiogenesis* hallmark. Mutations in all of the components of these two pathways are not common in SKCM and have not been highlighted in any genomic study so far. The first of these two pathway enrichments is characterised by patterns of highly mutually exclusive somatic mutations in Platelet-derived growth factor (PDGF) genes, and their corresponding receptors: a network that has been recently proposed as an autocrine endogenous mechanism involved in melanoma proliferation control^61^.

A final example is given by the enriched pathway *Regulating the activity of RAC1* (associated with the *Activating Invasion and Metastasis* hallmark) in COREAD (Figure 6E). The Ras-Related C3 Botulinum Toxin Substrate 1 (RAC1) gene is a member of the Rho family of GTPases, whose activity is pivotal for cell motility^62^. Previous *in vitro* and *in vivo* studies in prostate cancer demonstrated a marked increase in RAC1 activity in cell migration and invasion, and that RAC1 inhibition immediately stopped these processes^63,64^. However, although the role of RAC1 in enabling metastasis has already been suggested, the mechanisms underlying such aberrant behaviour are poorly understood, and our findings could be used as a starting point for further investigations^65^.

Another interesting case is the high level of mutual exclusivity observed in the mutation patterns involving members of the *TP53 network*, highly enriched in the filtered analysis of SKCM, encompassing TP63, TP73, TNSF10, MYC and SUMD1 (Figure 6F). Whereas alterations in some nodes of this network are known to be an alternative to p53 repression, conferring chemoresistance and poor prognosis^66^, dissecting the functional relations between them is still widely considered a formidable challenge^67^. Our results point out alternative players worthy to be looked at in this network (particularly, among the top frequently altered, TNSF10).

Taken together, these results show the effectiveness of our approach in identifying potential novel cancer driver networks composed by lowly frequently mutated genes.

## Discussion

We have presented a computational pipeline, with a paired statistical framework implemented in an open-source R package (SLAPenrich) to identify genomic alterations in biological pathways, which putatively contribute to the acquisition of the canonical cancer hallmarks. Our statistical framework does not seek pathways whose alterations are enriched at the individual sample level nor at the global level, i.e. considering the union of all the genes altered in at least one sample. Instead, it assumes that an individual mutation involving a given pathway in a given sample might be sufficient to deregulate the activity of that pathway in that sample and it allows enriched pathways to be mutated in a mutually exclusive manner across samples.

With this method we have performed a large-scale comparative analysis of the mutational landscape of different cancer types at the level of cancer hallmarks. Our results represent a first data-driven landmark exploration of the hallmarks of cancer showing that they might be acquired through preferential genomic alterations of heterogeneous sets of pathways across cancer types. This has confirmed the established predominance of certain hallmarks in defined cancer types, and it has highlighted peculiar patterns of altered pathways for several cancer lineages. Finally, by using the identified hallmark signatures as a ground truth signal, we have devised an approach to detect novel cancer driver genes and networks.

A number of possible limitations could hamper the derivation of definitive conclusions from our study, such as the use of only mutations, the possibility that some of the analysed cohorts of patients are representative only of well-defined disease subtypes, the limitation of our knowledge of pathways, and the possibility that pathways that we were not mapped onto cancer hallmarks in our curation could correspond to specific capabilities of cancer cell in certain tumour types. Possible future developments of our method could integrate different *omics*, such as transcriptional data, to better refine the set of functionally impacting variants considered in the analysis. Additionally further refinements could account for structural variants such as small indels and copy number alterations, known to play an important role in cancer.

Our computational pipeline should be of wide usability for the functional characterization of sparse genomic data from heterogeneous populations sharing common traits and subjected to strong selective pressure. As an example of its applicability we have studied large cohorts of publicly available cancer genomes that are publicly available from the TCGA. However, SLAPenrich is of great utility in other scenarios such as for characterizing genomic data generated upon chemical mutagenesis to identify somatic mutations involved in acquired drug resistance, as reported in a recent publication^68^. More generally, it can be used to characterize, at the pathway level, any type of biological dataset that can be modeled as a presence/absence matrix, where genes are on the rows and samples are on the columns.

## Methods

### Formal description of the SLAPenrich statistical framework

Let us consider the list of all the genes *G* = {*g*_1_,*g*_2_,…,*g_n_*}, whose somatic mutational status has been determined across a population of samples *S* = {*s*_1_,*s*_2_,…,*s_m_*}, and a function

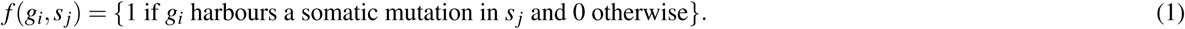

Given the set of all the genes whose products belong to the same pathway *P*, we aim at assessing if there is a statistically significant tendency for the samples in *S* to carry mutations in *P*. Importantly, we do not require the genes in *P* to be significantly enriched in those that are altered in any individual sample nor in the subset of *G* composed by all the genes harbouring at least one somatic mutation in at least one sample. In what follows *P* will be used to indicate the pathway under consideration as well as the corresponding set of genes, interchangeably. We assume that *P* is altered in sample *s_j_* if there is a gene *g_i_* belonging to *G* such that *g_i_* is a member of *P* and *f*(*g_i_*,*s_j_*) = 1, i.e. at least one gene in the pathway *P* is altered in the *j*-th sample (Supplementary Figure S1B). To quantify how likely it is to observe at least one gene belonging to *P* altered in sample *s_j_*, we introduce the variable *X_j_* = |{*g_i_* ∈ *G*: *g_i_* ∈ *P* and *f*(*g_i_*,*s_j_*) = 1}|, accounting for the number of genes in *P* altered in sample *s_j_*. Under the assumption of both a gene-wise and sample-wise statistical independence, the probability of *X_j_* assuming a value greater or equal than 1 is given by:

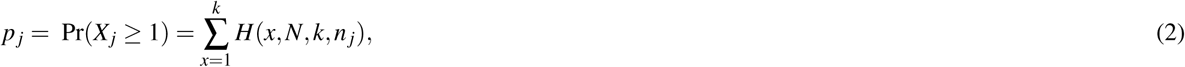

where *N* is the size of the gene background-population, *k* is the number of genes in *P*, *n_j_* is the total number of genes *g_i_* such that *f*(*g_i_*,*s_j_*) = 1, i.e. the total number of genes harbouring an alteration in sample *s_j_*, and *H* is the probability mass function of a hypergeometric distribution:

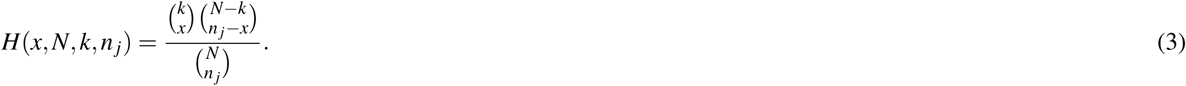

To take into account the impact of the exonic lengths *λ* (*g*) of the genes (*g*) on the estimation of the alteration probability of the pathway they are part of *P*, it is possible to redefine the *p_j_* probabilities (of observing at least one genes in the pathway *P* altered in sample *s_j_*) as follows:

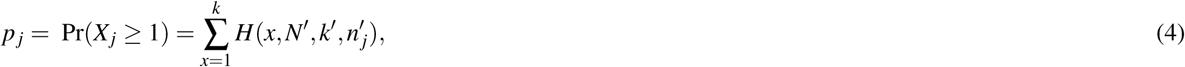

where 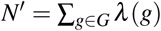 with *G* the gene background-population, i.e. the sum of all the exonic content block lengths of all the genes; 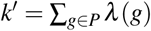 is the sum of the exonic block length of all the genes in the pathway *P*; *n′_j_* is the total number of individual point mutations involving genes belonging to *P* in sample *s_j_*, and *H* is defined as in equation 3, but with parameters *x*,*N′*,*k′*, and *n′_j_*. Similarly, the *p_j_* probabilities can be modeled accounting for the total exonic block lengths of all the genes belonging to *P* and the expected/observed background mutation rate^29^, as follows:

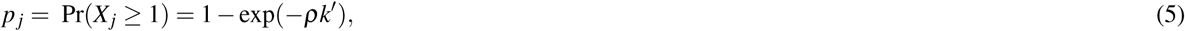

where *k′* is defined as for equation 4 and *ρ* is the background mutation rate, which can be estimated from the input dataset directly or set to established estimated values (such as 10^−6^*/*nucleotide)^29^.

If considering the event “the pathway *P* is altered in sample *s_j_*″ as the outcome of a single test in a set of Bernoulli trials {*j*} (with *j* = 1,…,*M*) (one for each sample in *S*), then each *p_j_* can be interpreted as the success probability of the *j*–*th* trial. By definition, summing these probabilities across all the elements of *S* (all the trials) gives the expected number of successes *E*(*P*), i.e. the expected number of samples harbouring a mutation in at least one gene belonging to *P*:

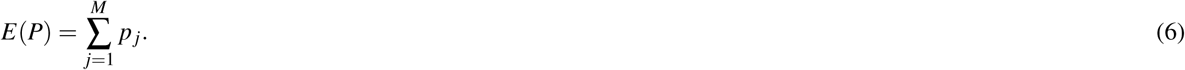

On the other hand, if we consider a function *ϕ* on the domain of the *X* variables, defined as *ϕ*(*X*) = 1 − *δ* (*X*), where *δ* (*X*) is the Dirac delta function (assuming null value for every *X* ≠ 0), i.e. *ϕ*(*X*) = {1 if *X* > 0, and 0 otherwise}, then summing the *ϕ*(*X_i_*) across all the samples in *S*, gives the observed number of samples harbouring a mutation in at least one gene belonging to *P*:

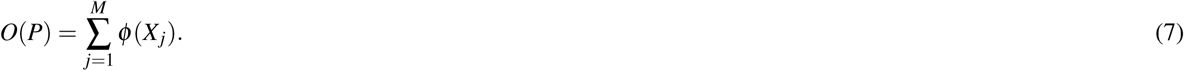

A pathway alteration index, quantifying the deviance of *O*(*P*) from its expectation, and thus how unexpected is to find so many samples with alterations in the pathway *P*, can be then quantified as:

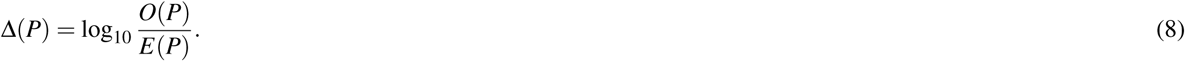

To assess the significance of such deviance, let us note that the probability of the event *O*(*P*) = *y*, with *y* ≤ *M*, i.e. the probability of observing exactly *y* samples harbouring alterations in the pathway *P*, distributes as a Poisson binomial *B* (a discrete probability distribution modeling the sum of a set of {*j*} independent Bernoulli trials where the success probabilities *p_j_* are not identical (with *j* = 1,…,*M*). In our case, the *j*-th Bernoulli trial accounts for the event “the pathway *P* is altered in the sample *s_j_*” and its success probability is given by the {*p_j_*} introduced above (and computed with one amongst 2, 4, or 5). The parameters of such *B* distribution are then the probabilities *π* = {*p_j_*}, and its mean is given by Equation 6. The probability of the event *O*(*P*) = *y* can be then written as

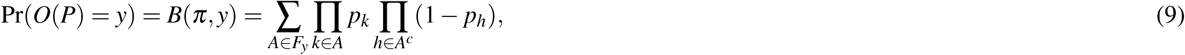

where *F_y_* is the set of all the possible subsets of *y* elements that can be selected from the trial 1,2,…,*M* (for example, if *M* = 3, then *F*_2_ = {{1,2},{1,3},{2,3}}, and *A^c^* is the complement of *A*, i.e. {1,2,…,*M*}\*A*. Therefore a *p-value* can be computed against the null hypothesis that *O*(*P*) is drawn from a Poisson binomial distribution parametrised through the vector of probabilities *π*. Such *p-value* can be derived for an observation *O*(*P*) = *z*, with *z* ≤ *M*, as (Supplementary Figure S1C):

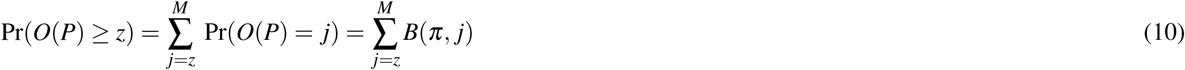

Finally, *p-values* resulting from testing all the pathways in the considered collection are corrected for multiple hypothesis testing with a user-selected method among (in decreasing order of stringency) Bonferroni, Benjamini-Hochberg, and Storey-Tibshirani^69^.

SLAPenrich is implemented as an R package publicly available and fully documented at (https://github.com/saezlab/SLAPenrich/). An overview of the exposed function of this package is also provided in the Additional File 8.

### Pathway gene sets collection and pre-processing

The hallmark signature analyses were performed on a large collection of pathway gene sets from the Pathway Commons data portal (v8, 2016/04)^33^ (http://www.pathwaycommons.org/archives/PC2/v4-201311/). This contained an initial catalogue of 2,794 gene sets (one for each pathway) that were assembled from multiple public available resources, and covering 15,281 unique genes.

From this pathway collection, those gene sets containing less than 4 or more than 1,000 genes, were discarded. Additionally, in order to remove redundancies, those gene sets (*i*) corresponding to the same pathway across different resources or (*ii*) with a large overlap (Jaccard index (*J*) > 0.8, as detailed below) were merged together by intersecting them. The gene sets resulting from this compression were then added to the collection (with a joint pathway label) and those participating in at least one of these merging were discarded. Finally, gene names were updated to their most recent HGCN^70^ approved symbols (this updating procedure is also executed by a dedicate function in of the SLAPenrich package, by default on each genomic datasets prior the analysis). The whole process yielded a final collection of 1,911 pathway gene sets, for a total number of 1,138 genes assigned to at least one gene set.

Given two gene sets *P*_1_ and *P*_2_ the corresponding *J*(*P*_1_,*P*_2_) is defined as:

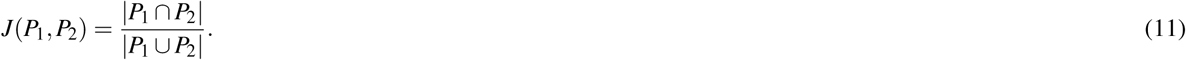

To guarantee results’ comparability with respect to previously published studies, for the case study analysis on the LUAD dataset we downloaded and used the whole collection of KEGG^31^ pathway gene sets from MsigDB^32^, encompassing 189 gene sets for a total number of 5,224 genes included in at least one set.

### Curation of a pathway/hallmark map

We implemented a simple routine (included in the SLAPenrich R package) that assigns to each of the 10 canonical cancer hallmarks a subset of the pathways in a given collection. To this aim this routine searches for determined keywords (typically processes or cellular components) known to be associated with each hallmark in the name of the pathway (such as for example: ‘DNA repair’ or ‘DNA damage’ for the *Genome instability and mutations* hallmark) or for key nodes in the set of included genes or keyword in their name prefix (such as for example ‘TGF’, ‘SMAD’, and ‘IFN’ for *Tumour-promoting inflammation*. The full list of keywords used in this analysis are reported in the Supplementary Table S7. Results of this data curation are reported in the Supplementary Table S8.

### Mutual exclusivity coverage

After correcting the *p-values* yielded by testing all the pathways in a given collection, the enriched pathways can be additionally filtered based on a mutual exclusivity criterion, as a further evidence of positive selection. To this aim, for a given enriched pathway *P*, an exclusive coverage score *C*(*P*) is computed as

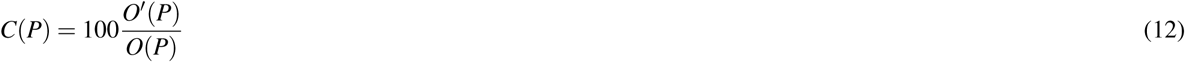

where *O*(*P*) is the number of samples in which at least one gene belonging to the pathway *P* is mutated, and *O*′(*P*) is the number of samples in which exactly one gene belonging to the pathway gene-set *P* is mutated. All the pathways *P* such that *C*(*P*) is at least equal to a chosen threshold value pass this final filter.

### Hallmark heterogeneity signature analysis: genomic datasets and high-confidence cancer genes

Tissue-specific catalogues of genomic variants for 10 different cancer types (breast invasive carcinoma, colon and rectum adenocarcinoma, glioblastoma multiforme, head and neck squamous cell carcinoma, kidney renal clear cell carcinoma, lung adenocarcinoma, ovarian serous cystadenocarcinoma, prostate adenocarcinoma, skin cutaneous melanoma, and thyroid carcinoma) were downloaded from the GDSC1000 data portal described in^34^ (http://www.cancerrxgene.org/gdsc1000/). This resource (available at http://www.cancerrxgene.org/gdsc1000/GDSC1000_WebResources//Data/suppData/TableS2B.xlsx) encompasses variants from sequencing of 6,815 tumor normal sample pairs derived from 48 different sequencing studies^36^ and reannotated using a pipeline consistent with the COSMIC database^71^ (Vagrent: https://zenodo.org/record/16732#.VbeVY2RViko).

Lists of tissue-specific high-confidence cancer genes^36^ were downloaded from the same data portal (http://www.cancerrxgene.org/gdsc1000/GDSC1000_WebResources//Data/suppData/TableS2A.xlsx). These were identified by combining complementary signals of positive selection detected through different state of the art methods^72,73^ and further filtered as described in^34^ (http://www.cell.com/cms/attachment/2062367827/2064170160/mmc1.pdf).

### Hallmark heterogeneity signature analysis: Individual SLAPenrich analysis parameters

All the individual SLAPenrich analyses were performed using the SLAPE.analyse function of the SLAPenrich R package (https://github.com/saezlab/SLAPenrich/) using a Bernoulli model for the individual pathway alteration probabilities across all the samples, the set of all the genes in the dataset under consideration as background population, selecting pathways with at least one gene point mutated in at least 5% of the samples and at least 2 different genes with at least one point mutation across the whole dataset, and and a pathway gene sets collection downloaded from pathway commons^33^, post-processed for redundancy reduction as explained in the previous sections, and embedded in the SLAPenrich package as R data object: PATHCOM HUMAN nr i hu 2016.RData.

A pathway in this collection was considered significantly enriched, and used in the following computation of the hallmark cumulative heterogeneity score, if the SLAPenrichment false discovery rate (FDR) was less than 5% and its mutually exclusive coverage (EC) was greater than 50%.

### Down-sampling analyses

To investigate how differences in sample size might bias the SLAPenrichment results due to a potential tendency for larger datasets to produce larger number of SLAPenriched pathways, down-sampled SLAPenrich analyses were conducted for the 5 datasets with more than 350 samples, i.e. BRCA, COREAD, GBM, HNSC, and LUAD. Particularly, for *n* ∈ {800,400,250} for BRCA and *n* = 250 for the other cancer types, 50 different SLAPenrich analyses were performed on *n* samples randomly selected from the genomic dataset of the cancer type under consideration, with the parameter specifications described in the previous section. The average number of enriched pathways (FDR < 5% and EC > 50%) across the 50 analysis was observed.

### Hallmark signature analysis: signature quantification

For a given cancer type *C* and a given hallmark *H* a cumulative heterogeneity score (CHS) was quantified as the ratio of the pathways associated to *H* in the SLAPenrich analysis of the *C* variants.

The CDS scores for all the 10 hallmark composed the hallmark signature of *C*.

## Acknowledgements

OpenTargets funding to JSR (Projects OpenTargets15 and OpenTargets16). We would like to thank Jorge Buendia, Mathew Garnett, and Annalisa “Lilla” Mupo for a number of insightful discussions, David Tamborero and Nuria Lopez-Bigas for critical feedback on the manuscript.

## Author contributions statement

FI designed the statistical framework underlying SLAPenrich, conceived the hallmark heterogeneity analysis, and designed the other heuristic algorithms, conceived the visualization framework, implemented the R package, and wrote the manuscript; LGA contributed to the implementation of the visualization functions, tested and contributed to implementing the R package, curated data, and contributed to manuscript writing and revising; JB contributed to testing the R package, interpreted results and findings, contributed to manuscript writing and revising; IM contributed to the design of the validation analyses, read and edited the manuscript; DRW contributed to the design of the statistical framework and supervised its mathematical formalization; UM contributed to the interpretation of results; JSR supervised the study and contributed to the manuscript writing and revising.

## Additional information

### Competing financial interests

The authors declare no conflict of interests.

