## Supplementary Materials for "Pathway-based dissection of the genomic heterogeneity of cancer hallmarks’ acquisition with SLAPenrich" (from another version of this preprint: version 4, 2018-03-16)

### Supplementary Methods

#### S1. Introduction

Here we describe in detail the mathematics underpinning SLAPenrich, its implementation, a case study, as well as a comparison with PathScore and PathScan, two related tools.

SLAPenrich is implemented as an R package (available at https://github.com/saezlab/SLAPenrich).
It includes different collections of pathway gene sets from multiple public available sources [1], together with all the data objects needed to run the analysis described in our manuscript. However, it can be also used with any user-defined collection of gene-sets. An overview of the exposed functions of the R package is provided in Additional File 8.
The statistical framework implemented by SLAPenrich is detailed in the Methods section of our manuscript. To visualize enriched pathways SLAPenrich makes use of presence/absence matrices visualised as *binary heatmaps* where columns indicate samples, rows indicate genes harboring at least one somatic mutation in at least one sample of the analyzed dataset, and colors indicate the absence or the presence of somatic mutations (respectively) in a given gene/sample combination. To emphasize mutual exclusivity trends among the row-wise mutation patterns, rows and columns of these heatmaps are sorted with a heuristic method (detailed below) that minimizes the superposition of mutated samples column-wisely, thus the overlaps of the mutation patterns across the rows (an example is provided in Supplementary Figure S2A). To finally summarize the results, an analysis of the enriched-pathway *core-component genes* can be performed. The aim of this final analysis is to visualize in the same heatmap enriched pathways that share a frequently mutated sub-set of genes (the core-component) that is supposed to lead the pathway enrichments, together with a membership matrix specifying to which enriched pathway each core-component gene belongs to (an example is provided in Supplementary Figure S2B, introduced in the next section). This allows filtering out from the results those pathways that are not directly relevant to the disease under consideration, in a supervised way. A final feature

of the package is the identification of pathways that are differentially enriched (thus frequently altered) across two sub-populations of samples of the same input dataset, as detailed in the following sections.

#### S2. Heuristic mutual exclusivity sorting and pathway visualization

The set of somatic mutations of a cancer genomic dataset can be easily modeled as a binary (or Boolean) matrix, whose entries can assume only two possible values, i.e. 0 or 1. In this case, the columns indicate samples, its rows indicate genes (or vice-versa) and a non-zero entry the presence of a somatic mutations in a given gene/sample combination. In a binary matrix, a run is a sequence of consecutive non-zero entries. Reordering rows and columns in a way that the number of runs on the rows and the column-wise marginal totals are minimized is an effective way to highlight patterns of mutual exclusivity among the runs of different rows, i.e. the genes of the considered sub-set. This is an NP-hard problem [2] here referred as *mutual-exclusivity sorting*. In SLAPenrich a heuristic implementation of the mutual-exclusivity sorting is provided in a dedicated R function used by the internal visualization routines, although this function is also available and usable on any user defined binary matrix. Here, for simplicity we will describe an execution of this heuristic applied to a binary matrix summarizing a genomic dataset (with genes on the rows, samples on the columns, and binary entries specifying the status of a gene in a given sample).

In the initial step of the algorithm all the samples and all the genes in the input matrix are declared as *uncovered* and an empty vector is initialized: this is the set of *covered* genes $G$. Then the algorithm proceeds through a series of iterations until the sets of uncovered genes and uncovered samples are both empty. In each of these iterations a *best in class gene* is identified. This is the uncovered gene with the maximal exclusive coverage, which is defined as the number of uncovered samples in which this gene is mutated minus the number of samples in which at least another uncovered gene is mutated. Finally, the identified best in class gene is removed from the set of the uncovered genes, it is attached to $G$, and the set of samples in which it is mutated are removed from the set of the uncovered samples.

After these iterations have been executed, an empty vector of samples $L$ is initialized and all the samples of the dataset are labeled again as uncovered. Then for each of the best in class gene $g$ (in the same order as they appear in $G$) and until there are uncovered samples, the uncovered samples in which $g$ is mutated are sorted according to the exclusive coverage of $g$ across them (in decreasing ordered), they are labeled as covered samples and attached in the resulting order to $L$.

To obtain the final mutual-exclusivity sorting of the initial dataset, the corresponding inputted binary matrix is rearranged by permuting the genes/rows in the same order as they appear in $G$ and the samples/columns in the same order as they appear in $L$.

#### S3. Identification and visualization of enriched pathway core-components

To identify shared core-components across significantly enriched pathways, the set of enriched pathways and their composing genes are modeled as a bipartite network, in which nodes in the first set correspond to enriched pathways and nodes in the second set to genes belonging to at least one of the enriched pathways. Finally a pathway node is connected with an edge to each of its composing gene nodes. The resulting bipartite network is then mined for communities, i.e. groups of densely interconnected nodes, by using a fast community detection algorithm based on a greedy strategy [3]. The resulting communities are finally visualized as independent heatmaps where nodes in the first set (pathways) are on the columns, nodes in the second set (genes) are on the rows and a not-empty cell in position $i,j$ indicates that the $i$-th gene belongs to the $j$-th pathway (an example is provided in Supplementary Figure S2B).

#### S.4 Differential pathway enrichment analysis

Similarly to differential gene expression analysis, the two sub-populations to be contrasted are defined through a contrast matrix. Then individual SLAPenrichment analyses are performed on these two populations, yielding two sets of results. The pathways that are significantly enriched in at least one of the two analyses (according to a user defined false discovery rate (FDR) threshold) are then selected and, for each of them, a differential enrichment score is computed as:

$$\Delta_{A,B}(P) = -\log_{10} FDR_{A(P)} + \log_{10} FDR_{B(P)}$$

where $A$ and $B$ are the two contrasted sub-populations (respectively, positive and negative) and $FDR_{A(P)}$ and $FDR_{B(P)}$ are the two SLAPenrichment FDRs obtained in the two corresponding individual analyses, and $P$ is the pathway under consideration. Graphic routines included in our package allow a pathway level visualization of the inputted alterations across the two contrasted population, on the domain of the differentially enriched pathways as well as heatmaps and barplots of the differential enrichment scores (see an example in Supplementary Figure S2C).

#### S.5 LUAD case study analysis, detailed results and comparison with other methods

To test the ability of SLAPenrich to recover pathways that are known to be associated to a given disease state and different clinico-pathological features, we re-analysed, a published dataset encompassing somatic mutations found in 188 lung adenocarcinoma (LUAD) patients, studied in [4]. To this aim we downloaded annotations of somatic variants and associated clinical information from http://genome.wustl.edu/pub/supplemental/tsp_nature_2008/ (files: supplementary_table_2.tsv and supplementary_table_15.tsv, respectively).

The variants annotations were converted into a genomic event matrix (EM) with altered genes on the rows, patient sample identifiers on the columns, and generic *i,j* entries specifying the number of observed point mutations hosted by the *i*-th gene in the *j*-th patient.

A SLAPenrich analysis on the resulting dataset was performed using the SLAPE.analyse function with default values for all the parameters (including a Bernoulli model [5] for the individual pathway alteration probabilities across all the samples, and the choice of the set of all the altered genes in the dataset as background population), and a pathway gene sets collection from KEGG [6] (embedded in the package as R data object: SLAPE.MSigDB_KEGG_hugoUpdated).

This analysis yielded 48 significantly enriched pathways, at a FDR < 5% and a mutual exclusive coverage (EC) > 50% (Supplementary Table S1). Among these, we found pathways whose deregulation is known to be involved in lung cancer, such as *Tight Junction* (alteration score (AS) = 0.37, EC = 89%)[7] (Supplementary Figure S2A), *Gap Junction* (AS = 0.45, EC = 75%)[8], and several pathways previously found with other computational methods in LUAD (such as PathScan[4], among others [9] - examples include *Focal Adhesion* (AS = 0.06, EC = 84%), *ERBB signaling pathway* (AS = 0.27, EC = 69%), and *Dorsoventral Axis Formation* (AS = 0.42, EC = 55%). Additionally, we found a number of pathways recently proposed as potential targets for lung cancer therapy such as *GNRH signaling pathway* (AS = 0.45, EC = 87\%) [10], *WNT signaling pathway* (AS = 0.29, EC = 74\%) [11], and *VEGF signaling pathway* (AS = 0.33, EC = 80\%) [12].

After applying the same result curation described in [4], i.e. removal of known cancer pathways whose mutation lists are invariably collectively dominated by mutations in TP53, KRAS and EGFR, we found a significant agreement between our results and those obtained with PathScan on the same cohort of cancer patients (and reported in the Supplementary Table 1 of [4]: 26 enriched pathways (FDR < 5\% for both SLAPenrich and PathScan), out of 36 pathways enriched for SLAPenrich and 31 enriched for PathScan (at the same FDR threshold), Fisher's exact test (FET) p-value = $2.10 \times 10^{-14}$ (Supplementary Tables S1 and S2).

Additionally, we observed a significant correlation (R = 0.66, p = 0.0002) between the significance levels of the 26 commonly enriched pathways across the two methods (Supplementary Figure S3A).

A similar, comparison was performed between the output obtained with SLAPenrich and PathScore [13] on the same LUAD dataset. For this analysis a collection of 1,392 canonical pathway signatures from the Molecular Signature Database (MsigDB) [14] was used, as this is the reference collection used by PathScore. We observed a significant overlap (181 pathways, FET p-value = $2.76 \times 10^{-70}$) between the enriched pathways outputted by SLAPenrich (at an FDR < 5\%) and those outputted by PathScore (adjusted p-value < 0.05) (Supplementary Table S3).

As most of the significantly enriched pathways outputted by PathScore have a null p-value it was not possible to check the correlation between the patterns of enrichment significance across the two methods. However when looking at the top enriched pathways across the two analyses (SLAPenrich FDR = 1.76 x $10^{\wedge -12}$ and PathScore adjusted p-value = 0) the results' concordance was even more pronounced (100 overlapping pathways out of the 117 outputted by SLAPenrich and the 176 outputted by PathScore, FET p-value 8.63 x $10^{-83}$), Supplementary Table S3.

To further validate the ability of SLAPenrich in identifying disease relevant pathways and highlight the possible analytical venues allowed by our tool, we considered the clinical information of the samples in the analyzed LUAD dataset. Using this data, we stratified the considered patients based on their smoking status (never-smoker and current-smokers) and their bronchioalveolar carcinoma type (mucinous and non-mucinous), and performed a differential SLAPenrich analysis contrasting the variant profiles of the obtained sub-populations, using the far larger publicly available collection of pathway gene sets from Pathway Commons [1], post-processed for redundancy removal as described in the Methods section of the main text.
Outcomes from the first analysis, comparing never-smoker vs. current-smokers, are reported in Supplementary Table S4 and summarized in Supplementary Figure S2C. In total we found 147 differentially enriched pathways (enriched at FDR < 5% in at least one of the two sub-populations). Ranking these pathways according to their differential pathway enrichments, in decreasing order (Supplementary Figure S2C) highlighted, consistently with previously reported findings, in the current-smokers population a prominent enrichment of alterations in the RAS/RAF/MEK signaling cascade [15], telomerase activity [16], NOXA and PUMA signaling [17]. On the other hand, in the never-smoker population we observed prominent enrichments in EGFR signaling and EGFR-dependent endothelin signaling pathways [18].
When contrasting mucinous versus non-mucinous BAC types (Supplementary Figure S3B and Supplementary Table S5), we observed again correct associations between the mucinous BAC type and pathway alteration enrichments in the RAS/RAF/MEK signaling cascade [19], signaling by leptin [20], PI3K and MTOR signaling pathways [21], and inflammation related pathways such as CXCR3 and GM-CSF mediated signaling. For the non-mucinous BAC type population prominent enrichments were observed in pathways involving EGFR signaling consistently with what reported in [22].
We also performed, with the same common collection of pathways as above, a systematic comparison between SLAPenrich and PathScore [13] on genomic datasets encompassing 4,415 patients across 10 different cancer types from The Cancer Genome Atlas (TCGA). Results confirmed that SLAPenrich and PathScore detect very similar sets of enriched pathways across all the different analysed cancer types (median $-\log_{10}$ (FET p-value) = 119.2, ranging from 29 to 202, Supplementary Figure S4A). We observed a slightly better ability of SLAPenrich in ranking highly pathways that include at least one tissue-specific high-confidence cancer gene (HCG) [23] median HCGs covered by the top 10 enriched pathway for SLAPenrich = 18% against 8% for PathScore; 21% and 14% for the top 20; 33% and 25% for the top 50; 45% and 34% for the 100% (Supplementary Figure S4BC). The median difference of HCGs covered by pathways enriched according to the two methods at the same significance level (5% FDR for SLAPenrich and adjusted $p < 0.05$ for PathScore) favoured PathScore for a 1%.

As explained in the main text, even if performing similarly to SLAPenrich a number of features of PathScan and PathScore make them unsuitable for the hallmark analyses presented here. Finally, SLAPenrich post-processes pathway collections for redundancy reduction: in this way pathways with large overlaps are merged together instead of being tested individually (Figure 3). In summary, while other tools, specially PathScore, are based on similar assumptions and perform comparably, SLAPenrich provides a more flexible environment enabling a wide range of possible large-scale analyses.

#### S.6 SLAPenrich output stability test

For each of the 10 considered cancer types *C* considered in the analysis described in the main text, and whose somatic mutations were included in the dataset *D*, we generated 10 new datasets *Ds* (respectively, *Dp*) simulating a reduction of mutation call sensitivity (respectively, specificity) to 95, 80, 70, and 50%, by removing (respectively, introducing) a corresponding amount of random mutations uniformly distributed across the genome. Therefore, we introduced in each simulated dataset, a ratio of 5, 20, 30 and 50%, respectively, of false positives (FP) and false negatives (FN). To simulate a uniform spread of the introduced noise on the genome, for each simulated dataset the amount of FP/FN to inflate was partitioned across all the genes proportionally to their total exonic block lengths, and across patient samples proportionally to their mutation burdens. This resulted into 80 noise-inflated analyses for each cancer type. Subsequently, we executed SLAPenrich on each of the noise-inflated datasets, for a total amount of 800 different runs. Then we compared the set of pathways outputted by each of these noise inflated SLAPenrich analyses with that outputted when running SLAPenrich on the corresponding original (non-perturbed) dataset *D,* by means of Receiver Operatic Characteristic (ROC) indicators obtaining the results shown in Supplementary Figure S6 and described in the main text.

#### Supplementary Figures

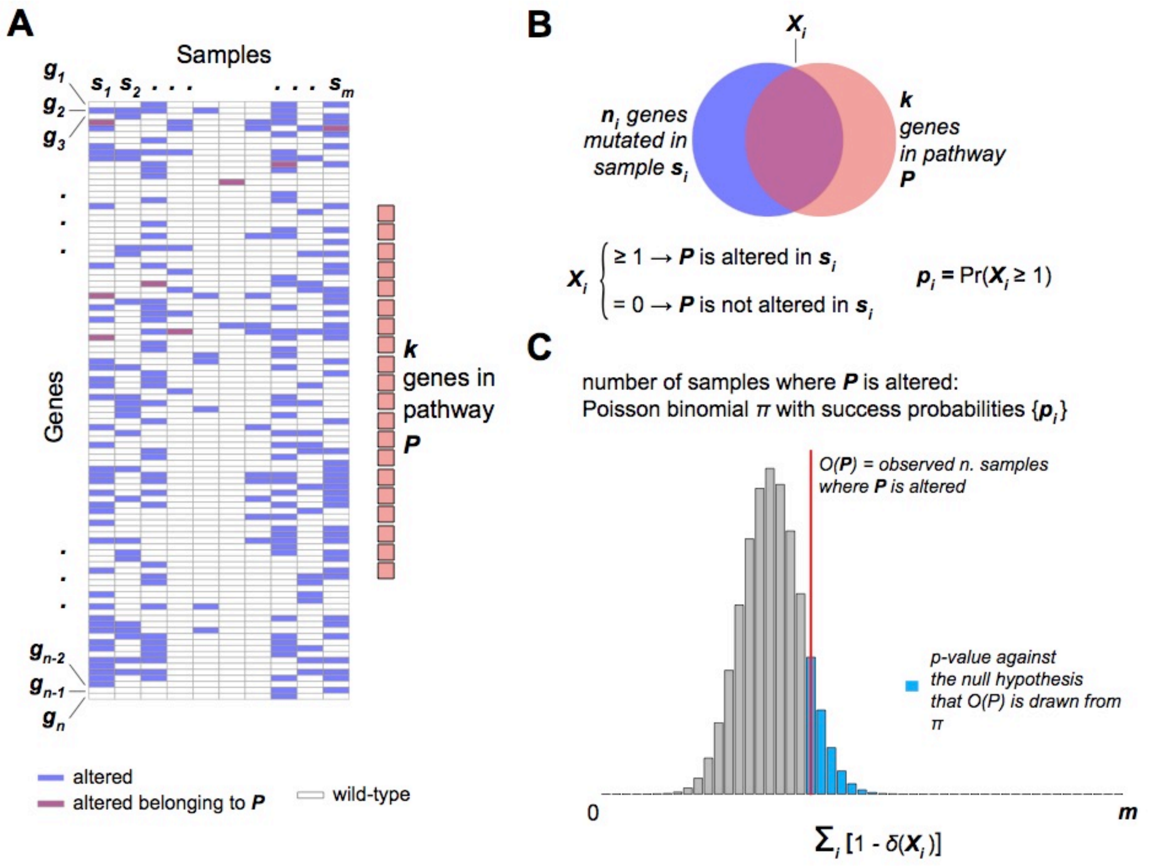

**Supplementary Figure S1: Schematic of the statistical framework underlying SLAPenrich.** The probability $\boldsymbol{p_i}$ of a pathway $\boldsymbol{P}$ being genomically altered in the individual sample $\boldsymbol{s_i}$ of the analyzed dataset is computed. This accounts for the somatic mutation rate of the sample and the sum of the total exonic length blocks of all the $\boldsymbol{k}$ genes in the pathway under consideration. $\boldsymbol{X_i}$ is a random variable quantifying the number of genes belonging to $\boldsymbol{P}$ that are altered in $\boldsymbol{s_i}$, hence the probability of $\boldsymbol{P}$ being altered is $\boldsymbol{p_i} = \Pr(\boldsymbol{X_i} \geq \boldsymbol{1})$. (B) A pathway $\boldsymbol{P}$ is assumed to be genomically altered in the sample $\boldsymbol{s_i}$ if at least one of its $\boldsymbol{k}$ genes is mutated in $\boldsymbol{s_i}$. (C) The number of samples for which $\boldsymbol{X_i}$ is greater than 0 is modeled through a Poisson binomial distribution $\boldsymbol{\pi}$. Here the success probabilities are the likelihoods computed in A. $\boldsymbol{\delta}$ is the Dirac delta function, equal to 1 only when its argument is equal to 0. A p-value against the null hypothesis that there is no association between $\boldsymbol{P}$ and the genomic somatic alterations in the analyzed dataset is computed as the complementary cumulative distribution function of $\boldsymbol{p_i}$ evaluated at $\boldsymbol{O}(\boldsymbol{P})$, which is the observed number of samples where $\boldsymbol{P}$ is genomically altered.

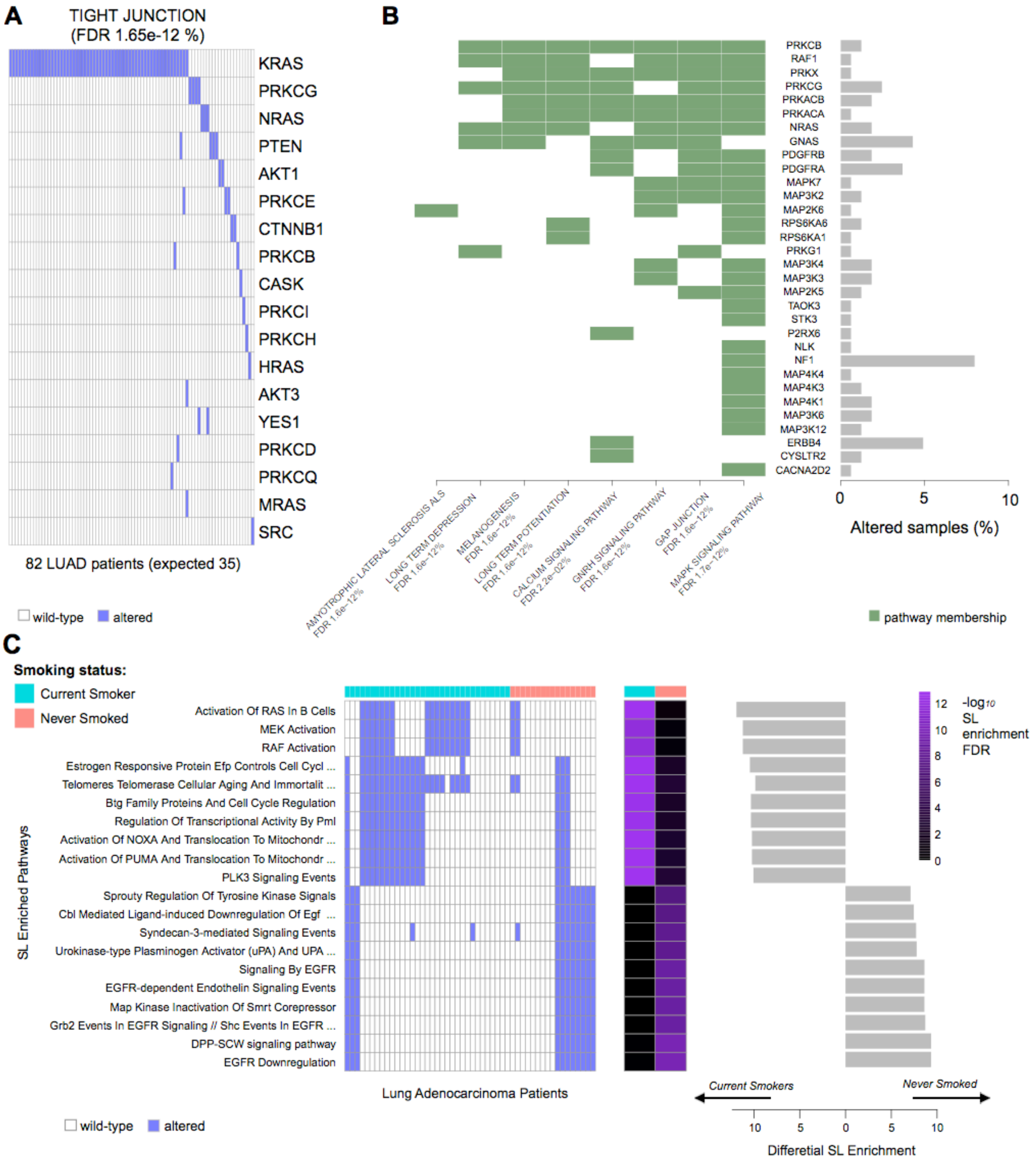

**Supplementary Figure S2: Visualization of enriched pathways, core-components and differential pathway enrichment analysis for the LUAD case study.** (A) Heatmap summarizing the status of the genes belonging to a pathway enriched at the population level in the case study of the lung adenocarcinoma dataset. Genes and patient samples (respectively on rows and columns) have been permuted with a dedicated function in order to highlight mutual exclusivity trends in the observed somatic alterations. (B) Heatmap showing a sub-set of genes (on the rows) shared by multiple significantly enriched pathways (on the columns), together with a bar plot diagram (on the right) showing the percentages of patient samples where each gene is altered. SLAPenrich automatically generates these figures. (C) Visual output of the differential enrichment analysis function using the case study lung adenocarcinoma dataset in input, and stratifying patients based on their smoking status. The heatmap on the left shows the alteration status of the top/bottom 10 most positively/negatively differentially enriched pathways between the groups of smokers vs non-smokers (on the column); the heatmap in the centre shows enrichment significance of individual pathways in the two sub-populations, and the barplot shows corresponding differential sample level enrichment scores.

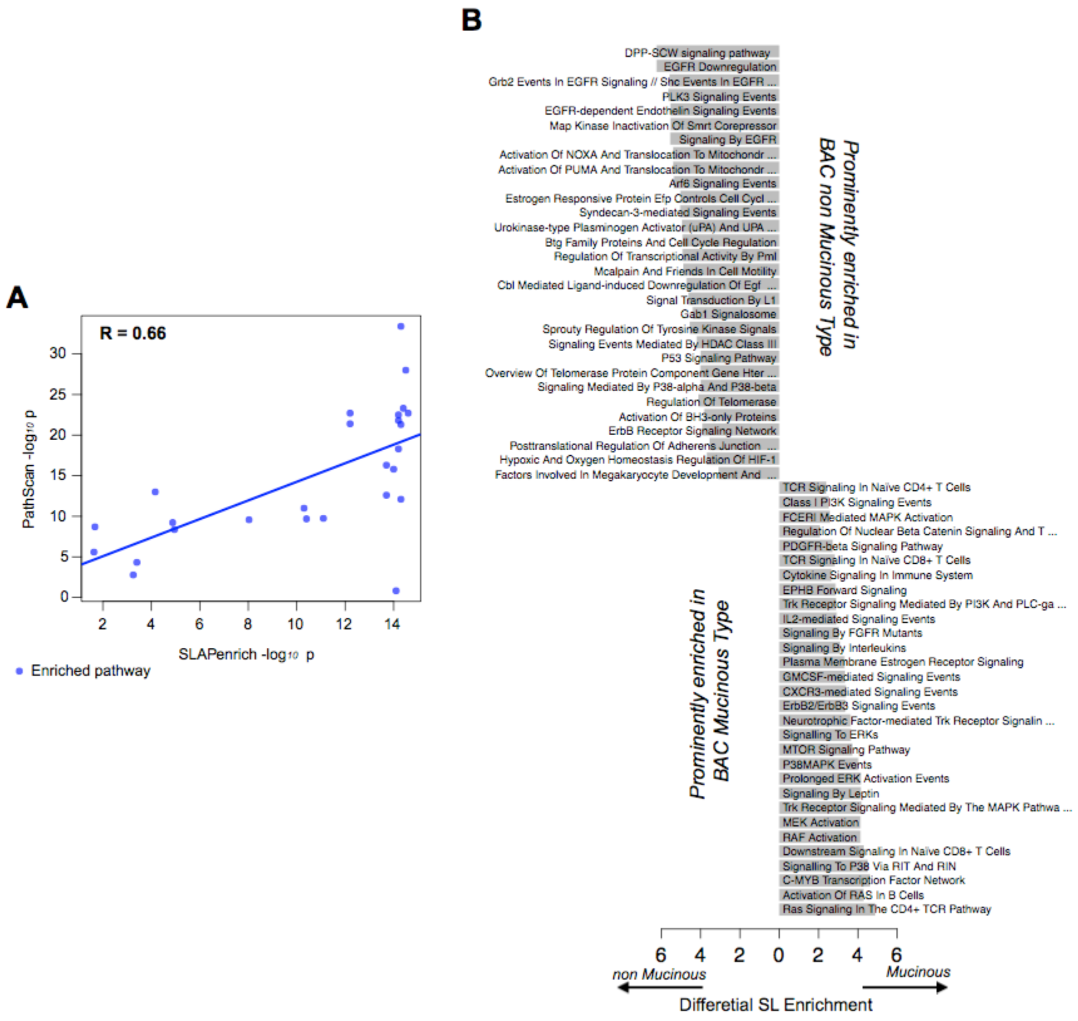

**Supplementary Figure S3: Differential pathway enrichment analysis results: non-mucinous vs mucinous bronchioloalveolar LUAD patients and comparison with PathScan.** (A) Comparison between the significance levels of the enriched pathways (blue dots) identified with both SLAPenrich (x-axis) and PathScan (y-axis) on the LUAD dataset; Results from a differential SLAPenrich analysis obtained contrasting two sub-pulations of LUAD patients based on their bronchioloalveolar type (non-mucinous vs. mucinous).

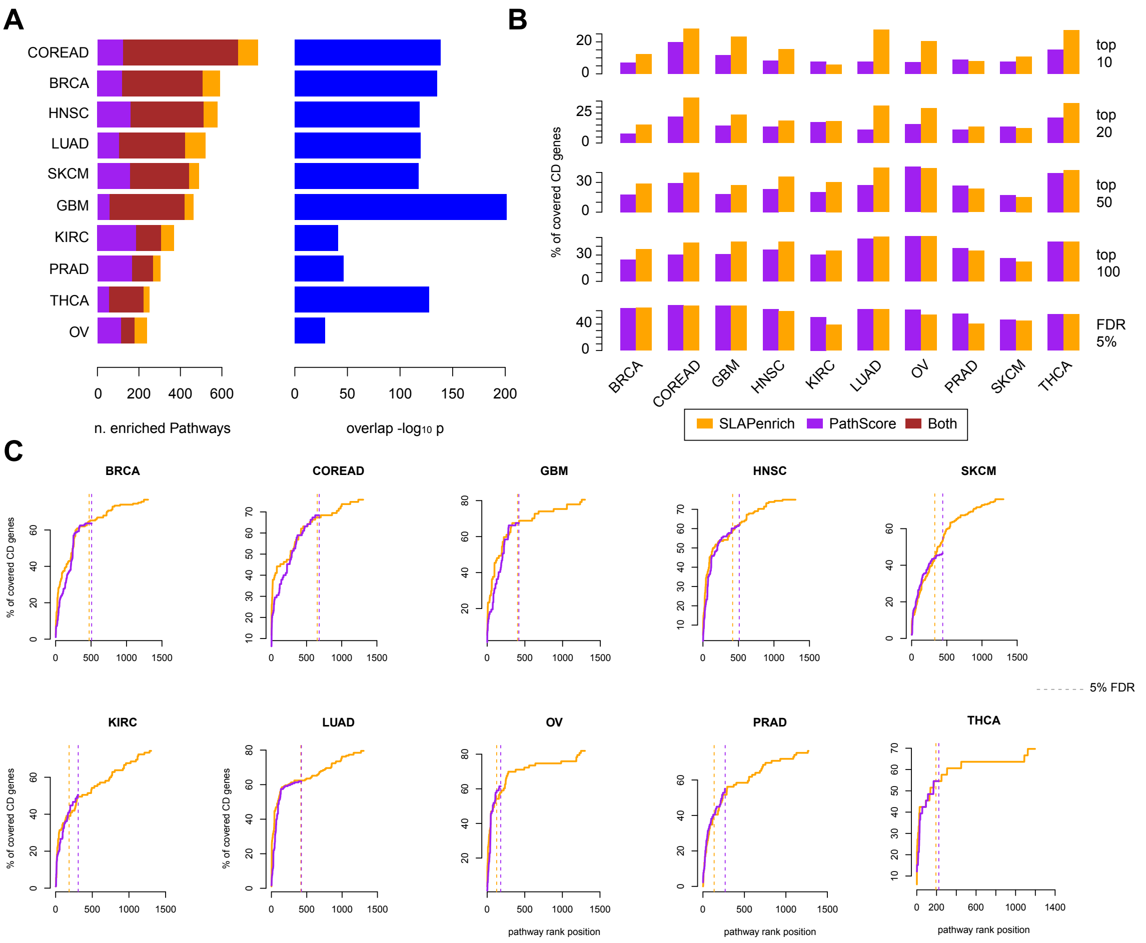

**Supplementary Figure S4: Pathway enrichment significance correlation between SLAPenrich and PathScore.** (A) Number of enriched pathways detected by SLAPenrich, PathScore and both methods across cancer types (left barplot), and overlap significance (right barplot); (B) Percentages of tissue specific high-confidence cancer driver genes included in the top 10, 20, 50 and 100 enriched pathways according to SLAPenrich and PathScore across cancer types (first 4 barplots), and in whole set of statistically significantly enriched pathway (FDR < 5% for SLAPenrich and adjusted p-value < 0.05 for PathScore; (C) Percentages of tissue specific high-confidence cancer driver genes included in the top *k* enriched pathways according to SLAPenrich and PathScore. For SLAPenrich, all the possible k values are considered; PathScore does not output results for all the tested pathways but only for the significantly enriched one, therefore in this case *k* ranges from 1 to the least significantly enriched pathway.

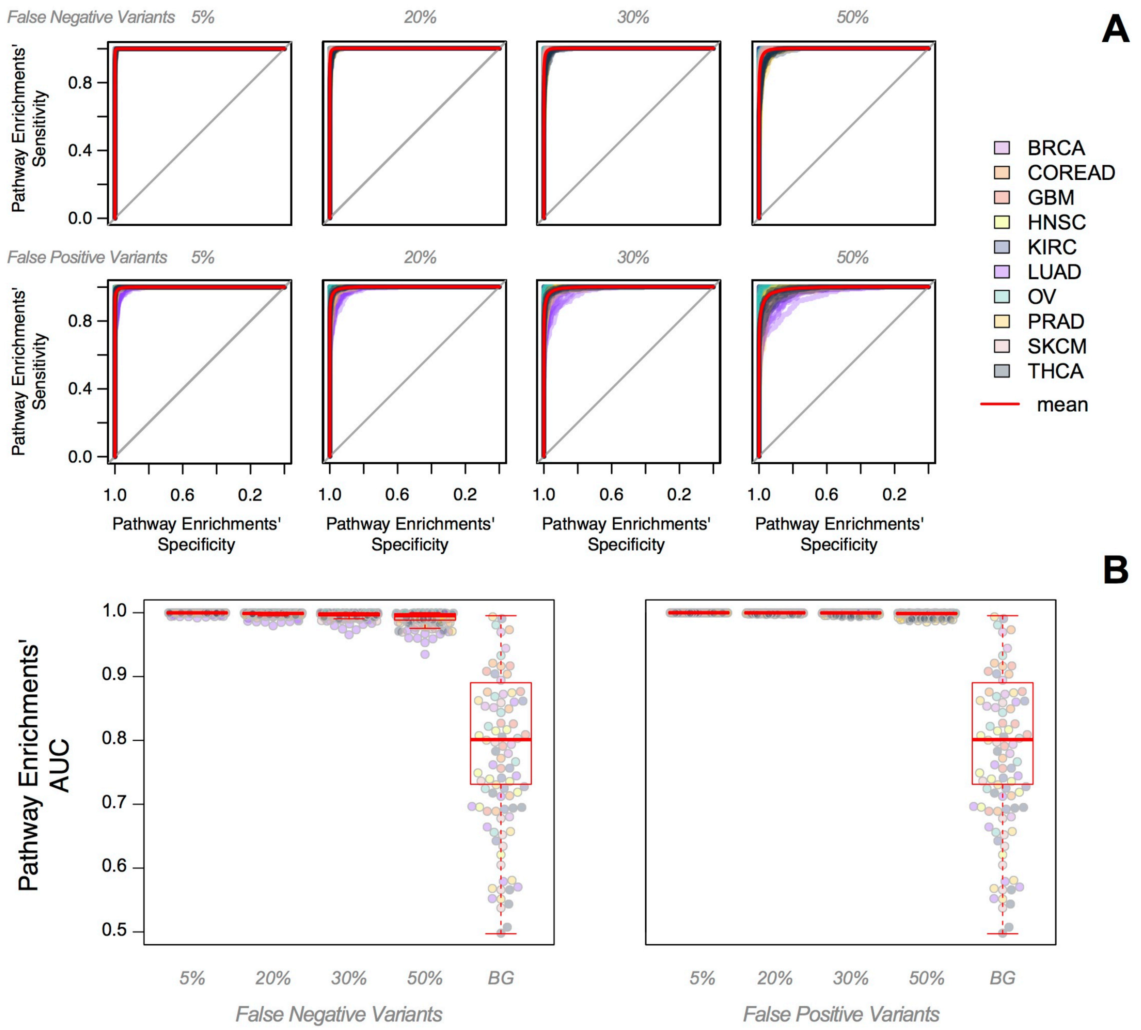

**Supplementary Figure S5: SLAPenrich stability with respect to random noise.** (A) Receiver Operating Characteristic (ROC) curves obtained by comparing the output of SLAPenrich executions on noise inflated versions of 10 genomics datasets (one per analysed cancer type) with that resulting from executing SLAPenrich on the corresponding original dataset. Plots on the first row show results from considering increasing ratios of inflated false negative variants, whereas those on the second row show results obtained when considering increasing ratios of inflated false positive variants. (B) Areas under the curves (AUCs) showed in the plots in A, grouped according to the ratio of inflated false negative variants (left plot) and false positive variants (right plot). In each plot a fifth background (BG) group, showing average AUCs from comparing results from each SLAPenrich analysis (on non noise inflated datasets) with all the others has been included for reference.

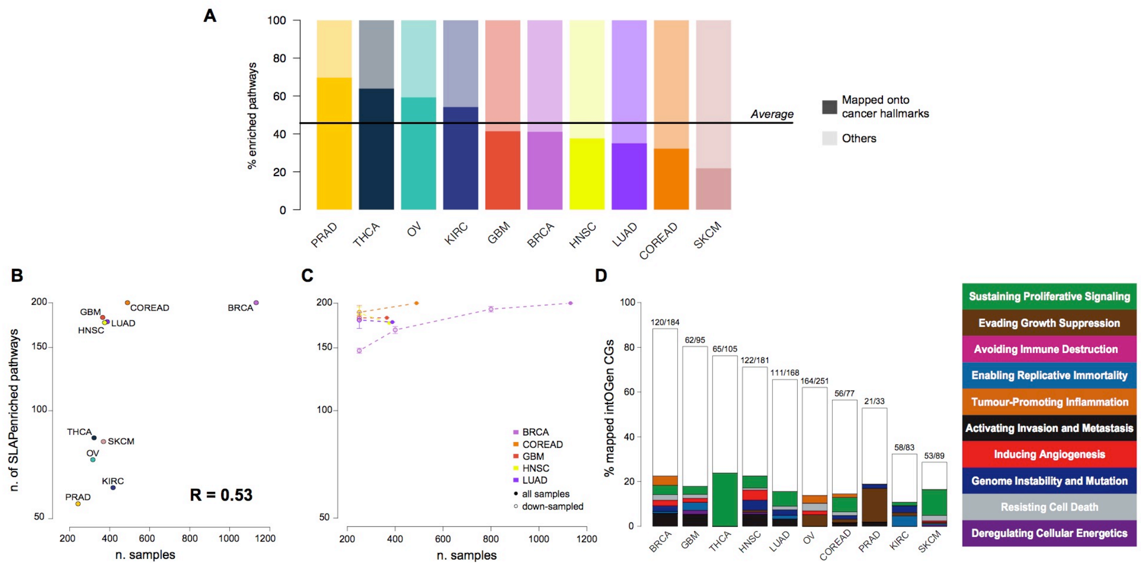

**Supplementary Figure S6: Enriched pathways versus sample size, downsampled analyses, and covered known cancer genes.** (A) Ratios of significantly enriched pathways that are mapped/not-mapped onto canonical cancer hallmarks across cancer types. (B) Number of significantly enriched pathway at the population versus the number of samples available in the analysed cohorts, across cancer type. (C) Number of significantly enriched pathway at the population level across 5 different cancer types (with more than 350 samples), indicated by different colors, and down-sampled trials. In each of this trials, for each cancer type and 50 different iterations, a set of $n$ samples is randomly selected and a SLAPenrich analysis is performed on this sub-set of data. Average number of SLAPenriched pathway (and standard deviations) are reported. n = 800, 400, and 250 for BRCA and n = 250 for the other four cancer type. For four of the tested tissues there is no tendency for increased number of samples to produce more SLAPenriched pathways. A mild dependency trend is observable for BRCA only, with a continuously increasing average number of enriched pathways as a function of sample size up to 800 samples, that plateaus above this size, with a very similar number of enriched pathways when analysing 1,132 samples or across 50 analysis on 800 pathways. (D) Each bar quantifies the ratio of high-confidence cancer genes contained in at least one pathway enriched at the population level (covered pathways), across cancer types. Different contained colored bars indicate the ratio of the genes included in covered pathways associated to different hallmarks, one colored bar per hallmark. The white bar at the top indicates the ratio of genes included in covered pathways associated to multiple hallmarks.

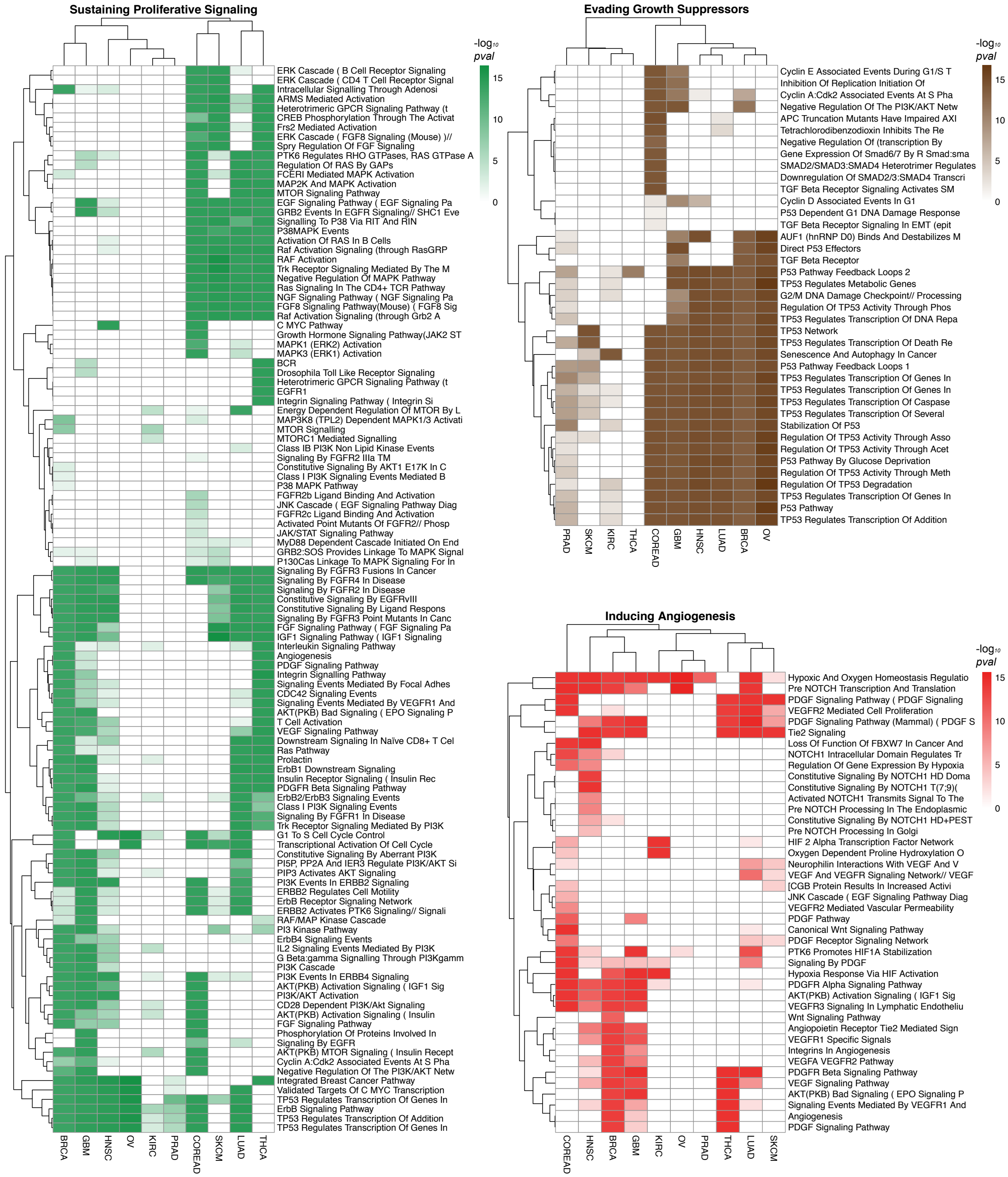

**Supplementary Figure S7: Hallmark heterogeneity across cancer types.** Heatmaps showing pathways enrichments at the population level across cancer types for individual hallmarks. Color intensities correspond to the enrichment significance. Cancer types and pathways are clustered using a correlation metric. See also Figure 4.

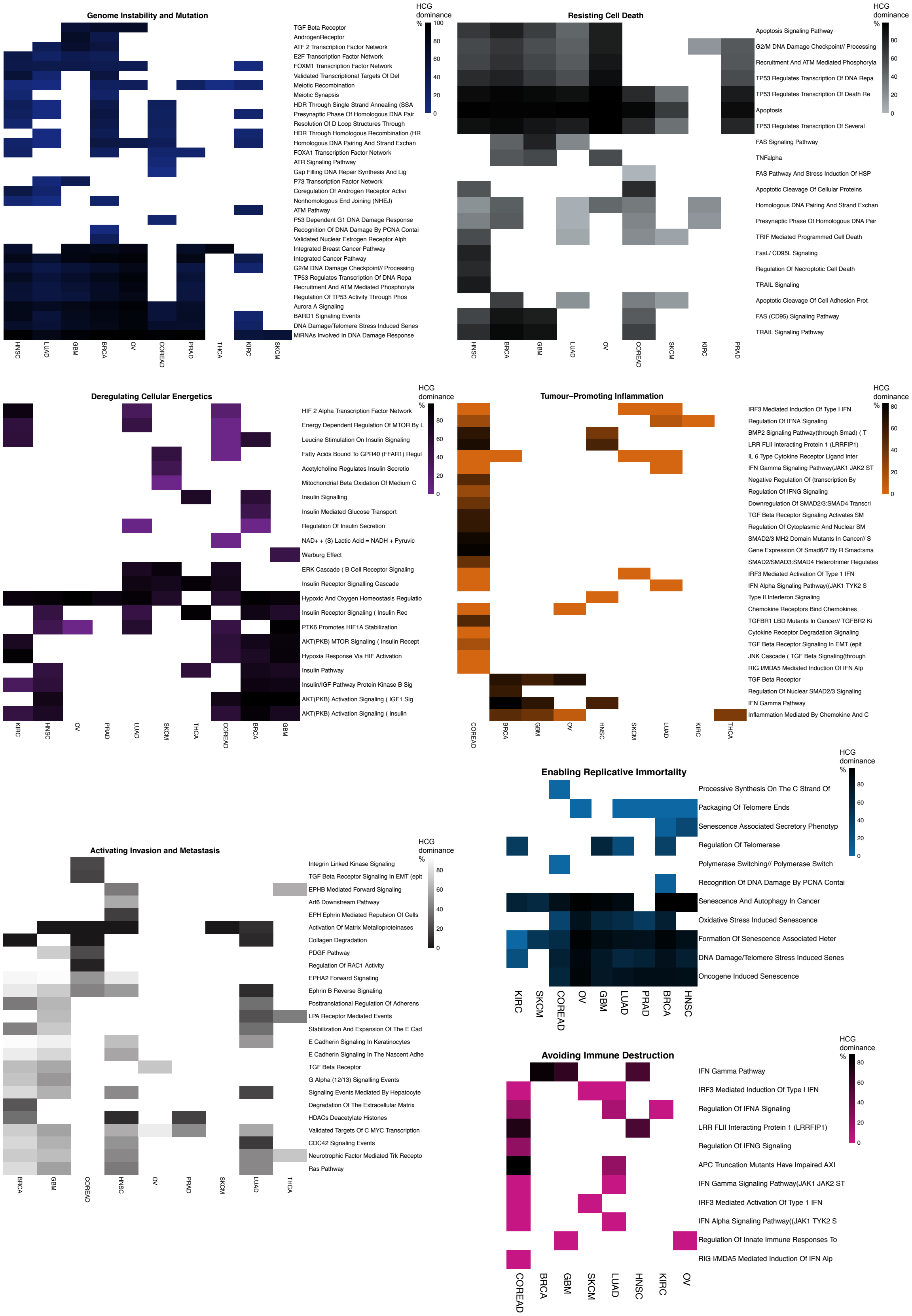
Genome Instability and Mutation
HCG dominance %
100
80
60
40
20
0
TGF Beta Receptor
AndrogenReceptor
ATF 2 Transcription Factor Network
E2F Transcription Factor Network
FOXM1 Transcription Factor Network
Validated Transcriptional Targets Of Del
Meiotic Recombination
Meiotic Synapsis
HDR Through Single Strand Annealing (SSA
Presynaptic Phase Of Homologous DNA Pair
Resolution Of D Loop Structures Through
HDR Through Homologous Recombination (HR
Homologous DNA Pairing And Strand Exchan
FOXA1 Transcription Factor Network
ATR Signaling Pathway
Gap Filling DNA Repair Synthesis And Lig
P73 Transcription Factor Network
Coregulation Of Androgen Receptor Activi
Nonhomologous End Joining (NHEJ)
ATM Pathway
P53 Dependent G1 DNA Damage Response
Recognition Of DNA Damage By PCNA Contai
Validated Nuclear Estrogen Receptor Alph
Integrated Breast Cancer Pathway
Integrated Cancer Pathway
G2/M DNA Damage Checkpoint// Processing
TP53 Regulates Transcription Of DNA Repa
Recruitment And ATM Mediated Phosphoryla
Regulation Of TP53 Activity Through Phos
Aurora A Signaling
BARD1 Signaling Events
DNA Damage/Telomere Stress Induced Senes
MiRNAs Involved In DNA Damage Response
HNSC
LUAD
GBM
BRCA
OV
COREAD
PRAD
THCA
KIRC
SKCM
Resisting Cell Death
HCG dominance %
80
60
40
20
0
Apoptosis Signaling Pathway
G2/M DNA Damage Checkpoint// Processing
Recruitment And ATM Mediated Phosphoryla
TP53 Regulates Transcription Of DNA Repa
TP53 Regulates Transcription Of Death Re
Apoptosis
TP53 Regulates Transcription Of Several
FAS Signaling Pathway
TNFalpha
FAS Pathway And Stress Induction Of HSP
Apoptotic Cleavage Of Cellular Proteins
Homologous DNA Pairing And Strand Exchan
Presynaptic Phase Of Homologous DNA Pair
TRIF Mediated Programmed Cell Death
FasL/ CD95L Signaling
Regulation Of Necroptotic Cell Death
TRAIL Signaling
Apoptotic Cleavage Of Cell Adhesion Prot
FAS (CD95) Signaling Pathway
TRAIL Signaling Pathway
HNSC
BRCA
GBM
LUAD
OV
COREAD
SKCM
KIRC
PRAD
Deregulating Cellular Energetics
HCG dominance %
80
60
40
20
0
HIF 2 Alpha Transcription Factor Network
Energy Dependent Regulation Of MTOR By L
Leucine Stimulation On Insulin Signaling
Fatty Acids Bound To GPR40 (FFAR1) Regul
Acetylcholine Regulates Insulin Secretio
Mitochondrial Beta Oxidation Of Medium C
Insulin Signalling
Insulin Mediated Glucose Transport
Regulation Of Insulin Secretion
NAD+ + (S) Lactic Acid = NADH + Pyruvic
Warburg Effect
ERK Cascade ( B Cell Receptor Signaling
Insulin Receptor Signalling Cascade
Hypoxic And Oxygen Homeostasis Regulatio
Insulin Receptor Signaling ( Insulin Rec
PTK6 Promotes HIF1A Stabilization
AKT(PKB) MTOR Signaling ( Insulin Recept
Hypoxia Response Via HIF Activation
Insulin Pathway
Insulin/IGF Pathway Protein Kinase B Sig
AKT(PKB) Activation Signaling ( IGF1 Sig
AKT(PKB) Activation Signaling ( Insulin
KIRC
HNSC
OV
PRAD
LUAD
SKCM
THCA
COREAD
BRCA
GBM
Tumour–Promoting Inflammation
HCG dominance %
80
60
40
20
0
IRF3 Mediated Induction Of Type I IFN
Regulation Of IFNA Signaling
BMP2 Signaling Pathway(through Smad) ( T
LRR FLII Interacting Protein 1 (LRRFIP1)
IL 6 Type Cytokine Receptor Ligand Inter
IFN Gamma Signaling Pathway(JAK1 JAK2 ST
Negative Regulation Of (transcription By
Regulation Of IFNG Signaling
Downregulation Of SMAD2/3:SMAD4 Transcri
TGF Beta Receptor Signaling Activates SM
Regulation Of Cytoplasmic And Nuclear SM
SMAD2/3 MH2 Domain Mutants In Cancer// S
Gene Expression Of Smad6/7 By R Smad:sma
SMAD2/SMAD3:SMAD4 Heterotrimer Regulates
IRF3 Mediated Activation Of Type 1 IFN
IFN Alpha Signaling Pathway((JAK1 TYK2 S
Type II Interferon Signaling
Chemokine Receptors Bind Chemokines
TGFBR1 LBD Mutants In Cancer// TGFBR2 Ki
Cytokine Receptor Degradation Signaling
TGF Beta Receptor Signaling In EMT (epit
JNK Cascade ( TGF Beta Signaling(through
RIG I/MDA5 Mediated Induction Of IFN Alp
TGF Beta Receptor
Regulation Of Nuclear SMAD2/3 Signaling
IFN Gamma Pathway
Inflammation Mediated By Chemokine And C
COREAD
BRCA
GBM
OV
HNSC
SKCM
LUAD
KIRC
THCA
Enabling Replicative Immortality
HCG dominance %
80
60
40
20
0
Processive Synthesis On The C Strand Of
Packaging Of Telomere Ends
Senescence Associated Secretory Phenotyp
Regulation Of Telomerase
Polymerase Switching// Polymerase Switch
Recognition Of DNA Damage By PCNA Contai
Senescence And Autophagy In Cancer
Oxidative Stress Induced Senescence
Formation Of Senescence Associated Heter
DNA Damage/Telomere Stress Induced Senes
Oncogene Induced Senescence
KIRC
SKCM
COREAD
OV
GBM
LUAD
PRAD
BRCA
HNSC
Activating Invasion and Metastasis
HCG dominance %
80
60
40
20
0
Integrin Linked Kinase Signaling
TGF Beta Receptor Signaling In EMT (epit
EPHB Mediated Forward Signaling
Arf6 Downstream Pathway
EPH Ephrin Mediated Repulsion Of Cells
Activation Of Matrix Metalloproteinases
Collagen Degradation
PDGF Pathway
Regulation Of RAC1 Activity
EPHA2 Forward Signaling
Ephrin B Reverse Signaling
Posttranslational Regulation Of Adherens
LPA Receptor Mediated Events
Stabilization And Expansion Of The E Cad
E Cadherin Signaling In Keratinocytes
E Cadherin Signaling In The Nascent Adhe
TGF Beta Receptor
G Alpha (12/13) Signalling Events
Signaling Events Mediated By Hepatocyte
Degradation Of The Extracellular Matrix
HDACs Deacetylate Histones
Validated Targets Of C MYC Transcription
CDC42 Signaling Events
Neurotrophic Factor Mediated Trk Recepto
Ras Pathway
BRCA
GBM
COREAD
HNSC
OV
PRAD
SKCM
LUAD
THCA
Avoiding Immune Destruction
HCG dominance %
80
60
40
20
0
IFN Gamma Pathway
IRF3 Mediated Induction Of Type I IFN
Regulation Of IFNA Signaling
LRR FLII Interacting Protein 1 (LRRFIP1)
Regulation Of IFNG Signaling
APC Truncation Mutants Have Impaired AXI
IFN Gamma Signaling Pathway(JAK1 JAK2 ST
IRF3 Mediated Activation Of Type 1 IFN
IFN Alpha Signaling Pathway((JAK1 TYK2 S
Regulation Of Innate Immune Responses To
RIG I/MDA5 Mediated Induction Of IFN Alp
COREAD
BRCA
GBM
SKCM
LUAD
HNSC
KIRC
OV

**Supplementary Figures S8: Impact of known cancer genes' mutations on the results.** Heatmaps showing, for each enriched-pathway/cancer-type, the ratio between the number samples harbouring mutations in known cancer genes belonging to the pathway under consideration and the total number of samples harbouring mutations in any gene belonging to the pathway under consideration. See also Figure 6.

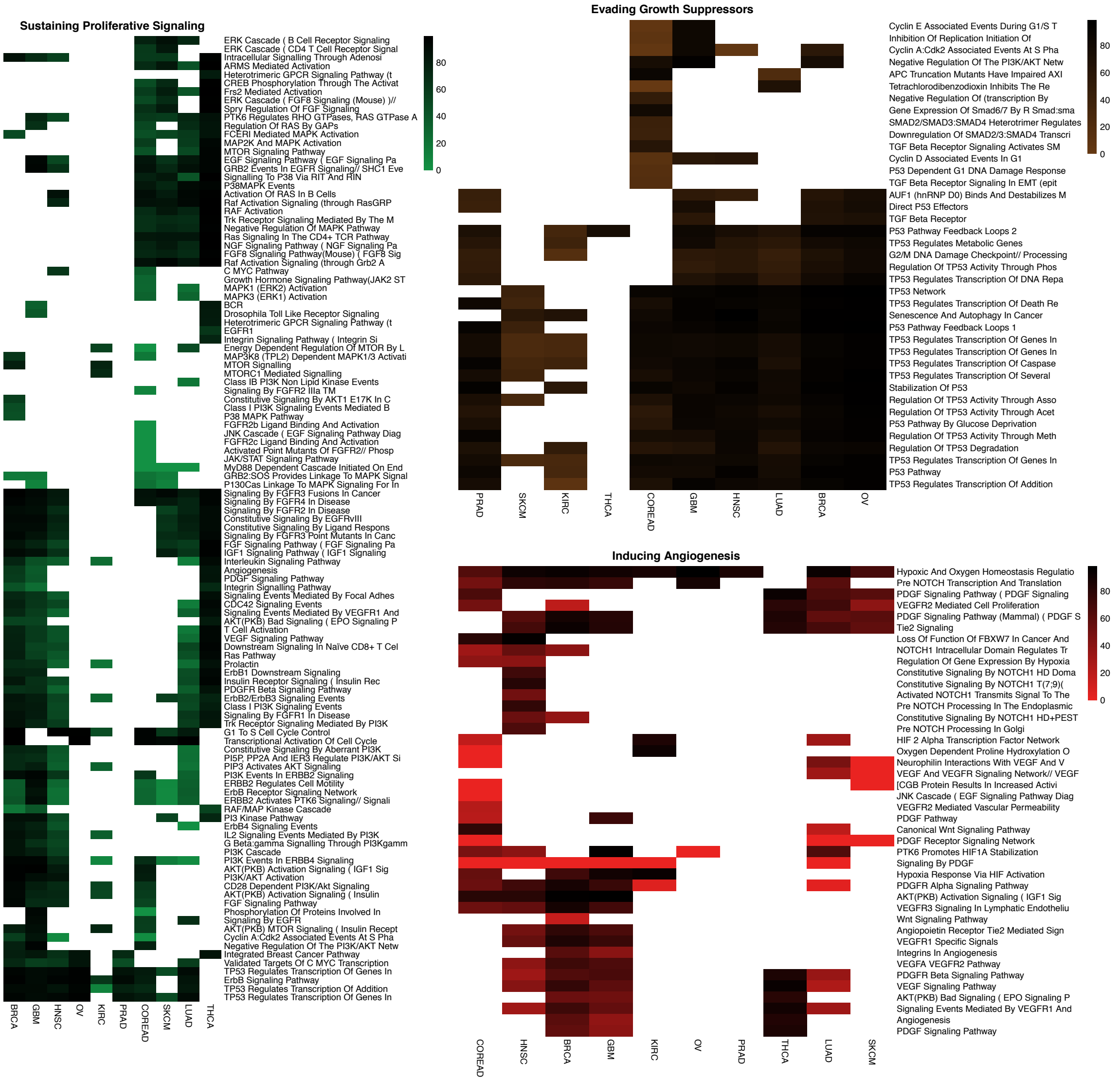

**Supplementary Figures S9: Impact of known cancer genes' mutations on the results.** Heatmaps showing, for each enriched-pathway/cancer-type, the ratio between the number samples harbouring mutations in known cancer genes belonging to the pathway under consideration and the total number of samples harbouring mutations in any gene belonging to the pathway under consideration. See also Figure 6.

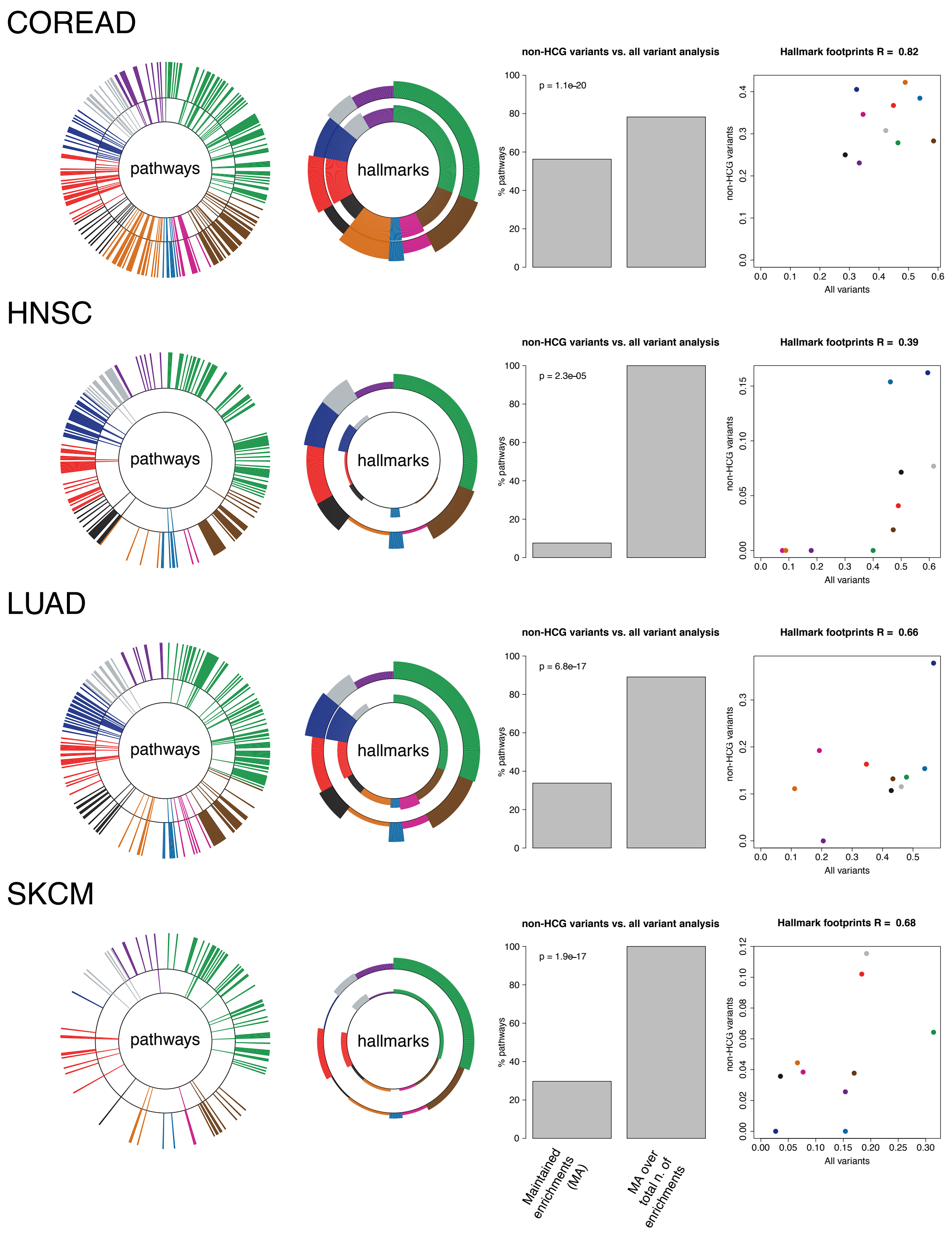

**Supplementary Figures S10: Hallmark signature analysis to discover new cancer driver networks.** In each row, first circle plots shows pathway enrichments at the population levels when considering all the somatic variants (bars on the external circle) and when considering only variants not involving known high-confidence cancer driver genes; second circle plot shows similarly a comparison between the hallmark signatures resulting from SLAPenrich analysis including (bars on the external circle) or excluding (bars on the internal circle) the variants involving known high-confidence cancer genes. The bar plot shows a comparison, in terms of true-positive-rate (TPR) and positive-predicted-value (PPV), of the SLAPenriched pathways across the two analysis and, finally, the scatter plots on the right shows a comparison between the resulting hallmark signatures.

#### List of Supplementary Tables

**Supplementary Table S1: KEGG pathways enriched in the LUAD case study dataset**

**Supplementary Table S2: SLAPenrich/PathScan results' comparison**

**Supplementary Table S3: SLAPenrich/PathScore results' comparison**

**Supplementary Table S4: Differential enrichment analysis comparing LUAD Smokers vs. Non-Smokers patients (Results)**

**Supplementary Table S5: Differential enrichment analysis comparing LUAD Mucinus vs. Non-Mucinus BAC types (Results)**

**Supplementary Table S6: SLAPenrich results across 10 different TCGA datasets**

**Supplementary Table S7: Keywords used to manually curated the mapping between genes, pathways and hallmarks**

**Supplementary Table S8: Manually curated mapping between genes, pathways and hallmarks**

**Supplementary Table S9: SLAPenrich results across 10 different TCGA datasets (excluding cancer driver genes)**

### Supplementary Note

#### Overview of the exposed functions in the SLAPenrich R package

SLAPenrich is implemented as an open-source Bioconductor R-package and it is public available on GitHub (at https://github.com/saezlab/SLAPenrich/). It contains seven exposed functions available to the user, nine internal functions and two data objects.

The referenced equations are contained in the formal description of the statistical framework underlying SLAPenrich, contained in the Methods section of the main text.

##### Input/Output

Of the thirteen exposed functions, two are for data input/output: the first one,

```
SLAPE.readDataset(filename),
```

reads a dataset stored in a .csv file as a sparse binary matrix; the second one,

```
SLAPE.write.table(PFP,EM,filename='', fdrth=Inf,exclcovth=0, PATH_COLLECTION,
GeneLenghts),
```

extracts from the `PFP` (pathway fingerprints) object (outputted by the `SLAPE.analysis` function, see below) the subset of pathways (from the collection specified in the pathway collection specified in `PATH_COLLECTION`) whose enrichment false discovery rate is below the threshold specified in the parameter `fdrth` and whose exclusive coverage (see methods) is above the threshold specified in the parameter `exclcovth`. The extracted enriched pathways are then assembled and written in the csv file specified in the parameter `filename`, together with other information such as, for example, the percentage of altered samples of the initial dataset (specified in the matrix `EM`) when considering individual genes in a given pathway, and total exonic block lengths of the enriched pathways (extracted from the data object specified in `GeneLenghts).`

##### Core analysis

The core analysis function implementing the statistical framework described in the methods section is

```
SLAPE.analyse(EM, show_progress=TRUE, correctionMethod='fdr', NSAMPLES=1,
NGENES=1, accExLength=TRUE, BACKGROUNDpopulation=NULL, PATH_COLLECTION,
path_probability='Bernoulli', Rho=10^-6,GeneLenghts)
```

This function takes in input a dataset (the parameter `EM`) stored in a sparse binary matrix, or a sparse matrix with integer non-null entries. In this matrix the columns correspond to samples, the rows correspond to genes and a non-zero entry indicates the presence of a somatic mutations harbored by a given sample in a given gene. If the matrix contains integer entries then they are deemed as the number of somatic point mutations harbored by a given sample in a given gene (these values will be considered to account for the sample mutation rate if the analysis takes into account of the gene exonic lengths, or converted in binary values otherwise, see below).

For each pathway gene-set $P$ in the pathway collection specified in the parameter `PATH_COLLECTION`, and an inputted genomic dataset (summarized by the parameter `EM`), this function computes first of all a vector of probabilities $\pi = \{p_i\}$ quantifying how likely each sample is to harbor at least one somatic mutation in a gene belonging to $P$, by random chance.

These probabilities are computed by default using a Bernoulli model accounting for the total exonic block lengths of all the genes belonging to $P$, and the expected or observed background mutation rate (Equation 5) [1,2] (as specified by the parameter `Rho`, which in the second case should be set equal to `NULL`). Alternatively, these probabilities can be computed through a complementary cumulative hypergeometric distribution evaluated at $X = 0$ and taking into account of the mutation burden of the samples, the size of $P$ in terms of number of genes (Equations 2 and 3, and `accExLength = FALSE`), or its total exonic content block length (AF Equation 4 and `accExLength = TRUE`, the default setting). In all the tests make use of a gene background population that can be defined by the user (through the parameter `BACKGROUNDpopulation`) or assembled pooling together all the genes belonging to at least one pathway of the collection specified in `PATH_COLLECTION`.

After $\pi$ has been computed, this function computes a *pathway alteration* score at the population level, quantifying the deviance of the number of samples in the datasets harbouring at least a somatic mutation in at least one gene of $P$, $O(P)$ (Equation 7) from its random expectation $E(P)$ (Equation 6, which is computed summing the $\{p_i\}$ across all the samples). Finally, it computes the significance of this score with a p-value against the null hypothesis: "$O(P)$ is drawn from a Poisson binomial distribution with $\{p_i\}$ success probabilities" (Equation 9). This comes from the observation that if there is no tendency for a given pathway to be recurrently mutated across $m$ samples of the datasets, then each of these samples can be considered as the observation of a single Bernoulli trial (in a series of $m$ of them), where the event under consideration in the $i$-th trial is "At least one gene belonging to $P$ is mutated in the $i$-th sample". The success probability of this event is given by $p_i$. Worthy of note is that a Poisson binomial distribution should be considered instead of a simple binomial distribution because the $p_i$ are, of course, not identical.

After alteration scores and corresponding significance have been assessed for all the pathways considered in the analysis, resulting p-values are corrected for multiple hypothesis testing with a user-defined method, specified by the parameter `correctionMethod`. Possible values for this parameter are all the admissible values of the parameter `method` in the built-in function `p.adjust` of R, plus *qvalue* through which the user can select the Storey-Tibshirani [3,4] correction method. Through the parameters `NSAMPLES` and `NGENES` the minimal values that the number of samples harbouring a mutation in the pathway $P$, and the number of genes in $P$ mutated in at least one sample should assume in order for $P$ to be included in the analysis can be specified, respectively. The default value for these two parameters is 1.

As mentioned, the two parameters `accExLength` and `BACKGROUNDpopulation` specify whether the gene exonic lengths should be taken into account while defining the probabilities $\{p_i\}$ described above, and the collection of official symbols of the genes that should be included in the background population in the used statistical framework, respectively. If the value of `accExLength` is `TRUE` (default) then the non-null values of the matrix coding for the inputted dataset (`EM`) are deemed to indicate the number of somatic point mutations harbored by a given gene in a give sample. `BACKGROUNDpopulation` could be, for example, all the genes whose mutational status is accounted in the inputted dataset `EM`. If the value of `BACKGROUNDpopulation` is `NULL` (default) then the set of all the genes included in at least one pathway of the collection included in the analysis is used as background population.

Finally, the parameter `show_progress` determines if a progress bar should be visualized during the execution of the analysis.

For an inputted dataset of $m$ samples and a collection of $p$ pathways included in the analysis, `SLAPE.Analyse` outputs also (i) a $p \times m$ binary *pathway alteration matrix* where rows indicate pathways, columns indicate samples and non-null entries indicate the presence of at least a somatic mutation in at least one gene of a given pathway in a given sample; (ii) a $p \times m$ *pathway mutation probability matrix*, where the $j$-th row contains the vector of probabilities $\{p_{j,i}\}$ of the $i$-th sample harboring at least a somatic mutation in at least one gene of the $j$-th pathway, by random chance; (iii) a vector of *pathway alteration expectations* (with an element for each pathway) with an estimation of the expected number of samples harbouring at least one somatic mutation in at least on gene of a given analysed pathway; (iv) a vector of *pathway exclusive coverage scores* quantifying the tendency of the genes composing each of the analysed pathway to be mutated in a mutual exclusive fashion; (v) a list of individual *binary pathway alteration matrices* (one for each analysed pathway), where the generic matrix $M_i$ has dimensions $k \times m$, where $k$ is the number of genes in the $i$-th pathway, $m$ is the number of samples in the analysed dataset and a generic non-null entry in position $h, j$ is equal to 1 if the $h$-th gene of the $i$-th pathway of the analyzed collection harbors at least one somatic mutation in the $j$-th sample; (vi) a vector of *numerical pathway identifiers*.

#### Visualisation

Storing the results outputted by the `SLAPE.Analyse` function in a list, it is possible to visualize them systematically and to produce pdf files with resulting plots using the function

```
SLAPE.serialPathVis(EM, PFP, fdrth=5, exCovTh=50, PATH='./', PATH_COLLECTION).
```

This function extracts from the list of results outputted by `SLAPE.Analyse` (specified by the `PFP` parameter) those pathways (from the collection specified by the parameter `PATH_COLLECTION`) with an enrichment false discovery rate (FDR) below the user defined threshold value specified by the parameter `fdrth`, and with an exclusive coverage score (Equation 12) above the threshold value specified by the parameter `exCovTh`. All the figures produced by this function are stored in the directory specified by the parameter `PATH`.
After selecting the pathways following the user definitions, this function systematically calls for each of them (distinguished by their numerical identifier, `Id`) the sub-routine:

```
SLAPE.pathVis(EM, PFP, Id, i=NULL,PATH='./', PATH_COLLECTION).
```

Before producing the plots, `SLAPE.pathVis` rearranges rows and columns of the *alteration matrix* of the `Id` pathway through a heuristic mutual-exclusivity sorting procedure (detailed in the method), which highlights the tendency of the composing genes to be mutated in a mutual exclusive fashion across the samples of the analyzed dataset. This sorting is implemented in the function

```
SLAPE.heuristic_mut_ex_sorting(EM),
```

which is exported, therefore available to the user and suitable for sorting any type of binary matrix. After this re-arrangement the *alteration matrix* of the `Id` pathway is visualized and stored in a pdf file as a binary heatmap with genes on the rows, samples on the columns, and blue entries indicating the presence of somatic mutations in given gene/sample combinations (an example is reported in Supplementary Figure S1A). Additionally, *mutation probabilities* and *alteration expectations* are visualized and saved as bar diagrams, together with other statistical scores in a separate figure file.

#### Extraction of core-components from the enriched pathways

The function

```
SLAPE.core_components(PFP, EM, PATH='./',
fdrth=Inf, exclcovth=0, PATH_COLLECTION),
```

identifies sets of core-components genes frequently altered and shared by multiple enriched pathways identified by the `SLAPE.Analyse` function (and specified in `PFP`). These core-component gene sets are supposed to lead the enrichment outcomes. To detect such core components, the function executes a fast greedy community detection algorithm [5,6], implemented in the `fastgreedy.community` function of the `iGraph` package [7-9]. The analysis performed by this function considers only enrichments at a false discovery rate lower than the threshold value specified in the parameter `fdrth` and corresponding to pathways with an exclusive coverage greater than the threshold value specified in the parameter `exclcovth`. After these core-component gene-sets have been identified, this function visualizes them together with the pathways they belong to through a set of *membership matrices*: binary heatmaps with pathways on the columns, a set of core-component genes on the rows and non-empty entries specifying to which enriched pathway each gene belongs to (an example is provided in Supplementary Figure S1B). These heatmaps are stored in individual pdf files and saved in the directory specified by the `PATH` parameter.

#### Differential pathway enrichment analysis

The function

```
SLAPE.diff_SLAPE_analysis (EM, contrastMatrix, positiveCondition,
negativeCondition, SLAPE.FDRth=5, ...)
```

performs a differential enrichment analysis of pathway alterations at the sample population level between two sample subsets of dataset specified in the parameter `EM` (defined as for the previous function). The parameter `contrastMatrix` specifies which samples are included in each sub-population and it is a binary matrix with sample identifiers on the rows and condition identifiers on the columns. The sample identifiers should match those of the initial datasets, i.e. the column headers of the `EM` matrix. A 1 in the position $i, j$ of such a matrix indicates that the i-th sample is included in the sub-population corresponding to the j-th condition.
The two sub-populations to be contrasted are specified by the parameters `positiveCondition` and `negativeCondition` that should match two different column headers of the `contrastMatrix`. This function first performs two independent SLAPenrich analyses on the two user-defined sub-populations of samples with an experimental setting specified by the additional parameters (not listed in the function signature above), which are the same of the `SLAPE.analyse` function. Then it selects the pathways with an enrichment FDR smaller than the value specified in the `SLAPE.FDRth=5` in at least one of the two independent analyses. For this set of pathways a differential enrichment score is computed, as detailed in the method, and summary heatmaps are visualised, as shown in Supplementary Figure S1C.

##### Accessory functions and data objects

The SLAPenrich package contains additional exposed functions allowing users to:

- check the consistency of (and possibly update the) gene symbol identifiers in both genomic datasets and pathway collection data object with the Hugo gene nomenclature (HGNC) catalogue, including approved gene symbols and previously used synonyms, contained in the `SLAPE.hgnc.table_20160210` data object;
- update or create a new HGNC catalogue, by downloading the most up-to-date version from the HUGO Gene Nomenclature Committee web-site (www.genenames.org);
- compute the total length of the exonic block of a given gene, making use of the gene exon attribute data object `SLAPE.all_genes_exonic_lengths_ensemble_20160209`, containing the genomic coordinates of all the exons for all the genes (the genome-wide total exonic block lengths are already precomputed and available in the

SLAPE.all_genes_exonic_content_block_lengths_ensemble_20160209 data object, and can be updated with a dedicated function);

- update the gene exon attribute data object, by making use of functions from the biomaRt R package [10-12].

Additionally, different collections of pathway gene sets from the Pathway Commons data portal (v4-201311) [7,9,13-16], and their post-processed versions computed as detailed in the methods to reduce redundancies are embedded in the package as R objects. These objects contain, for each pathway, multiple information such as uniprot identifiers of the composing genes, official data source, and pathway title/definition.

Finally the genomic datasets and corresponding patient clinical information used in the case study described in the following sections are also provided as R objects.
