## Supplementary Materials for "Pathway-based dissection of the genomic heterogeneity of cancer hallmarks’ acquisition with SLAPenrich" (from another version of this preprint: version 4, 2018-03-16)

### Additional File 7: Novel Cancer driver Networks

**Color legend:**

- Sustaining Proliferative Signaling
- Evading Growth Suppressors
- Avoiding Immune Destruction
- Enabling Replicative Immortality
- Tumour-Promoting Inflammation
- Activating Invasion and Metastasis
- Inducing Angiogenesis
- Genome Instability and Mutation
- Resisting Cell Death
- Deregulating Cellular Energetics

### BRCA

Collagen Degradation
FDR = 1.1%, FDR nod = 2.2%

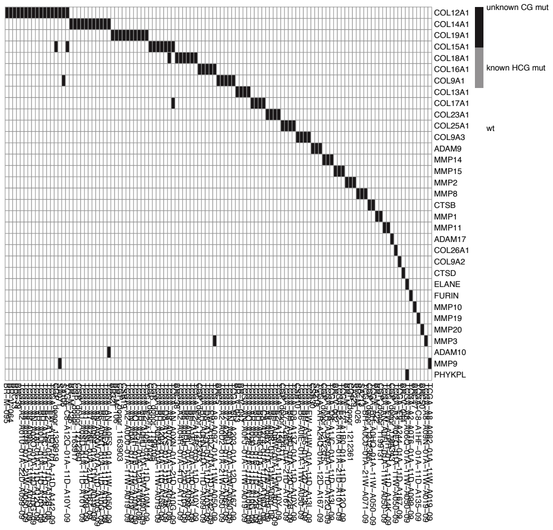


HDACs Deacetylate Histones
FDR = 1.6e−10%, FDR nod = 0.066%

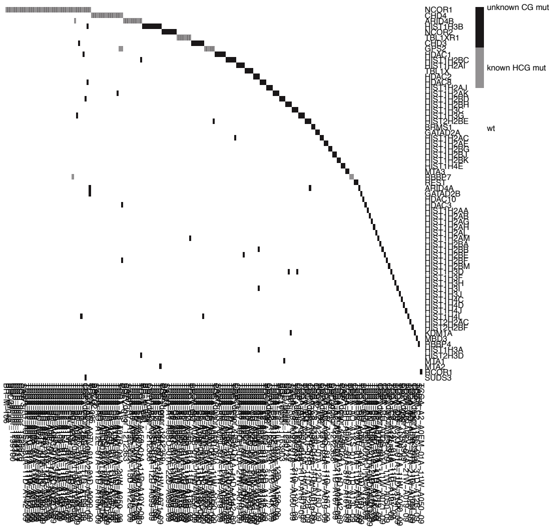


Regulation Of Insulin Secretion
FDR = 0.57%, FDR nod = 1.2%

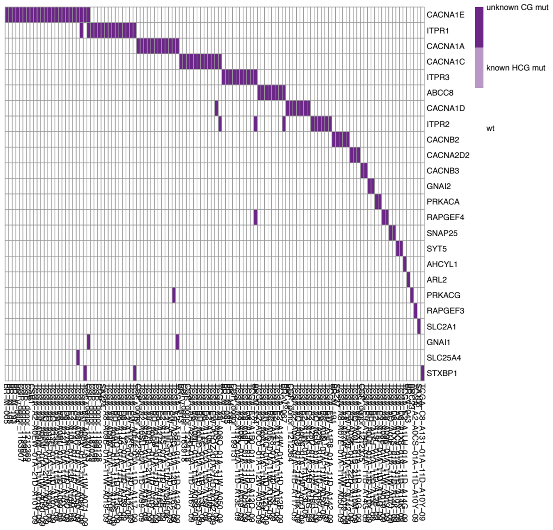


DNA Damage/Telomere Stress Induced Senescence
FDR = 7e−12%, FDR nod = 0.0024%

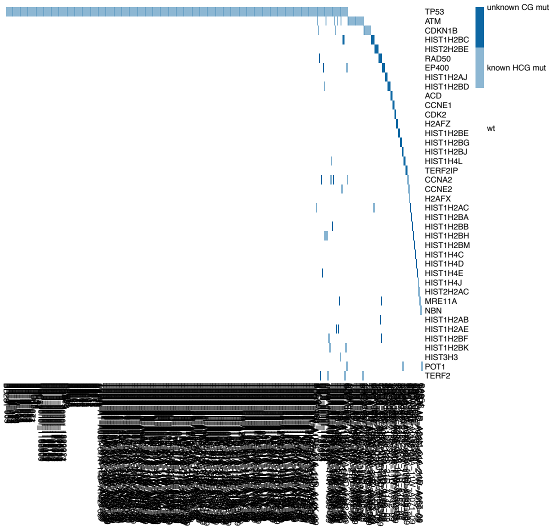


Packaging Of Telomere Ends
FDR = 1.3e−10%, FDR nod = 1.2e−09%

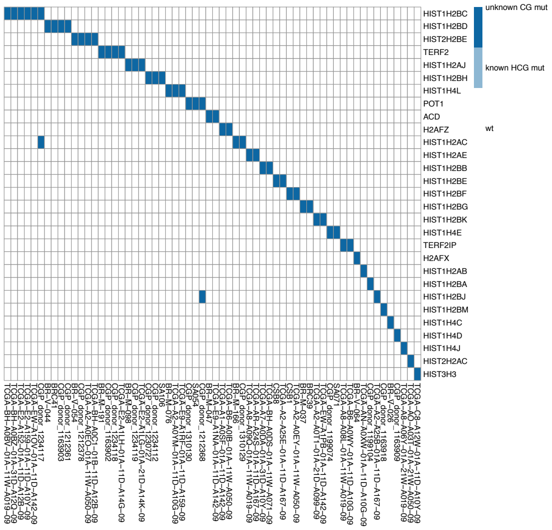


Senescence Associated Secretory Phenotype (SASP)
FDR = 0.00031%, FDR nod = 0.016%

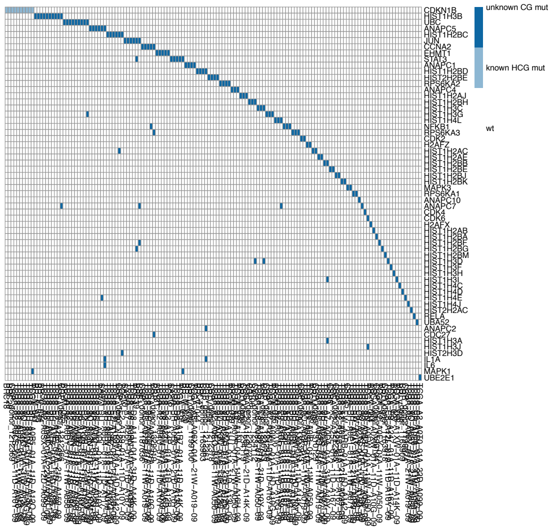

**G2/M DNA Damage Checkpoint**
**Processing Of DNA Double Strand Break Ends**
**FDR = 7e−12%, FDR nod = 1.1%**

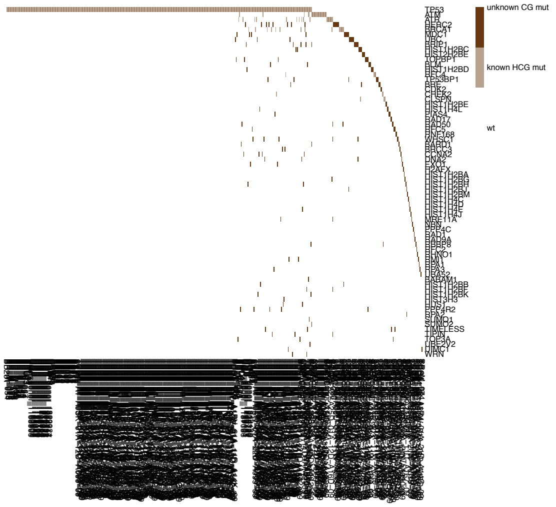


**DNA Damage/Telomere Stress Induced Senescence**
**FDR = 7e−12%, FDR nod = 0.0024%**

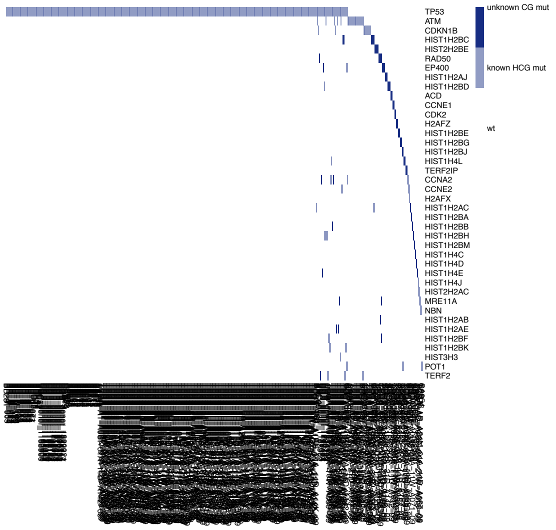


**G2/M DNA Damage Checkpoint**
**Processing Of DNA Double Strand Break Ends**
**FDR = 7e−12%, FDR nod = 1.1%**

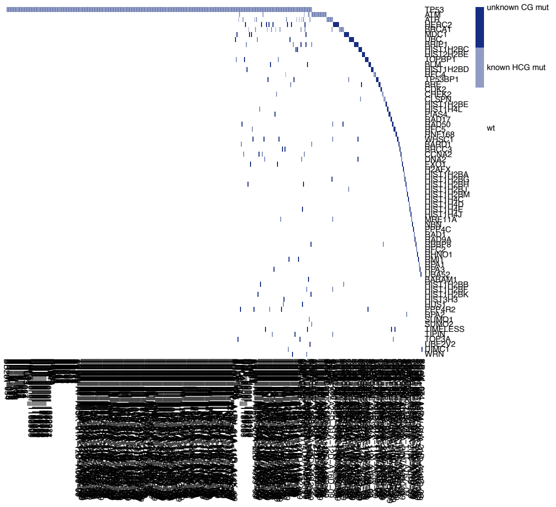


**Meiotic Recombination**
**FDR = 8.1e−12%, FDR nod = 3.6e−06%**

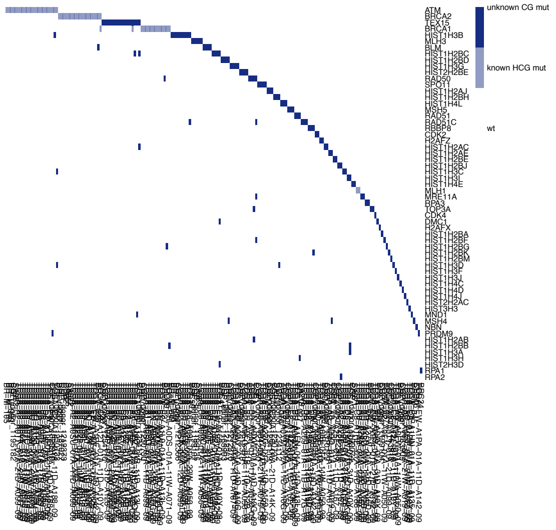


**Meiotic Synapsis**
**FDR = 7.3e−12%, FDR nod = 6.3e−10%**

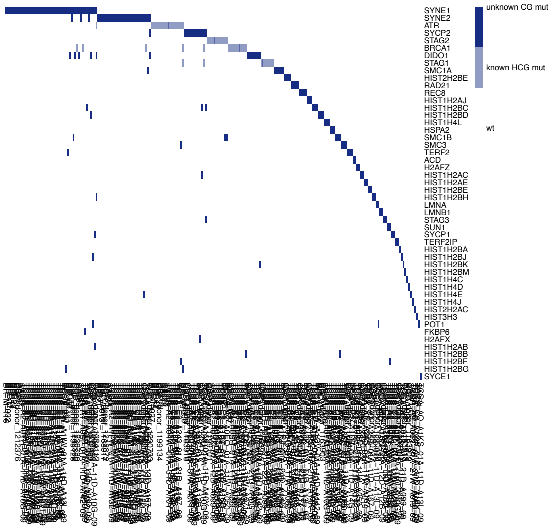


**Integrins In Angiogenesis**
**FDR = 7e−12%, FDR nod = 1.4%**

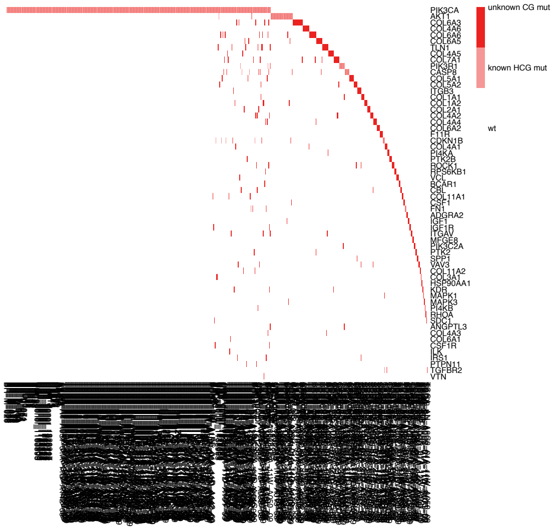

**Signaling By PDGF**
**FDR = 0.0026%, FDR nod = 0.0058%**

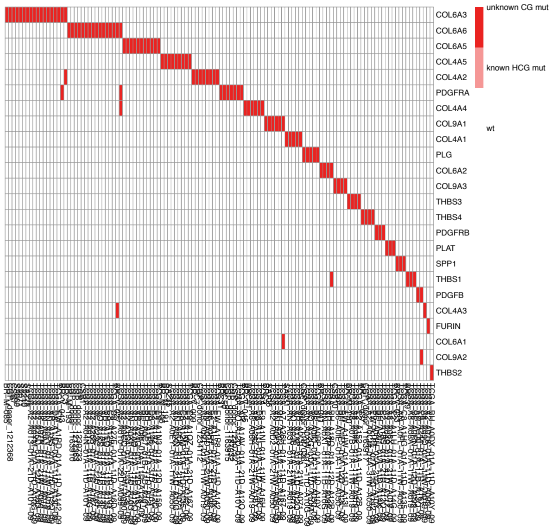


**Wnt Signaling Pathway**
**FDR = 1.4e−09%, FDR nod = 0.00094%**

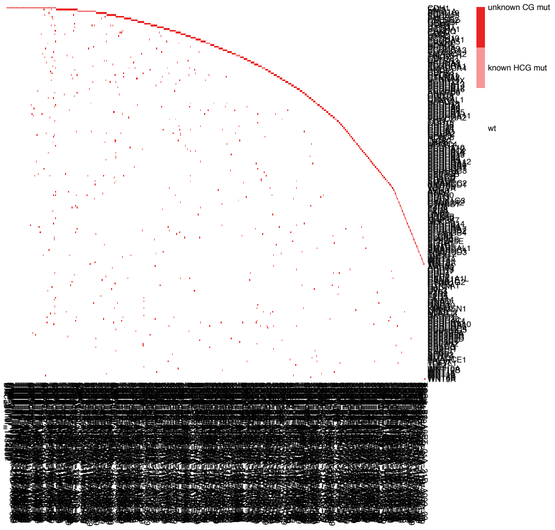


**G2/M DNA Damage Checkpoint**
**Processing Of DNA Double Strand Break Ends**
**FDR = 7e−12%, FDR nod = 1.1%**

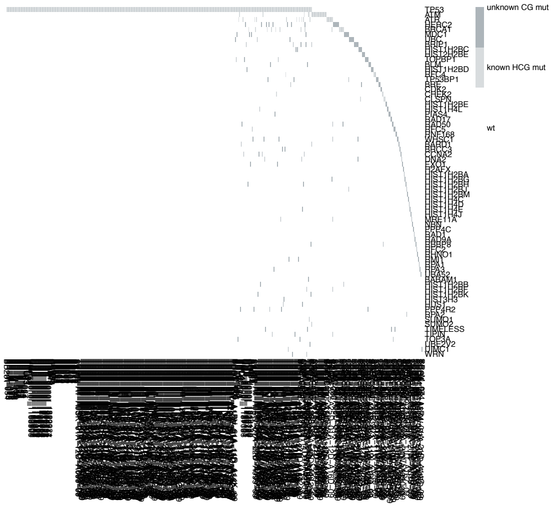


**Constitutive Signaling By Aberrant PI3K In Cancer**
**FDR = 7.3e−12%, FDR nod = 2.8%**

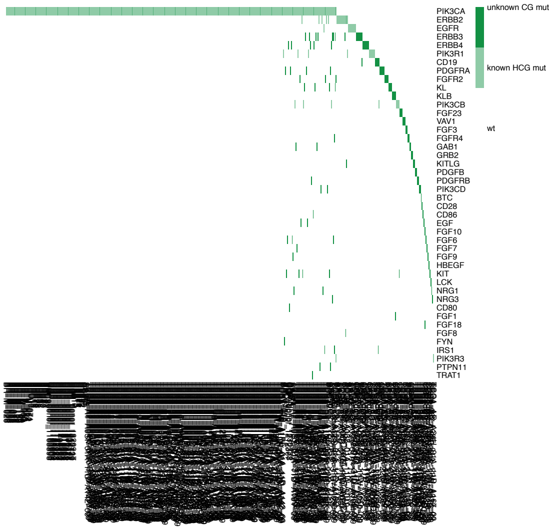


**GRB2:SOS Provides Linkage To MAPK Signaling For Integrins**
**FDR = 1.9%, FDR nod = 1.3%**

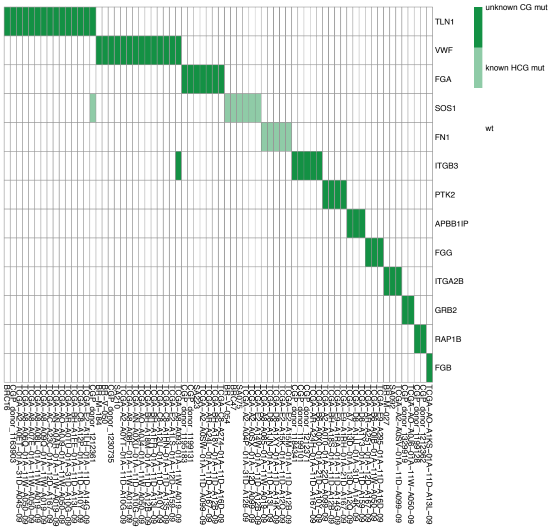


**Integrin Signalling Pathway**
**FDR = 7e−12%, FDR nod = 0.95%**

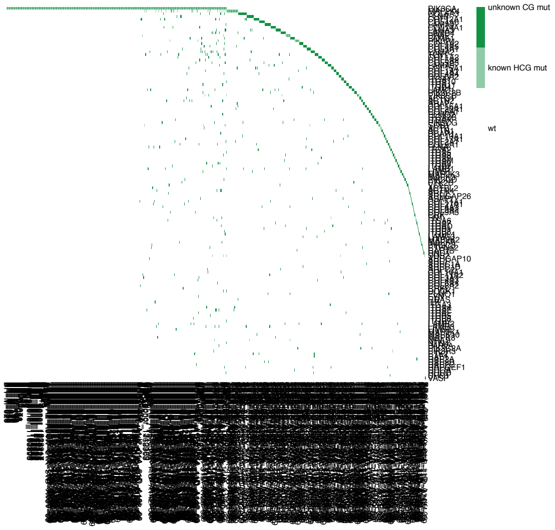

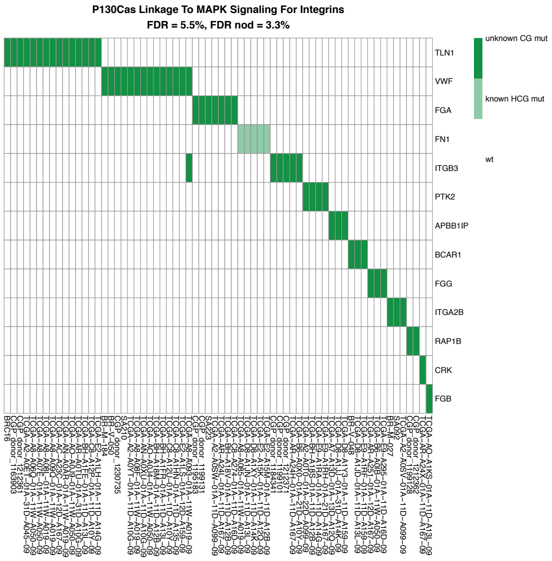
P130Cas Linkage To MAPK Signaling For Integrins
FDR = 5.5%, FDR nod = 3.3%
TLN1
VWF
FGA
FN1
ITGB3
PTK2
APBB1IP
BCAR1
FGG
ITGA2B
RAP1B
CRK
FGB
unknown CG mut
known HCG mut
wt

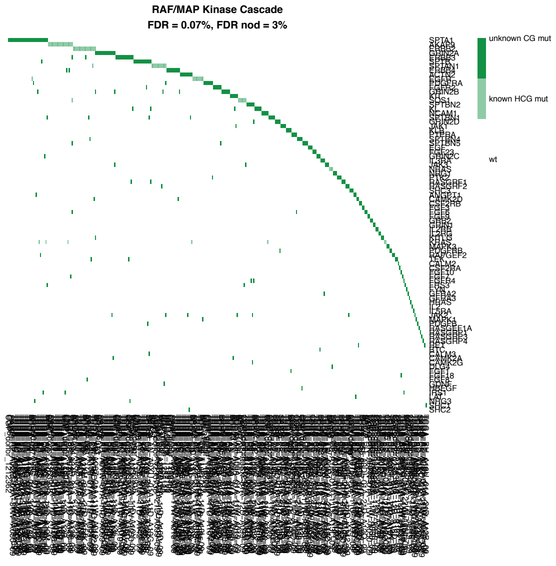
RAF/MAP Kinase Cascade
FDR = 0.07%, FDR nod = 3%
unknown CG mut
known HCG mut
wt

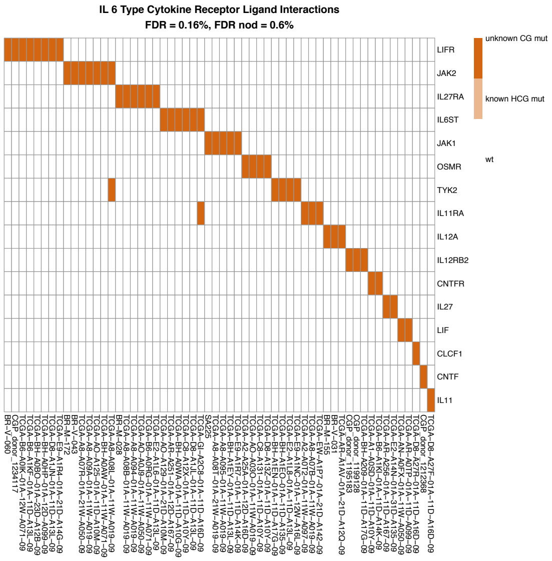
IL 6 Type Cytokine Receptor Ligand Interactions
FDR = 0.16%, FDR nod = 0.6%
LIFR
JAK2
IL27RA
IL6ST
JAK1
OSMR
TYK2
IL11RA
IL12A
IL12RB2
CNTFR
IL27
LIF
CLCF1
CNTF
IL11
unknown CG mut
known HCG mut
wt

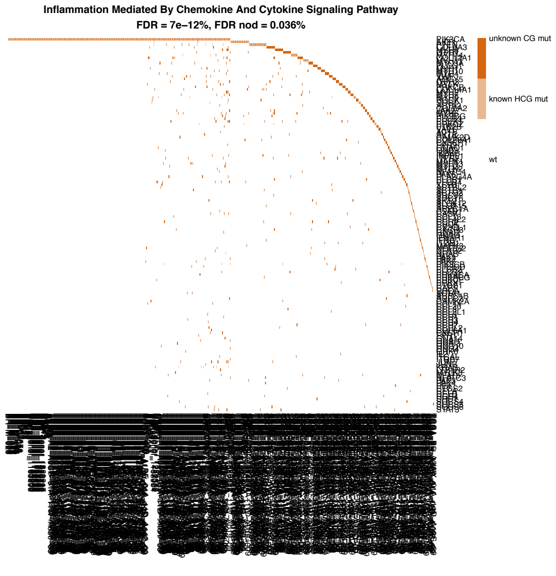
Inflammation Mediated By Chemokine And Cytokine Signaling Pathway
FDR = 7e−12%, FDR nod = 0.036%
unknown CG mut
known HCG mut
wt

### COREAD

**Activation Of Matrix Metalloproteinases**
**FDR = 0.003%, FDR nod = 0.0023%**

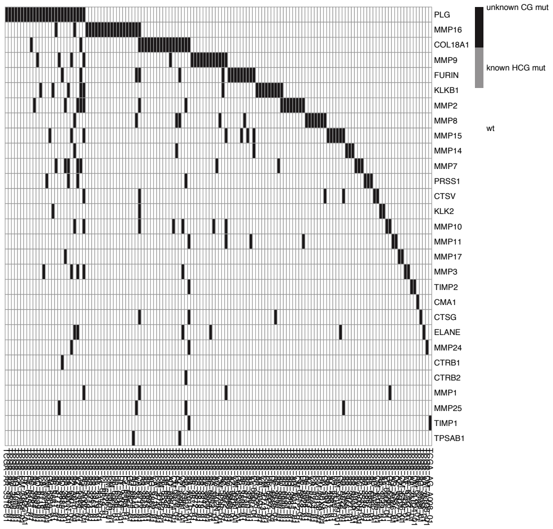


**Collagen Degradation**
**FDR = 0.054%, FDR nod = 0.028%**

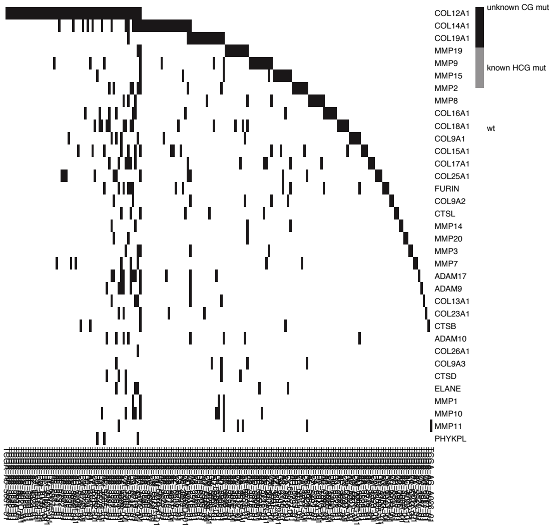


**E Cadherin Signaling In Keratinocytes**
**FDR = 1.9e−12%, FDR nod = 2e−04%**

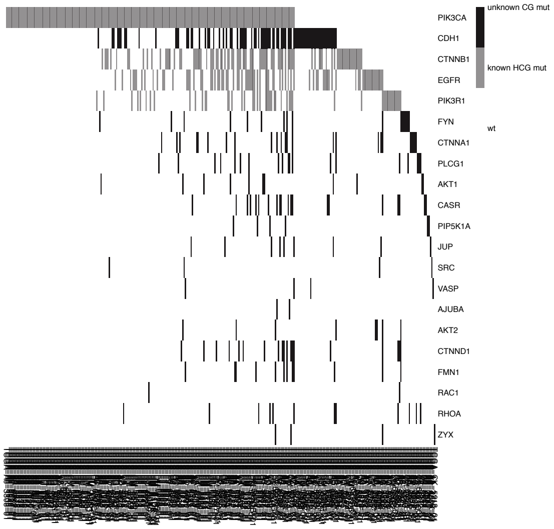


**EPHA2 Forward Signaling**
**FDR = 1.9e−12%, FDR nod = 5.3e−06%**

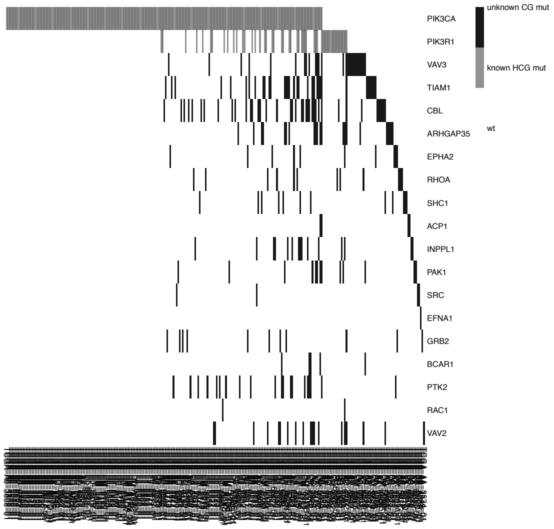


**Ephrin B Reverse Signaling**
**FDR = 1.9e−12%, FDR nod = 5.3e−06%**

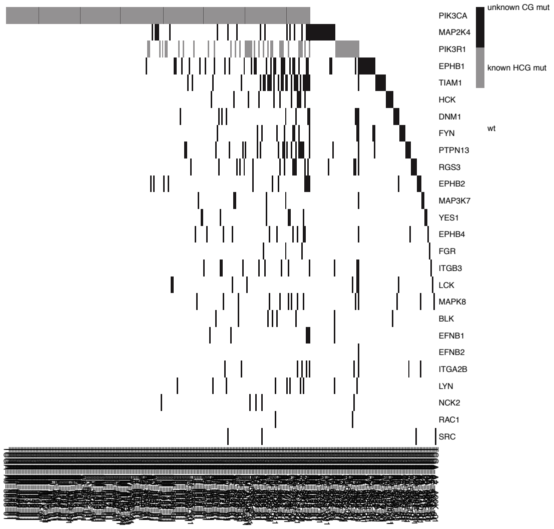


**PDGF Pathway**
**FDR = 8.9e−10%, FDR nod = 0.0023%**

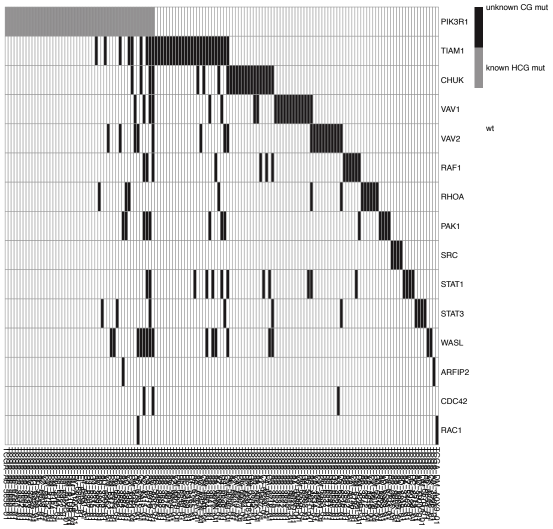

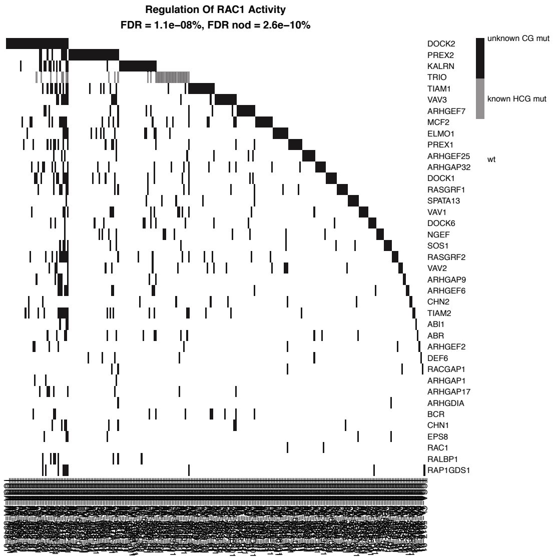
Regulation Of RAC1 Activity
FDR = 1.1e−08%, FDR nod = 2.6e−10%
DOCK2
PREX2
KALRN
TRIO
TIAM1
VAV3
ARHGEF7
MCF2
ELMO1
PREX1
ARHGEF25
ARHGAP32
DOCK1
RASGRF1
SPATA13
VAV1
DOCK6
NGEF
SOS1
RASGRF2
VAV2
ARHGAP9
ARHGEF6
CHN2
TIAM2
ABI1
ABR
ARHGEF2
DEF6
RACGAP1
ARHGAP1
ARHGAP17
ARHGDIA
BCR
CHN1
EPS8
RAC1
RALBP1
RAP1GDS1
unknown CG mut
known HCG mut
wt

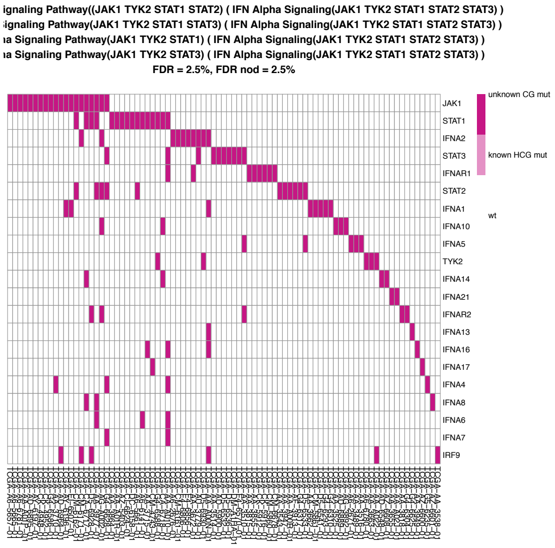
ignaling Pathway((JAK1 TYK2 STAT1 STAT2) ( IFN Alpha Signaling(JAK1 TYK2 STAT1 STAT2 STAT3) )
ignaling Pathway(JAK1 TYK2 STAT1 STAT3) ( IFN Alpha Signaling(JAK1 TYK2 STAT1 STAT2 STAT3) )
a Signaling Pathway(JAK1 TYK2 STAT1) ( IFN Alpha Signaling(JAK1 TYK2 STAT1 STAT2 STAT3) )
a Signaling Pathway(JAK1 TYK2 STAT3) ( IFN Alpha Signaling(JAK1 TYK2 STAT1 STAT2 STAT3) )
FDR = 2.5%, FDR nod = 2.5%
JAK1
STAT1
IFNA2
STAT3
IFNAR1
STAT2
IFNA1
IFNA10
IFNA5
TYK2
IFNA14
IFNA21
IFNAR2
IFNA13
IFNA16
IFNA17
IFNA4
IFNA8
IFNA6
IFNA7
IRF9
unknown CG mut
known HCG mut
wt

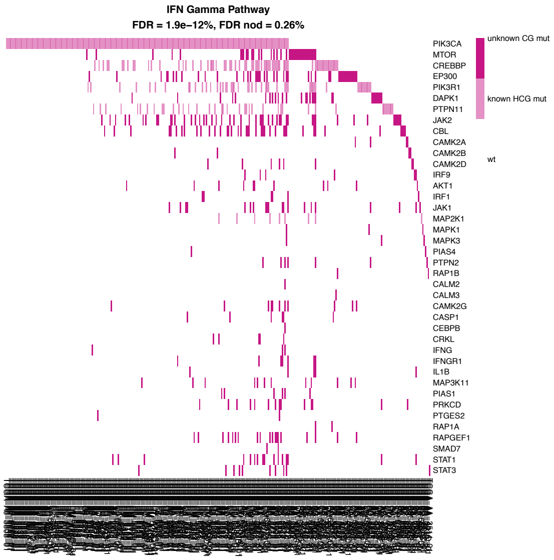
IFN Gamma Pathway
FDR = 1.9e−12%, FDR nod = 0.26%
PIK3CA
MTOR
CREBBP
EP300
PIK3R1
DAPK1
PTPN11
JAK2
CBL
CAMK2A
CAMK2B
CAMK2D
IRF9
AKT1
IRF1
JAK1
MAP2K1
MAPK1
MAPK3
PIAS4
PTPN2
RAP1B
CALM2
CALM3
CAMK2G
CASP1
CEBPB
CRKL
IFNG
IFNGR1
IL1B
MAP3K11
PIAS1
PRKCD
PTGES2
RAP1A
RAPGEF1
SMAD7
STAT1
STAT3
unknown CG mut
known HCG mut
wt

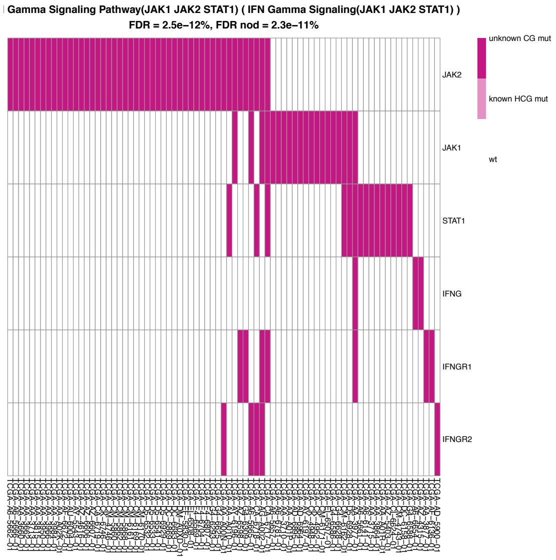
Gamma Signaling Pathway(JAK1 JAK2 STAT1) ( IFN Gamma Signaling(JAK1 JAK2 STAT1) )
FDR = 2.5e−12%, FDR nod = 2.3e−11%
JAK2
JAK1
STAT1
IFNG
IFNGR1
IFNGR2
unknown CG mut
known HCG mut
wt

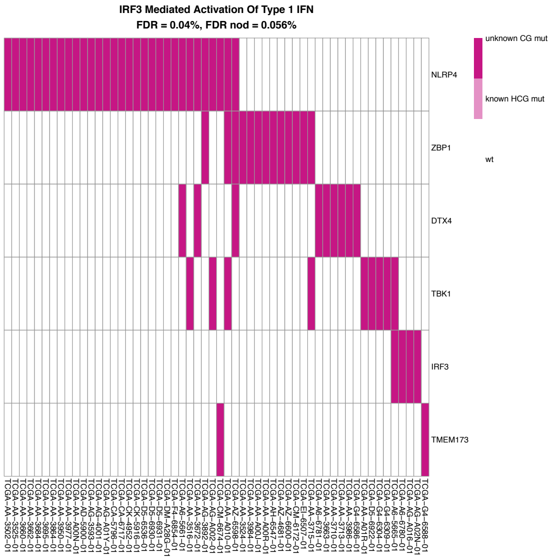
IRF3 Mediated Activation Of Type 1 IFN
FDR = 0.04%, FDR nod = 0.056%
NLRP4
ZBP1
DTX4
TBK1
IRF3
TMEM173
unknown CG mut
known HCG mut
wt

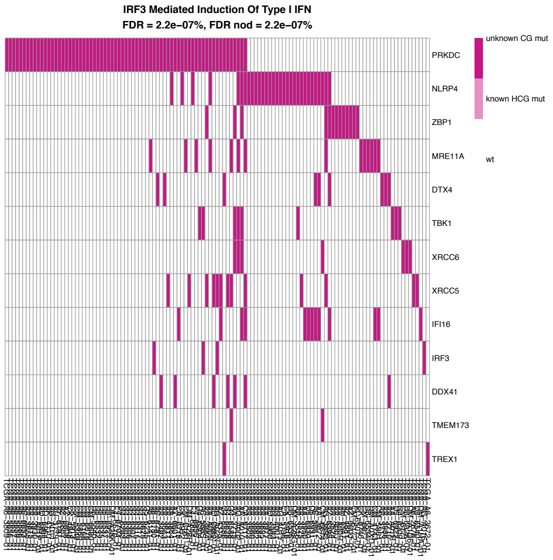
IRF3 Mediated Induction Of Type I IFN
FDR = 2.2e−07%, FDR nod = 2.2e−07%
PRKDC
NLRP4
ZBP1
MRE11A
DTX4
TBK1
XRCC6
XRCC5
IFI16
IRF3
DDX41
TMEM173
TREX1
unknown CG mut
known HCG mut
wt

**LRR FLII Interacting Protein 1 (LRRFIP1) Activates Type I IFN Production**
**FDR = 1.9e−12%, FDR nod = 0.021%**

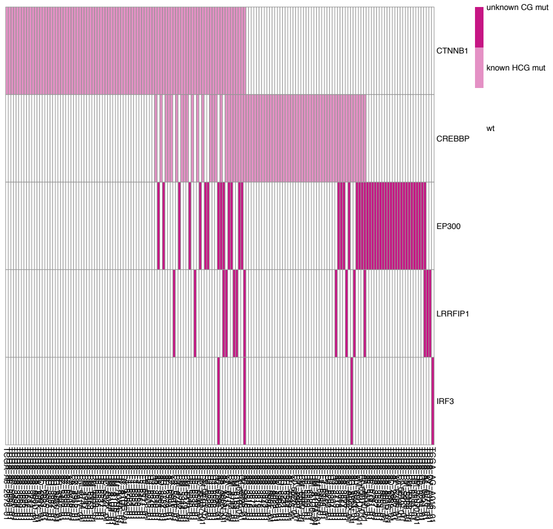


**Regulation Of IFNA Signaling**
**FDR = 1.9e−07%, FDR nod = 0.54%**

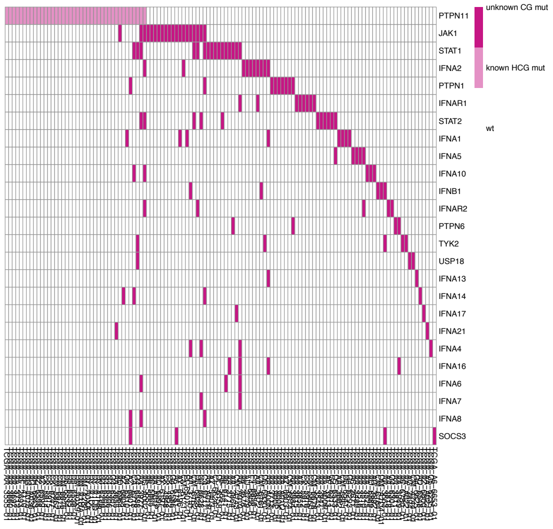


**Regulation Of IFNG Signaling**
**FDR = 1.5e−11%, FDR nod = 0.0025%**

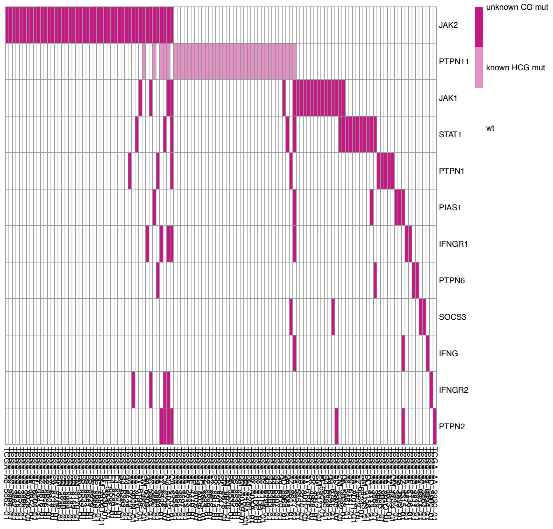


**RIG I/MDA5 Mediated Induction Of IFN Alpha/beta Pathways**
**FDR = 0.8%, FDR nod = 0.78%**

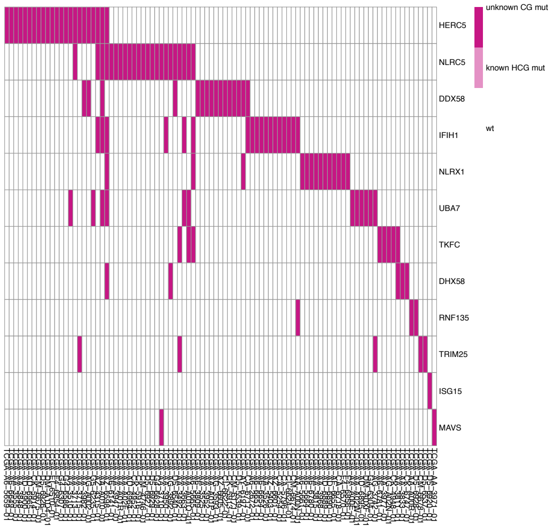


**Energy Dependent Regulation Of MTOR By LKB1 AMPK**
**FDR = 4.4%, FDR nod = 3.2%**

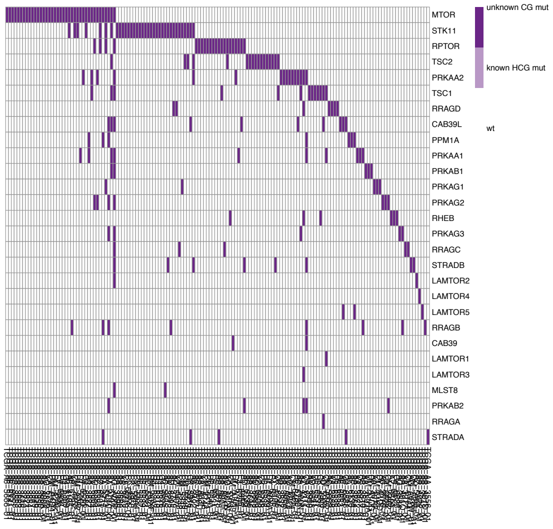


**Fatty Acids Bound To GPR40 (FFAR1) Regulate Insulin Secretion**
**FDR = 0.56%, FDR nod = 0.64%**

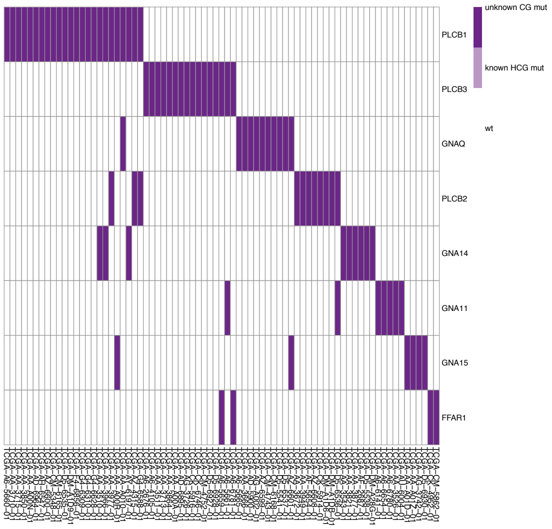

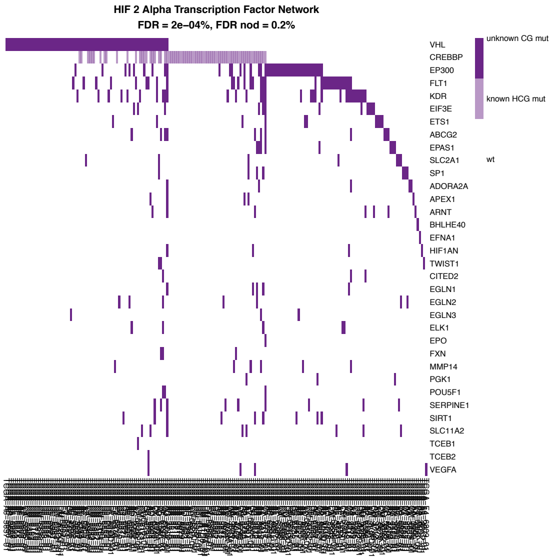
HIF 2 Alpha Transcription Factor Network
FDR = 2e−04%, FDR nod = 0.2%
VHL
CREBBP
EP300
FLT1
KDR
EIF3E
ETS1
ABCG2
EPAS1
SLC2A1
SP1
ADORA2A
APEX1
ARNT
BHLHE40
EFNA1
HIF1AN
TWIST1
CITED2
EGLN1
EGLN2
EGLN3
ELK1
EPO
FXN
MMP14
PGK1
POU5F1
SERPINE1
SIRT1
SLC11A2
TCEB1
TCEB2
VEGFA
unknown CG mut
known HCG mut
wt

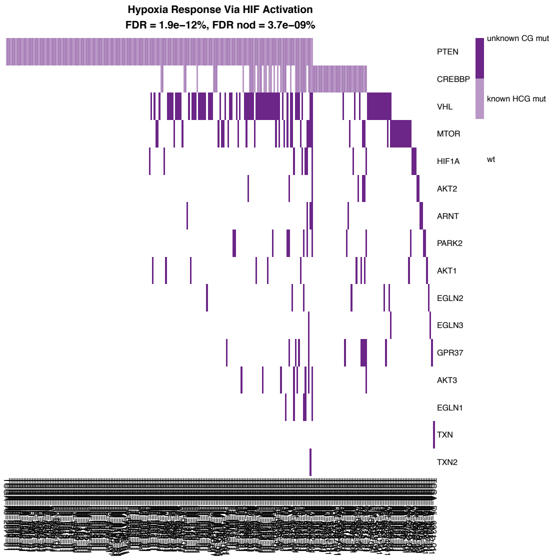
Hypoxia Response Via HIF Activation
FDR = 1.9e−12%, FDR nod = 3.7e−09%
PTEN
CREBBP
VHL
MTOR
HIF1A
AKT2
ARNT
PARK2
AKT1
EGLN2
EGLN3
GPR37
AKT3
EGLN1
TXN
TXN2
unknown CG mut
known HCG mut
wt

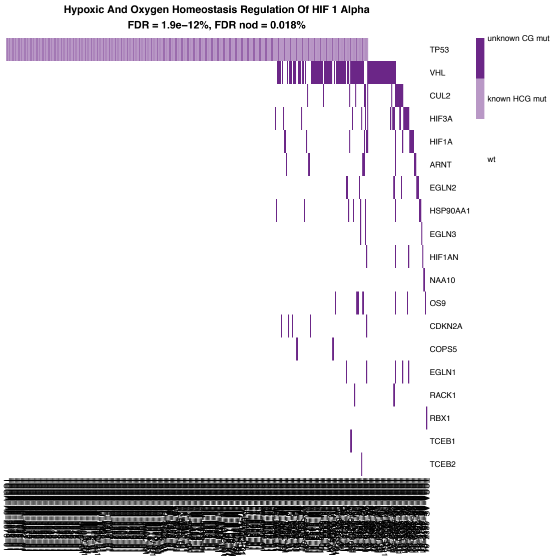
Hypoxic And Oxygen Homeostasis Regulation Of HIF 1 Alpha
FDR = 1.9e−12%, FDR nod = 0.018%
TP53
VHL
CUL2
HIF3A
HIF1A
ARNT
EGLN2
HSP90AA1
EGLN3
HIF1AN
NAA10
OS9
CDKN2A
COPS5
EGLN1
RACK1
RBX1
TCEB1
TCEB2
unknown CG mut
known HCG mut
wt

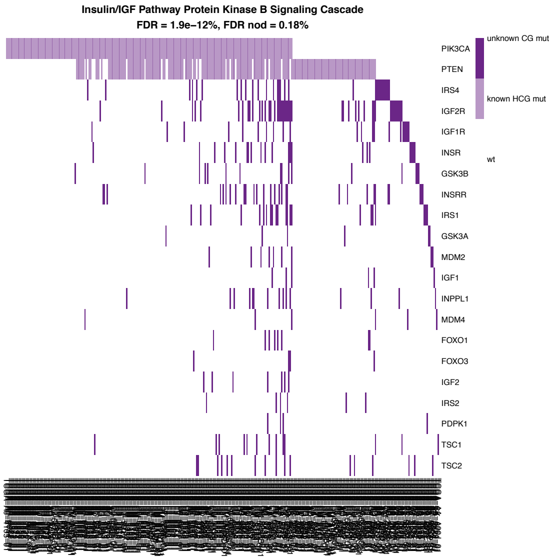
Insulin/IGF Pathway Protein Kinase B Signaling Cascade
FDR = 1.9e−12%, FDR nod = 0.18%
PIK3CA
PTEN
IRS4
IGF2R
IGF1R
INSR
GSK3B
INSRR
IRS1
GSK3A
MDM2
IGF1
INPPL1
MDM4
FOXO1
FOXO3
IGF2
IRS2
PDPK1
TSC1
TSC2
unknown CG mut
known HCG mut
wt

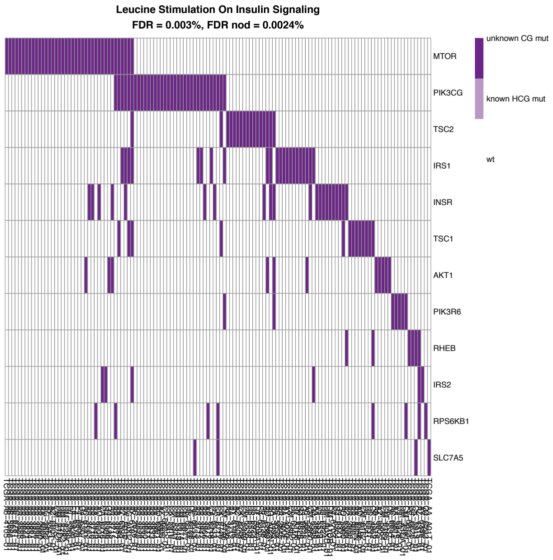
Leucine Stimulation On Insulin Signaling
FDR = 0.003%, FDR nod = 0.0024%
MTOR
PIK3CG
TSC2
IRS1
INSR
TSC1
AKT1
PIK3R6
RHEB
IRS2
RPS6KB1
SLC7A5
unknown CG mut
known HCG mut
wt

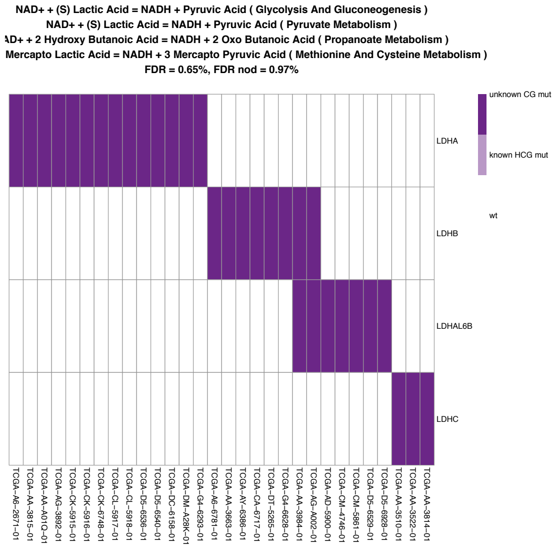
NAD+ + (S) Lactic Acid = NADH + Pyruvic Acid ( Glycolysis And Gluconeogenesis )
NAD+ + (S) Lactic Acid = NADH + Pyruvic Acid ( Pyruvate Metabolism )
AD+ + 2 Hydroxy Butanoic Acid = NADH + 2 Oxo Butanoic Acid ( Propanoate Metabolism )
Mercapto Lactic Acid = NADH + 3 Mercapto Pyruvic Acid ( Methionine And Cysteine Metabolism )
FDR = 0.65%, FDR nod = 0.97%
LDHA
LDHB
LDHAL6B
LDHC
unknown CG mut
known HCG mut
wt
TCGA−A6−2671−01
TCGA−AA−3815−01
TCGA−AA−A01Q−01
TCGA−AG−3892−01
TCGA−CK−5915−01
TCGA−CK−5916−01
TCGA−CK−6748−01
TCGA−CL−5917−01
TCGA−CL−5918−01
TCGA−D5−6536−01
TCGA−D5−6540−01
TCGA−DC−6158−01
TCGA−DM−A28K−01
TCGA−G4−6293−01
TCGA−A6−6781−01
TCGA−AA−3663−01
TCGA−AY−6386−01
TCGA−CA−6717−01
TCGA−DT−5265−01
TCGA−G4−6628−01
TCGA−AA−3984−01
TCGA−AG−A002−01
TCGA−AD−5900−01
TCGA−CM−4746−01
TCGA−CM−5861−01
TCGA−D5−6529−01
TCGA−D5−6928−01
TCGA−AA−3510−01
TCGA−AA−3522−01
TCGA−AA−3814−01

PTK6 Promotes HIF1A Stabilization
FDR = 2e−12%, FDR nod = 2.6e−08%

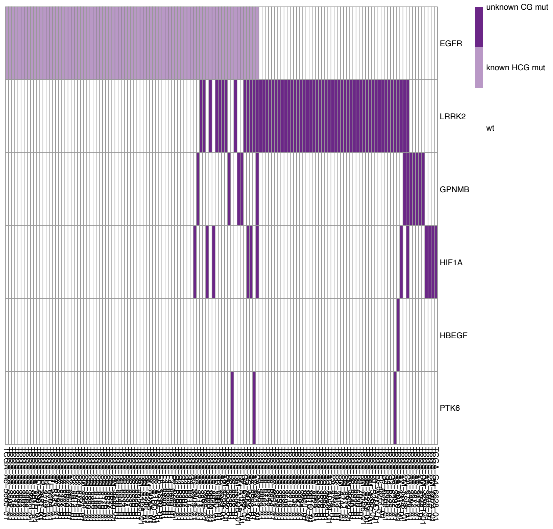


DNA Damage/Telomere Stress Induced Senescence
FDR = 1.9e−12%, FDR nod = 0.14%

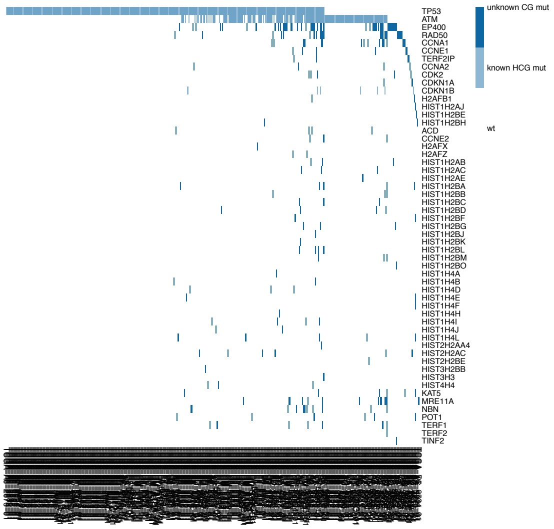


Formation Of Senescence Associated Heterochromatin Foci (SAHF)
FDR = 1.9e−12%, FDR nod = 2.3e−11%

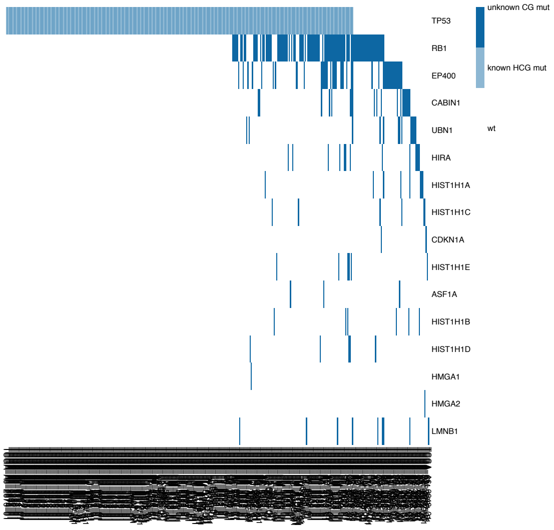


Polymerase Switching
Polymerase Switching On The C Strand Of The Telomere
FDR = 4.4%, FDR nod = 4.6%

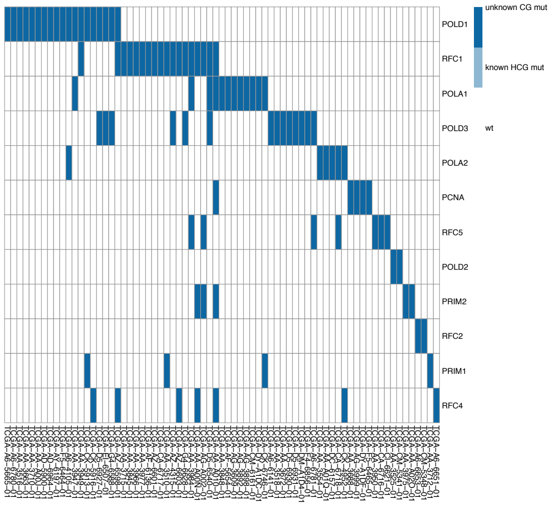


Processive Synthesis On The C Strand Of The Telomere
Telomere C Strand (Lagging Strand) Synthesis
FDR = 1.9e−12%, FDR nod = 2.3e−11%

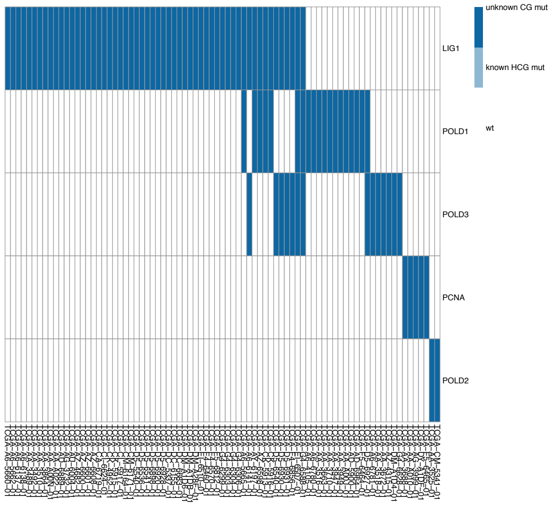


Senescence And Autophagy In Cancer
FDR = 1.9e−12%, FDR nod = 0.72%

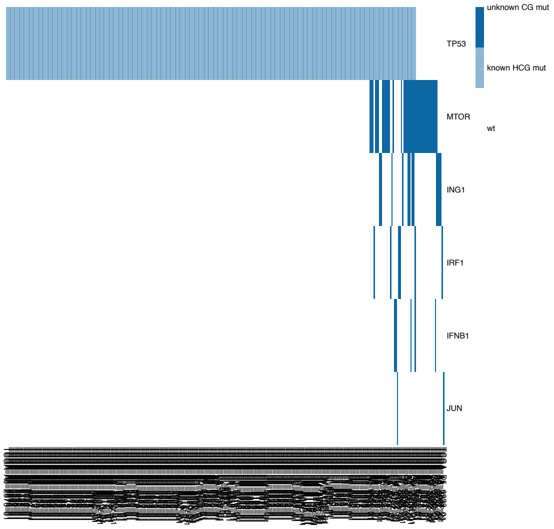

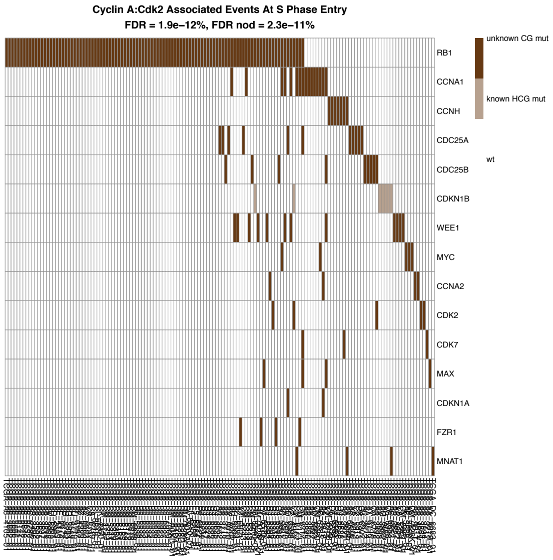
Cyclin A:Cdk2 Associated Events At S Phase Entry
FDR = 1.9e−12%, FDR nod = 2.3e−11%
RB1
CCNA1
CCNH
CDC25A
CDC25B
CDKN1B
WEE1
MYC
CCNA2
CDK2
CDK7
MAX
CDKN1A
FZR1
MNAT1
unknown CG mut
known HCG mut
wt

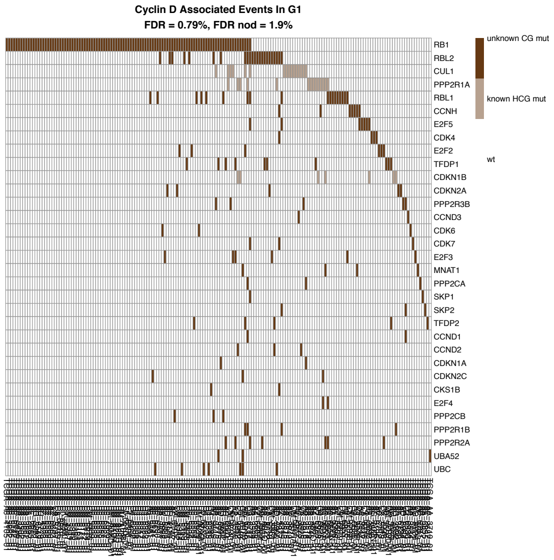
Cyclin D Associated Events In G1
FDR = 0.79%, FDR nod = 1.9%
RB1
RBL2
CUL1
PPP2R1A
RBL1
CCNH
E2F5
CDK4
E2F2
TFDP1
CDKN1B
CDKN2A
PPP2R3B
CCND3
CDK6
CDK7
E2F3
MNAT1
PPP2CA
SKP1
SKP2
TFDP2
CCND1
CCND2
CDKN1A
CDKN2C
CKS1B
E2F4
PPP2CB
PPP2R1B
PPP2R2A
UBA52
UBC
unknown CG mut
known HCG mut
wt

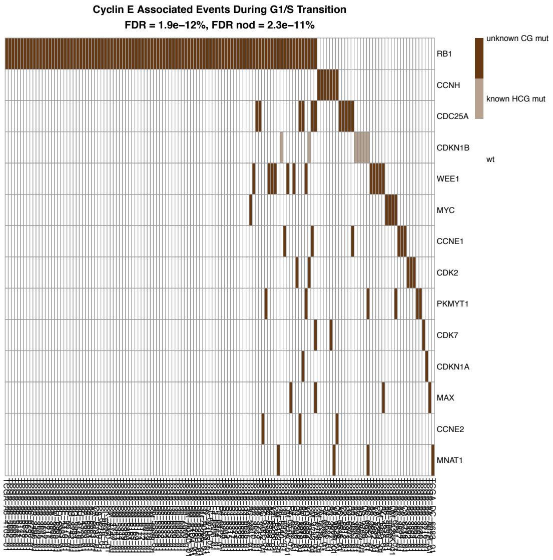
Cyclin E Associated Events During G1/S Transition
FDR = 1.9e−12%, FDR nod = 2.3e−11%
RB1
CCNH
CDC25A
CDKN1B
WEE1
MYC
CCNE1
CDK2
PKMYT1
CDK7
CDKN1A
MAX
CCNE2
MNAT1
unknown CG mut
known HCG mut
wt

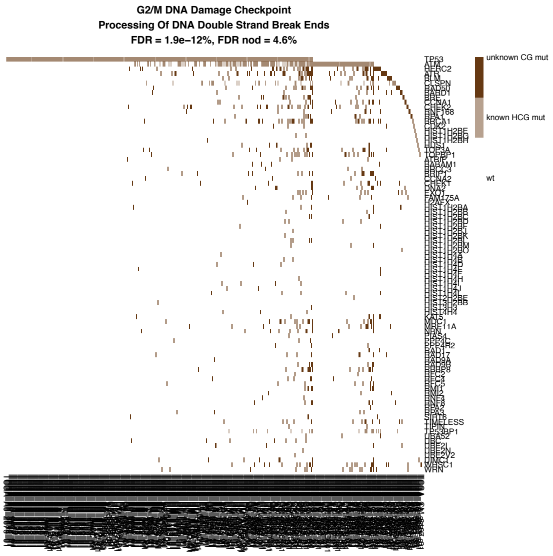
G2/M DNA Damage Checkpoint
Processing Of DNA Double Strand Break Ends
FDR = 1.9e−12%, FDR nod = 4.6%
unknown CG mut
known HCG mut
wt

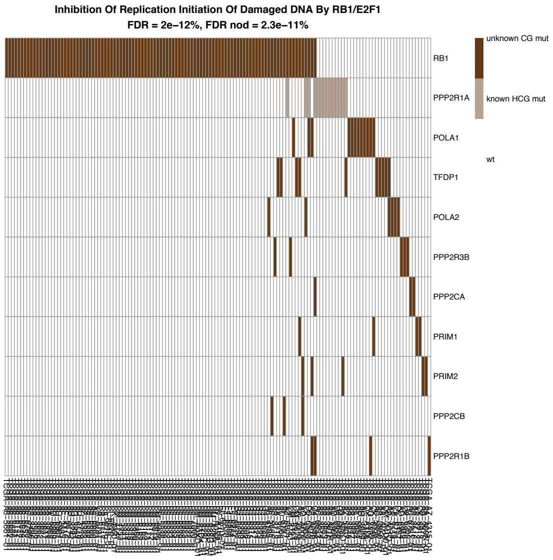
Inhibition Of Replication Initiation Of Damaged DNA By RB1/E2F1
FDR = 2e−12%, FDR nod = 2.3e−11%
RB1
PPP2R1A
POLA1
TFDP1
POLA2
PPP2R3B
PPP2CA
PRIM1
PRIM2
PPP2CB
PPP2R1B
unknown CG mut
known HCG mut
wt

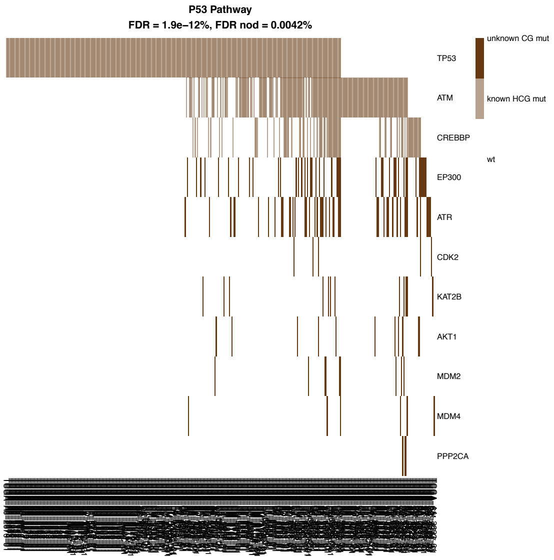
P53 Pathway
FDR = 1.9e−12%, FDR nod = 0.0042%
TP53
ATM
CREBBP
EP300
ATR
CDK2
KAT2B
AKT1
MDM2
MDM4
PPP2CA
unknown CG mut
known HCG mut
wt

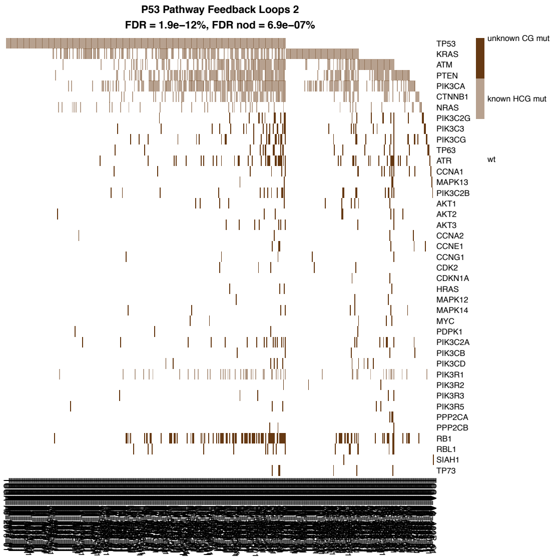
P53 Pathway Feedback Loops 2
FDR = 1.9e−12%, FDR nod = 6.9e−07%
TP53
KRAS
ATM
PTEN
PIK3CA
CTNNB1
NRAS
PIK3C2G
PIK3C3
PIK3CG
TP63
ATR
CCNA1
MAPK13
PIK3C2B
AKT1
AKT2
AKT3
CCNA2
CCNE1
CCNG1
CDK2
CDKN1A
HRAS
MAPK12
MAPK14
MYC
PDPK1
PIK3C2A
PIK3CB
PIK3CD
PIK3R1
PIK3R2
PIK3R3
PIK3R5
PPP2CA
PPP2CB
RB1
RBL1
SIAH1
TP73
unknown CG mut
known HCG mut
wt

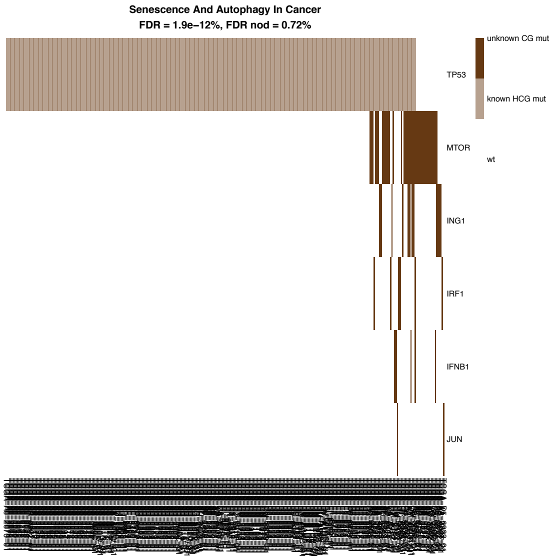
Senescence And Autophagy In Cancer
FDR = 1.9e−12%, FDR nod = 0.72%
TP53
MTOR
ING1
IRF1
IFNB1
JUN
unknown CG mut
known HCG mut
wt

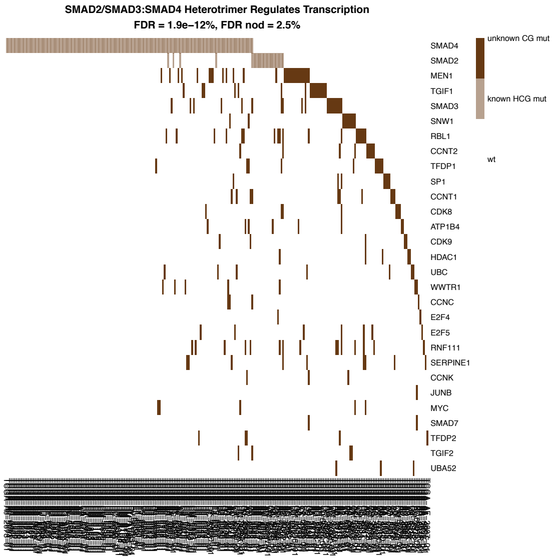
SMAD2/SMAD3:SMAD4 Heterotrimer Regulates Transcription
FDR = 1.9e−12%, FDR nod = 2.5%
SMAD4
SMAD2
MEN1
TGIF1
SMAD3
SNW1
RBL1
CCNT2
TFDP1
SP1
CCNT1
CDK8
ATP1B4
CDK9
HDAC1
UBC
WWTR1
CCNC
E2F4
E2F5
RNF111
SERPINE1
CCNK
JUNB
MYC
SMAD7
TFDP2
TGIF2
UBA52
unknown CG mut
known HCG mut
wt

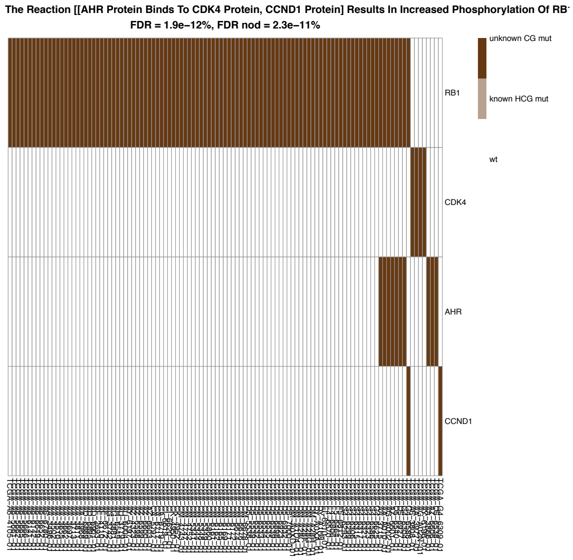
The Reaction [[AHR Protein Binds To CDK4 Protein, CCND1 Protein] Results In Increased Phosphorylation Of RB
FDR = 1.9e−12%, FDR nod = 2.3e−11%
RB1
CDK4
AHR
CCND1
unknown CG mut
known HCG mut
wt

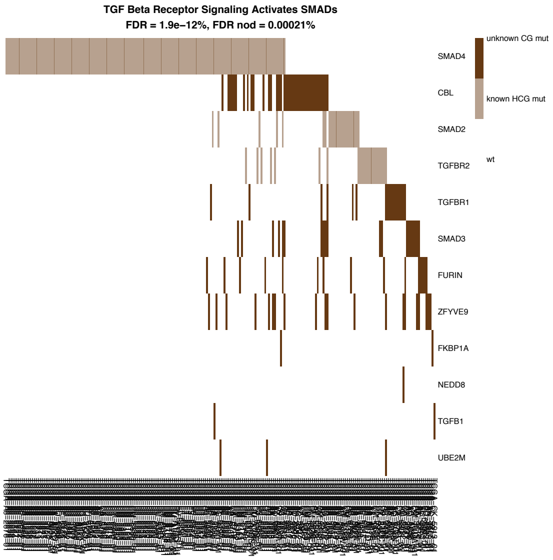
TGF Beta Receptor Signaling Activates SMADs
FDR = 1.9e−12%, FDR nod = 0.00021%
SMAD4
CBL
SMAD2
TGFBR2
TGFBR1
SMAD3
FURIN
ZFYVE9
FKBP1A
NEDD8
TGFB1
UBE2M
unknown CG mut
known HCG mut
wt

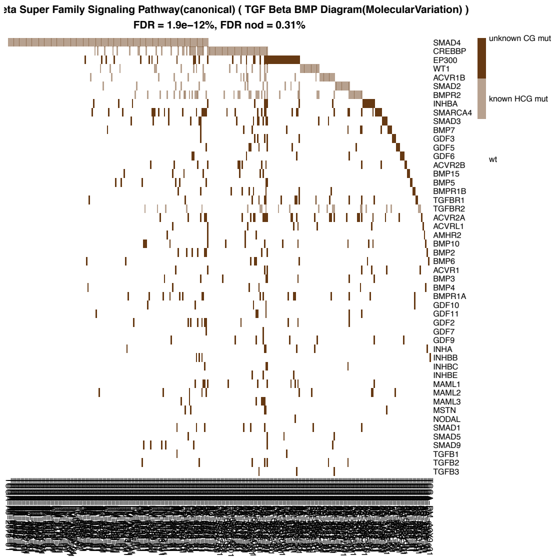
eta Super Family Signaling Pathway(canonical) ( TGF Beta BMP Diagram(MolecularVariation) )
FDR = 1.9e−12%, FDR nod = 0.31%
SMAD4
CREBBP
EP300
WT1
ACVR1B
SMAD2
BMPR2
INHBA
SMARCA4
SMAD3
BMP7
GDF3
GDF5
GDF6
ACVR2B
BMP15
BMP5
BMPR1B
TGFBR1
TGFBR2
ACVR2A
ACVRL1
AMHR2
BMP10
BMP2
BMP6
ACVR1
BMP3
BMP4
BMPR1A
GDF10
GDF11
GDF2
GDF7
GDF9
INHA
INHBB
INHBC
INHBE
MAML1
MAML2
MAML3
MSTN
NODAL
SMAD1
SMAD5
SMAD9
TGFB1
TGFB2
TGFB3
unknown CG mut
known HCG mut
wt

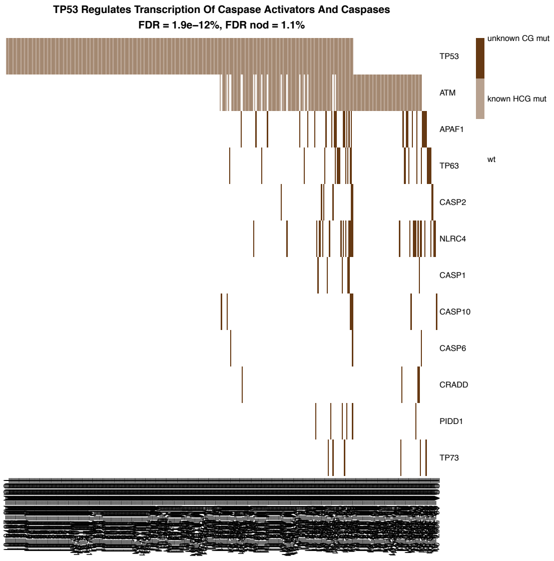
TP53 Regulates Transcription Of Caspase Activators And Caspases
FDR = 1.9e−12%, FDR nod = 1.1%
TP53
ATM
APAF1
TP63
CASP2
NLRC4
CASP1
CASP10
CASP6
CRADD
PIDD1
TP73
unknown CG mut
known HCG mut
wt

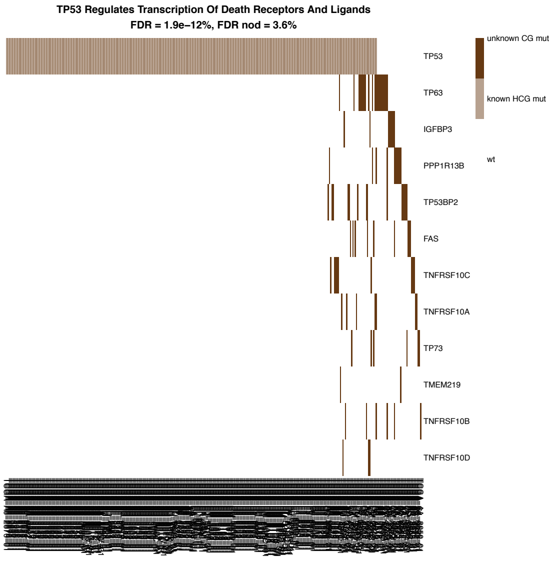
TP53 Regulates Transcription Of Death Receptors And Ligands
FDR = 1.9e−12%, FDR nod = 3.6%
TP53
TP63
IGFBP3
PPP1R13B
TP53BP2
FAS
TNFRSF10C
TNFRSF10A
TP73
TMEM219
TNFRSF10B
TNFRSF10D
unknown CG mut
known HCG mut
wt

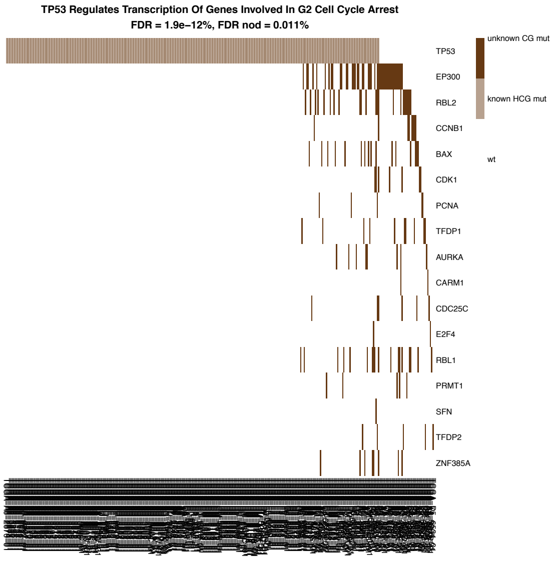
TP53 Regulates Transcription Of Genes Involved In G2 Cell Cycle Arrest
FDR = 1.9e−12%, FDR nod = 0.011%
TP53
EP300
RBL2
CCNB1
BAX
CDK1
PCNA
TFDP1
AURKA
CARM1
CDC25C
E2F4
RBL1
PRMT1
SFN
TFDP2
ZNF385A
unknown CG mut
known HCG mut
wt

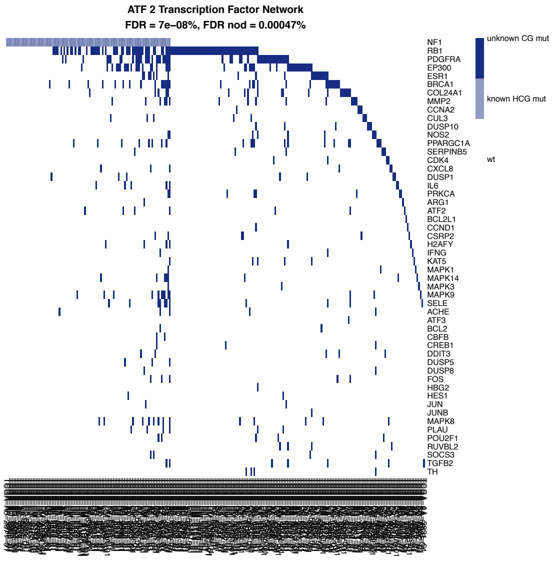
ATF 2 Transcription Factor Network
FDR = 7e−08%, FDR nod = 0.00047%
NF1
RB1
PDGFRA
EP300
ESR1
BRCA1
COL24A1
MMP2
CCNA2
CUL3
DUSP10
NOS2
PPARGC1A
SERPINB5
CDK4
CXCL8
DUSP1
IL6
PRKCA
ARG1
ATF2
BCL2L1
CCND1
CSRP2
H2AFY
IFNG
KAT5
MAPK1
MAPK14
MAPK3
MAPK9
SELE
ACHE
ATF3
BCL2
CBFB
CREB1
DDIT3
DUSP5
DUSP8
FOS
HBG2
HES1
JUN
JUNB
MAPK8
PLAU
POU2F1
RUVBL2
SOCS3
TGFB2
TH
unknown CG mut
known HCG mut
wt

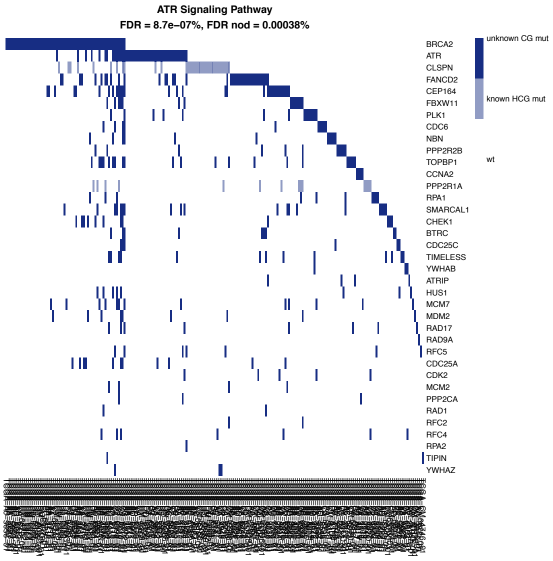
ATR Signaling Pathway
FDR = 8.7e−07%, FDR nod = 0.00038%
BRCA2
ATR
CLSPN
FANCD2
CEP164
FBXW11
PLK1
CDC6
NBN
PPP2R2B
TOPBP1
CCNA2
PPP2R1A
RPA1
SMARCAL1
CHEK1
BTRC
CDC25C
TIMELESS
YWHAB
ATRIP
HUS1
MCM7
MDM2
RAD17
RAD9A
RFC5
CDC25A
CDK2
MCM2
PPP2CA
RAD1
RFC2
RFC4
RPA2
TIPIN
YWHAZ
unknown CG mut
known HCG mut
wt

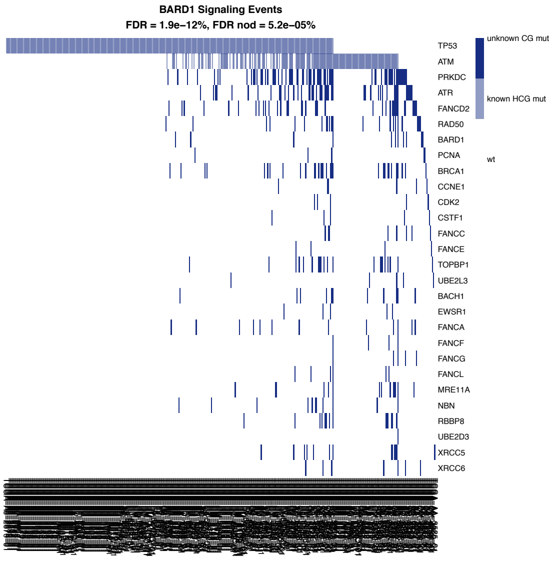
BARD1 Signaling Events
FDR = 1.9e−12%, FDR nod = 5.2e−05%
TP53
ATM
PRKDC
ATR
FANCD2
RAD50
BARD1
PCNA
BRCA1
CCNE1
CDK2
CSTF1
FANCC
FANCE
TOPBP1
UBE2L3
BACH1
EWSR1
FANCA
FANCF
FANCG
FANCL
MRE11A
NBN
RBBP8
UBE2D3
XRCC5
XRCC6
unknown CG mut
known HCG mut
wt

DNA Damage/Telomere Stress Induced Senescence

FDR = 1.9e−12%, FDR nod = 0.14%

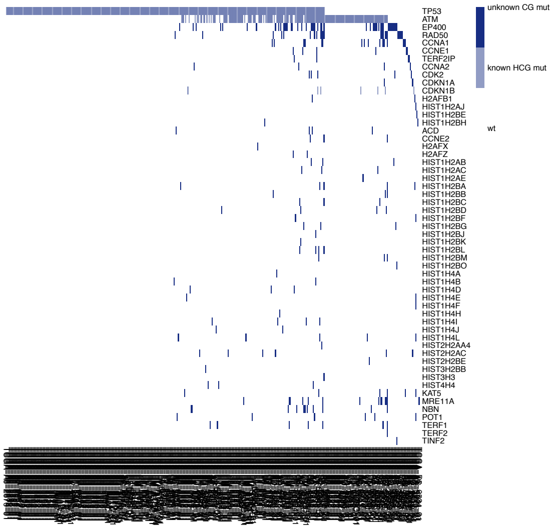


FOXA1 Transcription Factor Network

FDR = 2.3e−06%, FDR nod = 0.015%

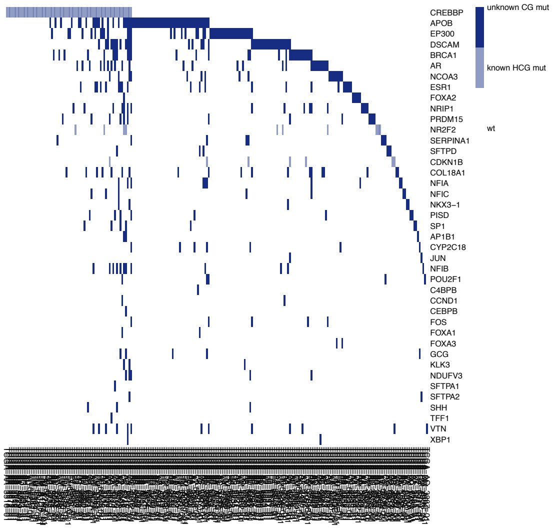


FOXM1 Transcription Factor Network

FDR = 1.9e−12%, FDR nod = 2.3e−11%

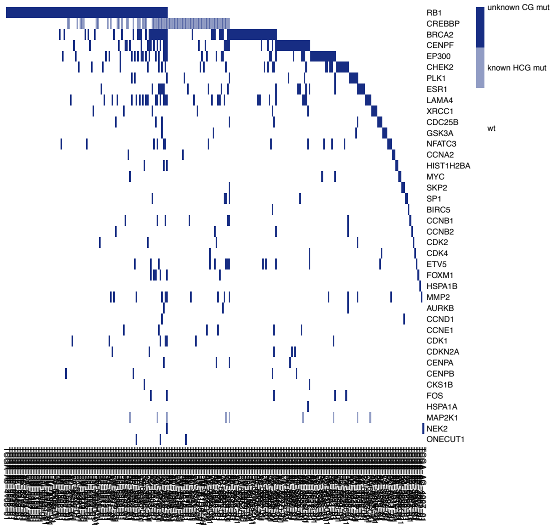


G2/M DNA Damage Checkpoint

Processing Of DNA Double Strand Break Ends

FDR = 1.9e−12%, FDR nod = 4.6%

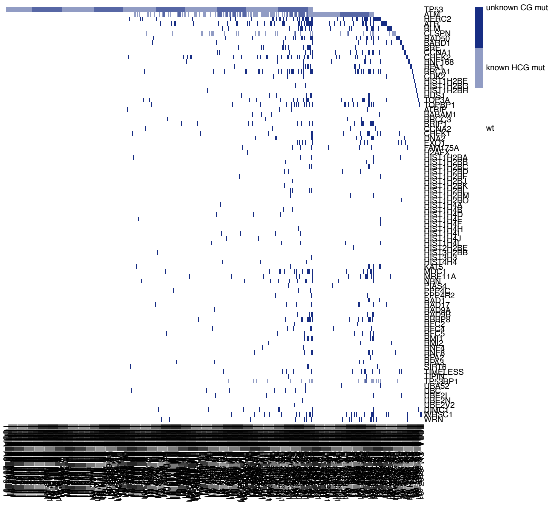


Gap Filling DNA Repair Synthesis And Ligation In GG NER

FDR = 0.002%, FDR nod = 0.0016%

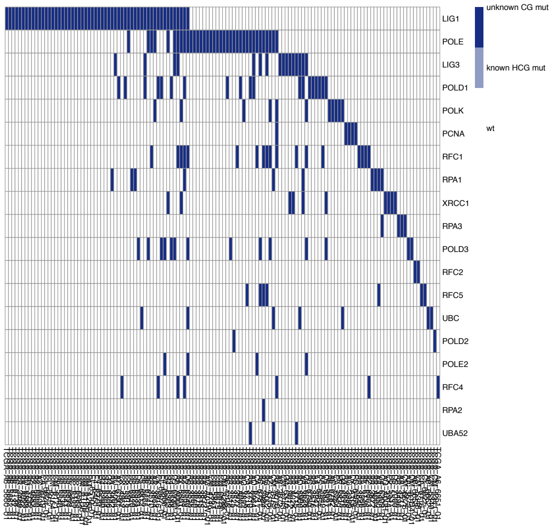


HDR Through Homologous Recombination (HRR)

FDR = 2.3e−12%, FDR nod = 0.51%

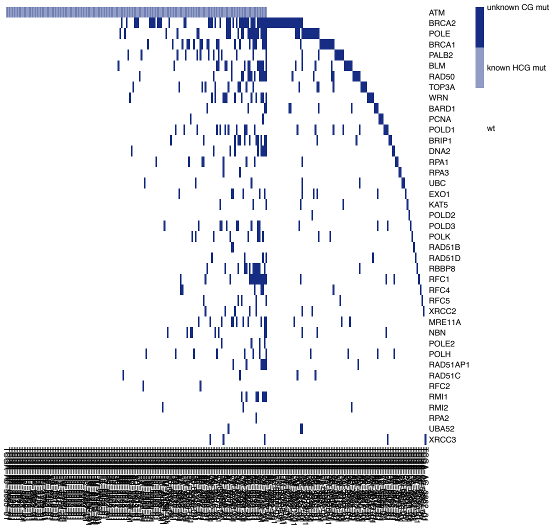

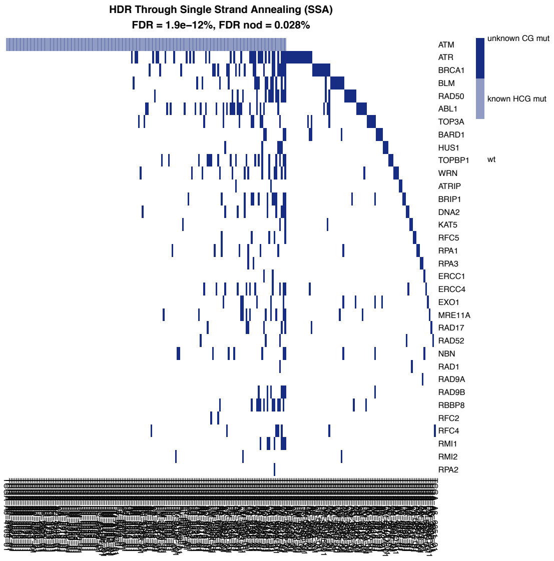
HDR Through Single Strand Annealing (SSA)
FDR = 1.9e−12%, FDR nod = 0.028%
ATM
ATR
BRCA1
BLM
RAD50
ABL1
TOP3A
BARD1
HUS1
TOPBP1
WRN
ATRIP
BRIP1
DNA2
KAT5
RFC5
RPA1
RPA3
ERCC1
ERCC4
EXO1
MRE11A
RAD17
RAD52
NBN
RAD1
RAD9A
RAD9B
RBBP8
RFC2
RFC4
RMI1
RMI2
RPA2
unknown CG mut
known HCG mut
wt

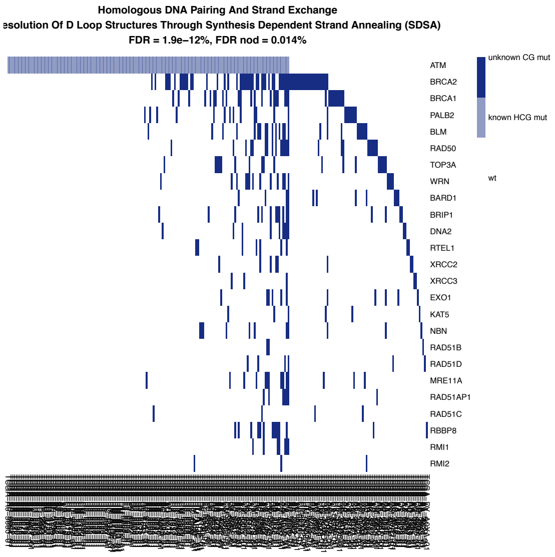
Homologous DNA Pairing And Strand Exchange
esolution Of D Loop Structures Through Synthesis Dependent Strand Annealing (SDSA)
FDR = 1.9e−12%, FDR nod = 0.014%
ATM
BRCA2
BRCA1
PALB2
BLM
RAD50
TOP3A
WRN
BARD1
BRIP1
DNA2
RTEL1
XRCC2
XRCC3
EXO1
KAT5
NBN
RAD51B
RAD51D
MRE11A
RAD51AP1
RAD51C
RBBP8
RMI1
RMI2
unknown CG mut
known HCG mut
wt

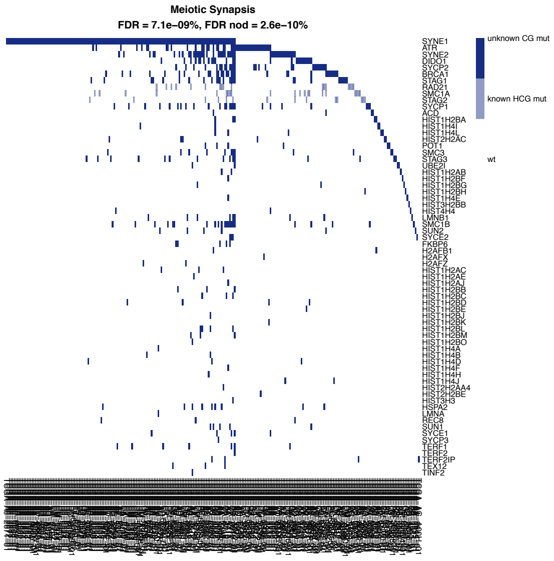
Meiotic Synapsis
FDR = 7.1e−09%, FDR nod = 2.6e−10%
SYNE1
ATR
SYNE2
DIDO1
SYCP2
BRCA1
STAG1
RAD21
SMC1A
STAG2
SYCP1
ACD
HIST1H2BA
HIST1H4I
HIST1H4L
HIST2H2AC
POT1
SMC3
STAG3
UBE2I
HIST1H2AB
HIST1H2BF
HIST1H2BG
HIST1H2BH
HIST1H4E
HIST3H2BB
HIST4H4
LMNB1
SMC1B
SUN2
SYCE2
FKBP6
H2AFB1
H2AFX
H2AFZ
HIST1H2AC
HIST1H2AE
HIST1H2AJ
HIST1H2BB
HIST1H2BC
HIST1H2BD
HIST1H2BE
HIST1H2BJ
HIST1H2BK
HIST1H2BL
HIST1H2BM
HIST1H2BO
HIST1H4A
HIST1H4B
HIST1H4D
HIST1H4F
HIST1H4H
HIST1H4J
HIST2H2AA4
HIST2H2BE
HIST3H3
HSPA2
LMNA
REC8
SUN1
SYCE1
SYCP3
TERF1
TERF2
TERF2IP
TEX12
TINF2
unknown CG mut
known HCG mut
wt

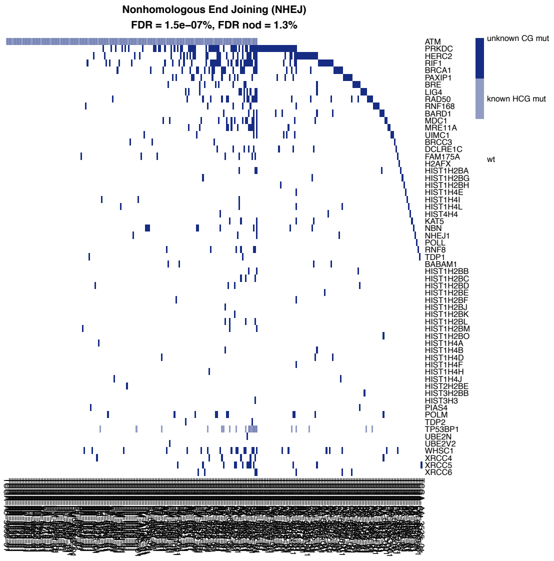
Nonhomologous End Joining (NHEJ)
FDR = 1.5e−07%, FDR nod = 1.3%
ATM
PRKDC
HERC2
RIF1
BRCA1
PAXIP1
BRE
LIG4
RAD50
RNF168
BARD1
MDC1
MRE11A
UIMC1
BRCC3
DCLRE1C
FAM175A
H2AFX
HIST1H2BA
HIST1H2BG
HIST1H2BH
HIST1H4E
HIST1H4I
HIST1H4L
HIST4H4
KAT5
NBN
NHEJ1
POLL
RNF8
TDP1
BABAM1
HIST1H2BB
HIST1H2BC
HIST1H2BD
HIST1H2BE
HIST1H2BF
HIST1H2BJ
HIST1H2BK
HIST1H2BL
HIST1H2BM
HIST1H2BO
HIST1H4A
HIST1H4B
HIST1H4D
HIST1H4F
HIST1H4H
HIST1H4J
HIST2H2BE
HIST3H2BB
HIST3H3
PIAS4
POLM
TDP2
TP53BP1
UBE2N
UBE2V2
WHSC1
XRCC4
XRCC5
XRCC6
unknown CG mut
known HCG mut
wt

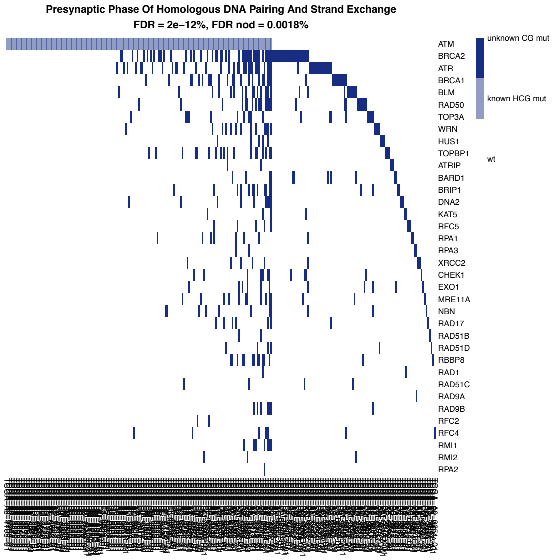
Presynaptic Phase Of Homologous DNA Pairing And Strand Exchange
FDR = 2e−12%, FDR nod = 0.0018%
ATM
BRCA2
ATR
BRCA1
BLM
RAD50
TOP3A
WRN
HUS1
TOPBP1
ATRIP
BARD1
BRIP1
DNA2
KAT5
RFC5
RPA1
RPA3
XRCC2
CHEK1
EXO1
MRE11A
NBN
RAD17
RAD51B
RAD51D
RBBP8
RAD1
RAD51C
RAD9A
RAD9B
RFC2
RFC4
RMI1
RMI2
RPA2
unknown CG mut
known HCG mut
wt

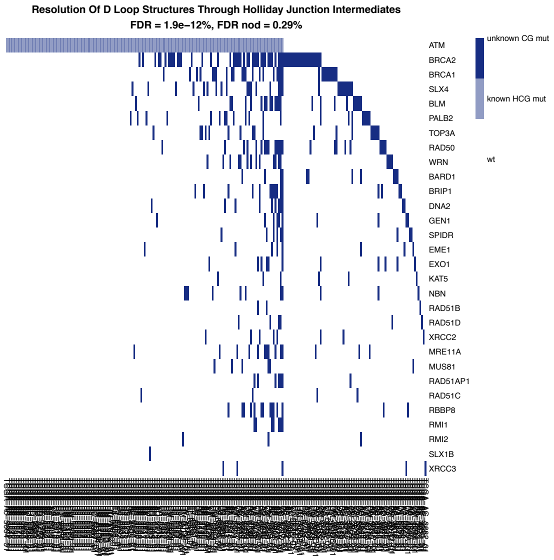
Resolution Of D Loop Structures Through Holliday Junction Intermediates
FDR = 1.9e−12%, FDR nod = 0.29%
ATM
BRCA2
BRCA1
SLX4
BLM
PALB2
TOP3A
RAD50
WRN
BARD1
BRIP1
DNA2
GEN1
SPIDR
EME1
EXO1
KAT5
NBN
RAD51B
RAD51D
XRCC2
MRE11A
MUS81
RAD51AP1
RAD51C
RBBP8
RMI1
RMI2
SLX1B
XRCC3
unknown CG mut
known HCG mut
wt

[CGB Protein Results In Increased Activity Of LHCGR Protein]
de Results In Decreased Activity Of PTGS2 Protein] Inhibits The Reaction [CGB Protein Results In Increased Chen
)(3, 2 F)quinazoline 1,3 Diamine Binds To F2R Protein] Promotes The Reaction [CGB Protein Results In Increased
ne Inhibits The Reaction [CGB Protein Results In Increased Chemical Synthesis Of Androgens]
itrazine Inhibits The Reaction [CGB Protein Results In Increased Secretion Of Cyclic AMP]
forsin Promotes The Reaction [CGB Protein Results In Increased Abundance Of Cyclic AMP]
Reaction [[Gonadotropins, Equine Co Treated With CGB Protein] Results In Increased Abundance Of Progesteron
ibits The Reaction [[CGB Protein Co Treated With Colforsin] Results In Increased Abundance Of Cyclic AMP]
Protein Promotes The Reaction [CGB Protein Results In Increased Abundance Of Progesterone]
ole Metabolite Inhibits The Reaction [CGB Protein Results In Increased Abundance Of Testosterone]
se Chloride Inhibits The Reaction [CGB Protein Results In Increased Expression Of STAR Protein]
terone Promotes The Reaction [CGB Protein Results In Increased Expression Of VEGFA MRNA]
one Enanthate Inhibits The Reaction [CGB Protein Results In Increased Abundance Of Androgens]
FDR = 0.0063%, FDR nod = 0.012%

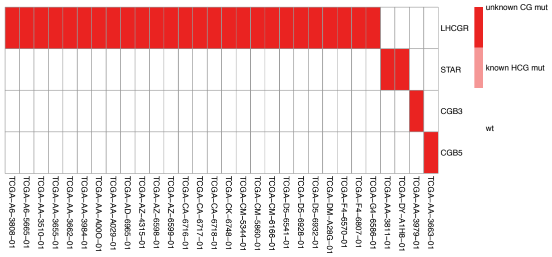


HIF 2 Alpha Transcription Factor Network
FDR = 2e−04%, FDR nod = 0.2%

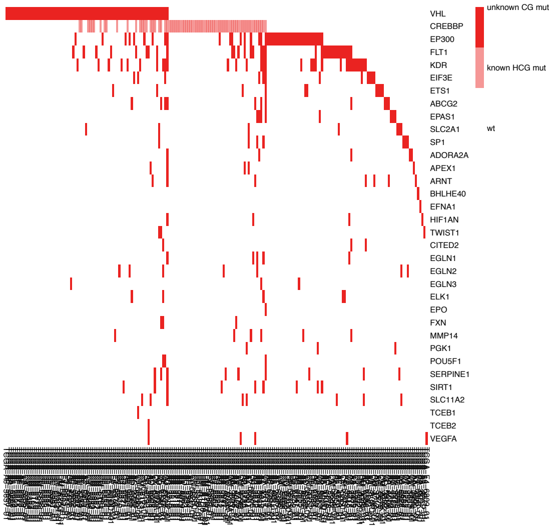


Hypoxia Response Via HIF Activation
FDR = 1.9e−12%, FDR nod = 3.7e−09%

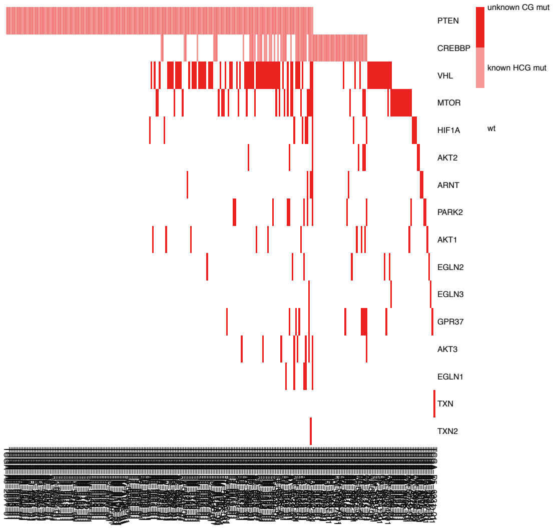


Hypoxic And Oxygen Homeostasis Regulation Of HIF 1 Alpha
FDR = 1.9e−12%, FDR nod = 0.018%

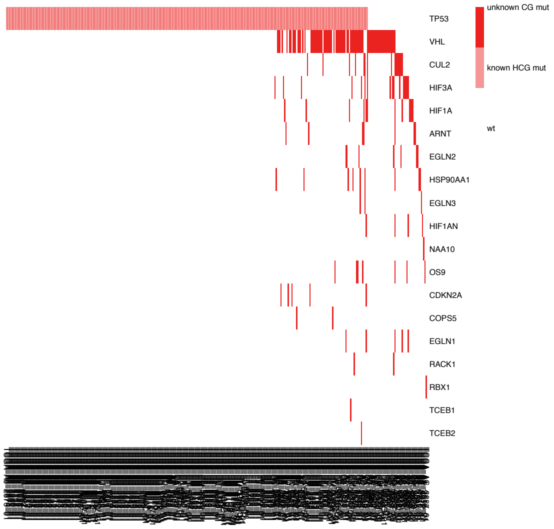


JNK Cascade ( EGF Signaling Pathway Diagram )
JNK Cascade ( PDGF Signaling Pathway )
FDR = 0.00054%, FDR nod = 0.0011%

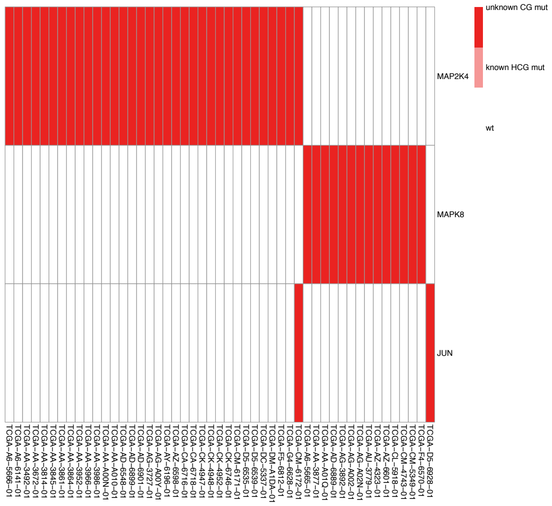


Neurophilin Interactions With VEGF And VEGFR
FDR = 0.68%, FDR nod = 0.78%

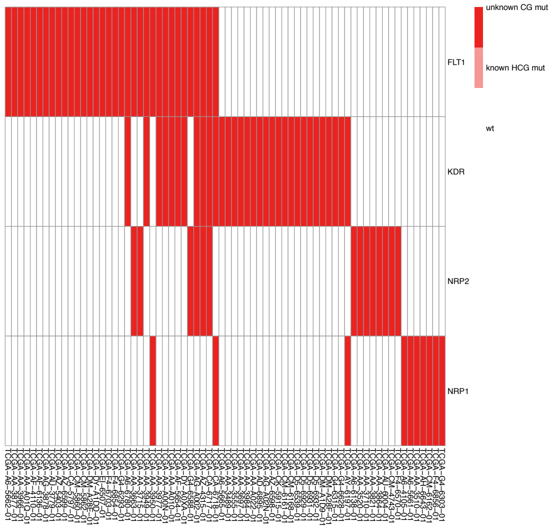

**Oxygen Dependent Proline Hydroxylation Of Hypoxia Inducible Factor Alpha**
**FDR = 0.01%, FDR nod = 0.0092%**

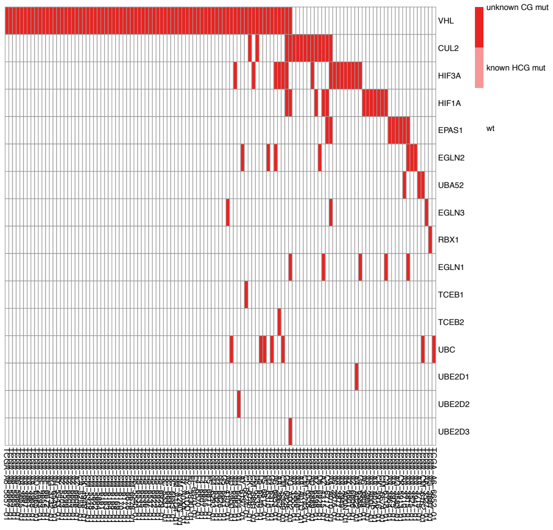


**PDGF Pathway**
**FDR = 8.9e–10%, FDR nod = 0.0023%**

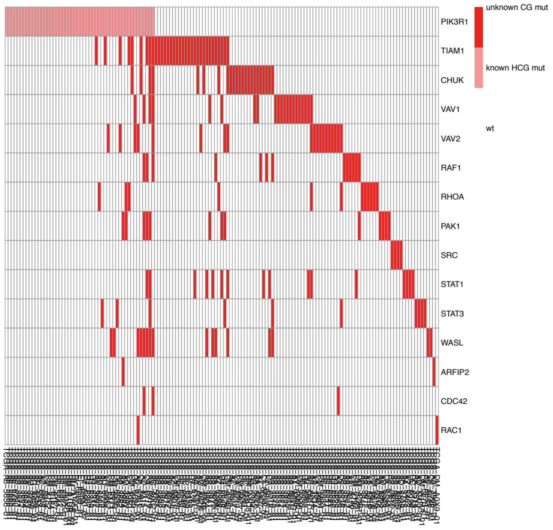


**PDGF Receptor Signaling Network**
**FDR = 7e–08%, FDR nod = 1.2e–07%**

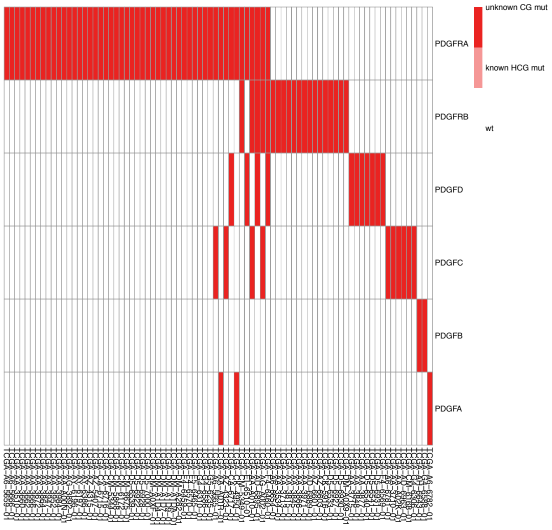


**PDGF Signaling Pathway ( PDGF Signaling Pathway )**
**FDR = 1.9e–12%, FDR nod = 0.00016%**

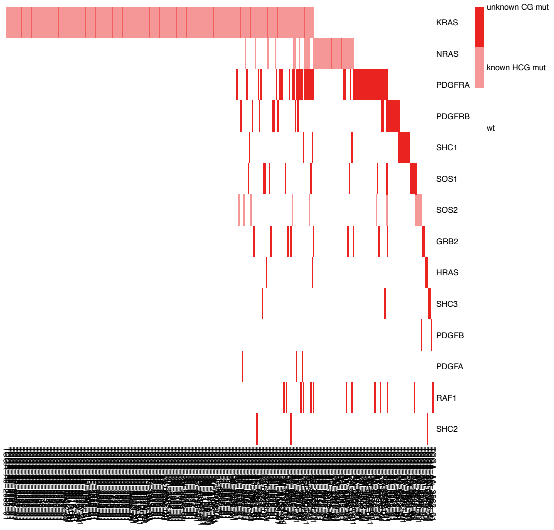


**PDGF Signaling Pathway (Mammal) ( PDGF Signaling Pathway )**
**FDR = 1.9e–12%, FDR nod = 0.16%**

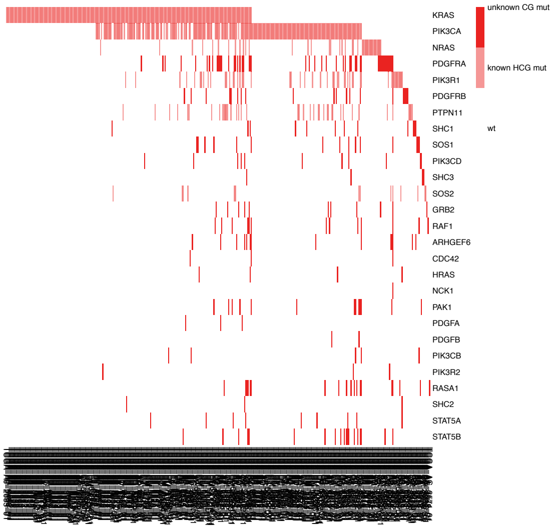


**PDGFR Alpha Signaling Pathway**
**FDR = 1.9e–12%, FDR nod = 0.0023%**

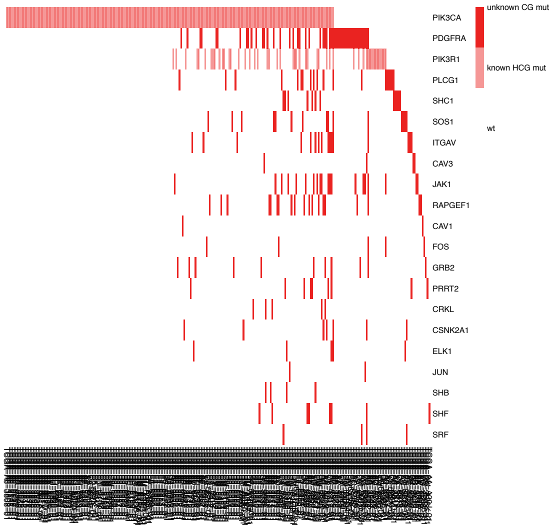

**PTK6 Promotes HIF1A Stabilization**
FDR = 2e−12%, FDR nod = 2.6e−08%

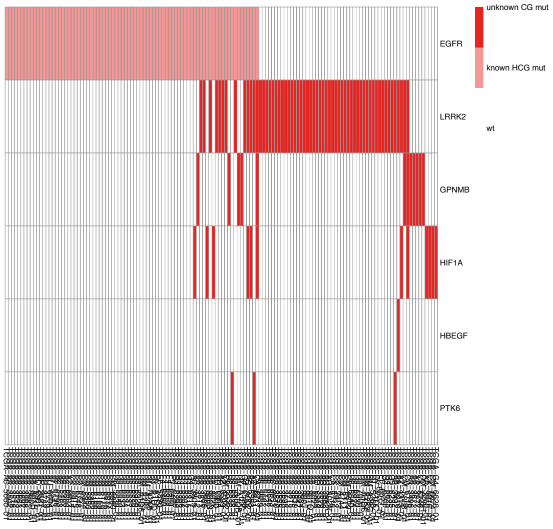


**Regulation Of Gene Expression By Hypoxia Inducible Factor**
FDR = 4.9e−09%, FDR nod = 1.8%

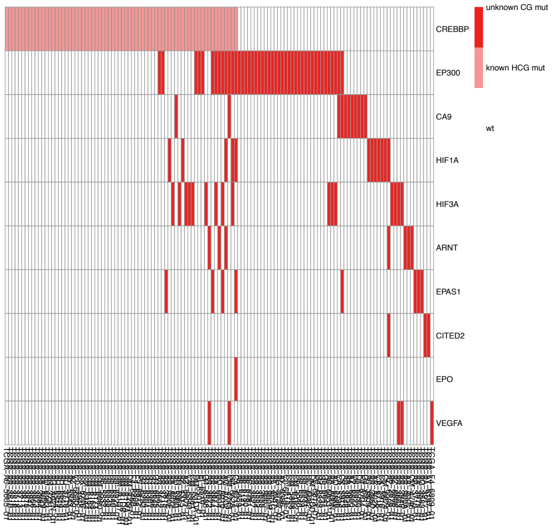


**Signaling By PDGF**
FDR = 1.9e−12%, FDR nod = 2.3e−11%

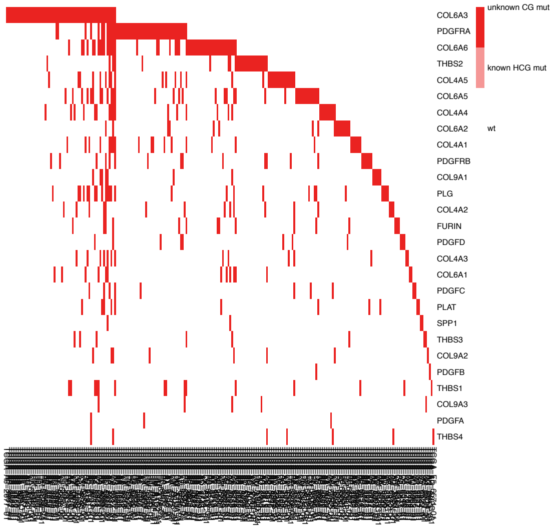


**Tie2 Signaling**
FDR = 1.9e−12%, FDR nod = 0.074%

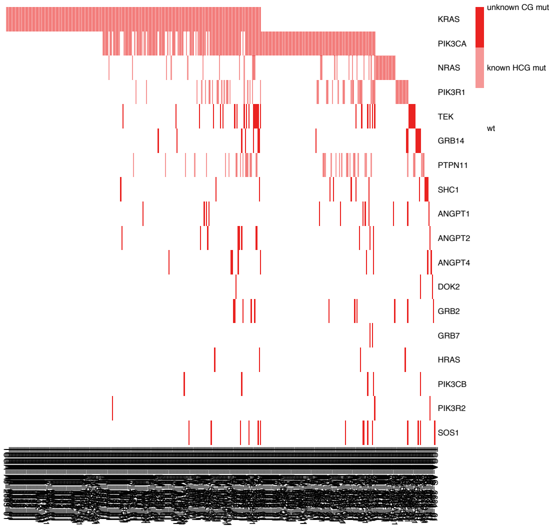


**VEGF Signaling Pathway**
FDR = 1.9e−12%, FDR nod = 0.053%

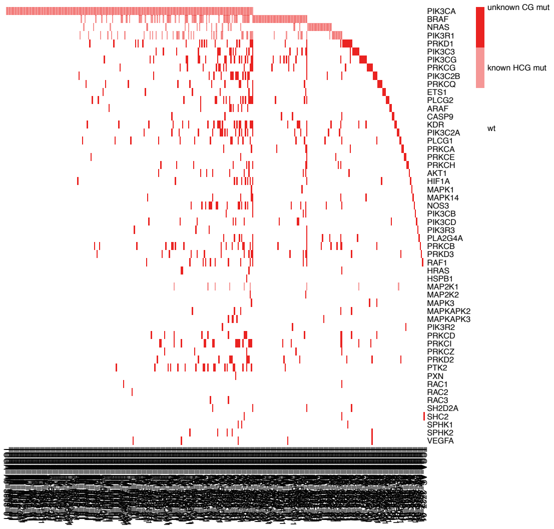


**VEGFR3 Signaling In Lymphatic Endothelium**
FDR = 1.9e−12%, FDR nod = 1.2%

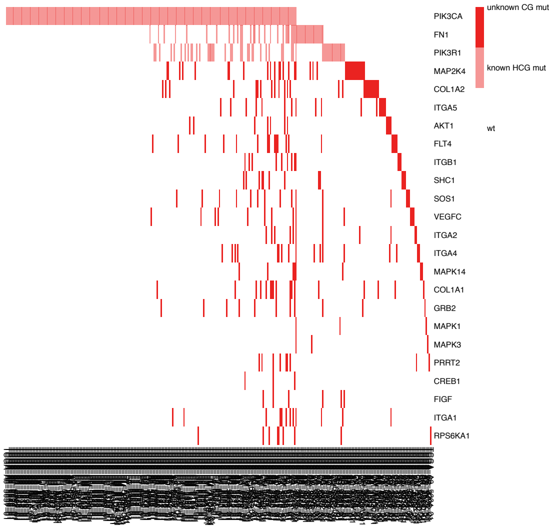

**Apoptotic Cleavage Of Cell Adhesion Proteins**
**FDR = 1.9e−12%, FDR nod = 2.3e−11%**

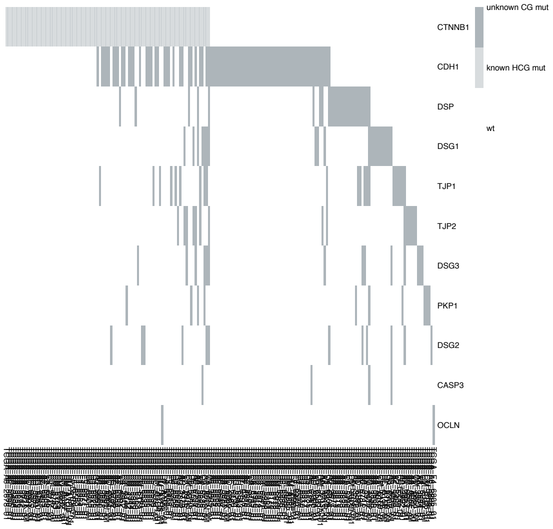


**FAS Pathway And Stress Induction Of HSP Regulation**
**FDR = 4e−06%, FDR nod = 5.5e−06%**

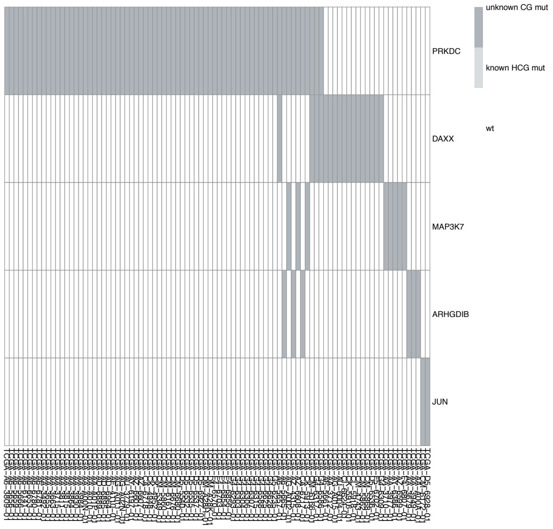


**FAS Signaling Pathway**
**FDR = 2.4e−12%, FDR nod = 3.7e−06%**

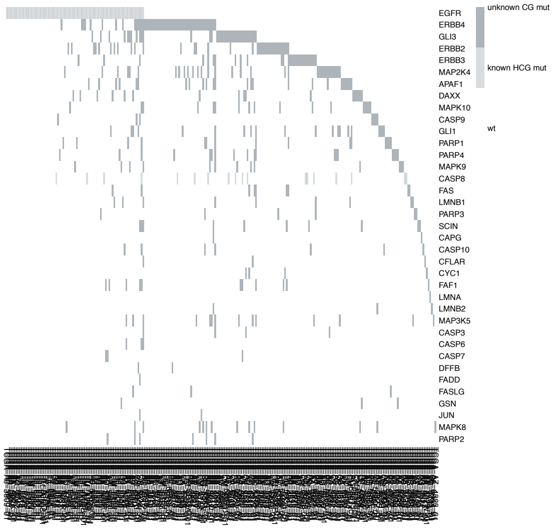


**G2/M DNA Damage Checkpoint**
**Processing Of DNA Double Strand Break Ends**
**FDR = 1.9e−12%, FDR nod = 4.6%**

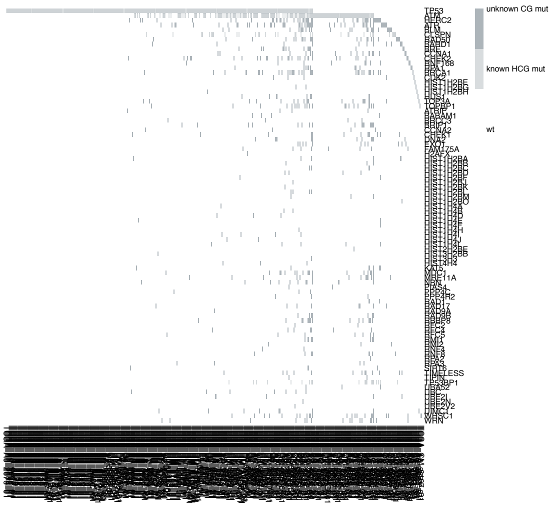


**Homologous DNA Pairing And Strand Exchange**
**esolution Of D Loop Structures Through Synthesis Dependent Strand Annealing (SDSA)**
**FDR = 1.9e−12%, FDR nod = 0.014%**

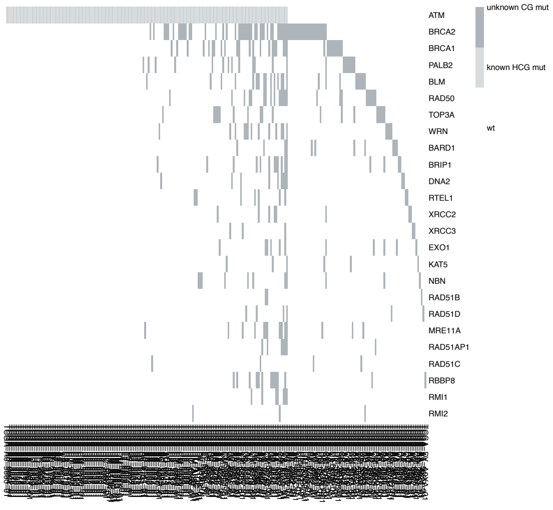


**Presynaptic Phase Of Homologous DNA Pairing And Strand Exchange**
**FDR = 2e−12%, FDR nod = 0.0018%**

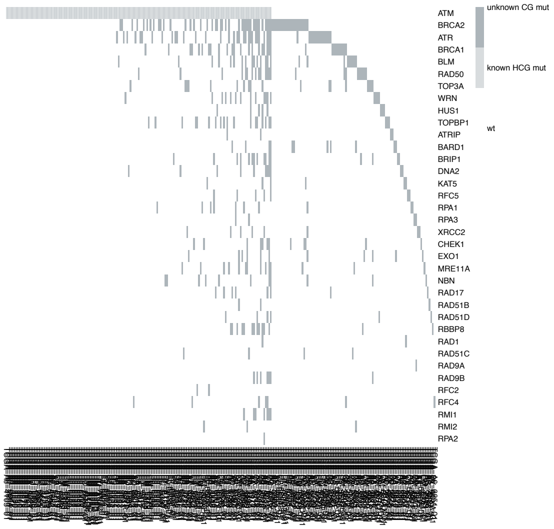

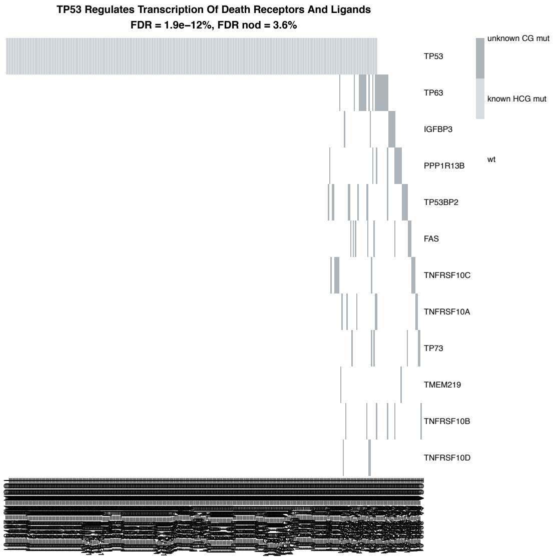
TP53 Regulates Transcription Of Death Receptors And Ligands
FDR = 1.9e−12%, FDR nod = 3.6%
TP53
TP63
IGFBP3
PPP1R13B
TP53BP2
FAS
TNFRSF10C
TNFRSF10A
TP73
TMEM219
TNFRSF10B
TNFRSF10D
unknown CG mut
known HCG mut
wt

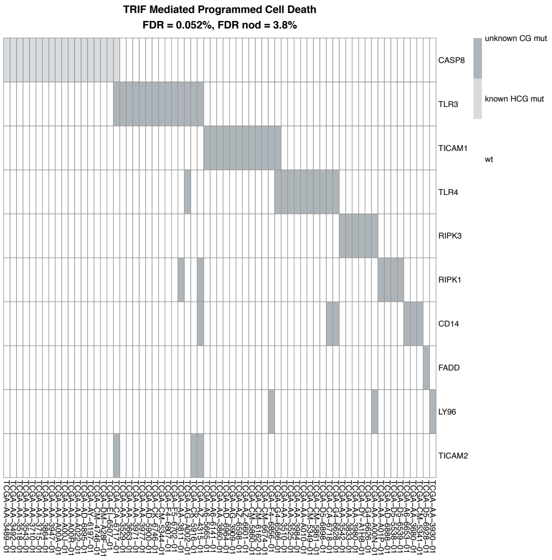
TRIF Mediated Programmed Cell Death
FDR = 0.052%, FDR nod = 3.8%
CASP8
TLR3
TICAM1
TLR4
RIPK3
RIPK1
CD14
FADD
LY96
TICAM2
unknown CG mut
known HCG mut
wt

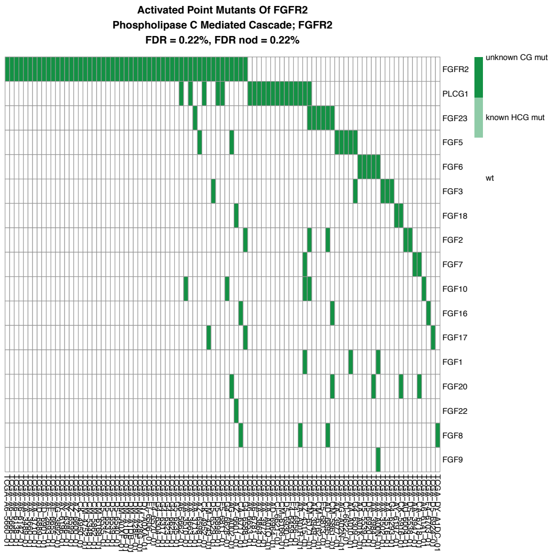
Activated Point Mutants Of FGFR2
Phospholipase C Mediated Cascade; FGFR2
FDR = 0.22%, FDR nod = 0.22%
FGFR2
PLCG1
FGF23
FGF5
FGF6
FGF3
FGF18
FGF2
FGF7
FGF10
FGF16
FGF17
FGF1
FGF20
FGF22
FGF8
FGF9
unknown CG mut
known HCG mut
wt

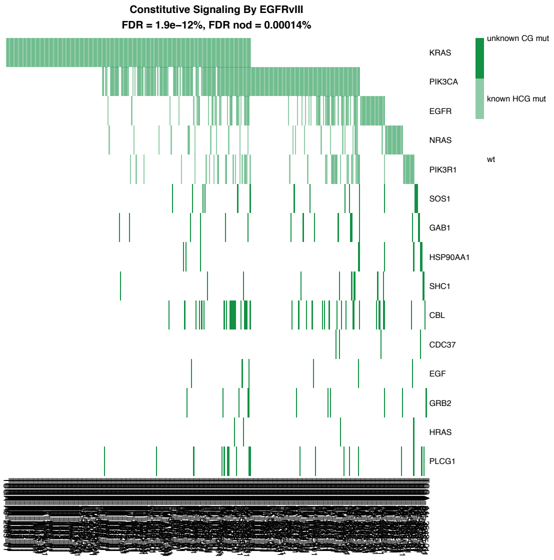
Constitutive Signaling By EGFRvIII
FDR = 1.9e−12%, FDR nod = 0.00014%
KRAS
PIK3CA
EGFR
NRAS
PIK3R1
SOS1
GAB1
HSP90AA1
SHC1
CBL
CDC37
EGF
GRB2
HRAS
PLCG1
unknown CG mut
known HCG mut
wt

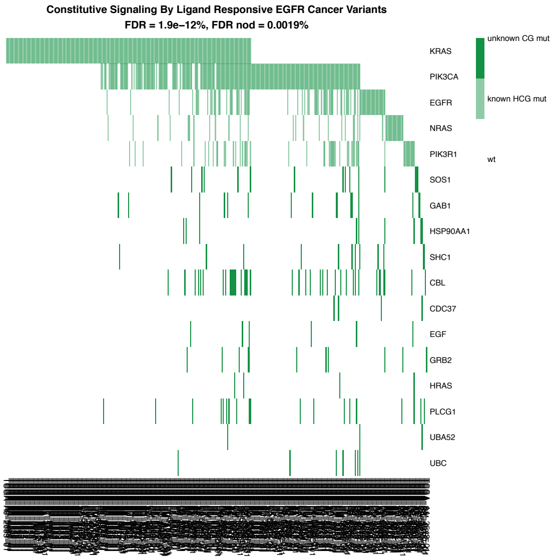
Constitutive Signaling By Ligand Responsive EGFR Cancer Variants
FDR = 1.9e−12%, FDR nod = 0.0019%
KRAS
PIK3CA
EGFR
NRAS
PIK3R1
SOS1
GAB1
HSP90AA1
SHC1
CBL
CDC37
EGF
GRB2
HRAS
PLCG1
UBA52
UBC
unknown CG mut
known HCG mut
wt

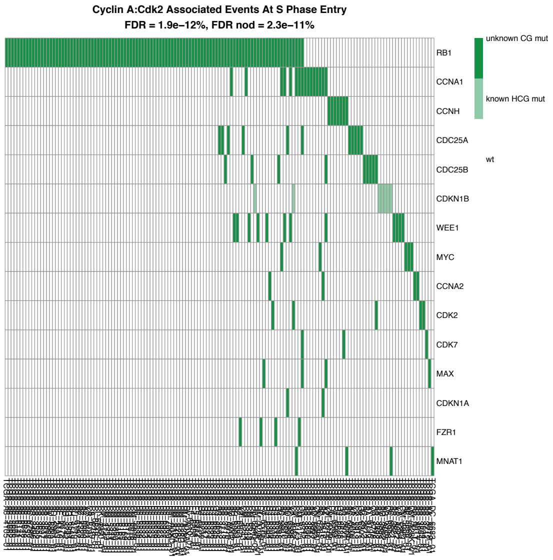
Cyclin A:Cdk2 Associated Events At S Phase Entry
FDR = 1.9e−12%, FDR nod = 2.3e−11%
RB1
CCNA1
CCNH
CDC25A
CDC25B
CDKN1B
WEE1
MYC
CCNA2
CDK2
CDK7
MAX
CDKN1A
FZR1
MNAT1
unknown CG mut
known HCG mut
wt

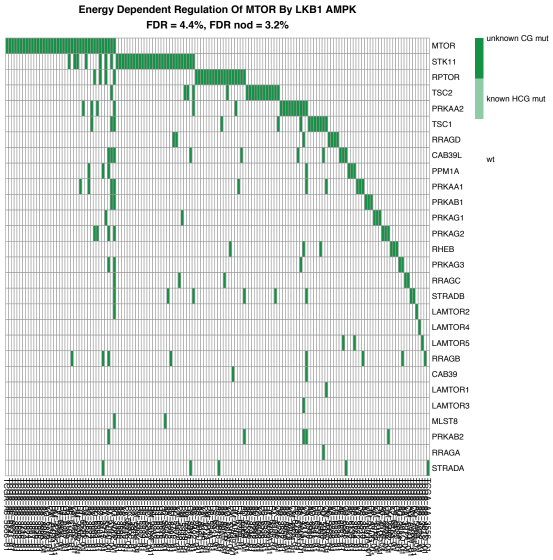
Energy Dependent Regulation Of MTOR By LKB1 AMPK
FDR = 4.4%, FDR nod = 3.2%
MTOR
STK11
RPTOR
TSC2
PRKAA2
TSC1
RRAGD
CAB39L
PPM1A
PRKAA1
PRKAB1
PRKAG1
PRKAG2
RHEB
PRKAG3
RRAGC
STRADB
LAMTOR2
LAMTOR4
LAMTOR5
RRAGB
CAB39
LAMTOR1
LAMTOR3
MLST8
PRKAB2
RRAGA
STRADA
unknown CG mut
known HCG mut
wt

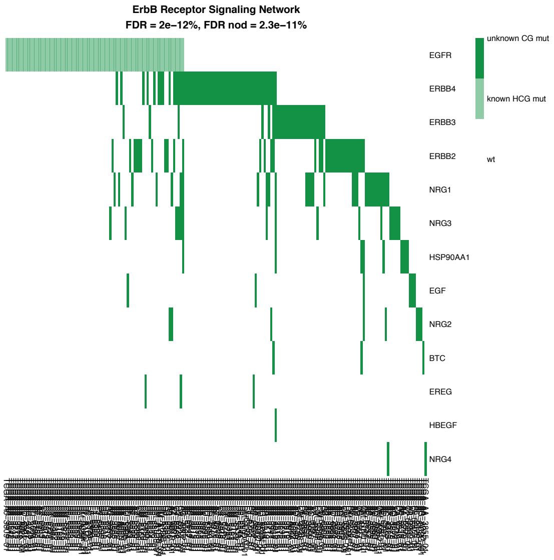
ErbB Receptor Signaling Network
FDR = 2e−12%, FDR nod = 2.3e−11%
EGFR
ERBB4
ERBB3
ERBB2
NRG1
NRG3
HSP90AA1
EGF
NRG2
BTC
EREG
HBEGF
NRG4
unknown CG mut
known HCG mut
wt

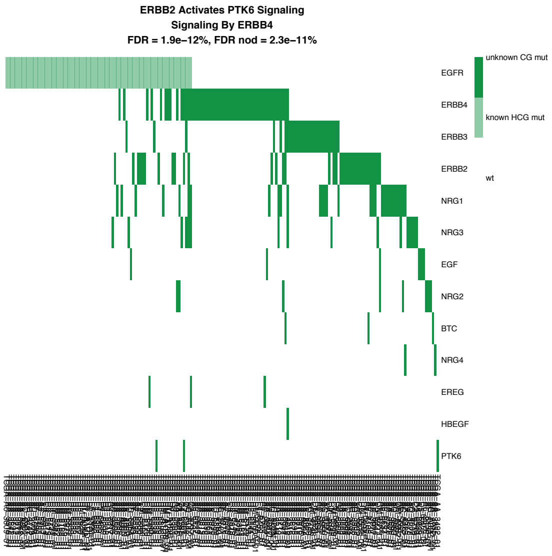
ERBB2 Activates PTK6 Signaling
Signaling By ERBB4
FDR = 1.9e−12%, FDR nod = 2.3e−11%
EGFR
ERBB4
ERBB3
ERBB2
NRG1
NRG3
EGF
NRG2
BTC
NRG4
EREG
HBEGF
PTK6
unknown CG mut
known HCG mut
wt

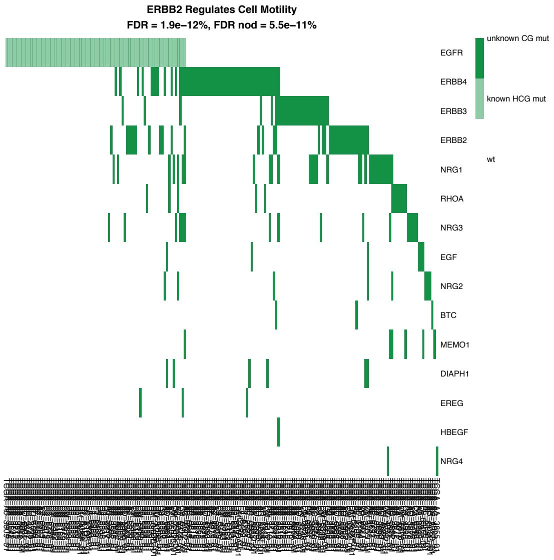
ERBB2 Regulates Cell Motility
FDR = 1.9e−12%, FDR nod = 5.5e−11%
EGFR
ERBB4
ERBB3
ERBB2
NRG1
RHOA
NRG3
EGF
NRG2
BTC
MEMO1
DIAPH1
EREG
HBEGF
NRG4
unknown CG mut
known HCG mut
wt

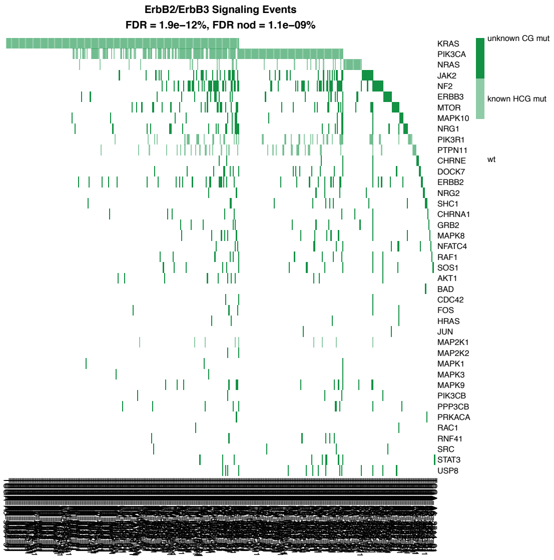
ErbB2/ErbB3 Signaling Events
FDR = 1.9e−12%, FDR nod = 1.1e−09%
KRAS
PIK3CA
NRAS
JAK2
NF2
ERBB3
MTOR
MAPK10
NRG1
PIK3R1
PTPN11
CHRNE
DOCK7
ERBB2
NRG2
SHC1
CHRNA1
GRB2
MAPK8
NFATC4
RAF1
SOS1
AKT1
BAD
CDC42
FOS
HRAS
JUN
MAP2K1
MAP2K2
MAPK1
MAPK3
MAPK9
PIK3CB
PPP3CB
PRKACA
RAC1
RNF41
SRC
STAT3
USP8
unknown CG mut
known HCG mut
wt

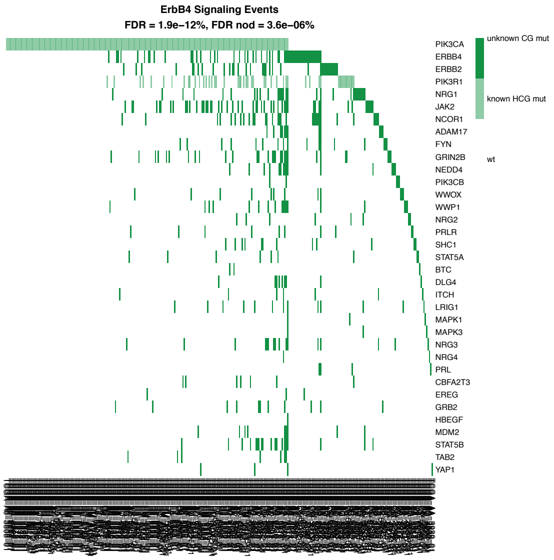
ErbB4 Signaling Events
FDR = 1.9e−12%, FDR nod = 3.6e−06%
PIK3CA
ERBB4
ERBB2
PIK3R1
NRG1
JAK2
NCOR1
ADAM17
FYN
GRIN2B
NEDD4
PIK3CB
WWOX
WWP1
NRG2
PRLR
SHC1
STAT5A
BTC
DLG4
ITCH
LRIG1
MAPK1
MAPK3
NRG3
NRG4
PRL
CBFA2T3
EREG
GRB2
HBEGF
MDM2
STAT5B
TAB2
YAP1
unknown CG mut
known HCG mut
wt

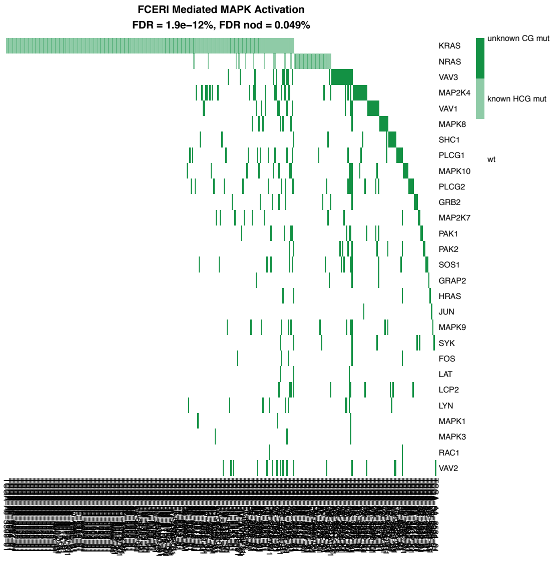
FCERI Mediated MAPK Activation
FDR = 1.9e−12%, FDR nod = 0.049%
KRAS
NRAS
VAV3
MAP2K4
VAV1
MAPK8
SHC1
PLCG1
MAPK10
PLCG2
GRB2
MAP2K7
PAK1
PAK2
SOS1
GRAP2
HRAS
JUN
MAPK9
SYK
FOS
LAT
LCP2
LYN
MAPK1
MAPK3
RAC1
VAV2
unknown CG mut
known HCG mut
wt

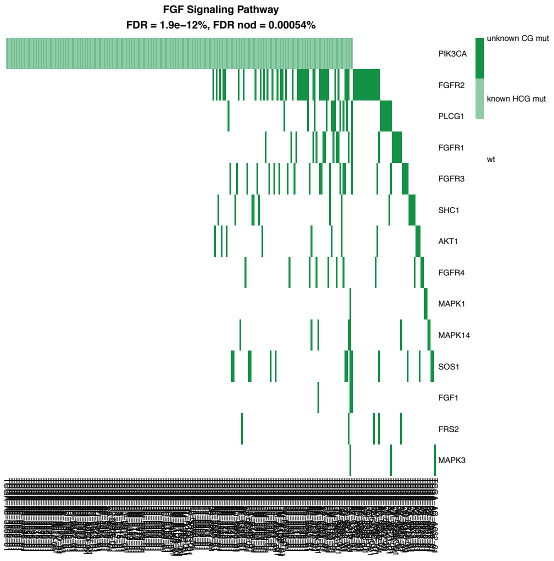
FGF Signaling Pathway
FDR = 1.9e−12%, FDR nod = 0.00054%
PIK3CA
FGFR2
PLCG1
FGFR1
FGFR3
SHC1
AKT1
FGFR4
MAPK1
MAPK14
SOS1
FGF1
FRS2
MAPK3
unknown CG mut
known HCG mut
wt

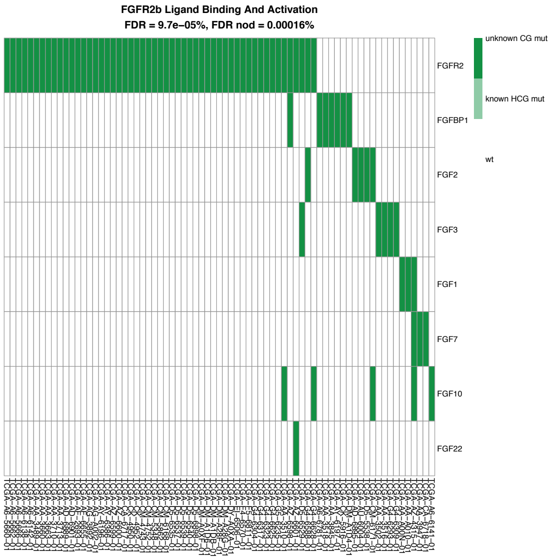
FGFR2b Ligand Binding And Activation
FDR = 9.7e−05%, FDR nod = 0.00016%
FGFR2
FGFBP1
FGF2
FGF3
FGF1
FGF7
FGF10
FGF22
unknown CG mut
known HCG mut
wt

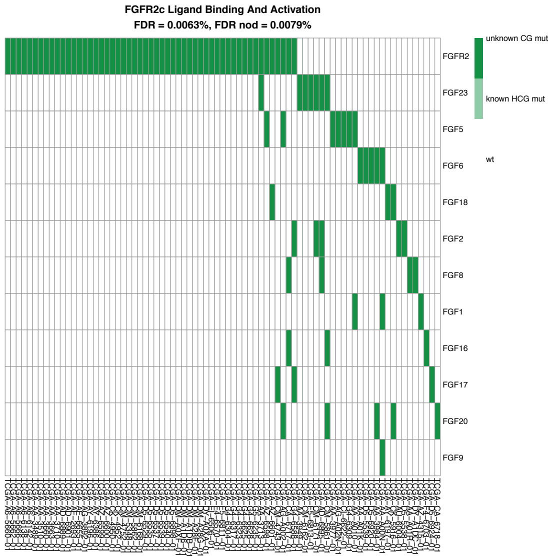
FGFR2c Ligand Binding And Activation
FDR = 0.0063%, FDR nod = 0.0079%
FGFR2
FGF23
FGF5
FGF6
FGF18
FGF2
FGF8
FGF1
FGF16
FGF17
FGF20
FGF9
unknown CG mut
known HCG mut
wt

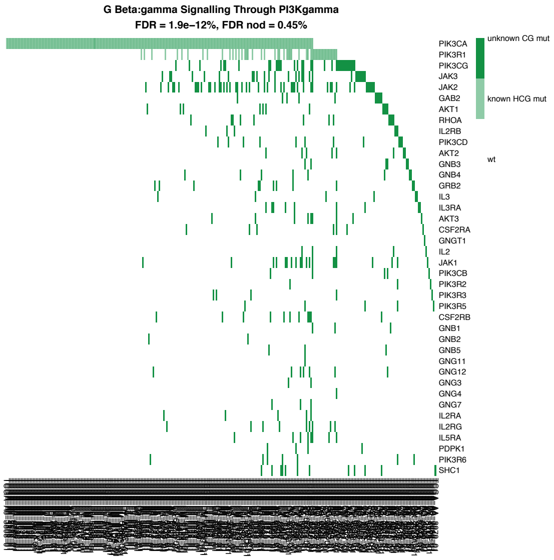
G Beta:gamma Signalling Through PI3Kgamma
FDR = 1.9e−12%, FDR nod = 0.45%
PIK3CA
PIK3R1
PIK3CG
JAK3
JAK2
GAB2
AKT1
RHOA
IL2RB
PIK3CD
AKT2
GNB3
GNB4
GRB2
IL3
IL3RA
AKT3
CSF2RA
GNGT1
IL2
JAK1
PIK3CB
PIK3R2
PIK3R3
PIK3R5
CSF2RB
GNB1
GNB2
GNB5
GNG11
GNG12
GNG3
GNG4
GNG7
IL2RA
IL2RG
IL5RA
PDPK1
PIK3R6
SHC1
unknown CG mut
known HCG mut
wt

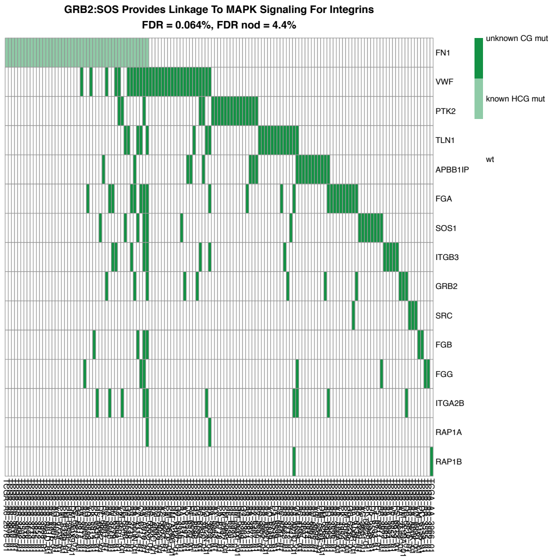
GRB2:SOS Provides Linkage To MAPK Signaling For Integrins
FDR = 0.064%, FDR nod = 4.4%
FN1
VWF
PTK2
TLN1
APBB1IP
FGA
SOS1
ITGB3
GRB2
SRC
FGB
FGG
ITGA2B
RAP1A
RAP1B
unknown CG mut
known HCG mut
wt

owth Hormone Signaling Pathway(JAK2 STAT5) ( Growth Hormone Signaling(JAK2 STAT5) )
FDR = 1.9e−12%, FDR nod = 2e−05%

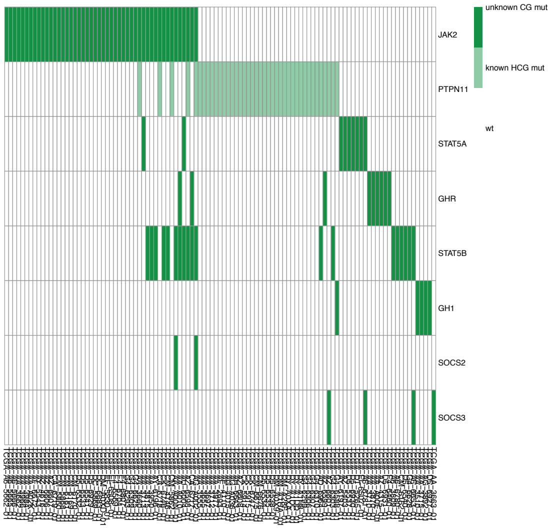


thway (through Glutamate, G Alpha Q And PLC Beta) ( GPCR GroupI Metabotropic Glutamate Receptor Signaling I
FDR = 1.9e−12%, FDR nod = 0.00018%

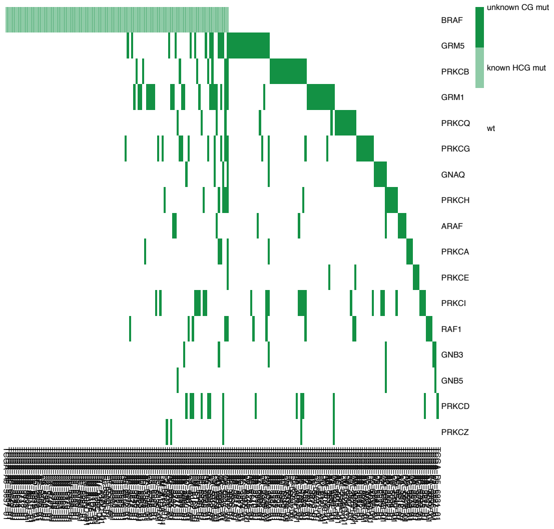


IL2 Signaling Events Mediated By PI3K
FDR = 2e−12%, FDR nod = 4.8%

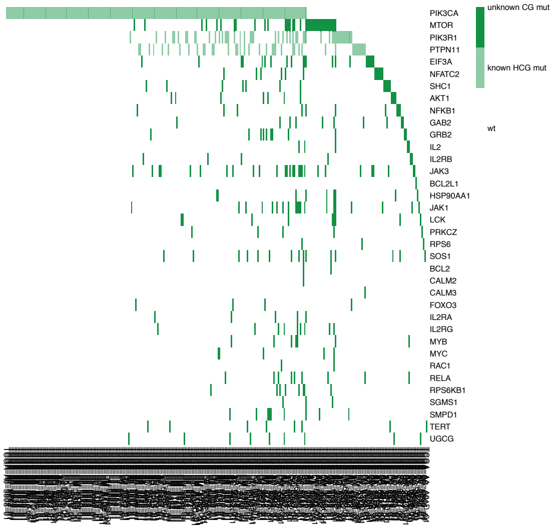


JAK/STAT Signaling Pathway
FDR = 0.06%, FDR nod = 0.048%

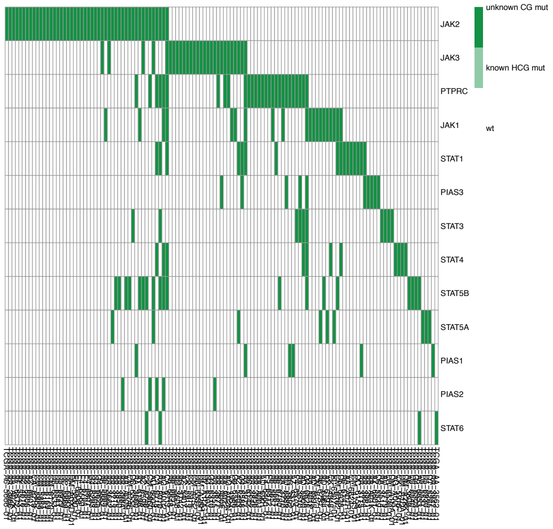


JNK Cascade ( EGF Signaling Pathway Diagram )
JNK Cascade ( PDGF Signaling Pathway )
FDR = 0.00054%, FDR nod = 0.0011%

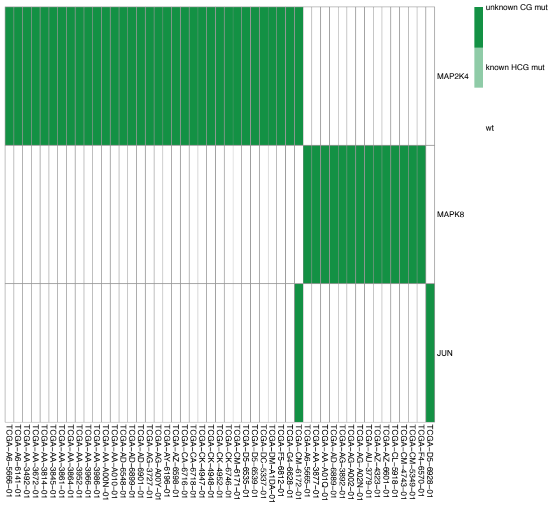


MAP3K8 (TPL2) Dependent MAPK1/3 Activation
FDR = 0.25%, FDR nod = 3.3%

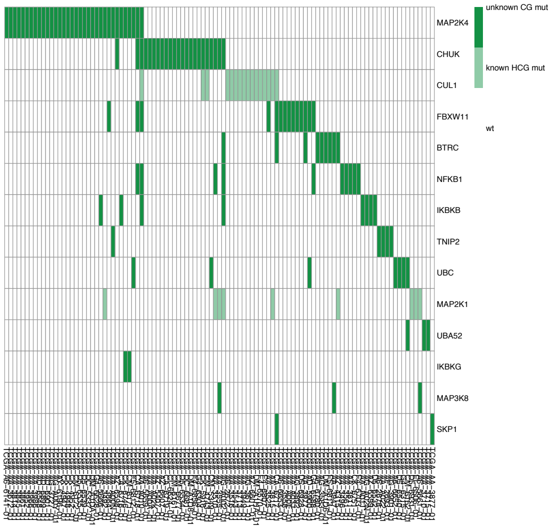

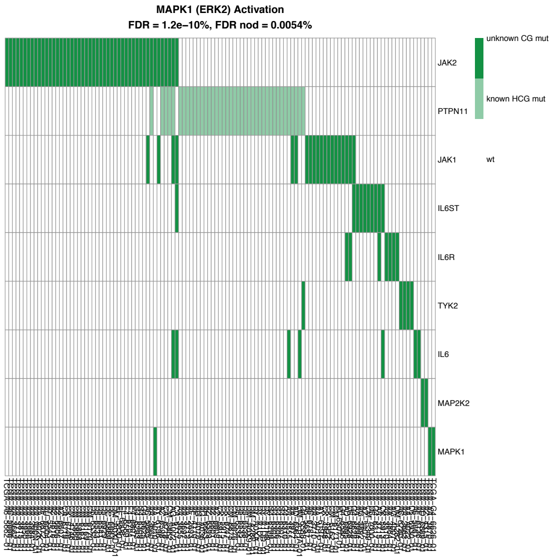
MAPK1 (ERK2) Activation
FDR = 1.2e−10%, FDR nod = 0.0054%
JAK2
PTPN11
JAK1
IL6ST
IL6R
TYK2
IL6
MAP2K2
MAPK1
unknown CG mut
known HCG mut
wt

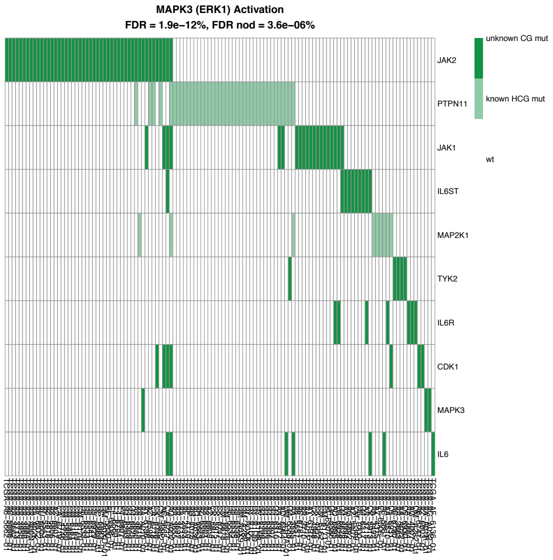
MAPK3 (ERK1) Activation
FDR = 1.9e−12%, FDR nod = 3.6e−06%
JAK2
PTPN11
JAK1
IL6ST
MAP2K1
TYK2
IL6R
CDK1
MAPK3
IL6
unknown CG mut
known HCG mut
wt

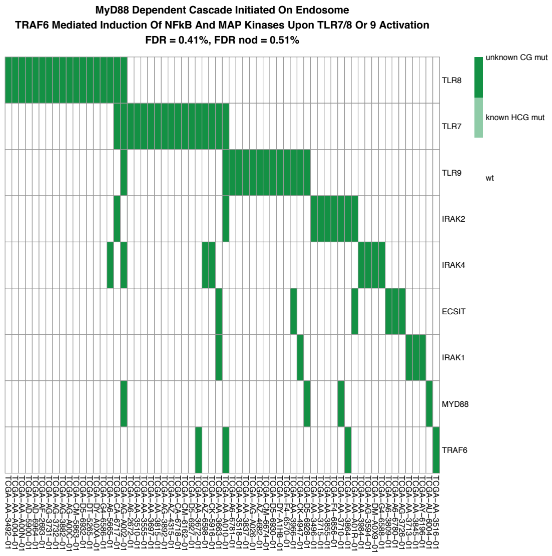
MyD88 Dependent Cascade Initiated On Endosome
TRAF6 Mediated Induction Of NFkB And MAP Kinases Upon TLR7/8 Or 9 Activation
FDR = 0.41%, FDR nod = 0.51%
TLR8
TLR7
TLR9
IRAK2
IRAK4
ECSIT
IRAK1
MYD88
TRAF6
unknown CG mut
known HCG mut
wt

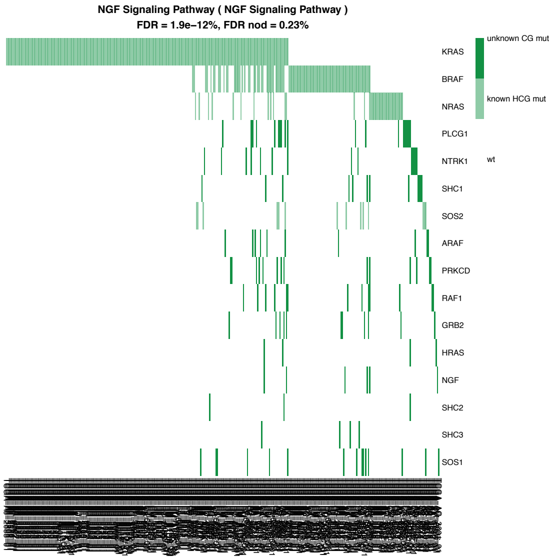
NGF Signaling Pathway ( NGF Signaling Pathway )
FDR = 1.9e−12%, FDR nod = 0.23%
KRAS
BRAF
NRAS
PLCG1
NTRK1
SHC1
SOS2
ARAF
PRKCD
RAF1
GRB2
HRAS
NGF
SHC2
SHC3
SOS1
unknown CG mut
known HCG mut
wt

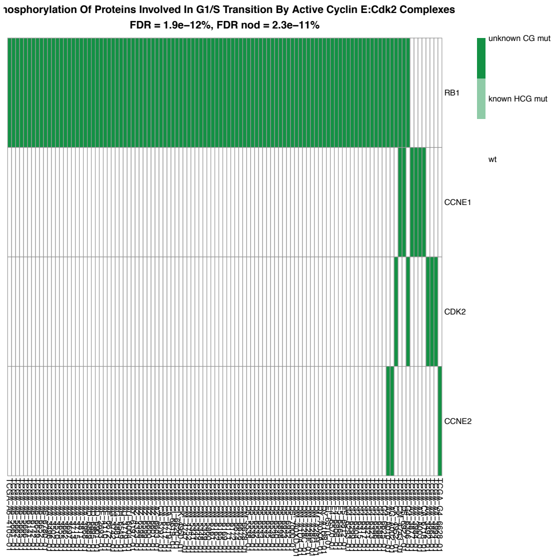
nosphorylation Of Proteins Involved In G1/S Transition By Active Cyclin E:Cdk2 Complexes
FDR = 1.9e−12%, FDR nod = 2.3e−11%
RB1
CCNE1
CDK2
CCNE2
unknown CG mut
known HCG mut
wt

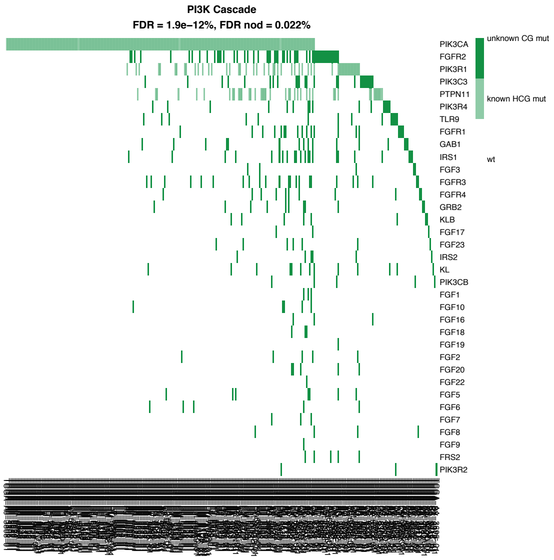
PI3K Cascade
FDR = 1.9e−12%, FDR nod = 0.022%
PIK3CA
FGFR2
PIK3R1
PIK3C3
PTPN11
PIK3R4
TLR9
FGFR1
GAB1
IRS1
FGF3
FGFR3
FGFR4
GRB2
KLB
FGF17
FGF23
IRS2
KL
PIK3CB
FGF1
FGF10
FGF16
FGF18
FGF19
FGF2
FGF20
FGF22
FGF5
FGF6
FGF7
FGF8
FGF9
FRS2
PIK3R2
unknown CG mut
known HCG mut
wt

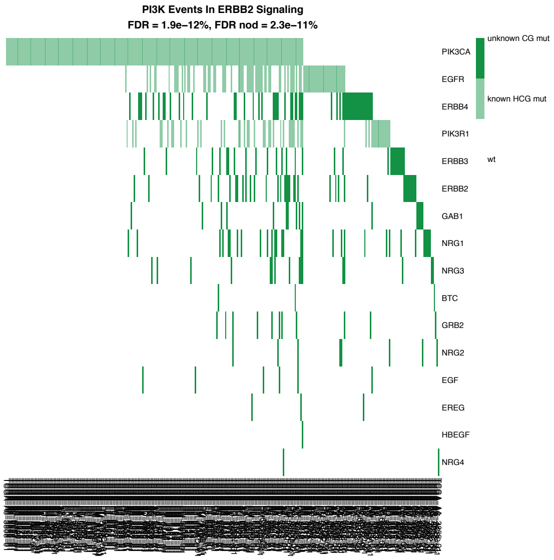
PI3K Events In ERBB2 Signaling
FDR = 1.9e−12%, FDR nod = 2.3e−11%
PIK3CA
EGFR
ERBB4
PIK3R1
ERBB3
ERBB2
GAB1
NRG1
NRG3
BTC
GRB2
NRG2
EGF
EREG
HBEGF
NRG4
unknown CG mut
known HCG mut
wt

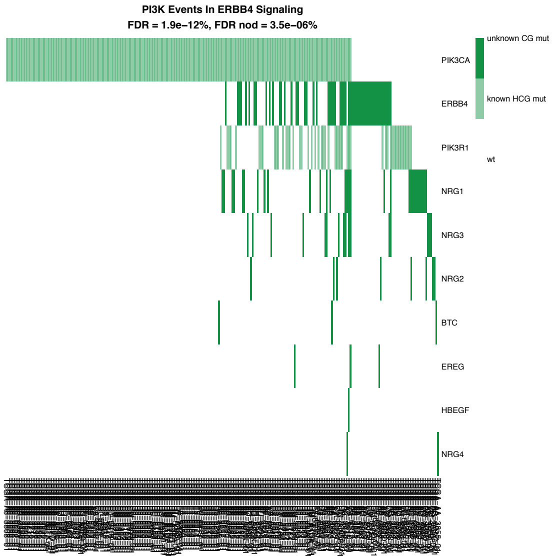
PI3K Events In ERBB4 Signaling
FDR = 1.9e−12%, FDR nod = 3.5e−06%
PIK3CA
ERBB4
PIK3R1
NRG1
NRG3
NRG2
BTC
EREG
HBEGF
NRG4
unknown CG mut
known HCG mut
wt

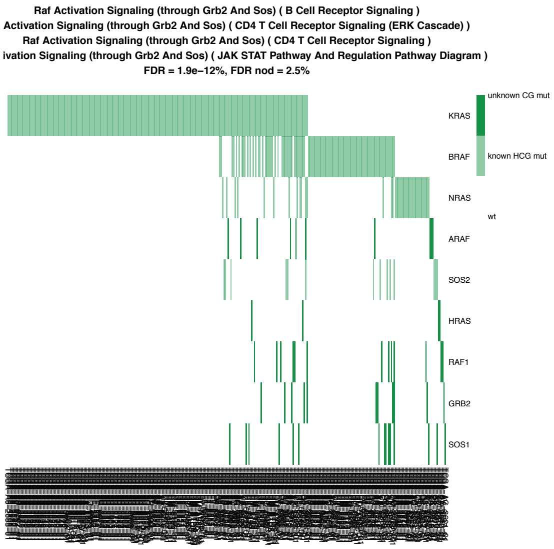
Raf Activation Signaling (through Grb2 And Sos) ( B Cell Receptor Signaling )
Activation Signaling (through Grb2 And Sos) ( CD4 T Cell Receptor Signaling (ERK Cascade) )
Raf Activation Signaling (through Grb2 And Sos) ( CD4 T Cell Receptor Signaling )
ivation Signaling (through Grb2 And Sos) ( JAK STAT Pathway And Regulation Pathway Diagram )
FDR = 1.9e−12%, FDR nod = 2.5%
KRAS
BRAF
NRAS
ARAF
SOS2
HRAS
RAF1
GRB2
SOS1
unknown CG mut
known HCG mut
wt

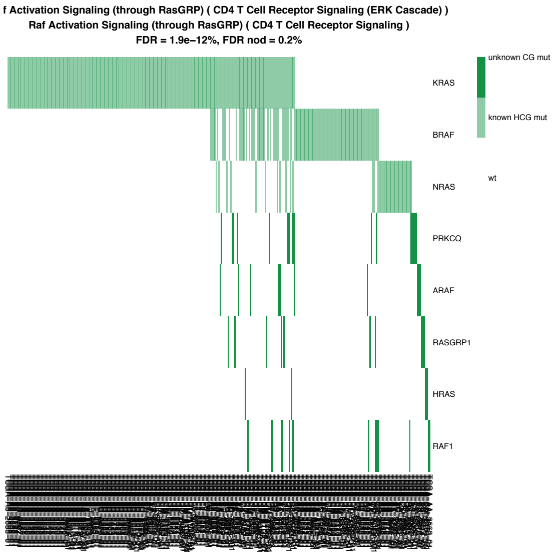
f Activation Signaling (through RasGRP) ( CD4 T Cell Receptor Signaling (ERK Cascade) )
Raf Activation Signaling (through RasGRP) ( CD4 T Cell Receptor Signaling )
FDR = 1.9e−12%, FDR nod = 0.2%
KRAS
BRAF
NRAS
PRKCQ
ARAF
RASGRP1
HRAS
RAF1
unknown CG mut
known HCG mut
wt

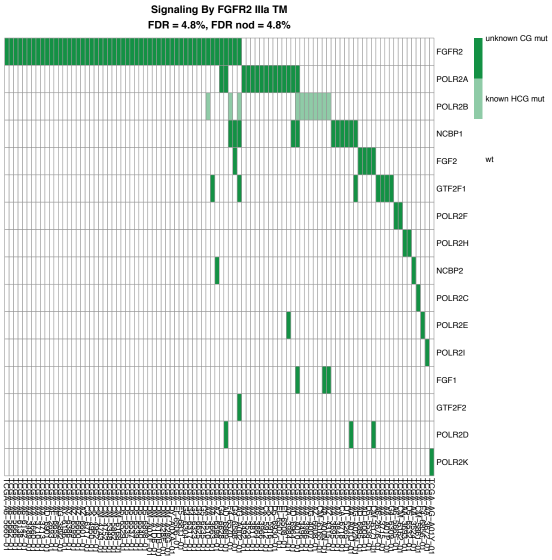
Signaling By FGFR2 IIIa TM
FDR = 4.8%, FDR nod = 4.8%
FGFR2
POLR2A
POLR2B
NCBP1
FGF2
GTF2F1
POLR2F
POLR2H
NCBP2
POLR2C
POLR2E
POLR2I
FGF1
GTF2F2
POLR2D
POLR2K
unknown CG mut
known HCG mut
wt

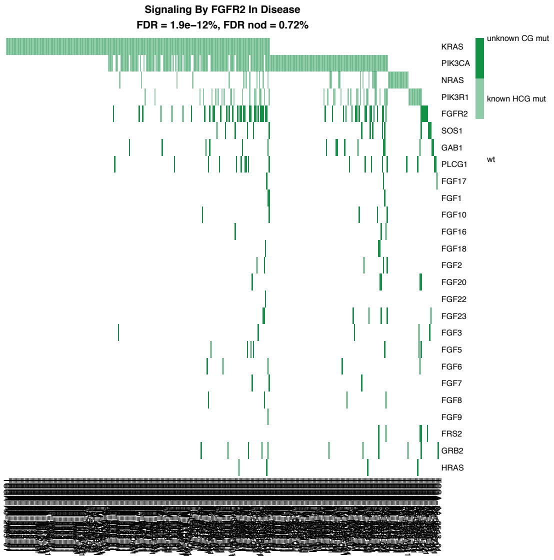
Signaling By FGFR2 In Disease
FDR = 1.9e−12%, FDR nod = 0.72%
KRAS
PIK3CA
NRAS
PIK3R1
FGFR2
SOS1
GAB1
PLCG1
FGF17
FGF1
FGF10
FGF16
FGF18
FGF2
FGF20
FGF22
FGF23
FGF3
FGF5
FGF6
FGF7
FGF8
FGF9
FRS2
GRB2
HRAS
unknown CG mut
known HCG mut
wt

**Signaling By FGFR3 Fusions In Cancer**
**FDR = 1.9e−12%, FDR nod = 2.1%**

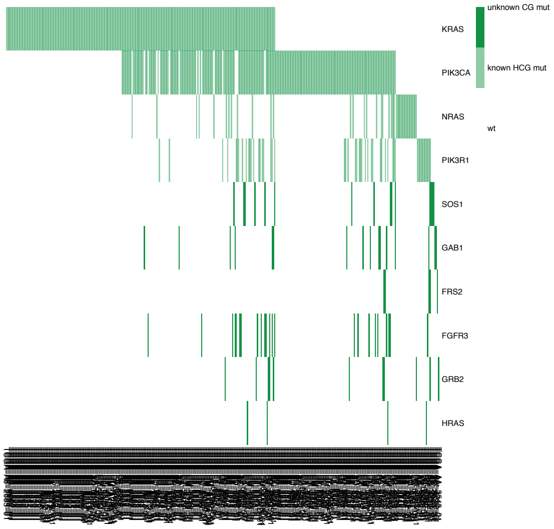


**Signalling To P38 Via RIT And RIN**
**FDR = 1.9e−12%, FDR nod = 0.072%**

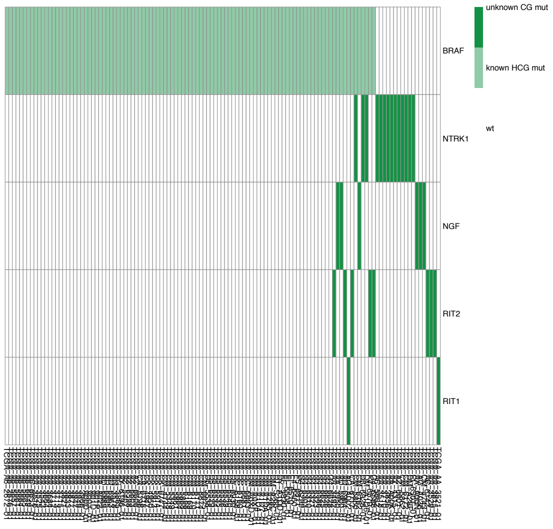


**T Cell Activation**
**FDR = 2.1e−12%, FDR nod = 1.9%**

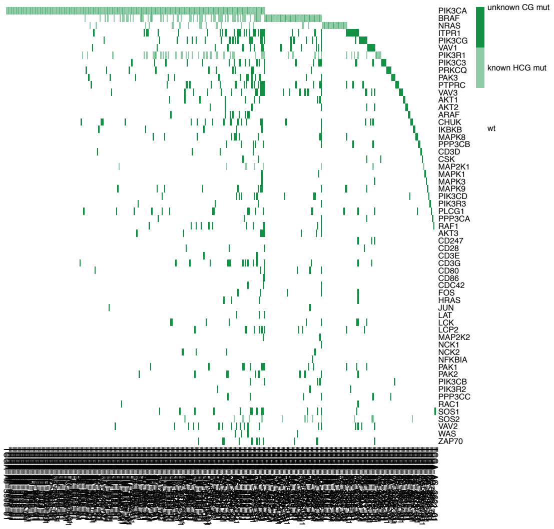


**TP53 Regulates Transcription Of Genes Involved In G2 Cell Cycle Arrest**
**FDR = 1.9e−12%, FDR nod = 0.011%**

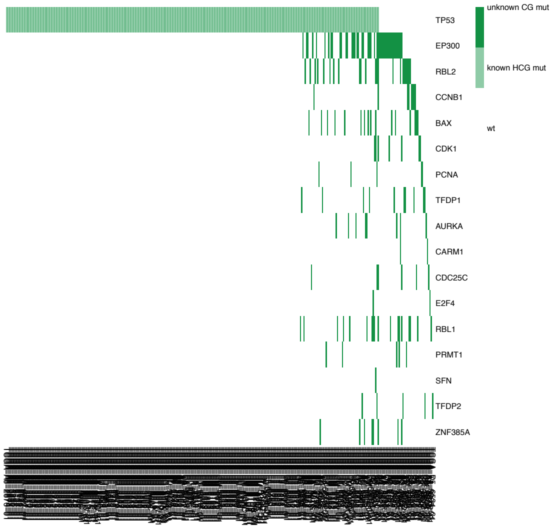


**VEGF Signaling Pathway**
**FDR = 1.9e−12%, FDR nod = 0.053%**

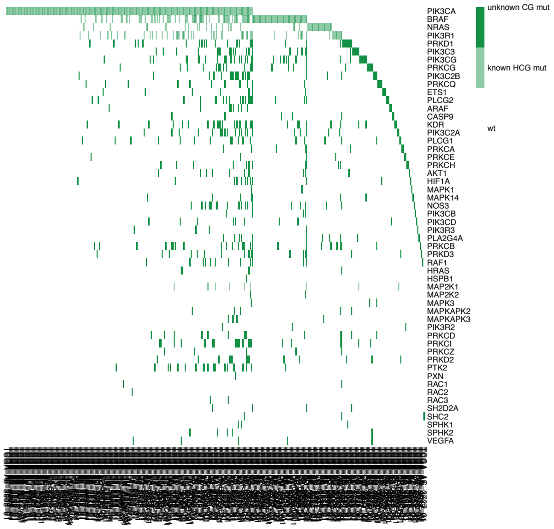


**BMP Signaling Pathway ( TGF Beta BMP Diagram(MolecularVariation) )**
**FDR = 1.9e−12%, FDR nod = 0.014%**

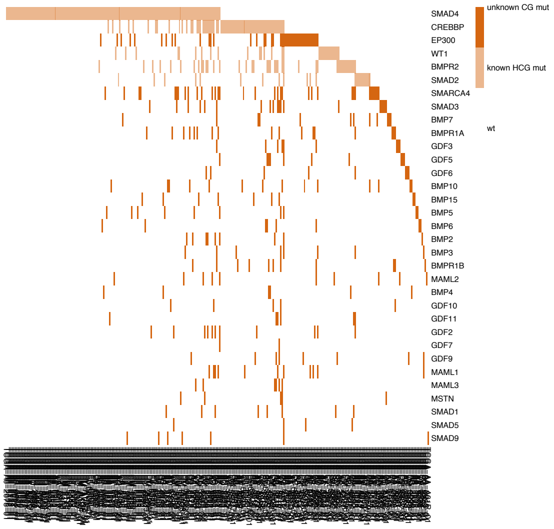

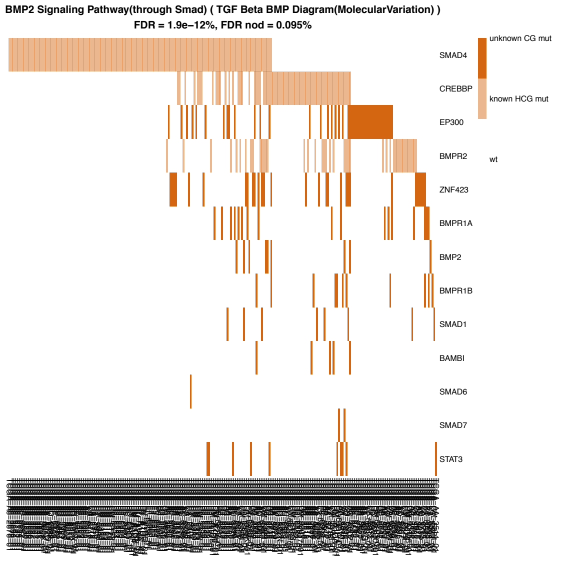
BMP2 Signaling Pathway(through Smad) ( TGF Beta BMP Diagram(MolecularVariation) )
FDR = 1.9e−12%, FDR nod = 0.095%
SMAD4
CREBBP
EP300
BMPR2
ZNF423
BMPR1A
BMP2
BMPR1B
SMAD1
BAMBI
SMAD6
SMAD7
STAT3
unknown CG mut
known HCG mut
wt

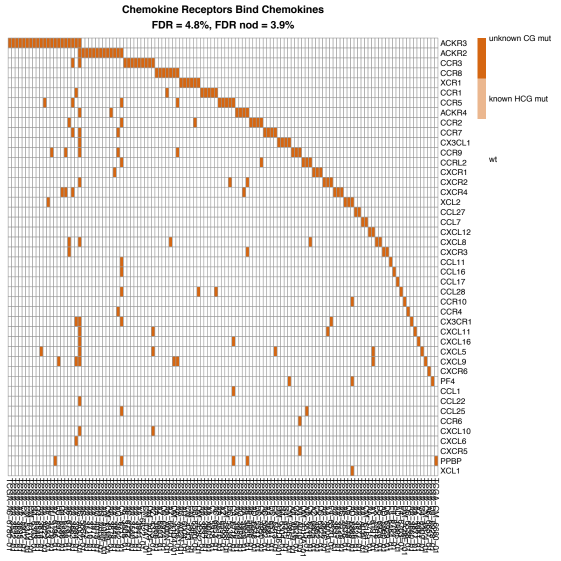
Chemokine Receptors Bind Chemokines
FDR = 4.8%, FDR nod = 3.9%
ACKR3
ACKR2
CCR3
CCR8
XCR1
CCR1
CCR5
ACKR4
CCR2
CCR7
CX3CL1
CCR9
CCRL2
CXCR1
CXCR2
CXCR4
XCL2
CCL27
CCL7
CXCL12
CXCL8
CXCR3
CCL11
CCL16
CCL17
CCL28
CCR10
CCR4
CX3CR1
CXCL11
CXCL16
CXCL5
CXCL9
CXCR6
PF4
CCL1
CCL22
CCL25
CCR6
CXCL10
CXCL6
CXCR5
PPBP
XCL1
unknown CG mut
known HCG mut
wt

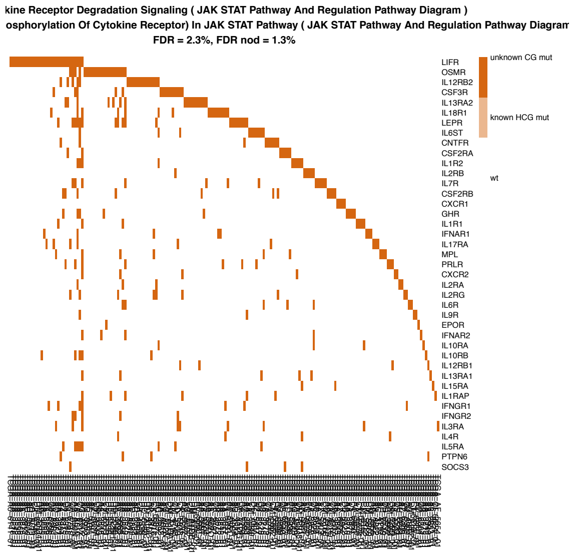
kine Receptor Degradation Signaling ( JAK STAT Pathway And Regulation Pathway Diagram )
osphorylation Of Cytokine Receptor) In JAK STAT Pathway ( JAK STAT Pathway And Regulation Pathway Diagram
FDR = 2.3%, FDR nod = 1.3%
LIFR
OSMR
IL12RB2
CSF3R
IL13RA2
IL18R1
LEPR
IL6ST
CNTFR
CSF2RA
IL1R2
IL2RB
IL7R
CSF2RB
CXCR1
GHR
IL1R1
IFNAR1
IL17RA
MPL
PRLR
CXCR2
IL2RA
IL2RG
IL6R
IL9R
EPOR
IFNAR2
IL10RA
IL10RB
IL12RB1
IL13RA1
IL15RA
IL1RAP
IFNGR1
IFNGR2
IL3RA
IL4R
IL5RA
PTPN6
SOCS3
unknown CG mut
known HCG mut
wt

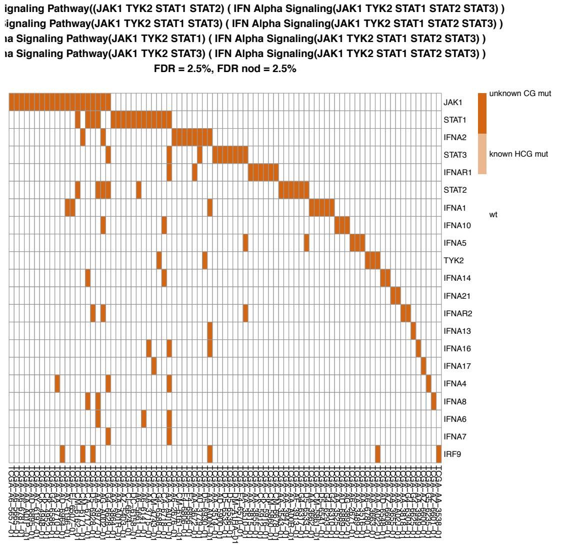
ignaling Pathway((JAK1 TYK2 STAT1 STAT2) ( IFN Alpha Signaling(JAK1 TYK2 STAT1 STAT2 STAT3) )
ignaling Pathway(JAK1 TYK2 STAT1 STAT3) ( IFN Alpha Signaling(JAK1 TYK2 STAT1 STAT2 STAT3) )
ıa Signaling Pathway(JAK1 TYK2 STAT1) ( IFN Alpha Signaling(JAK1 TYK2 STAT1 STAT2 STAT3) )
ıa Signaling Pathway(JAK1 TYK2 STAT3) ( IFN Alpha Signaling(JAK1 TYK2 STAT1 STAT2 STAT3) )
FDR = 2.5%, FDR nod = 2.5%
JAK1
STAT1
IFNA2
STAT3
IFNAR1
STAT2
IFNA1
IFNA10
IFNA5
TYK2
IFNA14
IFNA21
IFNAR2
IFNA13
IFNA16
IFNA17
IFNA4
IFNA8
IFNA6
IFNA7
IRF9
unknown CG mut
known HCG mut
wt

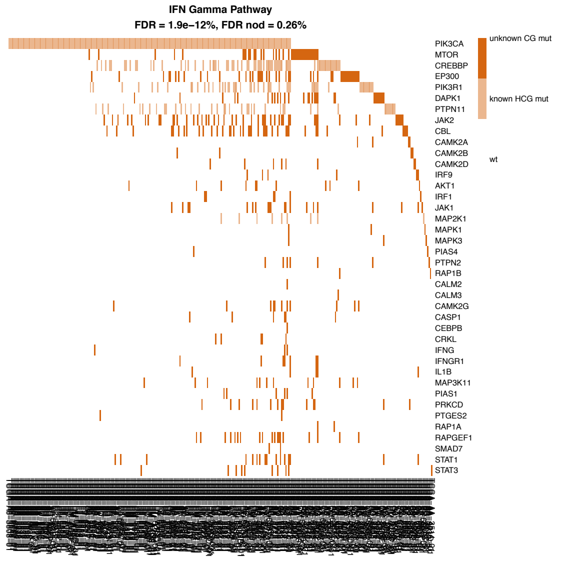
IFN Gamma Pathway
FDR = 1.9e−12%, FDR nod = 0.26%
PIK3CA
MTOR
CREBBP
EP300
PIK3R1
DAPK1
PTPN11
JAK2
CBL
CAMK2A
CAMK2B
CAMK2D
IRF9
AKT1
IRF1
JAK1
MAP2K1
MAPK1
MAPK3
PIAS4
PTPN2
RAP1B
CALM2
CALM3
CAMK2G
CASP1
CEBPB
CRKL
IFNG
IFNGR1
IL1B
MAP3K11
PIAS1
PRKCD
PTGES2
RAP1A
RAPGEF1
SMAD7
STAT1
STAT3
unknown CG mut
known HCG mut
wt

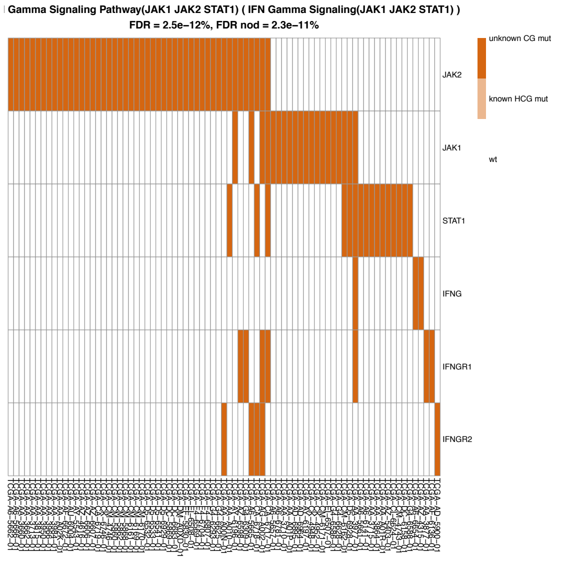
Gamma Signaling Pathway(JAK1 JAK2 STAT1) ( IFN Gamma Signaling(JAK1 JAK2 STAT1) )
FDR = 2.5e−12%, FDR nod = 2.3e−11%
JAK2
JAK1
STAT1
IFNG
IFNGR1
IFNGR2
unknown CG mut
known HCG mut
wt

**IL 6 Type Cytokine Receptor Ligand Interactions**
**FDR = 1.4e−09%, FDR nod = 1.2e−09%**

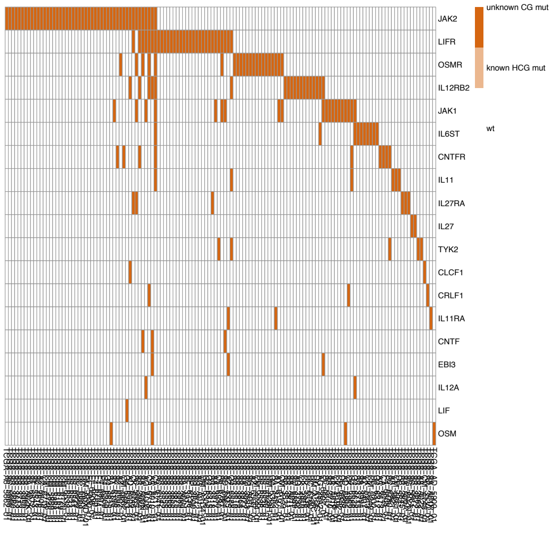


**IRF3 Mediated Activation Of Type 1 IFN**
**FDR = 0.04%, FDR nod = 0.056%**

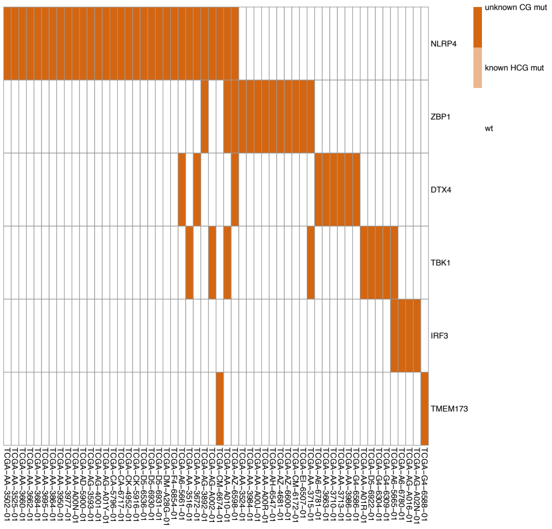


**IRF3 Mediated Induction Of Type I IFN**
**FDR = 2.2e−07%, FDR nod = 2.2e−07%**

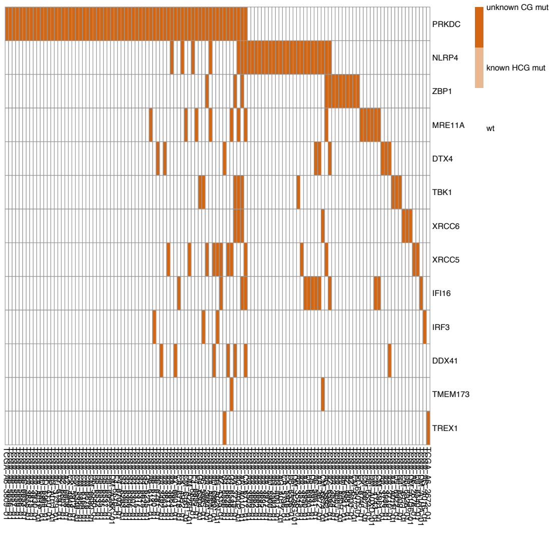


**JNK Cascade ( TGF Beta Signaling(through TAK1) )**
**FDR = 0.38%, FDR nod = 0.45%**

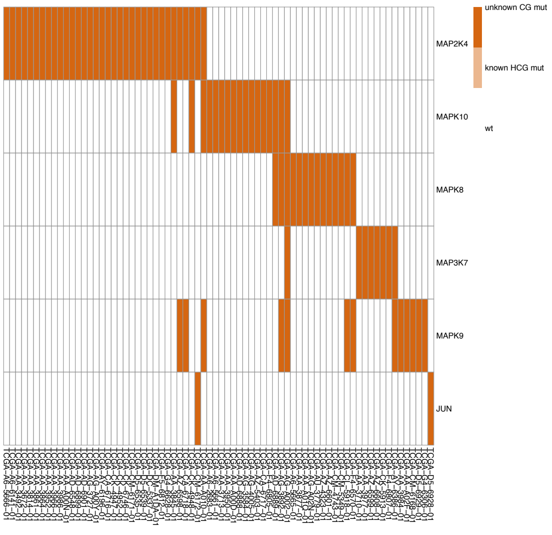


**LRR FLII Interacting Protein 1 (LRRFIP1) Activates Type I IFN Production**
**FDR = 1.9e−12%, FDR nod = 0.021%**

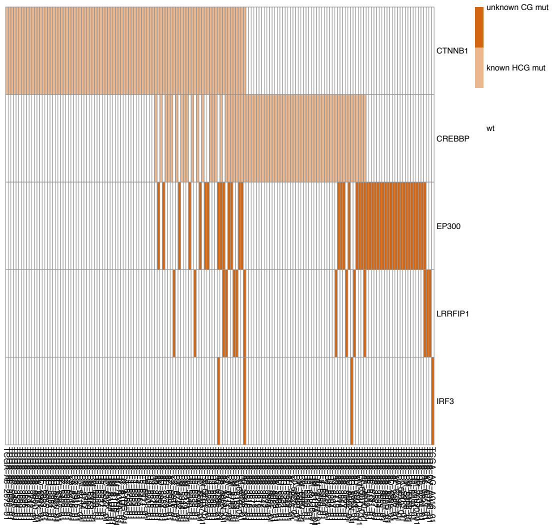


**Regulation Of IFNA Signaling**
**FDR = 1.9e−07%, FDR nod = 0.54%**

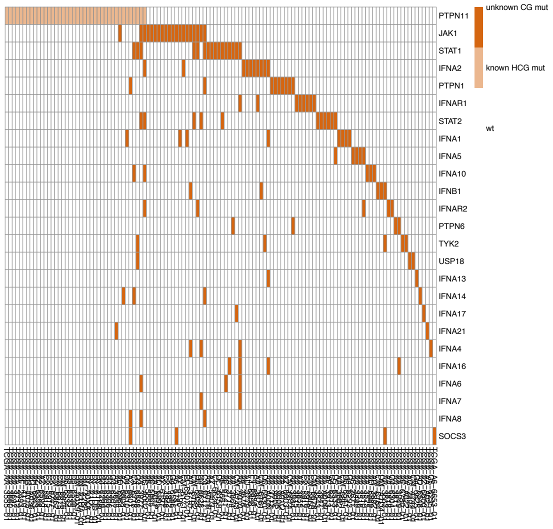

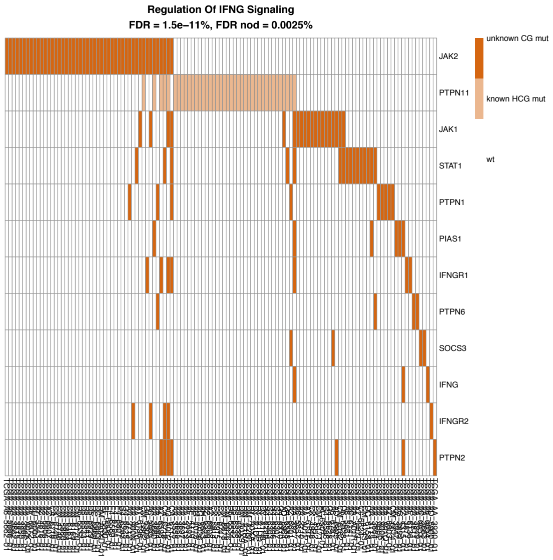
Regulation Of IFNG Signaling
FDR = 1.5e−11%, FDR nod = 0.0025%
JAK2
PTPN11
JAK1
STAT1
PTPN1
PIAS1
IFNGR1
PTPN6
SOCS3
IFNG
IFNGR2
PTPN2
unknown CG mut
known HCG mut
wt

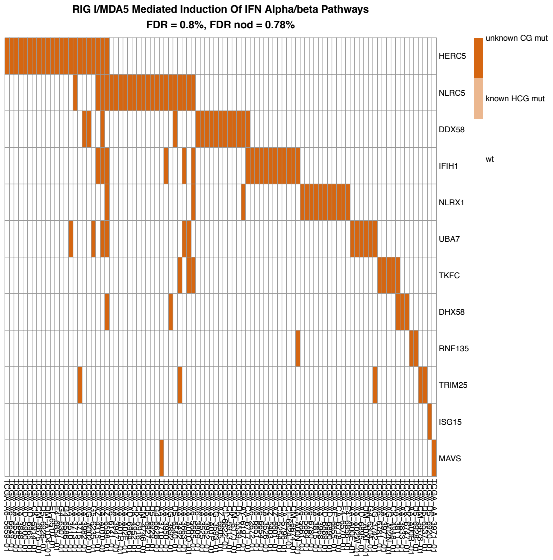
RIG I/MDA5 Mediated Induction Of IFN Alpha/beta Pathways
FDR = 0.8%, FDR nod = 0.78%
HERC5
NLRC5
DDX58
IFIH1
NLRX1
UBA7
TKFC
DHX58
RNF135
TRIM25
ISG15
MAVS
unknown CG mut
known HCG mut
wt

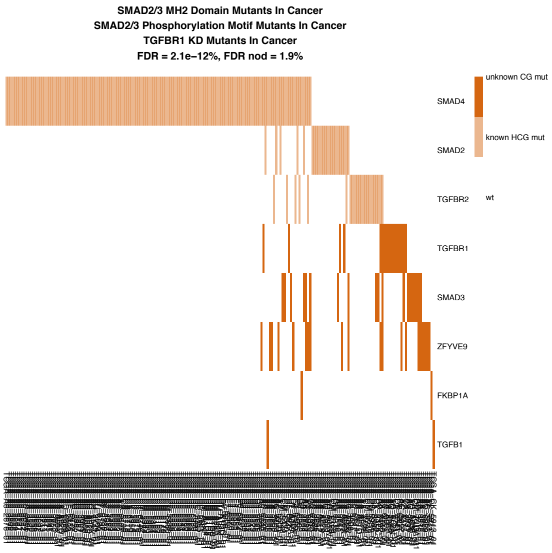
SMAD2/3 MH2 Domain Mutants In Cancer
SMAD2/3 Phosphorylation Motif Mutants In Cancer
TGFBR1 KD Mutants In Cancer
FDR = 2.1e−12%, FDR nod = 1.9%
SMAD4
SMAD2
TGFBR2
TGFBR1
SMAD3
ZFYVE9
FKBP1A
TGFB1
unknown CG mut
known HCG mut
wt

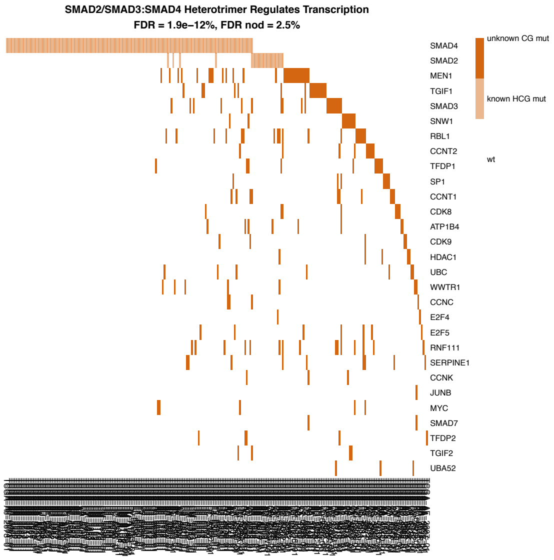
SMAD2/SMAD3:SMAD4 Heterotrimer Regulates Transcription
FDR = 1.9e−12%, FDR nod = 2.5%
SMAD4
SMAD2
MEN1
TGIF1
SMAD3
SNW1
RBL1
CCNT2
TFDP1
SP1
CCNT1
CDK8
ATP1B4
CDK9
HDAC1
UBC
WWTR1
CCNC
E2F4
E2F5
RNF111
SERPINE1
CCNK
JUNB
MYC
SMAD7
TFDP2
TGIF2
UBA52
unknown CG mut
known HCG mut
wt

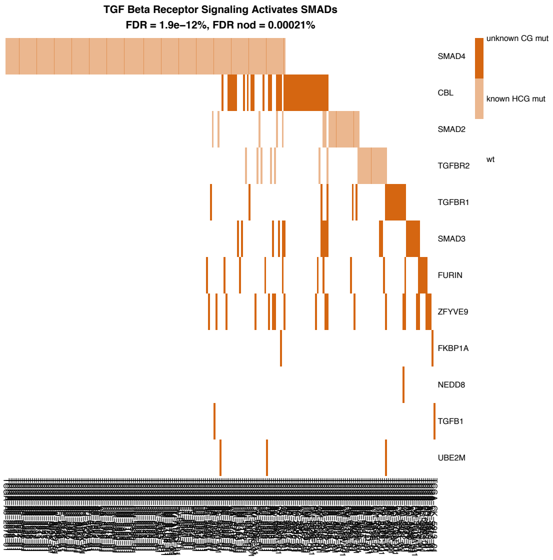
TGF Beta Receptor Signaling Activates SMADs
FDR = 1.9e−12%, FDR nod = 0.00021%
SMAD4
CBL
SMAD2
TGFBR2
TGFBR1
SMAD3
FURIN
ZFYVE9
FKBP1A
NEDD8
TGFB1
UBE2M
unknown CG mut
known HCG mut
wt

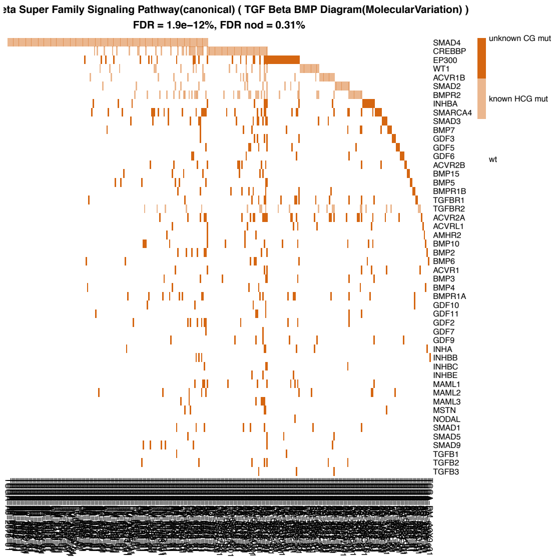
eta Super Family Signaling Pathway(canonical) ( TGF Beta BMP Diagram(MolecularVariation) )
FDR = 1.9e−12%, FDR nod = 0.31%
SMAD4
CREBBP
EP300
WT1
ACVR1B
SMAD2
BMPR2
INHBA
SMARCA4
SMAD3
BMP7
GDF3
GDF5
GDF6
ACVR2B
BMP15
BMP5
BMPR1B
TGFBR1
TGFBR2
ACVR2A
ACVRL1
AMHR2
BMP10
BMP2
BMP6
ACVR1
BMP3
BMP4
BMPR1A
GDF10
GDF11
GDF2
GDF7
GDF9
INHA
INHBB
INHBC
INHBE
MAML1
MAML2
MAML3
MSTN
NODAL
SMAD1
SMAD5
SMAD9
TGFB1
TGFB2
TGFB3
unknown CG mut
known HCG mut
wt

### GBM

**Signaling By PDGF**
**FDR = 0.0055%, FDR nod = 0.22%**

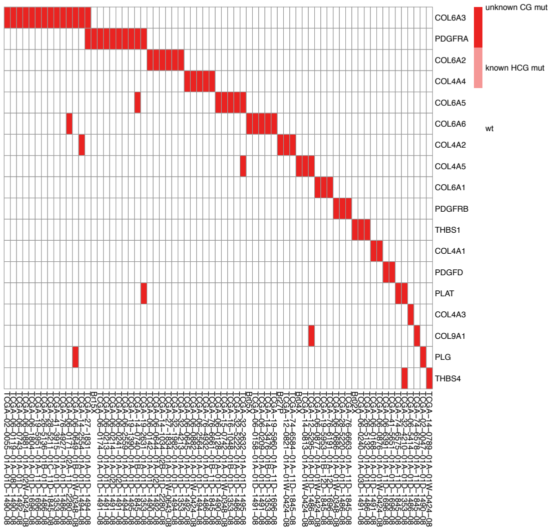


**GRB2:SOS Provides Linkage To MAPK Signaling For Integrins**
**FDR = 3.7%, FDR nod = 3.9%**

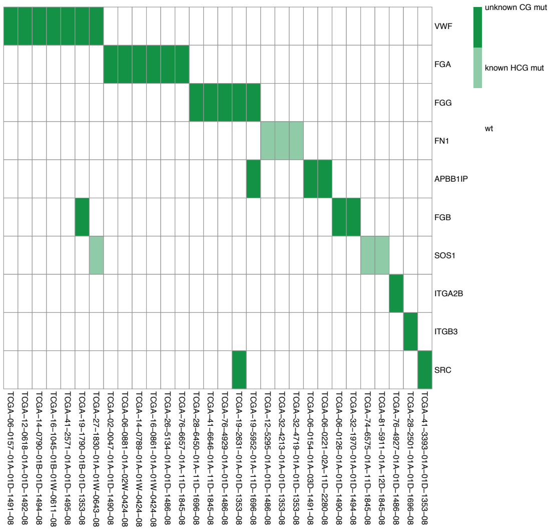


**P130Cas Linkage To MAPK Signaling For Integrins**
**FDR = 1.9%, FDR nod = 3.9%**

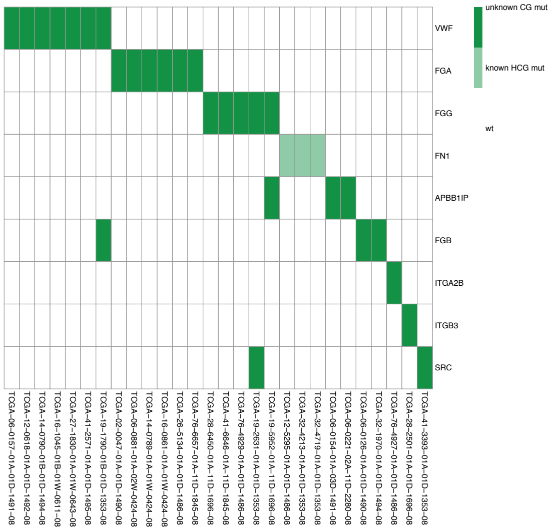

### HSNC

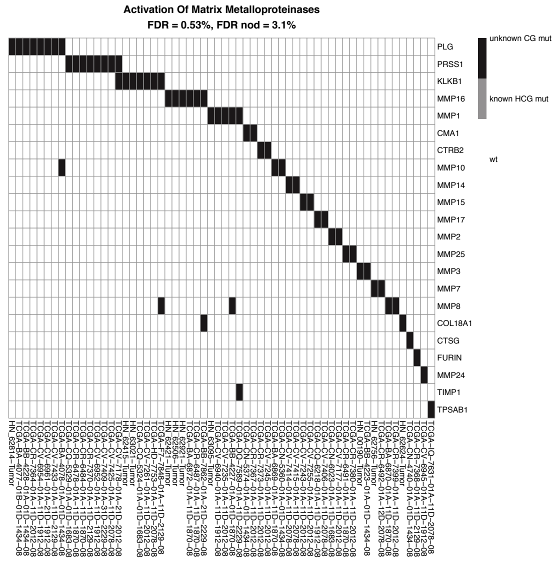
Activation Of Matrix Metalloproteinases
FDR = 0.53%, FDR nod = 3.1%
PLG
PRSS1
KLKB1
MMP16
MMP1
CMA1
CTRB2
MMP10
MMP14
MMP15
MMP17
MMP2
MMP25
MMP3
MMP7
MMP8
COL18A1
CTSG
FURIN
MMP24
TIMP1
TPSAB1
unknown CG mut
known HCG mut
wt

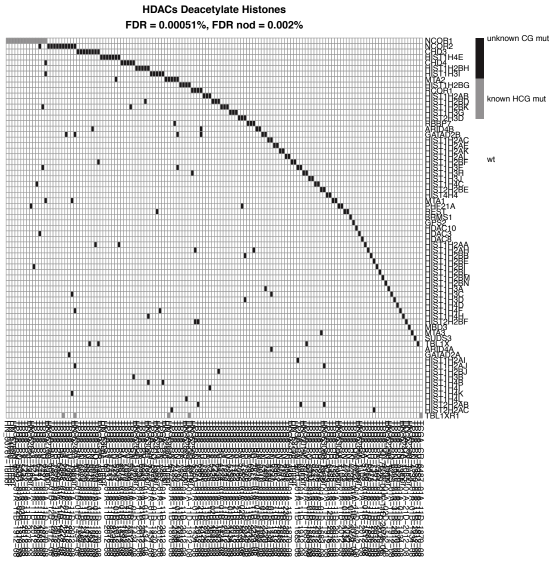
HDACs Deacetylate Histones
FDR = 0.00051%, FDR nod = 0.002%
unknown CG mut
known HCG mut
wt

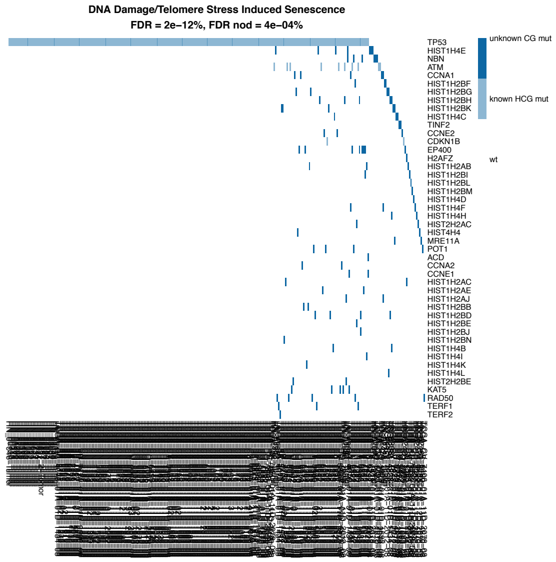
DNA Damage/Telomere Stress Induced Senescence
FDR = 2e−12%, FDR nod = 4e−04%
TP53
HIST1H4E
NBN
ATM
CCNA1
HIST1H2BF
HIST1H2BG
HIST1H2BH
HIST1H2BK
HIST1H4C
TINF2
CCNE2
CDKN1B
EP400
H2AFZ
HIST1H2AB
HIST1H2BI
HIST1H2BL
HIST1H2BM
HIST1H4D
HIST1H4F
HIST1H4H
HIST2H2AC
HIST4H4
MRE11A
POT1
ACD
CCNA2
CCNE1
HIST1H2AC
HIST1H2AE
HIST1H2AJ
HIST1H2BB
HIST1H2BD
HIST1H2BE
HIST1H2BJ
HIST1H2BN
HIST1H4B
HIST1H4I
HIST1H4K
HIST1H4L
HIST2H2BE
KAT5
RAD50
TERF1
TERF2
unknown CG mut
known HCG mut
wt

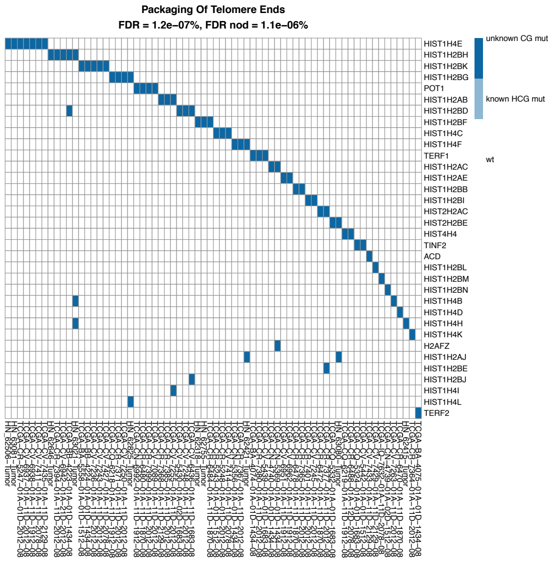
Packaging Of Telomere Ends
FDR = 1.2e−07%, FDR nod = 1.1e−06%
HIST1H4E
HIST1H2BH
HIST1H2BK
HIST1H2BG
POT1
HIST1H2AB
HIST1H2BD
HIST1H2BF
HIST1H4C
HIST1H4F
TERF1
HIST1H2AC
HIST1H2AE
HIST1H2BB
HIST1H2BI
HIST2H2AC
HIST2H2BE
HIST4H4
TINF2
ACD
HIST1H2BL
HIST1H2BM
HIST1H2BN
HIST1H4B
HIST1H4D
HIST1H4H
HIST1H4K
H2AFZ
HIST1H2AJ
HIST1H2BE
HIST1H2BJ
HIST1H4I
HIST1H4L
TERF2
unknown CG mut
known HCG mut
wt

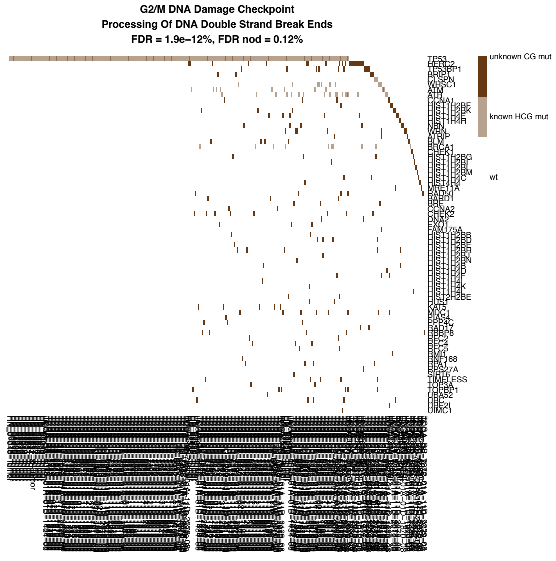
G2/M DNA Damage Checkpoint
Processing Of DNA Double Strand Break Ends
FDR = 1.9e−12%, FDR nod = 0.12%
unknown CG mut
known HCG mut
wt

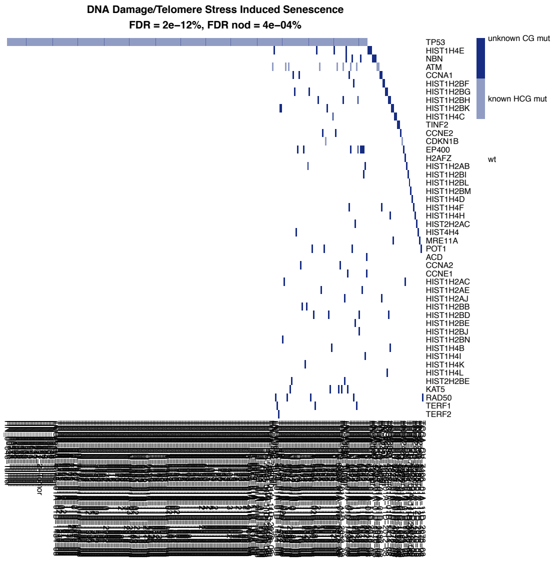
DNA Damage/Telomere Stress Induced Senescence
FDR = 2e−12%, FDR nod = 4e−04%
TP53
HIST1H4E
NBN
ATM
CCNA1
HIST1H2BF
HIST1H2BG
HIST1H2BH
HIST1H2BK
HIST1H4C
TINF2
CCNE2
CDKN1B
EP400
H2AFZ
HIST1H2AB
HIST1H2BI
HIST1H2BL
HIST1H2BM
HIST1H4D
HIST1H4F
HIST1H4H
HIST2H2AC
HIST4H4
MRE11A
POT1
ACD
CCNA2
CCNE1
HIST1H2AC
HIST1H2AE
HIST1H2AJ
HIST1H2BB
HIST1H2BD
HIST1H2BE
HIST1H2BJ
HIST1H2BN
HIST1H4B
HIST1H4I
HIST1H4K
HIST1H4L
HIST2H2BE
KAT5
RAD50
TERF1
TERF2
unknown CG mut
known HCG mut
wt

**FOXM1 Transcription Factor Network**
**FDR = 2e−12%, FDR nod = 0.006%**

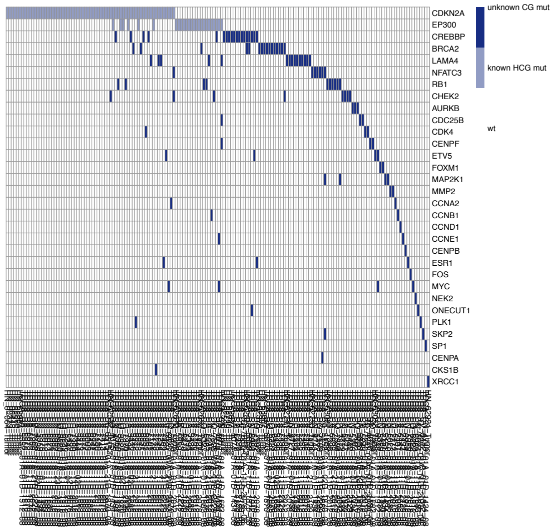


**G2/M DNA Damage Checkpoint**
**Processing Of DNA Double Strand Break Ends**
**FDR = 1.9e−12%, FDR nod = 0.12%**

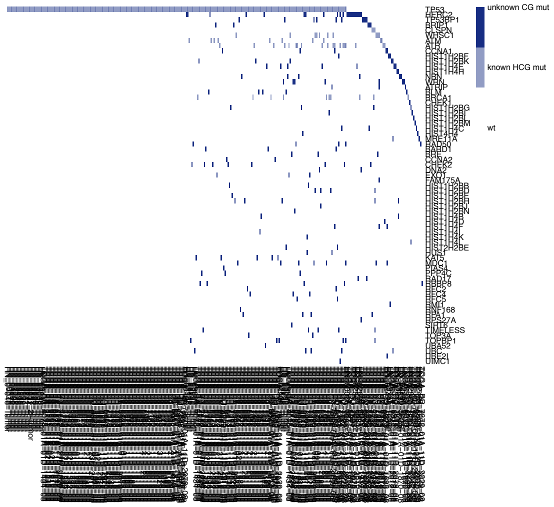


**Meiotic Recombination**
**FDR = 1.7e−11%, FDR nod = 1.4e−10%**

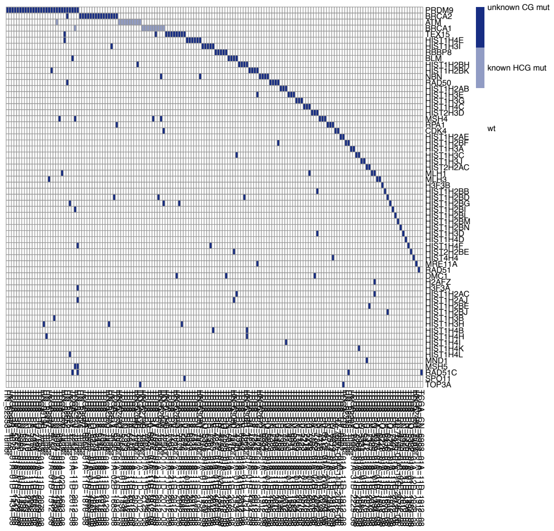


**Meiotic Synapsis**
**FDR = 1.7e−07%, FDR nod = 9e−09%**

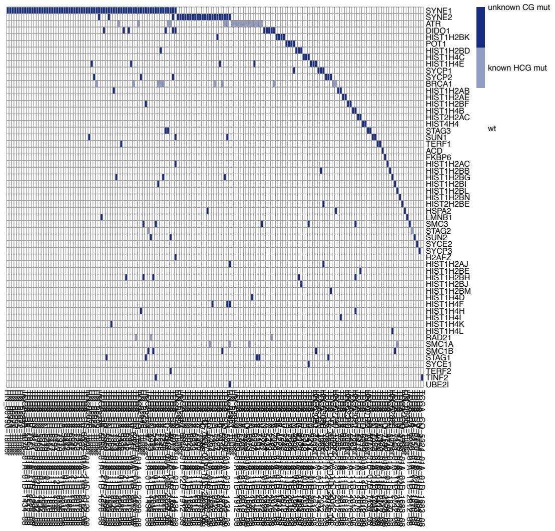


**nd ATM Mediated Phosphorylation Of Repair And Signaling Proteins At DNA Double Strand Breaks**
**FDR = 1.9e−12%, FDR nod = 0.028%**

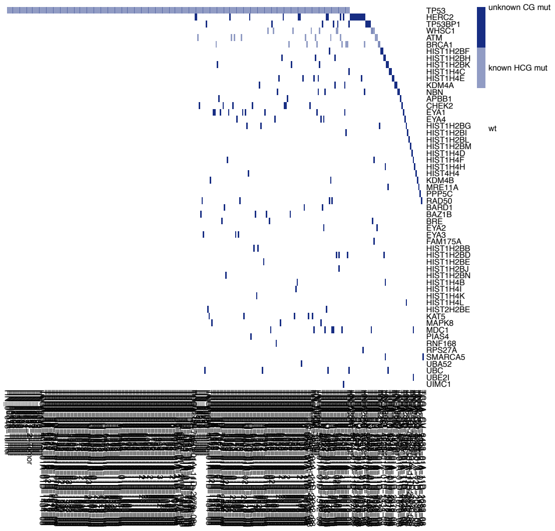


**Regulation Of Gene Expression By Hypoxia Inducible Factor**
**FDR = 1.8e−06%, FDR nod = 2.3%**

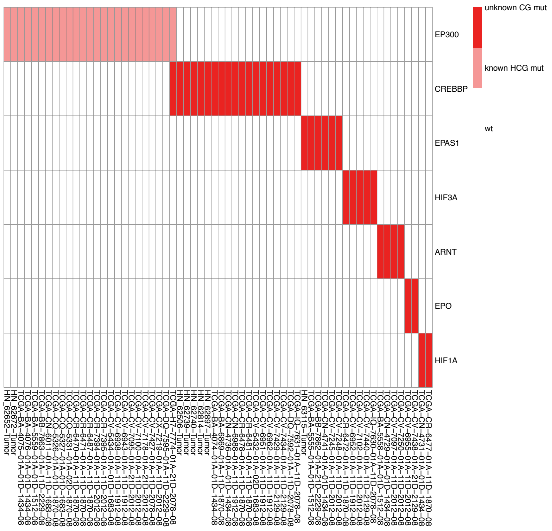

**Signaling By PDGF**
**FDR = 0.65%, FDR nod = 3.1%**

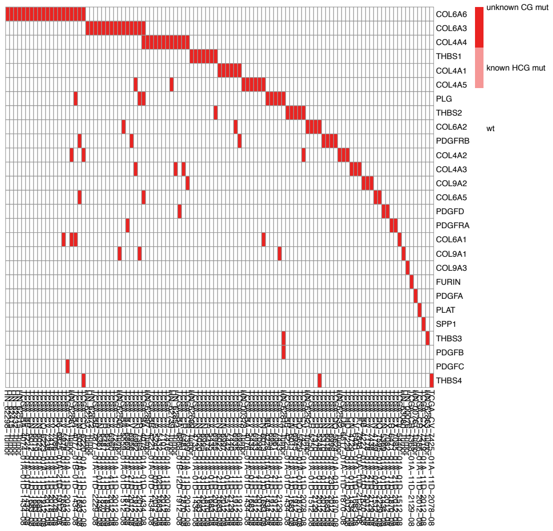


**G2/M DNA Damage Checkpoint**
**Processing Of DNA Double Strand Break Ends**
**FDR = 1.9e−12%, FDR nod = 0.12%**

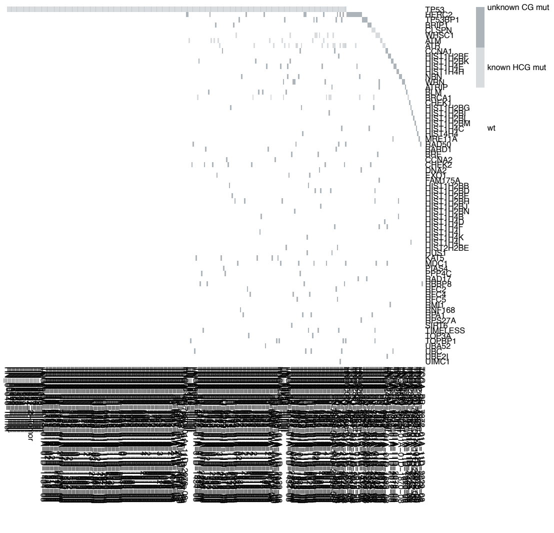


**ınd ATM Mediated Phosphorylation Of Repair And Signaling Proteins At DNA Double Strand Breaks**
**FDR = 1.9e−12%, FDR nod = 0.028%**

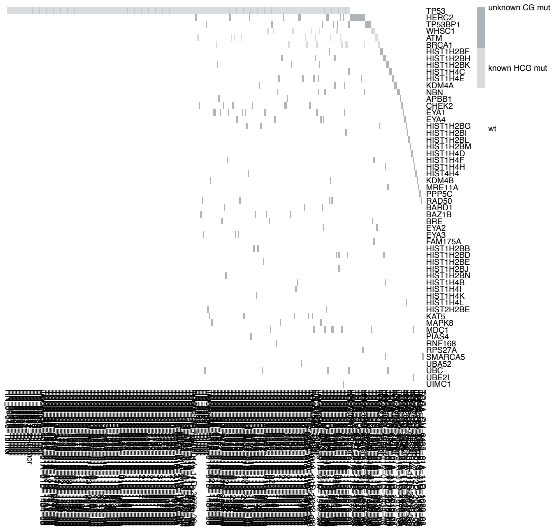

### KIRC

**Regulation Of IFNA Signaling**
**FDR = 1.1%, FDR nod = 2.1%**

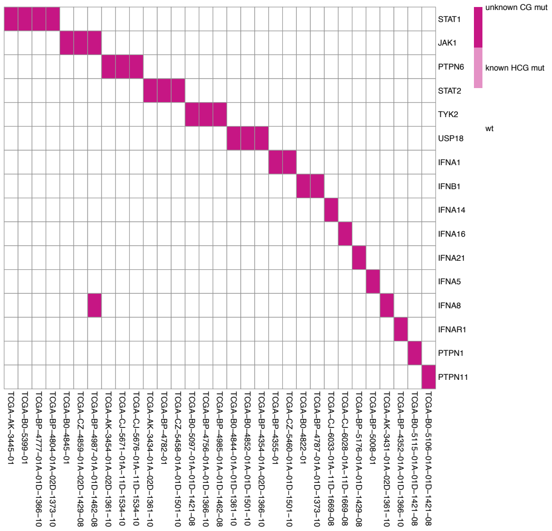


**DNA Damage/Telomere Stress Induced Senescence**
**FDR = 0.14%, FDR nod = 1.9%**

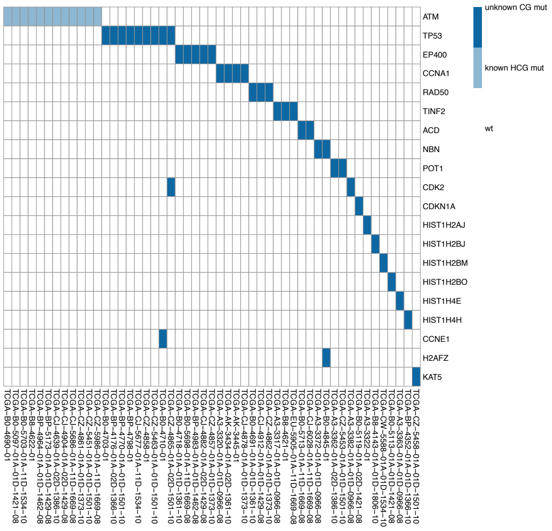


**Formation Of Senescence Associated Heterochromatin Foci (SAHF)**
**FDR = 0.97%, FDR nod = 2.1%**

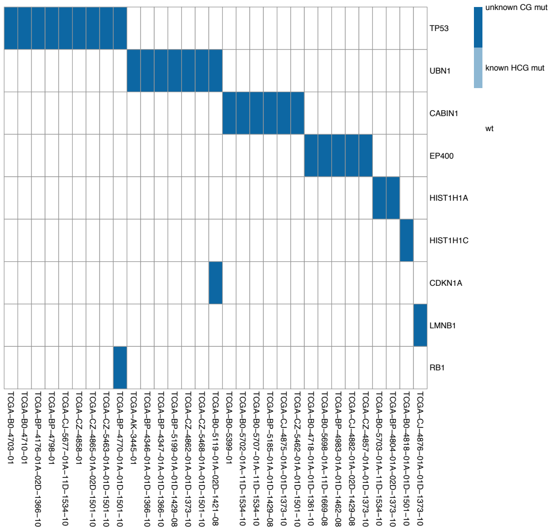


**G2/M DNA Damage Checkpoint**
**Processing Of DNA Double Strand Break Ends**
**FDR = 0.97%, FDR nod = 2.1%**

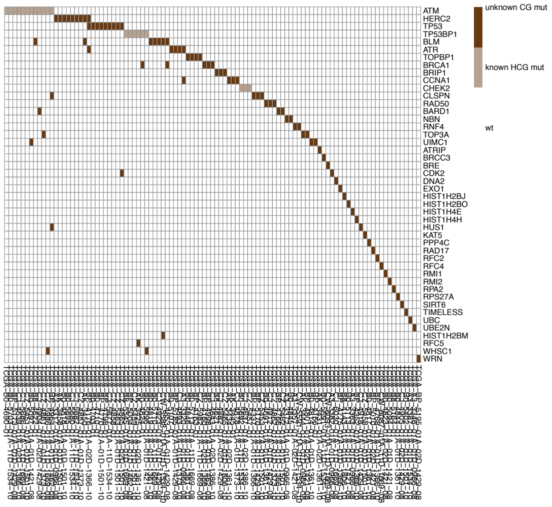


**Transcription Of Additional Cell Cycle Genes Whose Exact Role In The P53 Pathway Remain Uncertain**
**FDR = 0.67%, FDR nod = 0.97%**

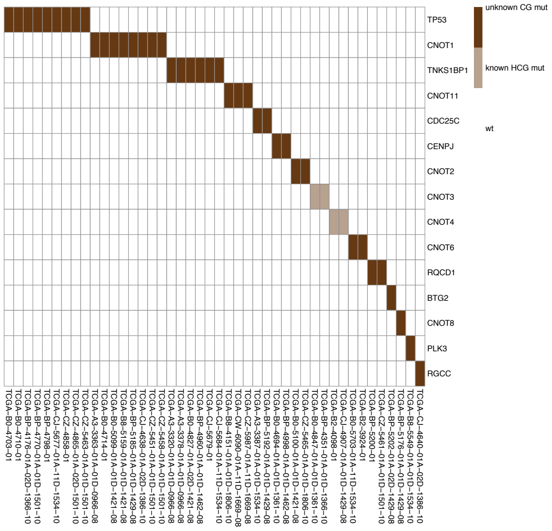


**TP53 Regulates Transcription Of Genes Involved In G2 Cell Cycle Arrest**
**FDR = 1%, FDR nod = 2.1%**

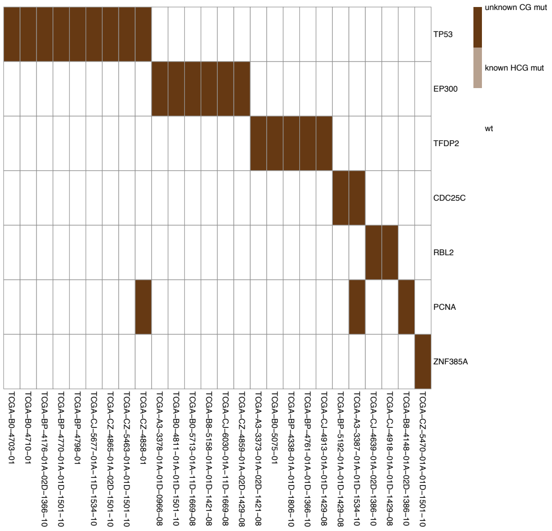

**DNA Damage/Telomere Stress Induced Senescence**
**FDR = 0.14%, FDR nod = 1.9%**

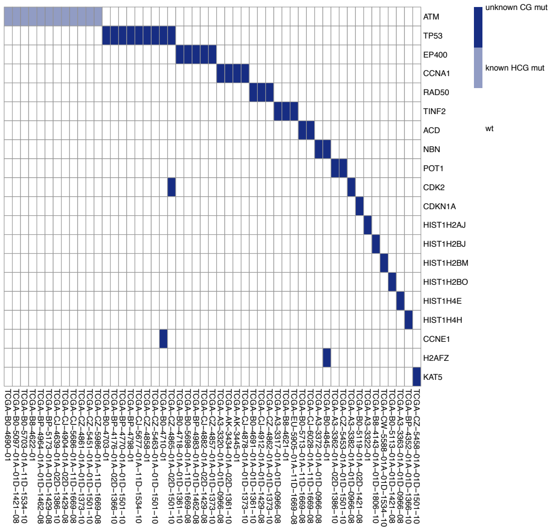


**FOXM1 Transcription Factor Network**
**FDR = 0.29%, FDR nod = 1.2%**

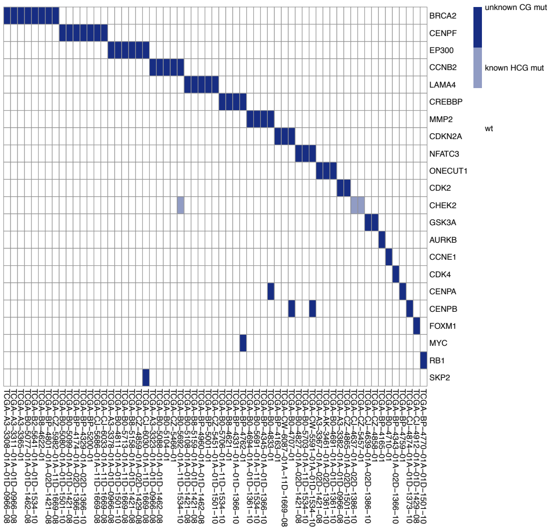


**G2/M DNA Damage Checkpoint**
**Processing Of DNA Double Strand Break Ends**
**FDR = 0.97%, FDR nod = 2.1%**

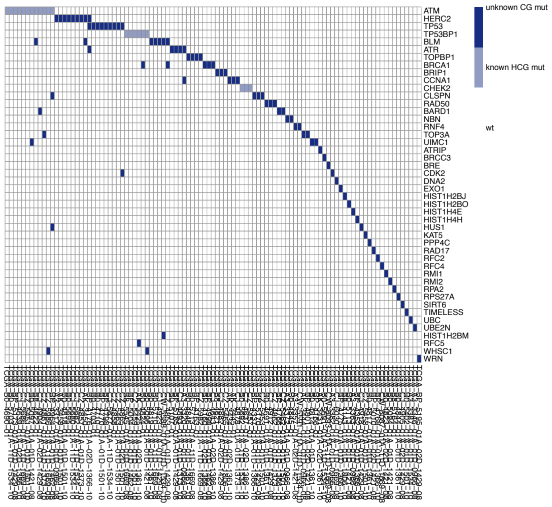


**Meiotic Recombination**
**FDR = 0.22%, FDR nod = 1.8%**

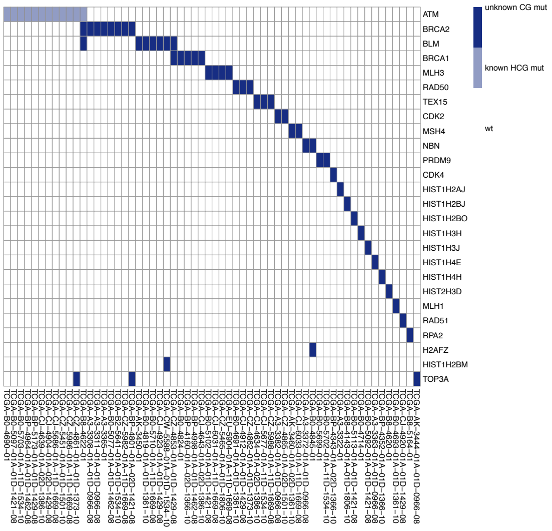


**PDGFR Alpha Signaling Pathway**
**FDR = 3%, FDR nod = 2.1%**

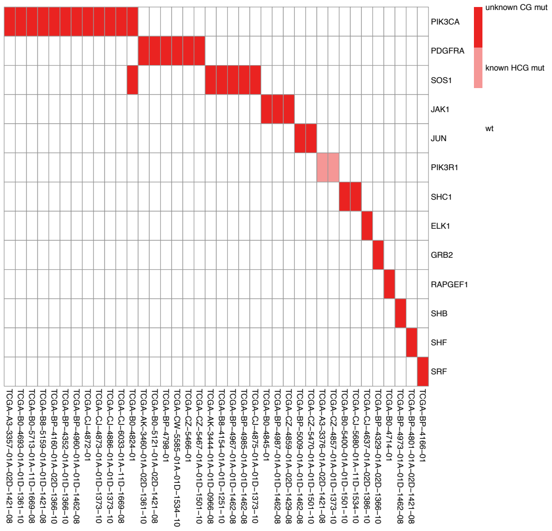


**Signaling By PDGF**
**FDR = 0.21%, FDR nod = 0.97%**

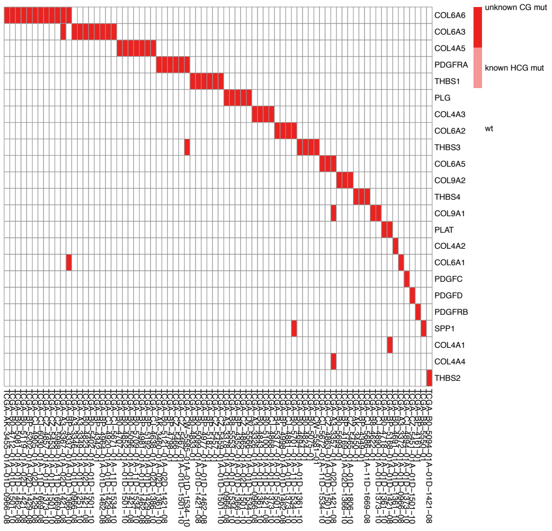

**G2/M DNA Damage Checkpoint**
**Processing Of DNA Double Strand Break Ends**
**FDR = 0.97%, FDR nod = 2.1%**

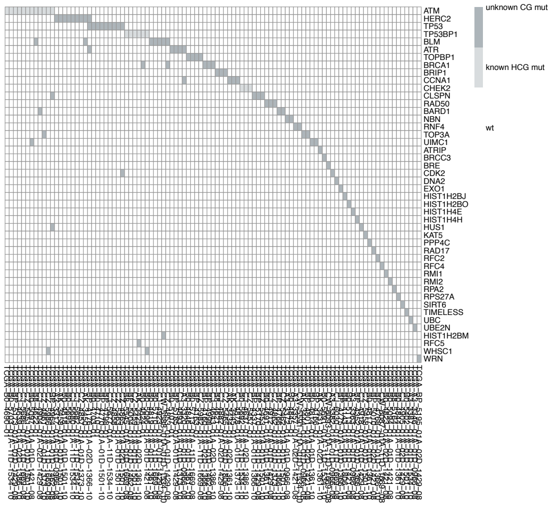


**PI3K Events In ERBB4 Signaling**
**FDR = 4.8%, FDR nod = 3%**

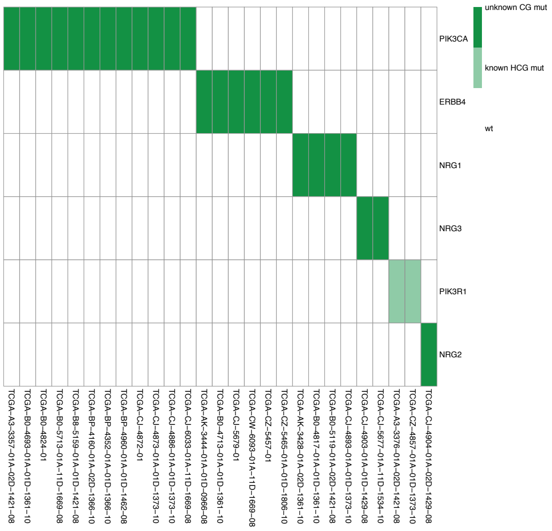


**Transcription Of Additional Cell Cycle Genes Whose Exact Role In The P53 Pathway Remain Uncertain**
**FDR = 0.67%, FDR nod = 0.97%**

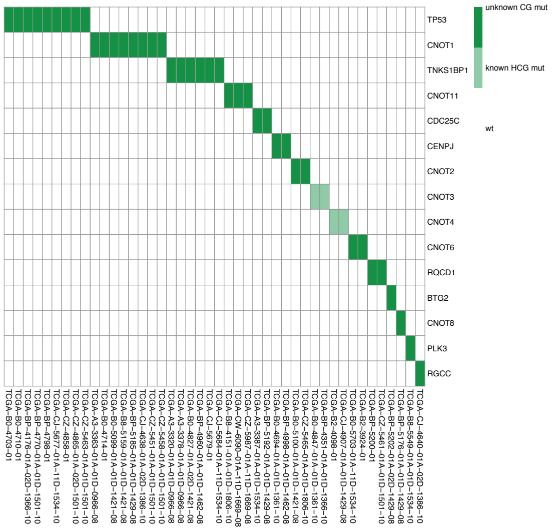


**TP53 Regulates Transcription Of Genes Involved In G2 Cell Cycle Arrest**
**FDR = 1%, FDR nod = 2.1%**

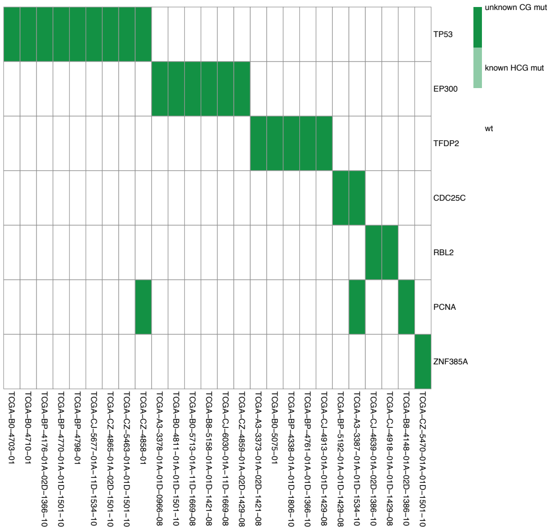


**Regulation Of IFNA Signaling**
**FDR = 1.1%, FDR nod = 2.1%**

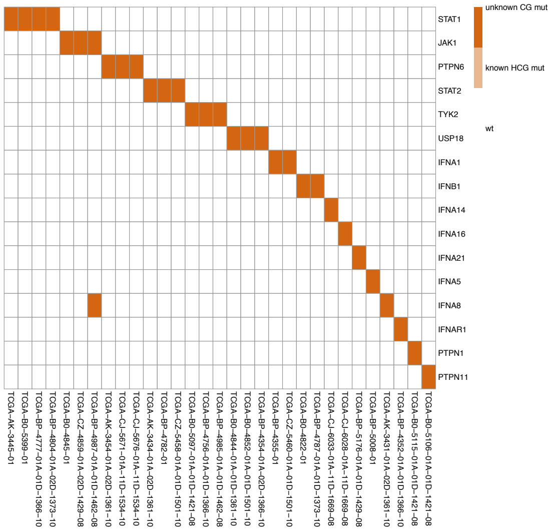

### LUAD

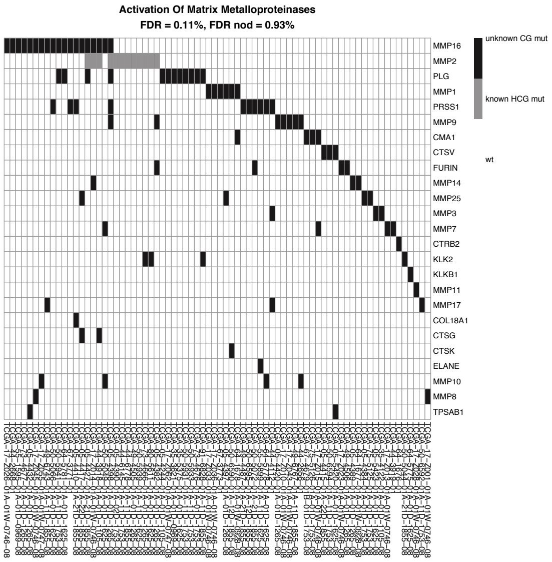
Activation Of Matrix Metalloproteinases
FDR = 0.11%, FDR nod = 0.93%
MMP16
MMP2
PLG
MMP1
PRSS1
MMP9
CMA1
CTSV
FURIN
MMP14
MMP25
MMP3
MMP7
CTRB2
KLK2
KLKB1
MMP11
MMP17
COL18A1
CTSG
CTSK
ELANE
MMP10
MMP8
TPSAB1
unknown CG mut
known HCG mut
wt

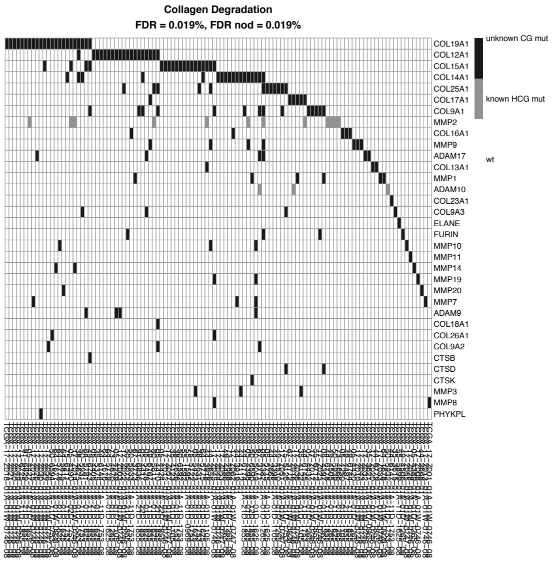
Collagen Degradation
FDR = 0.019%, FDR nod = 0.019%
COL19A1
COL12A1
COL15A1
COL14A1
COL25A1
COL17A1
COL9A1
MMP2
COL16A1
MMP9
ADAM17
COL13A1
MMP1
ADAM10
COL23A1
COL9A3
ELANE
FURIN
MMP10
MMP11
MMP14
MMP19
MMP20
MMP7
ADAM9
COL18A1
COL26A1
COL9A2
CTSB
CTSD
CTSK
MMP3
MMP8
PHYKPL
unknown CG mut
known HCG mut
wt

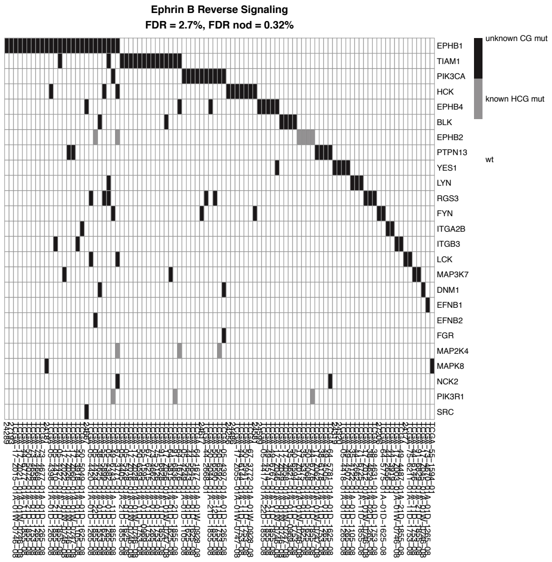
Ephrin B Reverse Signaling
FDR = 2.7%, FDR nod = 0.32%
EPHB1
TIAM1
PIK3CA
HCK
EPHB4
BLK
EPHB2
PTPN13
YES1
LYN
RGS3
FYN
ITGA2B
ITGB3
LCK
MAP3K7
DNM1
EFNB1
EFNB2
FGR
MAP2K4
MAPK8
NCK2
PIK3R1
SRC
unknown CG mut
known HCG mut
wt

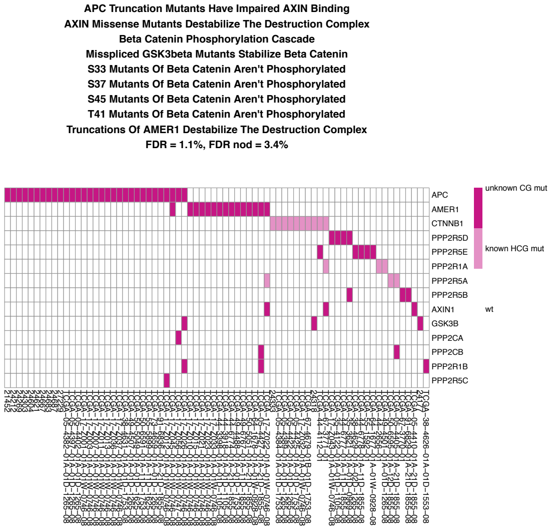
APC Truncation Mutants Have Impaired AXIN Binding
AXIN Missense Mutants Destabilize The Destruction Complex
Beta Catenin Phosphorylation Cascade
Misspliced GSK3beta Mutants Stabilize Beta Catenin
S33 Mutants Of Beta Catenin Aren't Phosphorylated
S37 Mutants Of Beta Catenin Aren't Phosphorylated
S45 Mutants Of Beta Catenin Aren't Phosphorylated
T41 Mutants Of Beta Catenin Aren't Phosphorylated
Truncations Of AMER1 Destabilize The Destruction Complex
FDR = 1.1%, FDR nod = 3.4%
APC
AMER1
CTNNB1
PPP2R5D
PPP2R5E
PPP2R1A
PPP2R5A
PPP2R5B
AXIN1
GSK3B
PPP2CA
PPP2CB
PPP2R1B
PPP2R5C
unknown CG mut
known HCG mut
wt

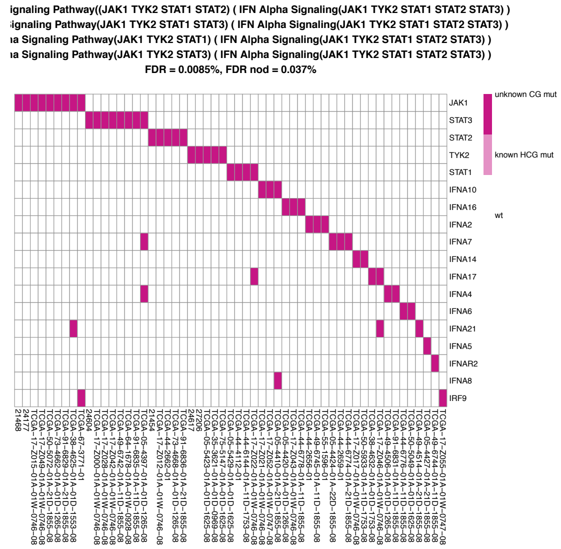
ignaling Pathway((JAK1 TYK2 STAT1 STAT2) ( IFN Alpha Signaling(JAK1 TYK2 STAT1 STAT2 STAT3) )
ignaling Pathway(JAK1 TYK2 STAT1 STAT3) ( IFN Alpha Signaling(JAK1 TYK2 STAT1 STAT2 STAT3) )
a Signaling Pathway(JAK1 TYK2 STAT1) ( IFN Alpha Signaling(JAK1 TYK2 STAT1 STAT2 STAT3) )
a Signaling Pathway(JAK1 TYK2 STAT3) ( IFN Alpha Signaling(JAK1 TYK2 STAT1 STAT2 STAT3) )
FDR = 0.0085%, FDR nod = 0.037%
JAK1
STAT3
STAT2
TYK2
STAT1
IFNA10
IFNA16
IFNA2
IFNA7
IFNA14
IFNA17
IFNA4
IFNA6
IFNA21
IFNA5
IFNAR2
IFNA8
IRF9
unknown CG mut
known HCG mut
wt

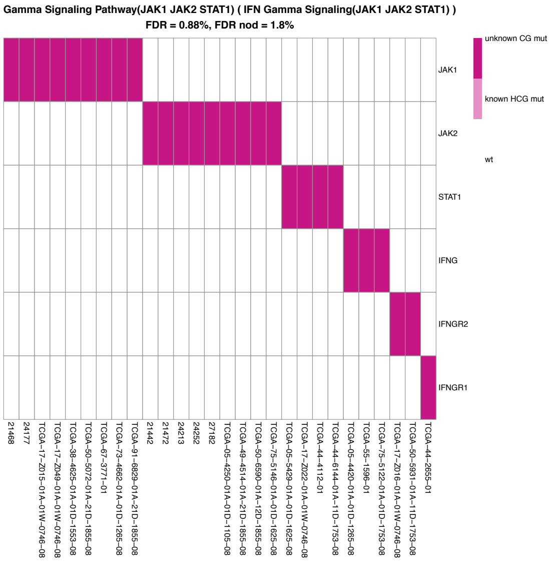
Gamma Signaling Pathway(JAK1 JAK2 STAT1) ( IFN Gamma Signaling(JAK1 JAK2 STAT1) )
FDR = 0.88%, FDR nod = 1.8%
JAK1
JAK2
STAT1
IFNG
IFNGR2
IFNGR1
unknown CG mut
known HCG mut
wt

**IRF3 Mediated Induction Of Type I IFN**
**FDR = 0.02%, FDR nod = 0.074%**

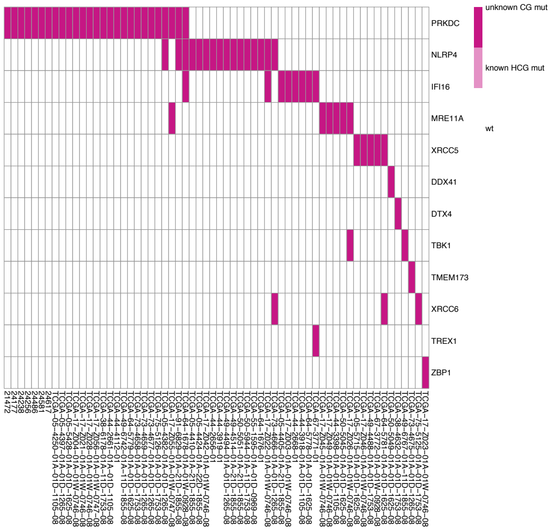


**Regulation Of IFNA Signaling**
**FDR = 0.0013%, FDR nod = 0.019%**

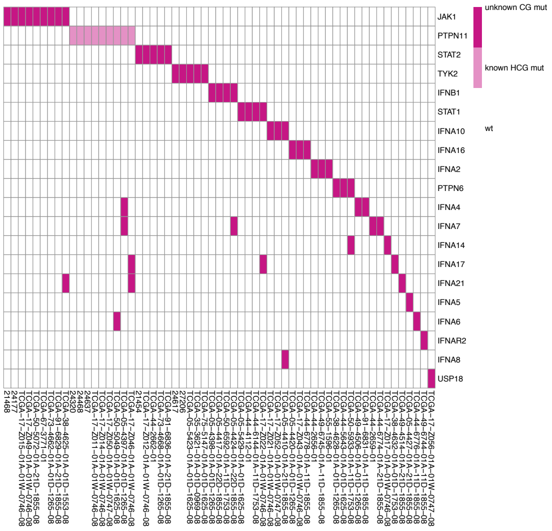


**DNA Damage/Telomere Stress Induced Senescence**
**FDR = 2e−12%, FDR nod = 0.0053%**

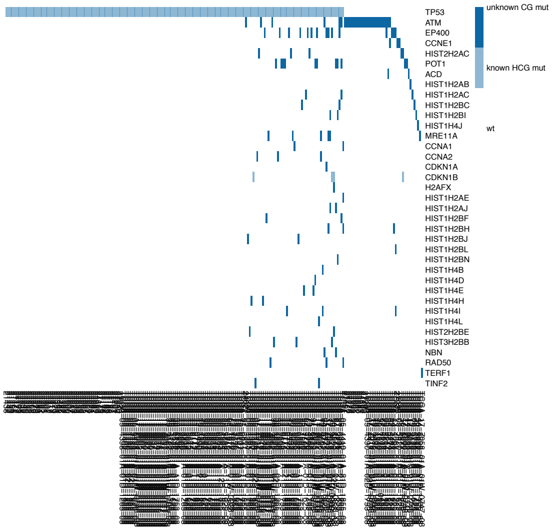


**Packaging Of Telomere Ends**
**FDR = 0.07%, FDR nod = 0.2%**

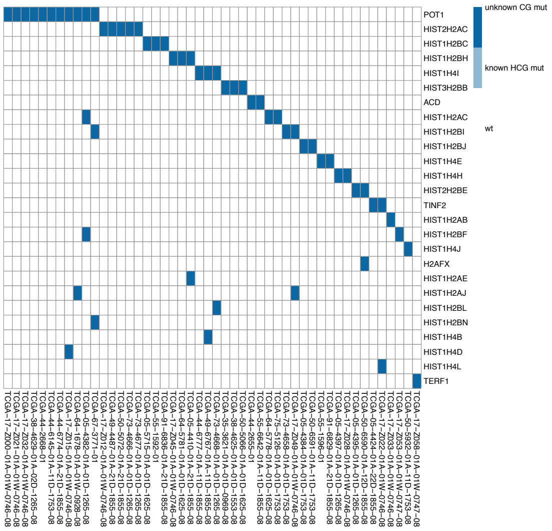


**APC Truncation Mutants Have Impaired AXIN Binding**
**AXIN Missense Mutants Destabilize The Destruction Complex**
**Beta Catenin Phosphorylation Cascade**
**Misspliced GSK3beta Mutants Stabilize Beta Catenin**
**S33 Mutants Of Beta Catenin Aren't Phosphorylated**
**S37 Mutants Of Beta Catenin Aren't Phosphorylated**
**S45 Mutants Of Beta Catenin Aren't Phosphorylated**
**T41 Mutants Of Beta Catenin Aren't Phosphorylated**
**Truncations Of AMER1 Destabilize The Destruction Complex**
**FDR = 1.1%, FDR nod = 3.4%**

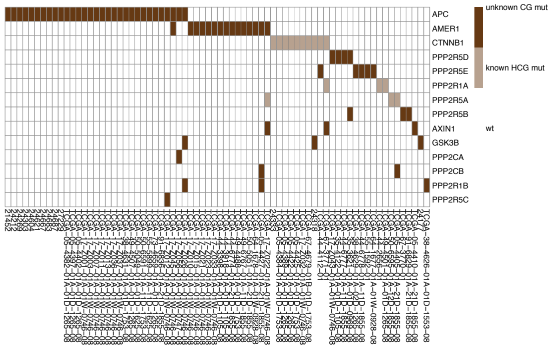


**AUF1 (hnRNP D0) Binds And Destabilizes MRNA**
**Autodegradation Of The E3 Ubiquitin Ligase COP1**
**CDK Mediated Phosphorylation And Removal Of Cdc6**
**CDT1 Association With The CDC6:ORC:origin Complex**
**Cross Presentation Of Soluble Exogenous Antigens (endosomes)**
**Dectin 1 Mediated Noncanonical NF KB Signaling**
**Degradation Of AXIN**
**Degradation Of DVL**
**Degradation Of GLI1 By The Proteasome**
**Degradation Of GLI2 By The Proteasome**
**G2/M Checkpoints**
**GLI3 Is Processed To GLI3R By The Proteasome**
**Hedgehog Ligand Biogenesis**
**Hh Mutants That Don't Undergo Autocatalytic Processing Are Degraded By ERAD**
**NIK >noncanonical NF KB Signaling**
**Orc1 Removal From Chromatin**
**Regulation Of Activated PAK 2p34 By Proteasome Mediated Degradation**
**Regulation Of Ornithine Decarboxylase (ODC)**
**SCF Beta TrCP Mediated Degradation Of Emi1**
**The Role Of GTSE1 In G2/M Progression After G2 Checkpoint**
**Ubiquitin Dependent Degradation Of Cyclin D1**
**Ubiquitin Mediated Degradation Of Phosphorylated Cdc25A**

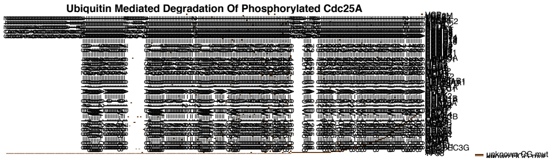

P53 Pathway
FDR = 2e−12%, FDR nod = 1%

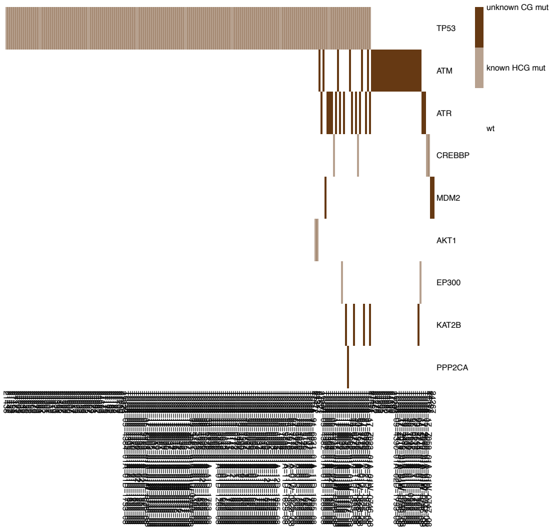


P53 Pathway Feedback Loops 2
FDR = 2e−12%, FDR nod = 2.9e−05%

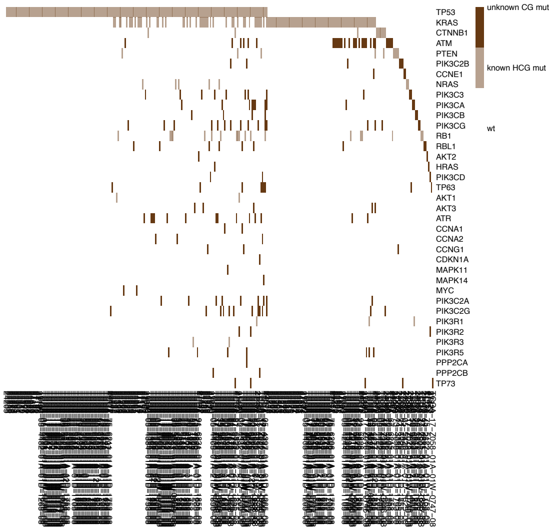


Regulation Of TP53 Activity Through Phosphorylation
FDR = 2e−12%, FDR nod = 1.1%

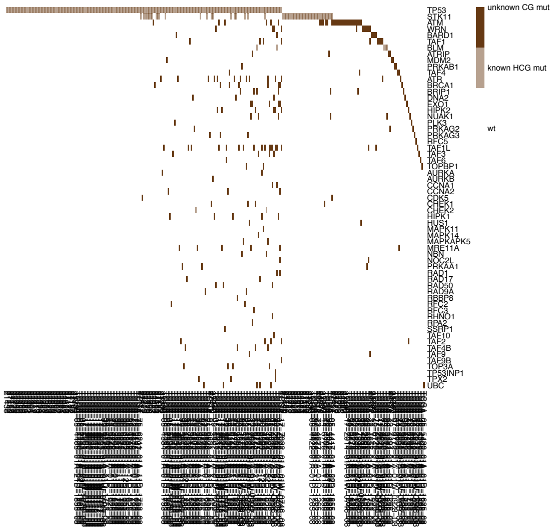


Stabilization Of P53
FDR = 2e−12%, FDR nod = 1.2%

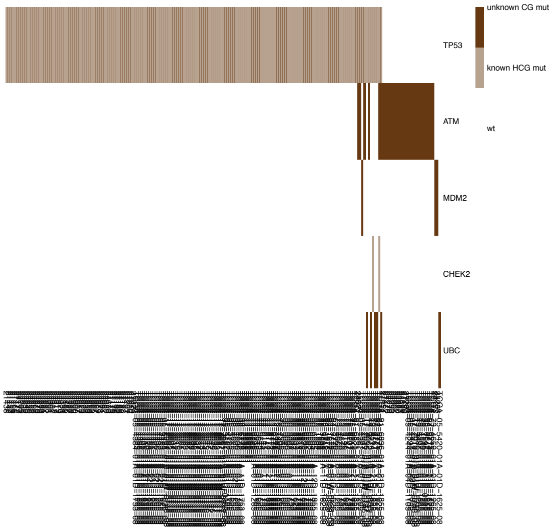


TP53 Regulates Transcription Of Caspase Activators And Caspases
FDR = 2e−12%, FDR nod = 0.14%

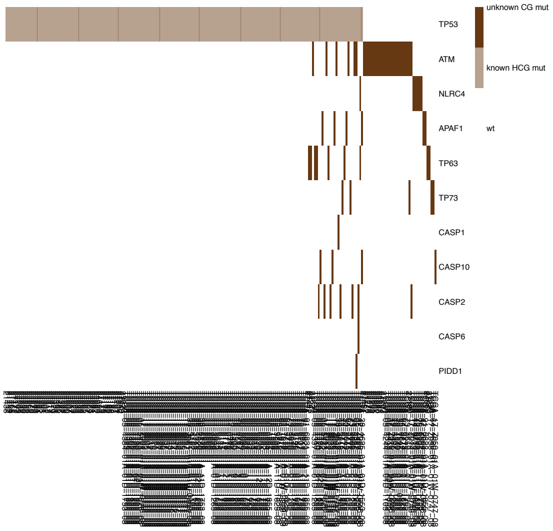


BARD1 Signaling Events
FDR = 2e−12%, FDR nod = 0.031%

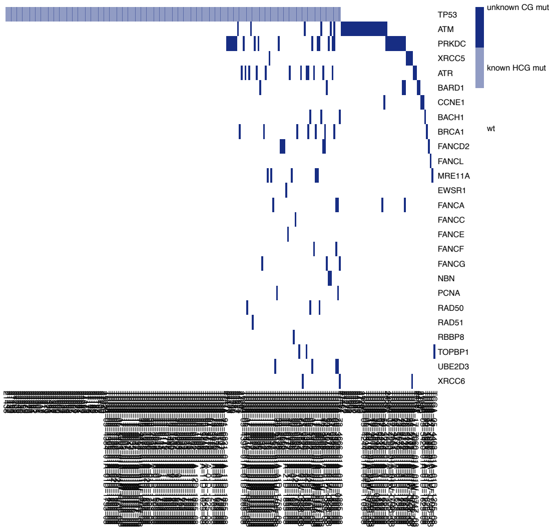

**DNA Damage/Telomere Stress Induced Senescence**
**FDR = 2e−12%, FDR nod = 0.0053%**

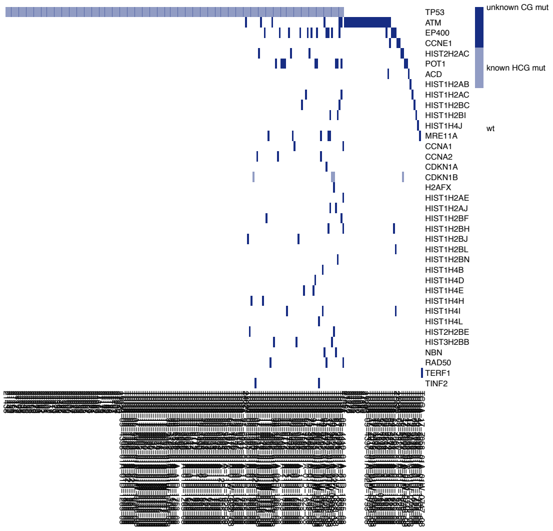


**E2F Transcription Factor Network**
**FDR = 9.5e−07%, FDR nod = 1.1%**

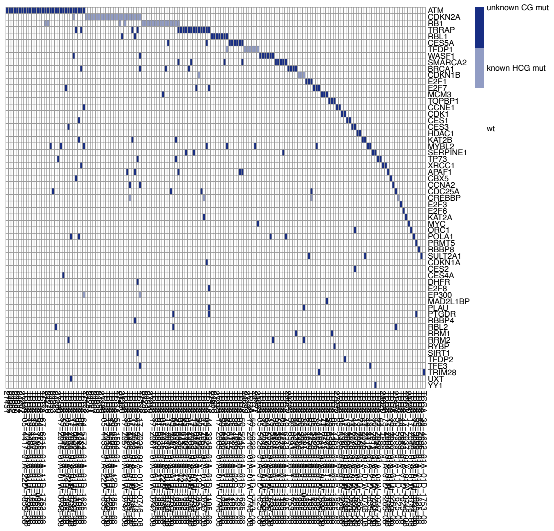


**HDR Through Homologous Recombination (HRR)**
**FDR = 1.2%, FDR nod = 1.3%**

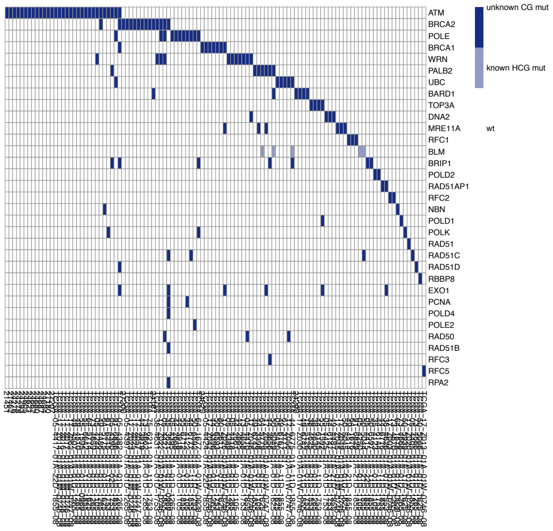


**HDR Through Single Strand Annealing (SSA)**
**FDR = 0.14%, FDR nod = 0.41%**

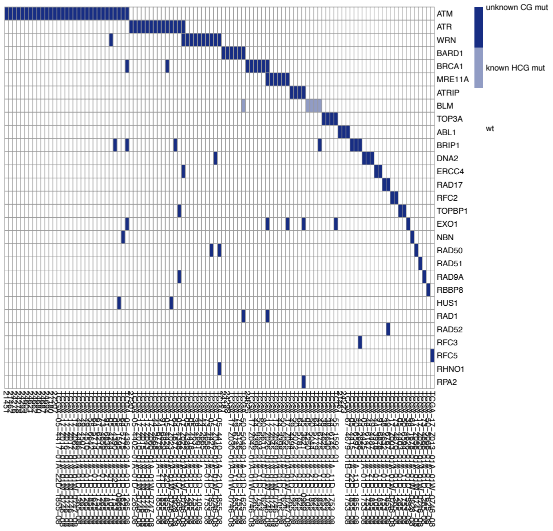


**Homologous DNA Pairing And Strand Exchange**
**esolution Of D Loop Structures Through Synthesis Dependent Strand Annealing (SDSA)**
**FDR = 1.1%, FDR nod = 1.1%**

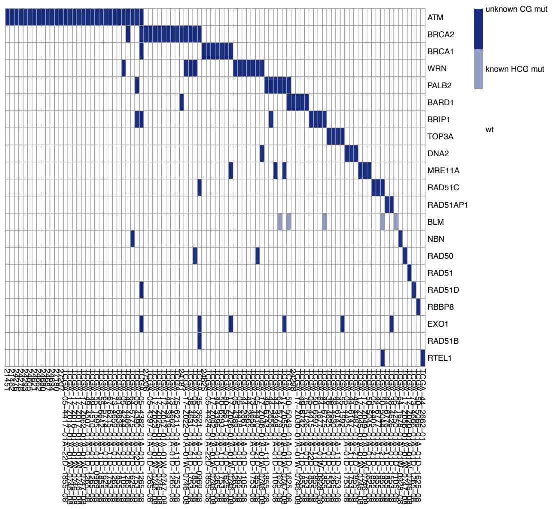


**Integrated Breast Cancer Pathway**
**FDR = 2e−12%, FDR nod = 0.23%**

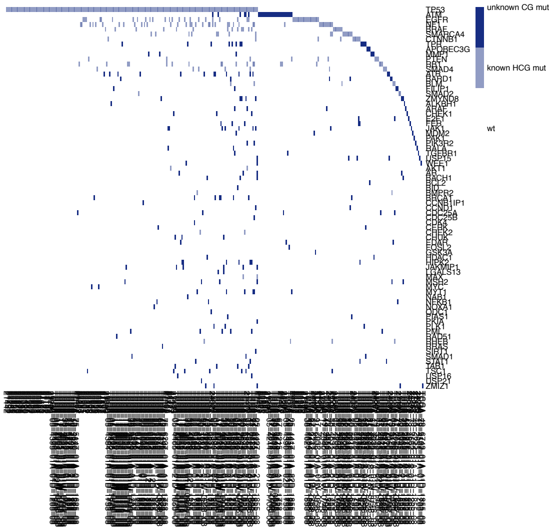

**Integrated Cancer Pathway**
**FDR = 2e−12%, FDR nod = 0.23%**

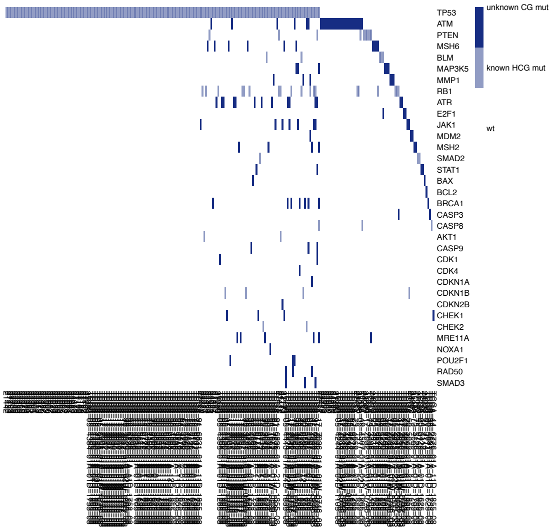


**Meiotic Recombination**
**FDR = 8.2e−06%, FDR nod = 2.7e−05%**

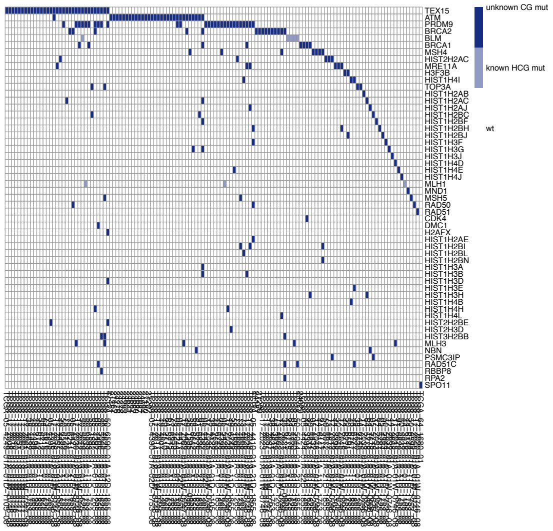


**Meiotic Synapsis**
**FDR = 5.3%, FDR nod = 1.4%**

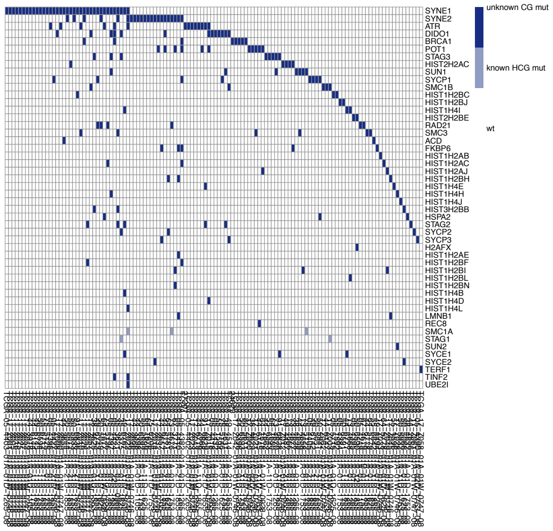


**Nonhomologous End Joining (NHEJ)**
**FDR = 0.35%, FDR nod = 0.23%**

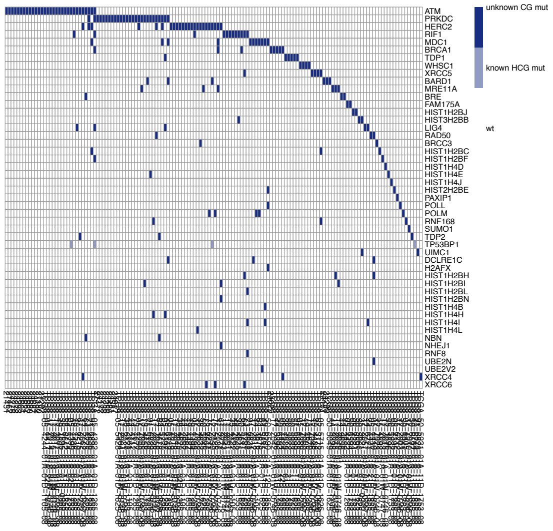


**P73 Transcription Factor Network**
**FDR = 0.11%, FDR nod = 1.2%**

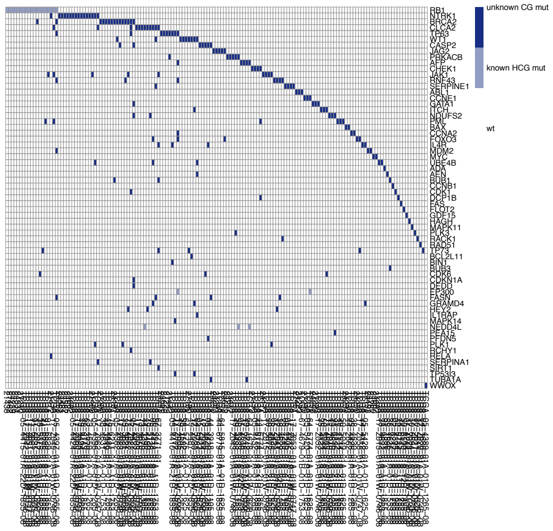


**Presynaptic Phase Of Homologous DNA Pairing And Strand Exchange**
**FDR = 0.13%, FDR nod = 0.23%**

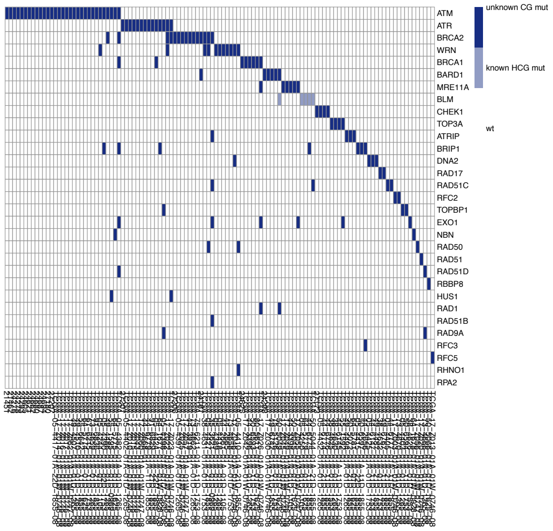

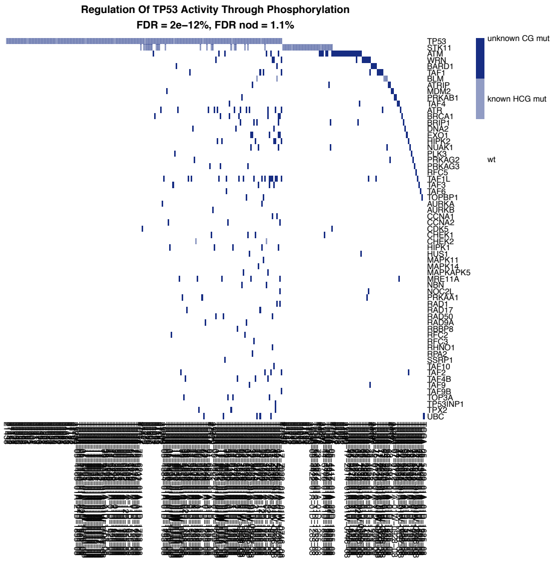

Regulation Of TP53 Activity Through Phosphorylation
FDR = 2e−12%, FDR nod = 1.1%
unknown CG mut
known HCG mut
wt
TP53
STK11
ATM
WRN
BARD1
TAF1
BLM
ATRIP
MDM2
PRKAB1
TAF4
ATR
BRCA1
BRIP1
DNA2
EXO1
HIPK2
NUAK1
PLK3
PRKAG2
PRKAG3
RFC5
TAF1L
TAF3
TAF6
TOPBP1
AURKA
AURKB
CCNA1
CCNA2
CDK5
CHEK1
CHEK2
HIPK1
HUS1
MAPK11
MAPK14
MAPKAPK5
MRE11A
NBN
NOC2L
PRKAA1
RAD1
RAD17
RAD50
RAD9A
RBBP8
RFC2
RFC3
RHNO1
RPA2
SSRP1
TAF10
TAF2
TAF4B
TAF9
TAF9B
TOP3A
TP53INP1
TPX2
UBC


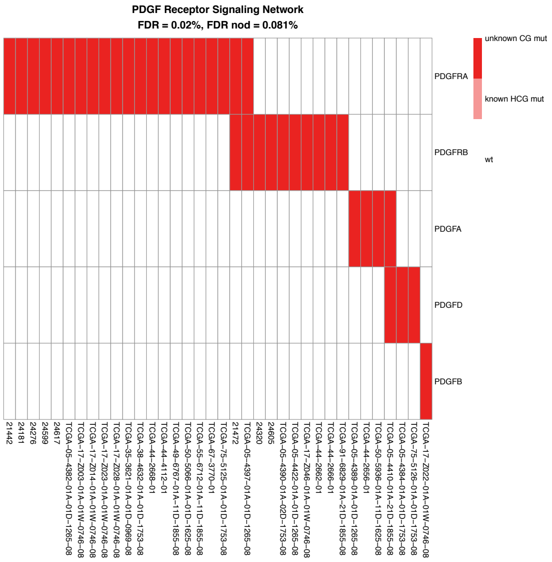

PDGF Receptor Signaling Network
FDR = 0.02%, FDR nod = 0.081%
unknown CG mut
known HCG mut
wt
PDGFRA
PDGFRB
PDGFA
PDGFD
PDGFB


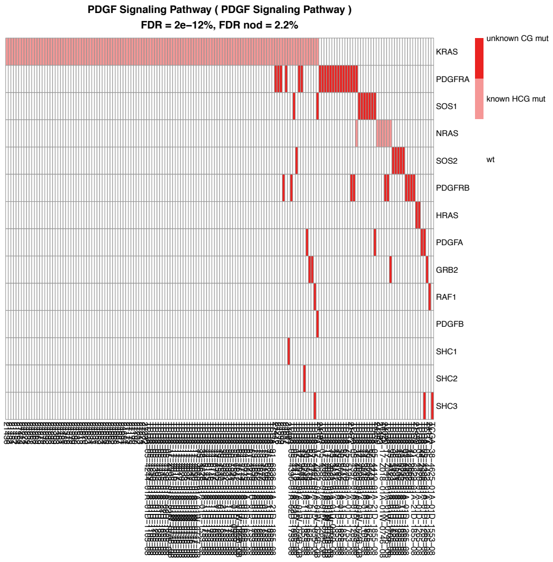

PDGF Signaling Pathway ( PDGF Signaling Pathway )
FDR = 2e−12%, FDR nod = 2.2%
unknown CG mut
known HCG mut
wt
KRAS
PDGFRA
SOS1
NRAS
SOS2
PDGFRB
HRAS
PDGFA
GRB2
RAF1
PDGFB
SHC1
SHC2
SHC3


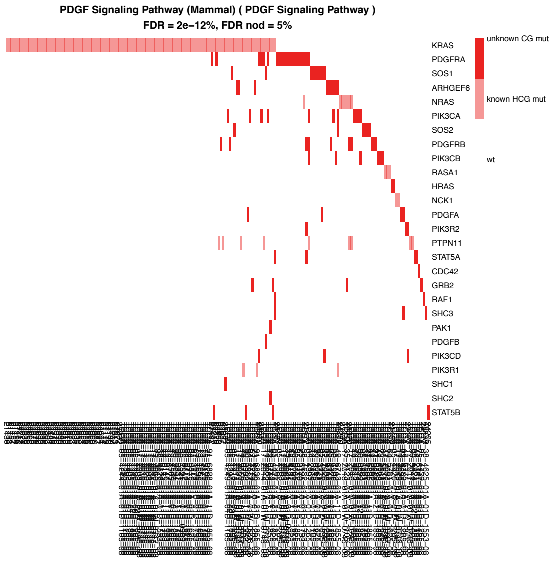

PDGF Signaling Pathway (Mammal) ( PDGF Signaling Pathway )
FDR = 2e−12%, FDR nod = 5%
unknown CG mut
known HCG mut
wt
KRAS
PDGFRA
SOS1
ARHGEF6
NRAS
PIK3CA
SOS2
PDGFRB
PIK3CB
RASA1
HRAS
NCK1
PDGFA
PIK3R2
PTPN11
STAT5A
CDC42
GRB2
RAF1
SHC3
PAK1
PDGFB
PIK3CD
PIK3R1
SHC1
SHC2
STAT5B


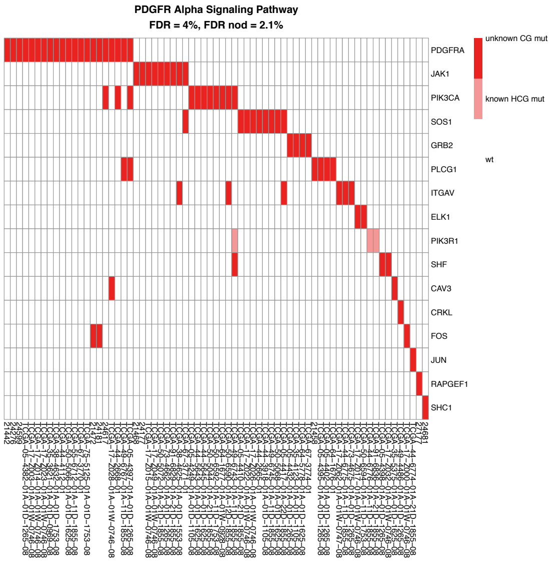

PDGFR Alpha Signaling Pathway
FDR = 4%, FDR nod = 2.1%
unknown CG mut
known HCG mut
wt
PDGFRA
JAK1
PIK3CA
SOS1
GRB2
PLCG1
ITGAV
ELK1
PIK3R1
SHF
CAV3
CRKL
FOS
JUN
RAPGEF1
SHC1


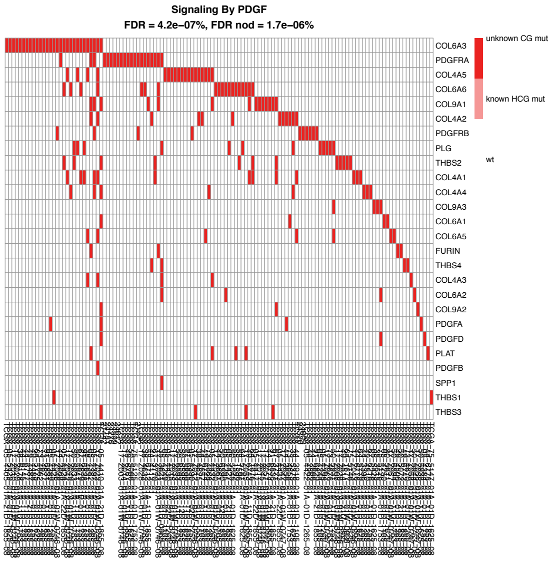

Signaling By PDGF
FDR = 4.2e−07%, FDR nod = 1.7e−06%
unknown CG mut
known HCG mut
wt
COL6A3
PDGFRA
COL4A5
COL6A6
COL9A1
COL4A2
PDGFRB
PLG
THBS2
COL4A1
COL4A4
COL9A3
COL6A1
COL6A5
FURIN
THBS4
COL4A3
COL6A2
COL9A2
PDGFA
PDGFD
PLAT
PDGFB
SPP1
THBS1
THBS3

**VEGF And VEGFR Signaling Network**
**VEGF Binds To VEGFR Leading To Receptor Dimerization**
**FDR = 1.7e−08%, FDR nod = 0.2%**

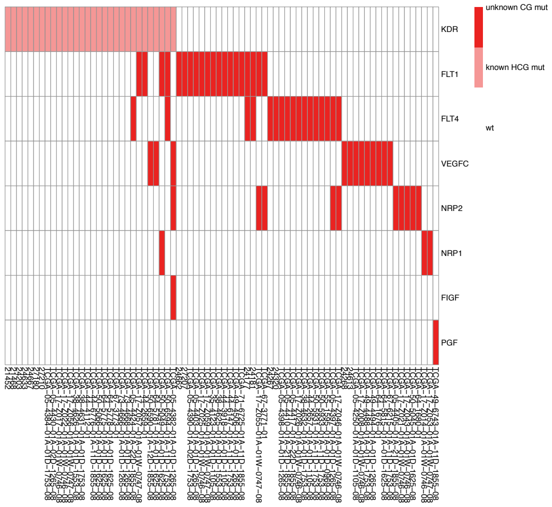


**VEGF Signaling Pathway**
**FDR = 9e−06%, FDR nod = 0.14%**

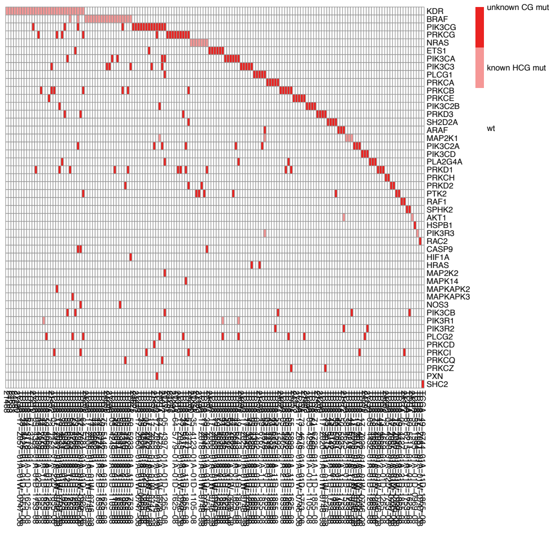


**VEGFR3 Signaling In Lymphatic Endothelium**
**FDR = 5.6%, FDR nod = 0.73%**

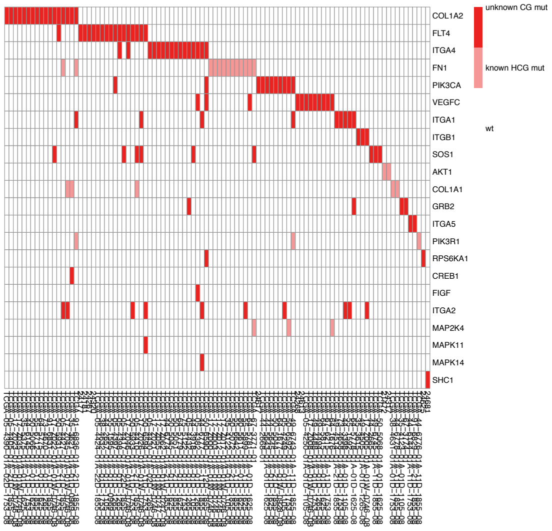


**Homologous DNA Pairing And Strand Exchange**
**esolution Of D Loop Structures Through Synthesis Dependent Strand Annealing (SDSA)**
**FDR = 1.1%, FDR nod = 1.1%**

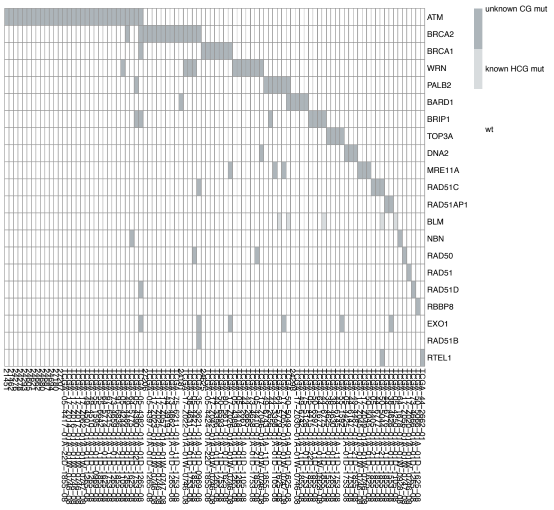


**Presynaptic Phase Of Homologous DNA Pairing And Strand Exchange**
**FDR = 0.13%, FDR nod = 0.23%**

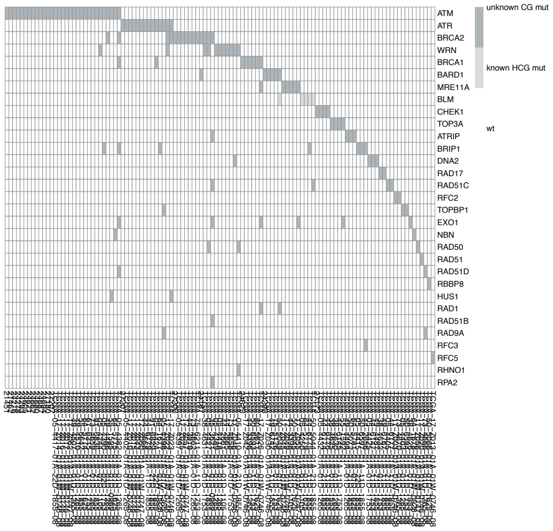


**TRIF Mediated Programmed Cell Death**
**FDR = 0.00019%, FDR nod = 2.7e−05%**

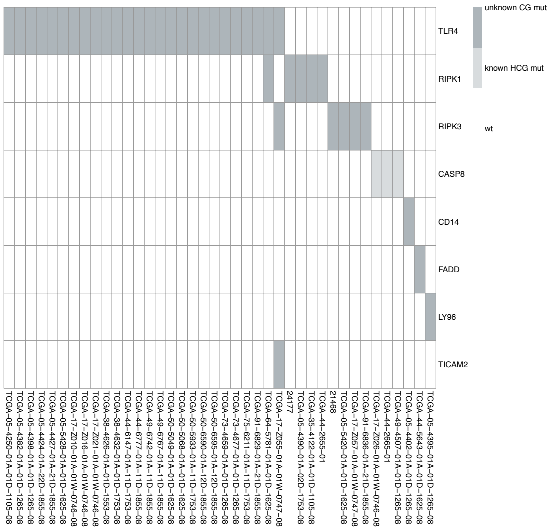

**ARMS Mediated Activation**
**FDR = 0.00051%, FDR nod = 2.1%**

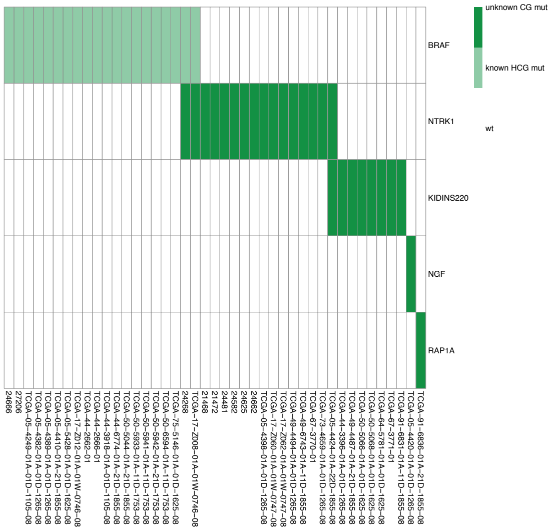


**Constitutive Signaling By Aberrant PI3K In Cancer**
**FDR = 2e−12%, FDR nod = 0.08%**

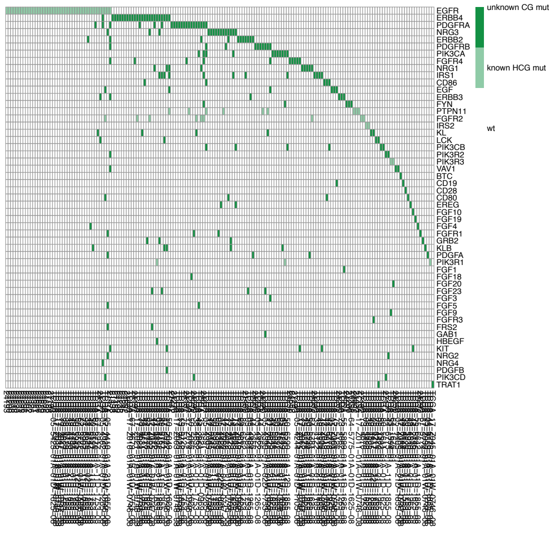


**ErbB Receptor Signaling Network**
**FDR = 2.3e−12%, FDR nod = 0.16%**

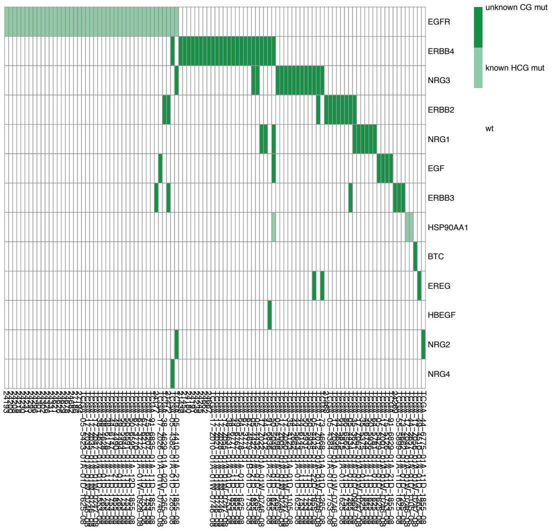


**ERBB2 Activates PTK6 Signaling**
**Signaling By ERBB4**
**FDR = 2.2e−12%, FDR nod = 0.2%**

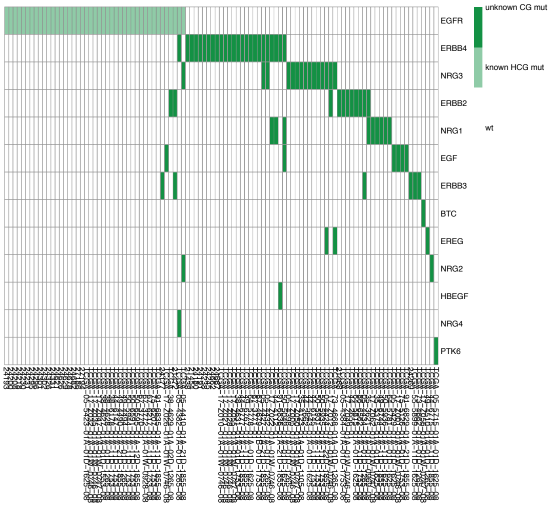


**ERBB2 Regulates Cell Motility**
**FDR = 9.4e−12%, FDR nod = 1%**

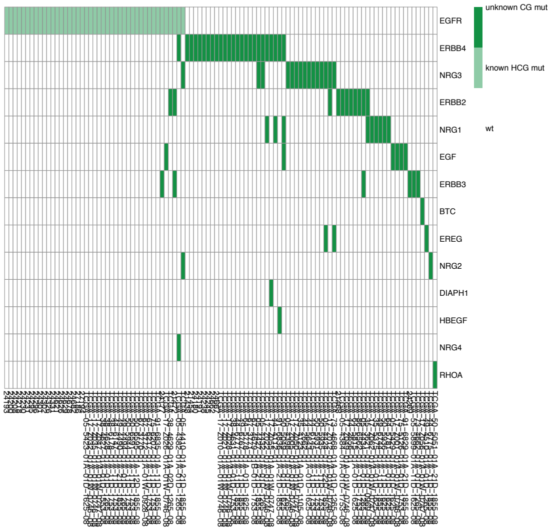


**ErbB4 Signaling Events**
**FDR = 3%, FDR nod = 2.1%**

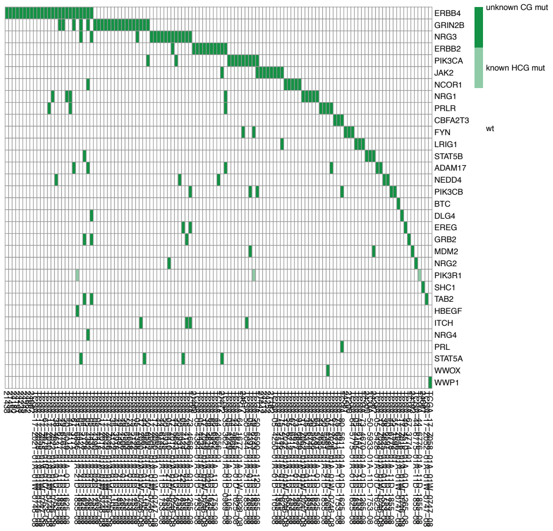

**G1 To S Cell Cycle Control**
**FDR = 2e−12%, FDR nod = 0.046%**

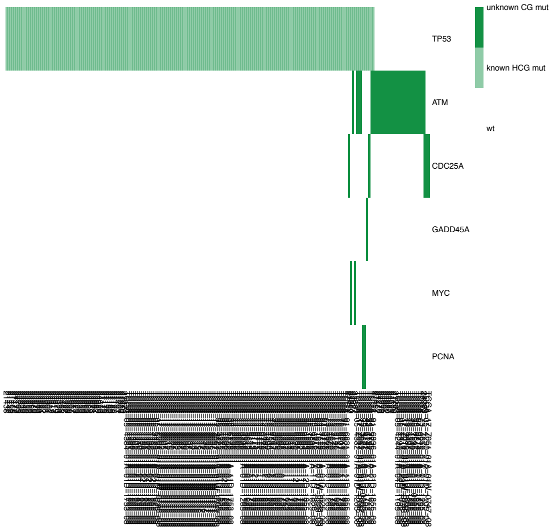


**thway (through Glutamate, G Alpha Q And PLC Beta) ( GPCR GroupI Metabotropic Glutamate Receptor Signaling I**
**FDR = 0.00071%, FDR nod = 0.46%**

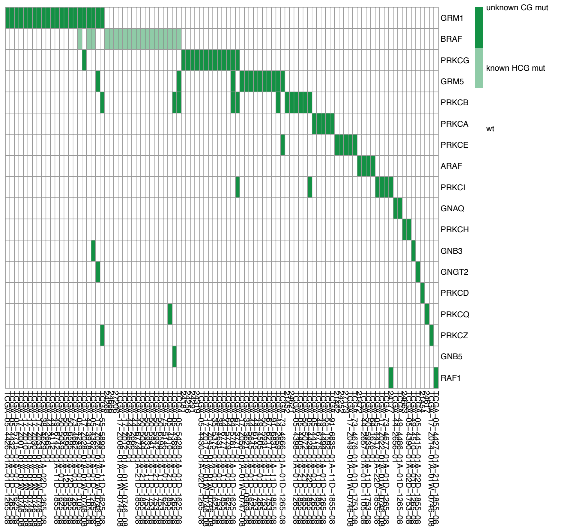


**Integrated Breast Cancer Pathway**
**FDR = 2e−12%, FDR nod = 0.23%**

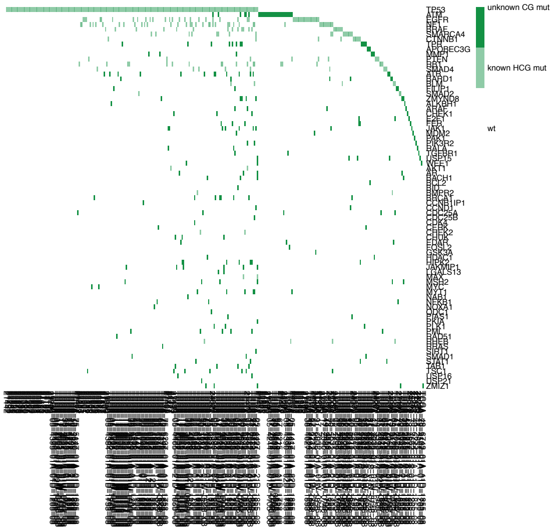


**Interleukin Signaling Pathway**
**FDR = 0.24%, FDR nod = 4.8%**

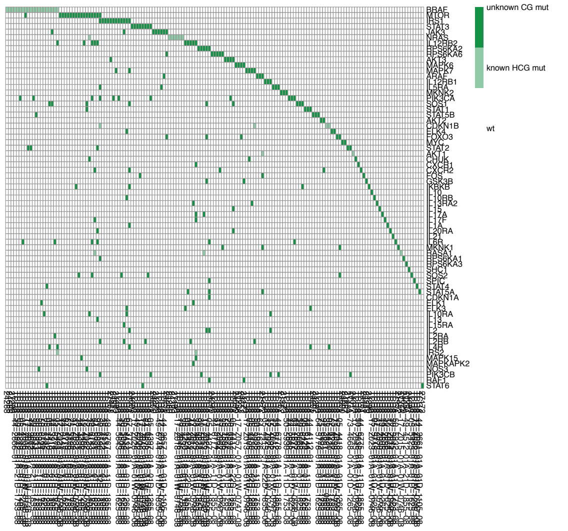


**MAPK1 (ERK2) Activation**
**FDR = 0.23%, FDR nod = 2.8%**

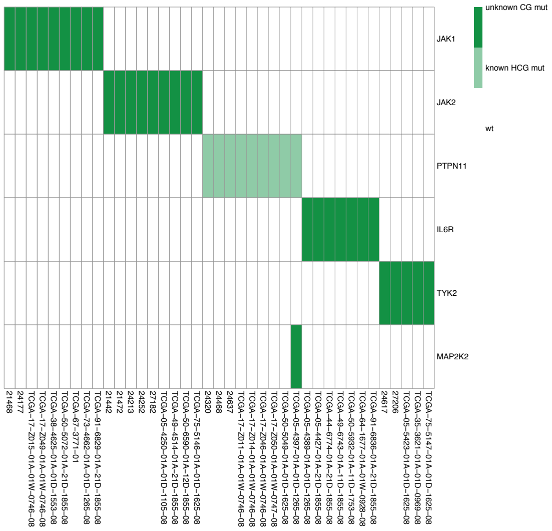


**MAPK3 (ERK1) Activation**
**FDR = 0.012%, FDR nod = 1%**

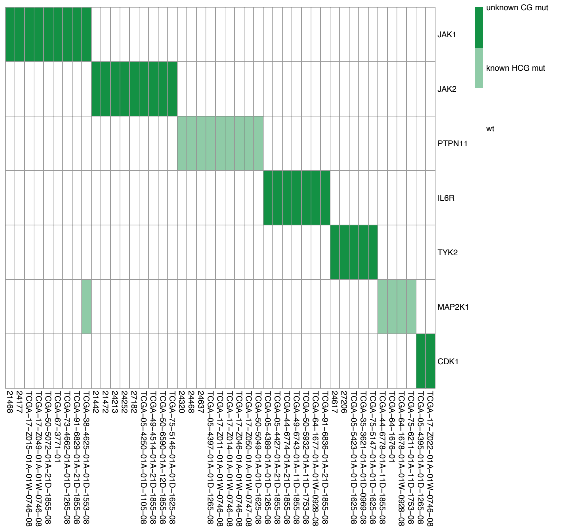

**PI3K Events In ERBB2 Signaling**
**FDR = 9.6e−12%, FDR nod = 0.18%**

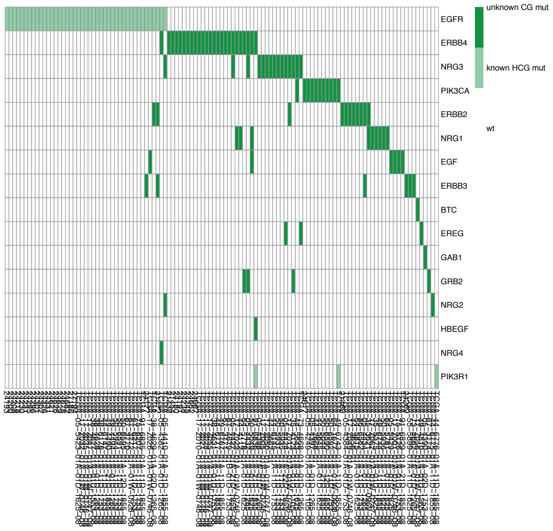


**PI3K Events In ERBB4 Signaling**
**FDR = 0.56%, FDR nod = 0.091%**

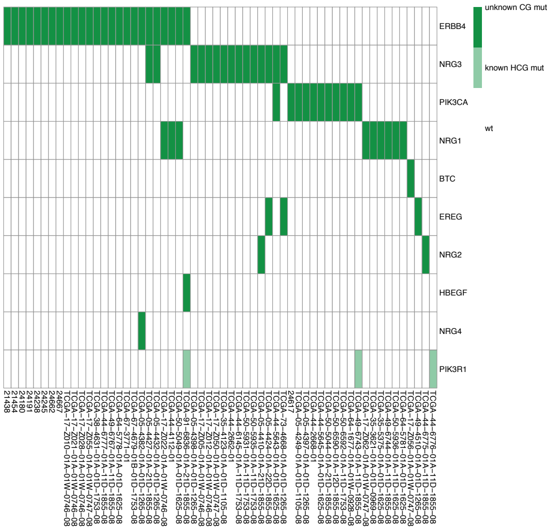


**PI3K/AKT Activation**
**FDR = 5.5%, FDR nod = 0.23%**

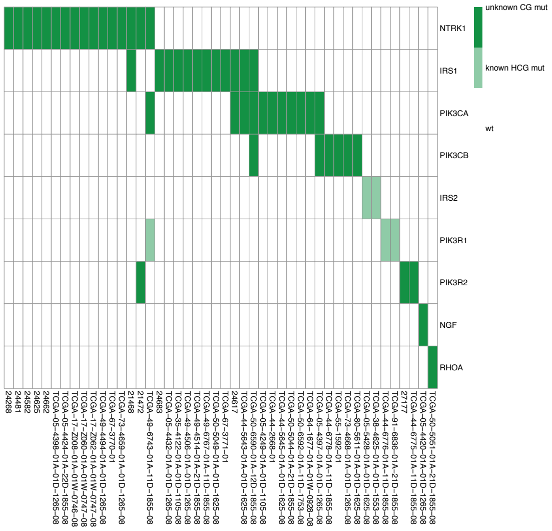


**PIP3 Activates AKT Signaling**
**FDR = 1.6e−10%, FDR nod = 1.4%**

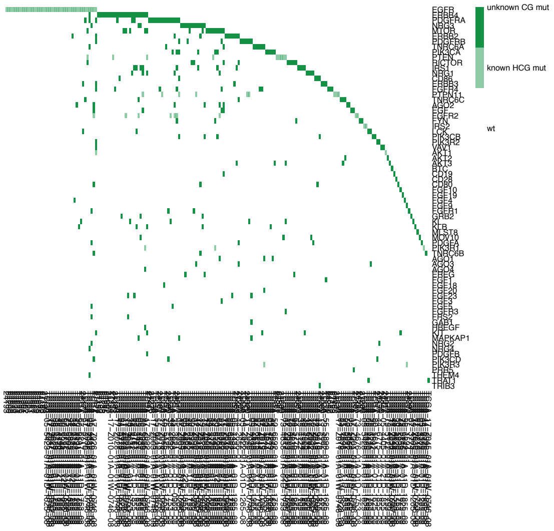


**Prolactin**
**FDR = 2e−12%, FDR nod = 0.00055%**

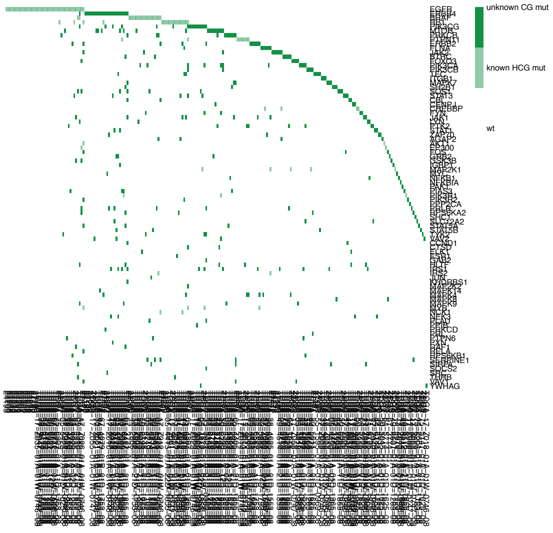


**Signalling To P38 Via RIT And RIN**
**FDR = 1e−09%, FDR nod = 1.1e−05%**

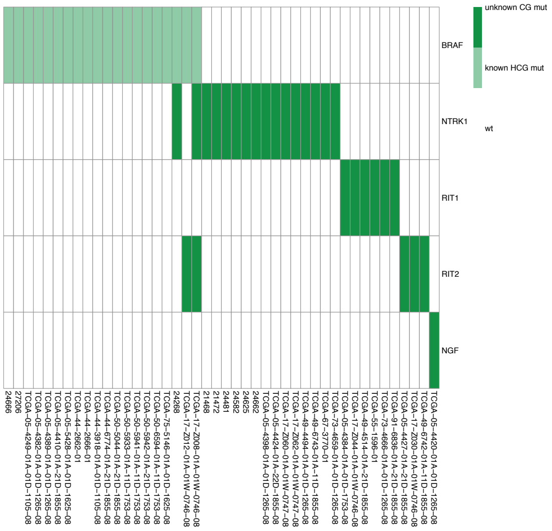

**VEGF Signaling Pathway**
**FDR = 9e−06%, FDR nod = 0.14%**

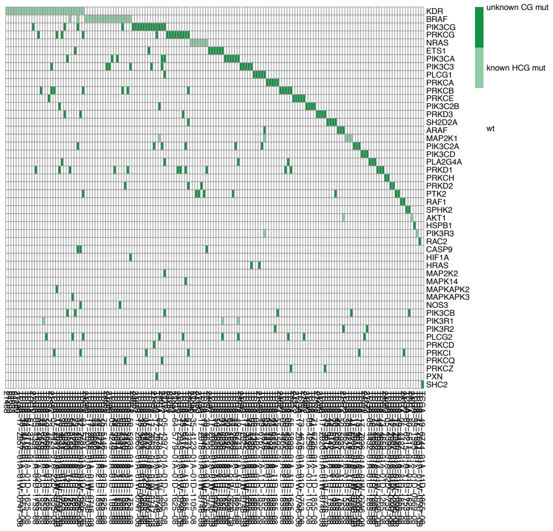


**ignaling Pathway((JAK1 TYK2 STAT1 STAT2) ( IFN Alpha Signaling(JAK1 TYK2 STAT1 STAT2 STAT3) )**
**ignaling Pathway(JAK1 TYK2 STAT1 STAT3) ( IFN Alpha Signaling(JAK1 TYK2 STAT1 STAT2 STAT3) )**
**a Signaling Pathway(JAK1 TYK2 STAT1) ( IFN Alpha Signaling(JAK1 TYK2 STAT1 STAT2 STAT3) )**
**a Signaling Pathway(JAK1 TYK2 STAT3) ( IFN Alpha Signaling(JAK1 TYK2 STAT1 STAT2 STAT3) )**
**FDR = 0.0085%, FDR nod = 0.037%**

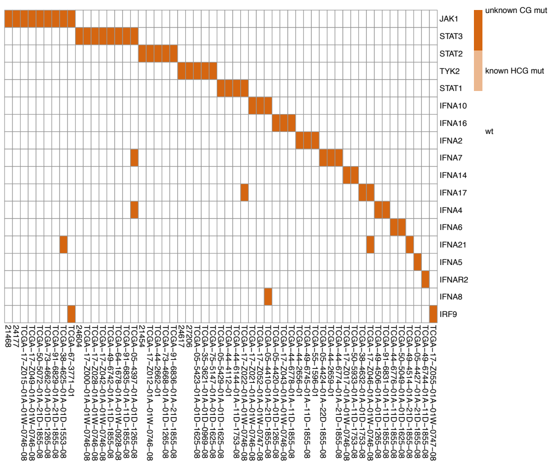


**Gamma Signaling Pathway(JAK1 JAK2 STAT1) ( IFN Gamma Signaling(JAK1 JAK2 STAT1) )**
**FDR = 0.88%, FDR nod = 1.8%**

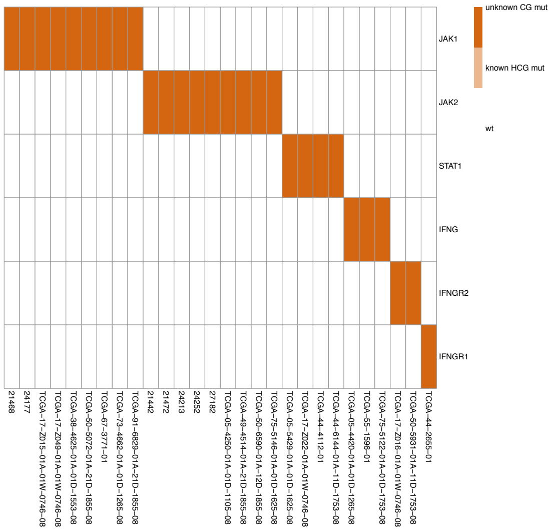


**IL 6 Type Cytokine Receptor Ligand Interactions**
**FDR = 0.74%, FDR nod = 1.4%**

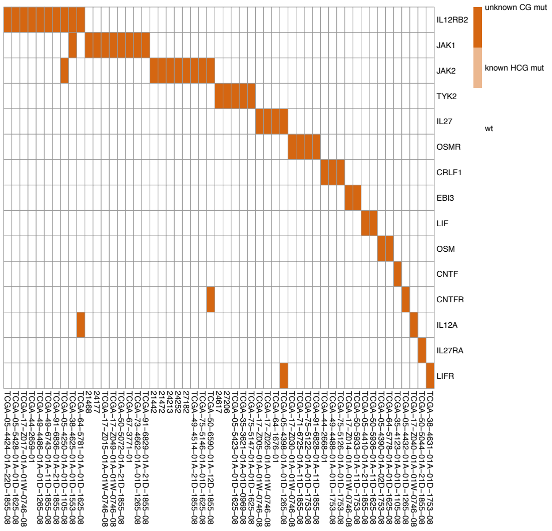


**IRF3 Mediated Induction Of Type I IFN**
**FDR = 0.02%, FDR nod = 0.074%**

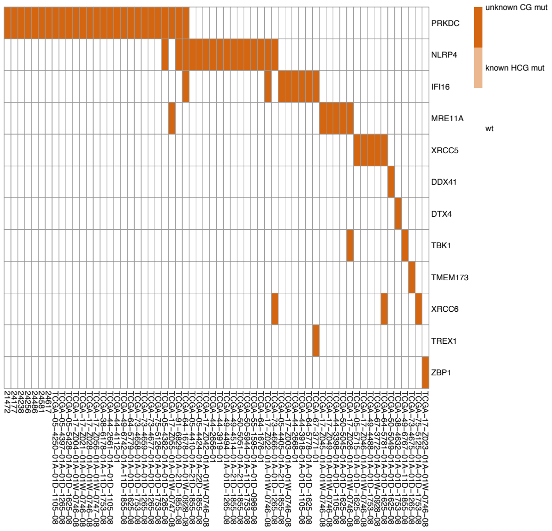


**Regulation Of IFNA Signaling**
**FDR = 0.0013%, FDR nod = 0.019%**

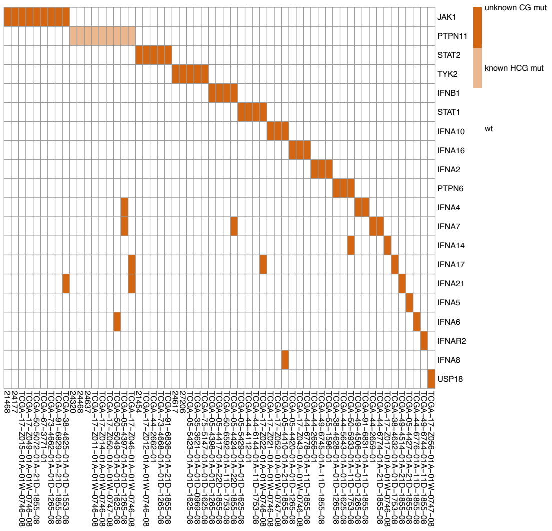

### PRAD

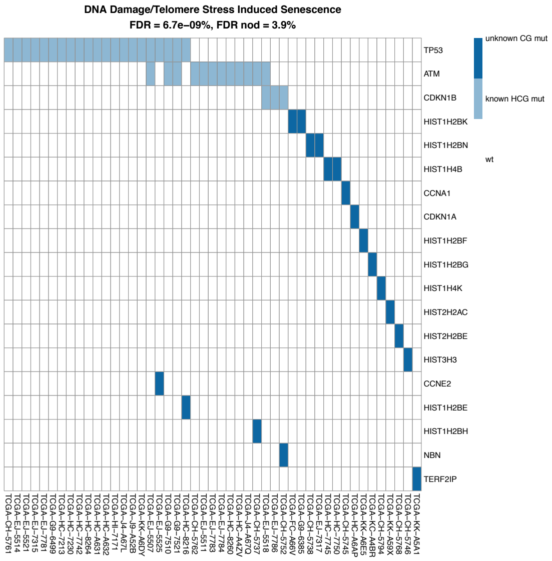
DNA Damage/Telomere Stress Induced Senescence
FDR = 6.7e−09%, FDR nod = 3.9%
TP53
ATM
CDKN1B
HIST1H2BK
HIST1H2BN
HIST1H4B
CCNA1
CDKN1A
HIST1H2BF
HIST1H2BG
HIST1H4K
HIST2H2AC
HIST2H2BE
HIST3H3
CCNE2
HIST1H2BE
HIST1H2BH
NBN
TERF2IP
unknown CG mut
known HCG mut
wt

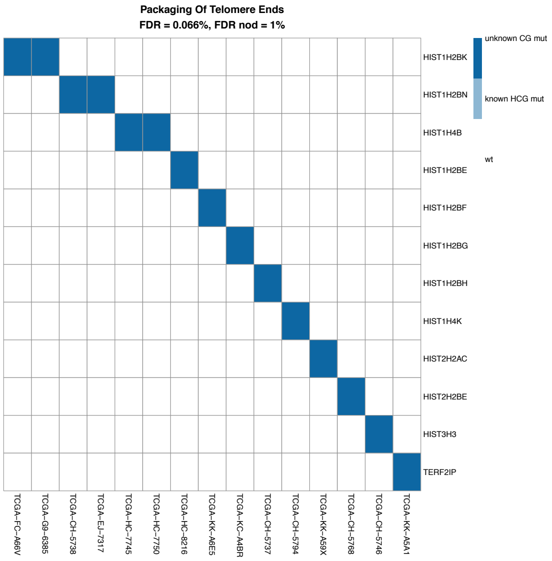
Packaging Of Telomere Ends
FDR = 0.066%, FDR nod = 1%
HIST1H2BK
HIST1H2BN
HIST1H4B
HIST1H2BE
HIST1H2BF
HIST1H2BG
HIST1H2BH
HIST1H4K
HIST2H2AC
HIST2H2BE
HIST3H3
TERF2IP
unknown CG mut
known HCG mut
wt
TCGA−FC−A66V
TCGA−G9−6385
TCGA−CH−5738
TCGA−EJ−7317
TCGA−HC−7745
TCGA−HC−7750
TCGA−HC−8216
TCGA−KK−A8E5
TCGA−KC−A4BR
TCGA−CH−5737
TCGA−CH−5794
TCGA−KK−A59X
TCGA−CH−5768
TCGA−CH−5746
TCGA−KK−A5A1

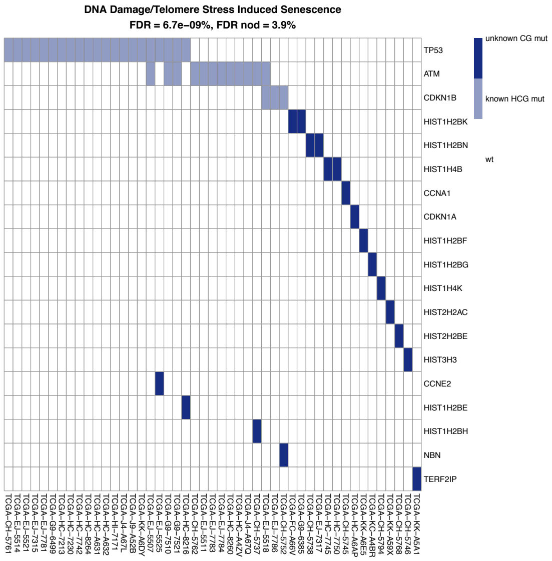
DNA Damage/Telomere Stress Induced Senescence
FDR = 6.7e−09%, FDR nod = 3.9%
TP53
ATM
CDKN1B
HIST1H2BK
HIST1H2BN
HIST1H4B
CCNA1
CDKN1A
HIST1H2BF
HIST1H2BG
HIST1H4K
HIST2H2AC
HIST2H2BE
HIST3H3
CCNE2
HIST1H2BE
HIST1H2BH
NBN
TERF2IP
unknown CG mut
known HCG mut
wt

### SKCM

**Activation Of Matrix Metalloproteinases**
**FDR = 0.022%, FDR nod = 0.093%**

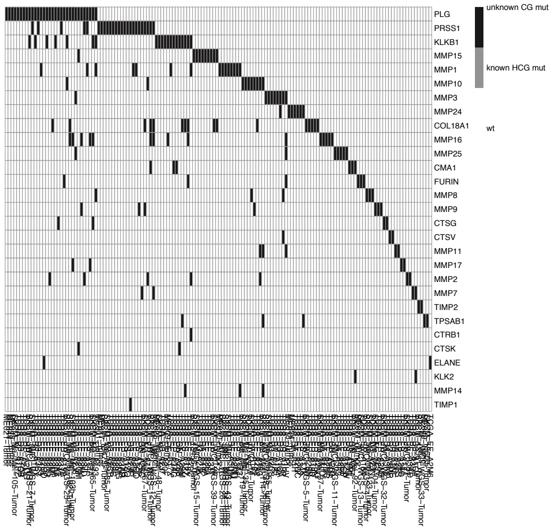


**IRF3 Mediated Activation Of Type 1 IFN**
**FDR = 1.2e−06%, FDR nod = 1.2e−05%**

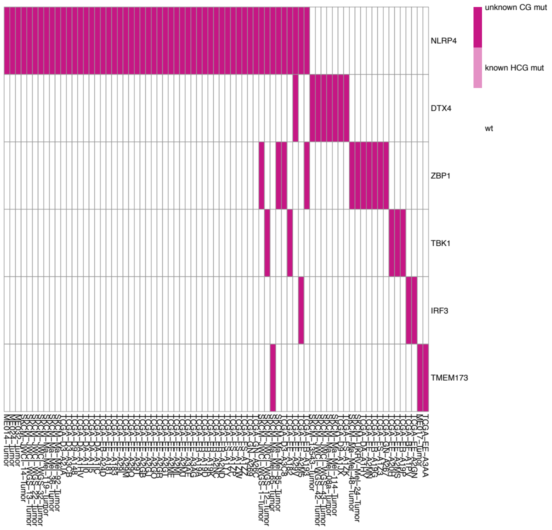


**Mitochondrial Beta Oxidation Of Medium Chain Saturated Fatty Acids**
**FDR = 0.42%, FDR nod = 1.5%**

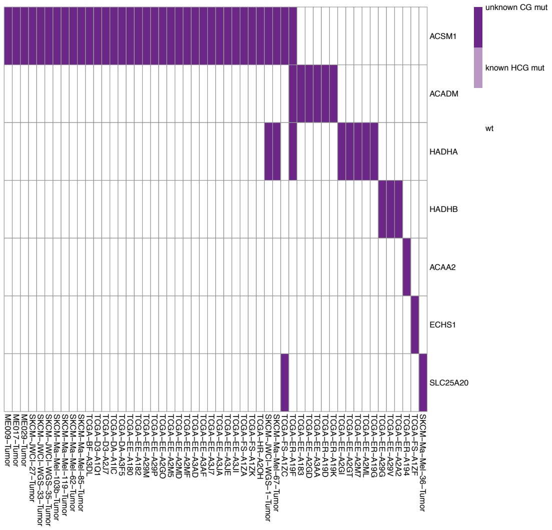


**TP53 Network**
**FDR = 4.2e−12%, FDR nod = 2.1e−05%**

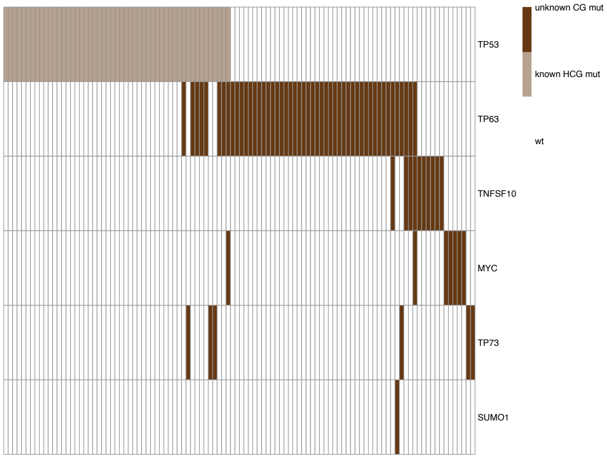


**TP53 Regulates Transcription Of Death Receptors And Ligands**
**FDR = 1.6e−11%, FDR nod = 0.43%**

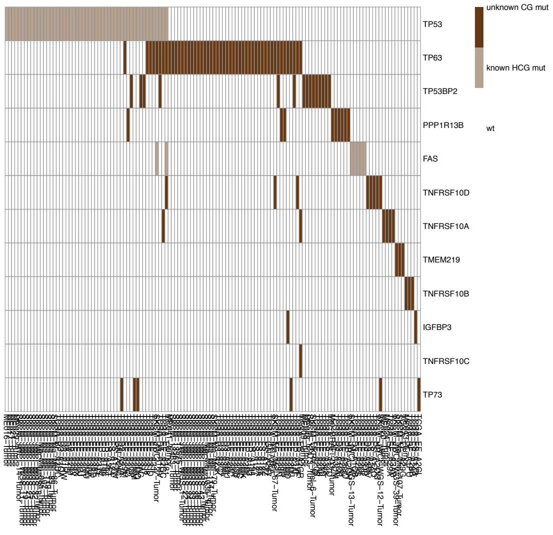


**[CGB Protein Results In Increased Activity Of LHCGR Protein]**
**de Results In Decreased Activity Of PTGS2 Protein] Inhibits The Reaction [CGB Protein Results In Increased Chen**
**(3, 2 F)quinazoline 1,3 Diamine Binds To F2R Protein] Promotes The Reaction [CGB Protein Results In Increased**
**ne Inhibits The Reaction [CGB Protein Results In Increased Chemical Synthesis Of Androgens]**
**trazine Inhibits The Reaction [CGB Protein Results In Increased Secretion Of Cyclic AMP]**
**forsin Promotes The Reaction [CGB Protein Results In Increased Abundance Of Cyclic AMP]**
**Reaction [[Gonadotropins, Equine Co Treated With CGB Protein] Results In Increased Abundance Of Progesteron**
**ibits The Reaction [[CGB Protein Co Treated With Colforsin] Results In Increased Abundance Of Cyclic AMP]**
**Protein Promotes The Reaction [CGB Protein Results In Increased Abundance Of Progesterone]**
**ole Metabolite Inhibits The Reaction [CGB Protein Results In Increased Abundance Of Testosterone]**
**se Chloride Inhibits The Reaction [CGB Protein Results In Increased Expression Of STAR Protein]**
**terone Promotes The Reaction [CGB Protein Results In Increased Expression Of VEGFA MRNA]**
**one Enanthate Inhibits The Reaction [CGB Protein Results In Increased Abundance Of Androgens]**
**FDR = 0.19%, FDR nod = 0.72%**

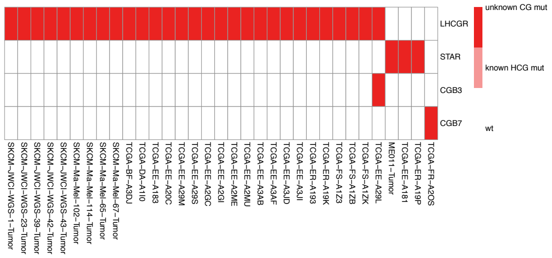

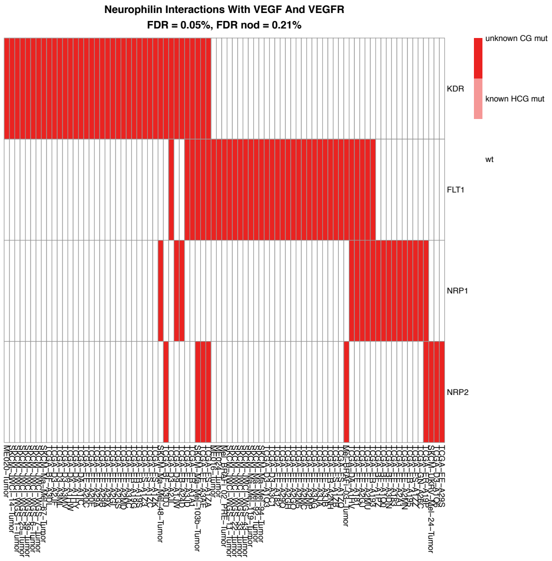
Neurophilin Interactions With VEGF And VEGFR
FDR = 0.05%, FDR nod = 0.21%
KDR
FLT1
NRP1
NRP2
unknown CG mut
known HCG mut
wt

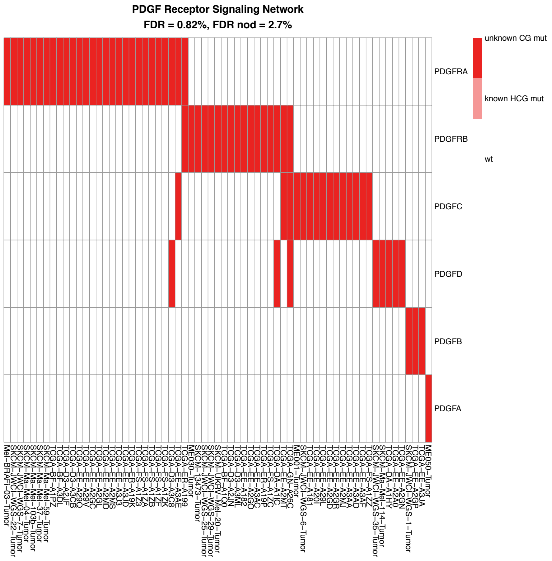
PDGF Receptor Signaling Network
FDR = 0.82%, FDR nod = 2.7%
PDGFRA
PDGFRB
PDGFC
PDGFD
PDGFB
PDGFA
unknown CG mut
known HCG mut
wt

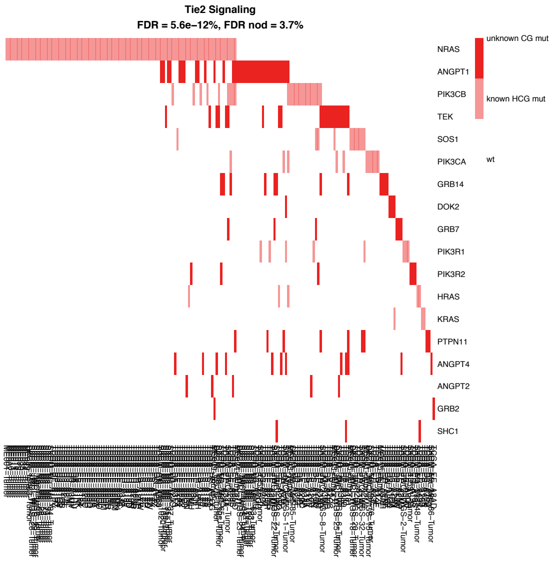
Tie2 Signaling
FDR = 5.6e−12%, FDR nod = 3.7%
NRAS
ANGPT1
PIK3CB
TEK
SOS1
PIK3CA
GRB14
DOK2
GRB7
PIK3R1
PIK3R2
HRAS
KRAS
PTPN11
ANGPT4
ANGPT2
GRB2
SHC1
unknown CG mut
known HCG mut
wt

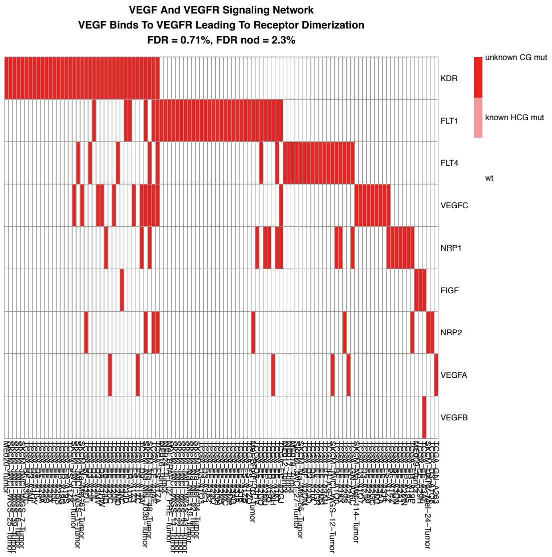
VEGF And VEGFR Signaling Network
VEGF Binds To VEGFR Leading To Receptor Dimerization
FDR = 0.71%, FDR nod = 2.3%
KDR
FLT1
FLT4
VEGFC
NRP1
FIGF
NRP2
VEGFA
VEGFB
unknown CG mut
known HCG mut
wt

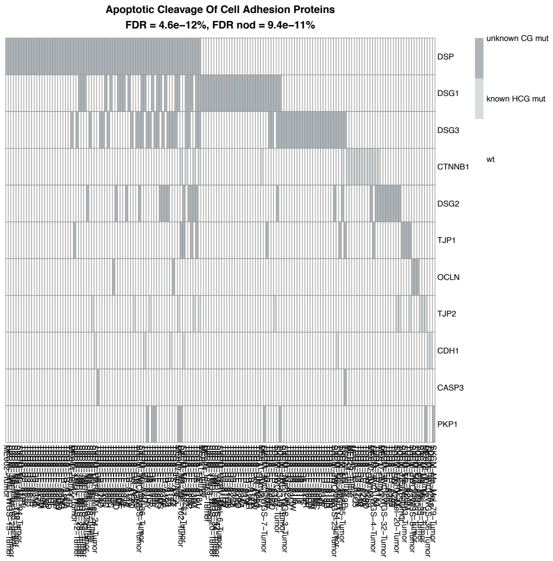
Apoptotic Cleavage Of Cell Adhesion Proteins
FDR = 4.6e−12%, FDR nod = 9.4e−11%
DSP
DSG1
DSG3
CTNNB1
DSG2
TJP1
OCLN
TJP2
CDH1
CASP3
PKP1
unknown CG mut
known HCG mut
wt

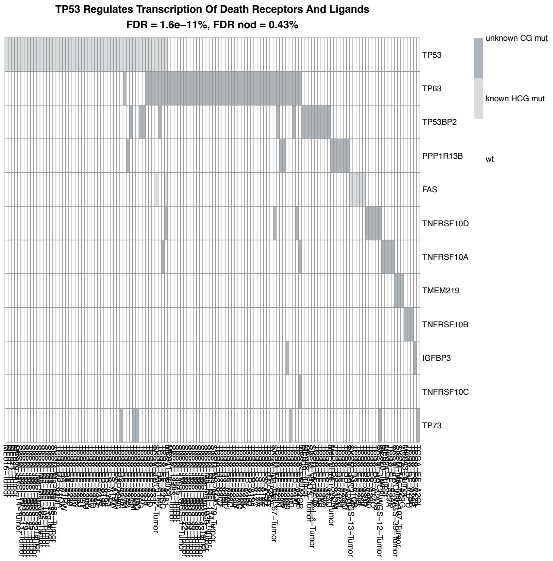
TP53 Regulates Transcription Of Death Receptors And Ligands
FDR = 1.6e−11%, FDR nod = 0.43%
TP53
TP63
TP53BP2
PPP1R13B
FAS
TNFRSF10D
TNFRSF10A
TMEM219
TNFRSF10B
IGFBP3
TNFRSF10C
TP73
unknown CG mut
known HCG mut
wt

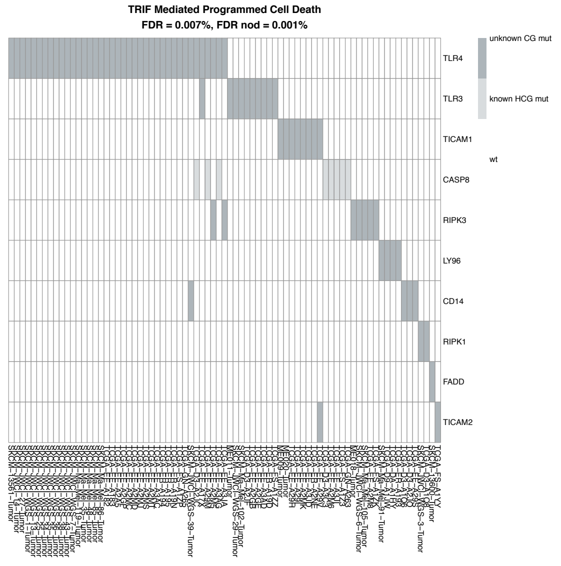
TRIF Mediated Programmed Cell Death
FDR = 0.007%, FDR nod = 0.001%
TLR4
TLR3
TICAM1
CASP8
RIPK3
LY96
CD14
RIPK1
FADD
TICAM2
unknown CG mut
known HCG mut
wt

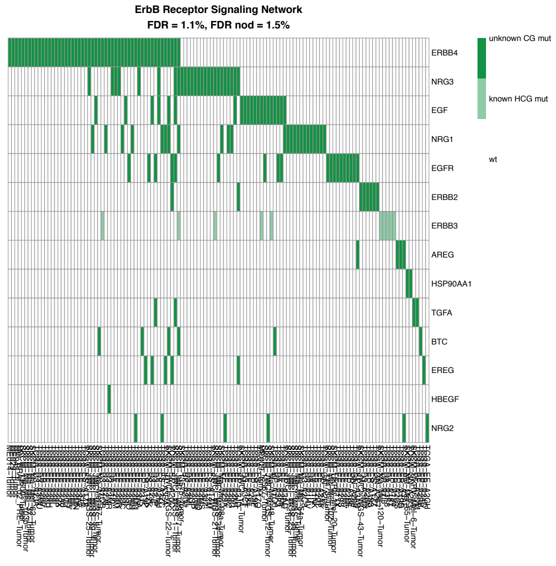
ErbB Receptor Signaling Network
FDR = 1.1%, FDR nod = 1.5%
ERBB4
NRG3
EGF
NRG1
EGFR
ERBB2
ERBB3
AREG
HSP90AA1
TGFA
BTC
EREG
HBEGF
NRG2
unknown CG mut
known HCG mut
wt

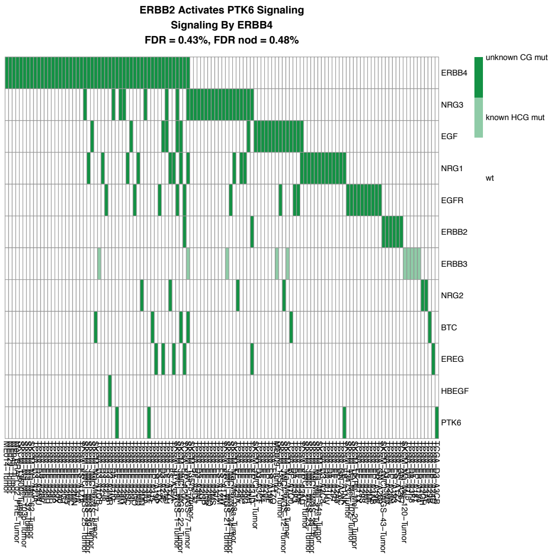
ERBB2 Activates PTK6 Signaling
Signaling By ERBB4
FDR = 0.43%, FDR nod = 0.48%
ERBB4
NRG3
EGF
NRG1
EGFR
ERBB2
ERBB3
NRG2
BTC
EREG
HBEGF
PTK6
unknown CG mut
known HCG mut
wt

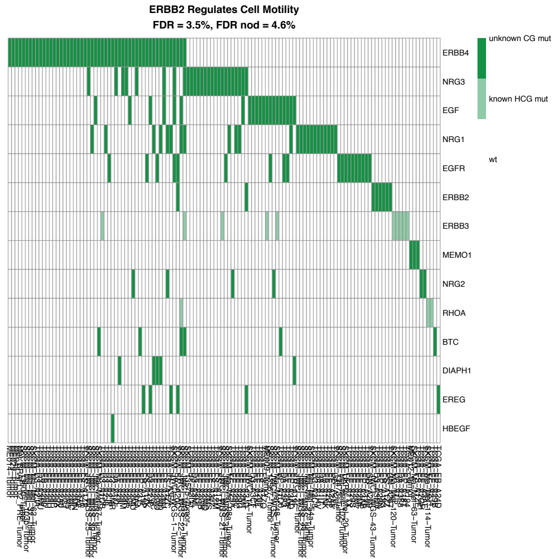
ERBB2 Regulates Cell Motility
FDR = 3.5%, FDR nod = 4.6%
ERBB4
NRG3
EGF
NRG1
EGFR
ERBB2
ERBB3
MEMO1
NRG2
RHOA
BTC
DIAPH1
EREG
HBEGF
unknown CG mut
known HCG mut
wt

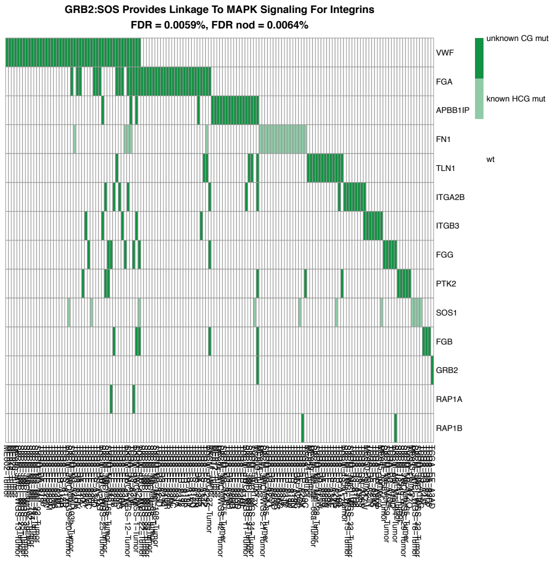
GRB2:SOS Provides Linkage To MAPK Signaling For Integrins
FDR = 0.0059%, FDR nod = 0.0064%
VWF
FGA
APBB1IP
FN1
TLN1
ITGA2B
ITGB3
FGG
PTK2
SOS1
FGB
GRB2
RAP1A
RAP1B
unknown CG mut
known HCG mut
wt

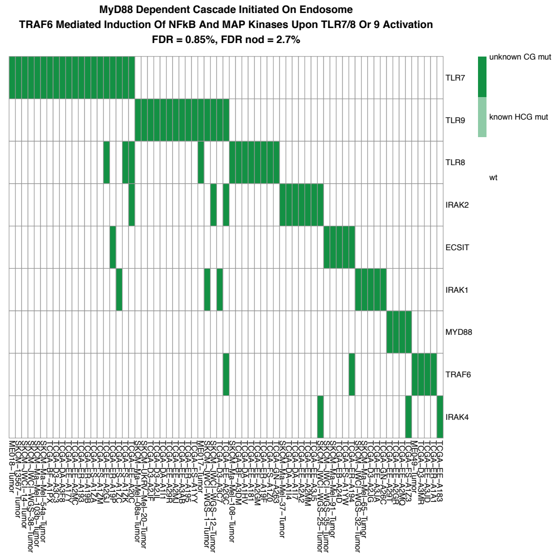
MyD88 Dependent Cascade Initiated On Endosome
TRAF6 Mediated Induction Of NFkB And MAP Kinases Upon TLR7/8 Or 9 Activation
FDR = 0.85%, FDR nod = 2.7%
TLR7
TLR9
TLR8
IRAK2
ECSIT
IRAK1
MYD88
TRAF6
IRAK4
unknown CG mut
known HCG mut
wt

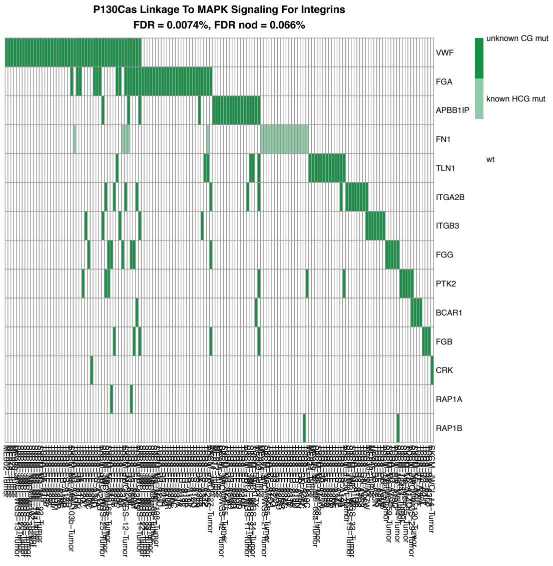
P130Cas Linkage To MAPK Signaling For Integrins
FDR = 0.0074%, FDR nod = 0.066%
VWF
FGA
APBB1IP
FN1
TLN1
ITGA2B
ITGB3
FGG
PTK2
BCAR1
FGB
CRK
RAP1A
RAP1B
unknown CG mut
known HCG mut
wt

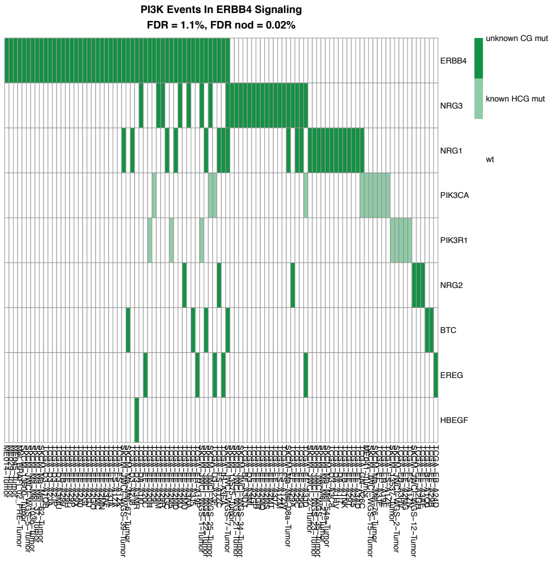
PI3K Events In ERBB4 Signaling
FDR = 1.1%, FDR nod = 0.02%
ERBB4
NRG3
NRG1
PIK3CA
PIK3R1
NRG2
BTC
EREG
HBEGF
unknown CG mut
known HCG mut
wt

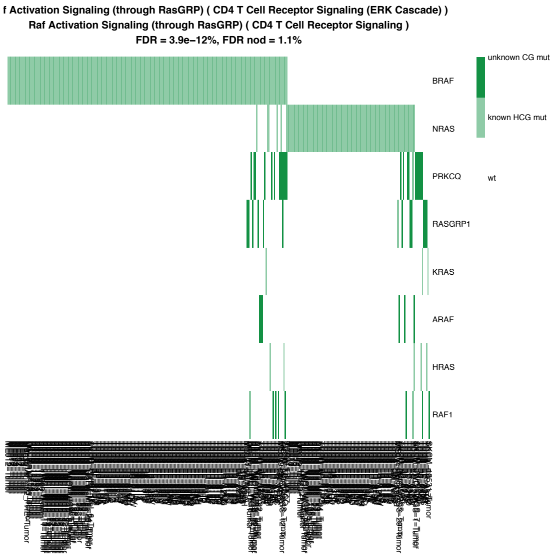
f Activation Signaling (through RasGRP) ( CD4 T Cell Receptor Signaling (ERK Cascade) )
Raf Activation Signaling (through RasGRP) ( CD4 T Cell Receptor Signaling )
FDR = 3.9e−12%, FDR nod = 1.1%
BRAF
NRAS
PRKCQ
RASGRP1
KRAS
ARAF
HRAS
RAF1
unknown CG mut
known HCG mut
wt

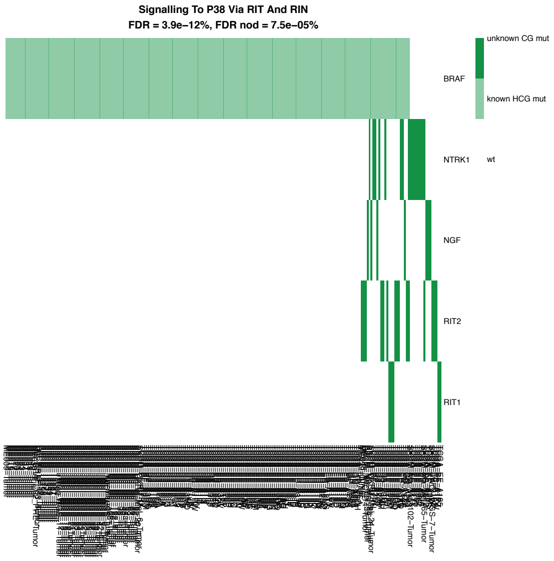
Signalling To P38 Via RIT And RIN
FDR = 3.9e−12%, FDR nod = 7.5e−05%
BRAF
NTRK1
NGF
RIT2
RIT1
unknown CG mut
known HCG mut
wt

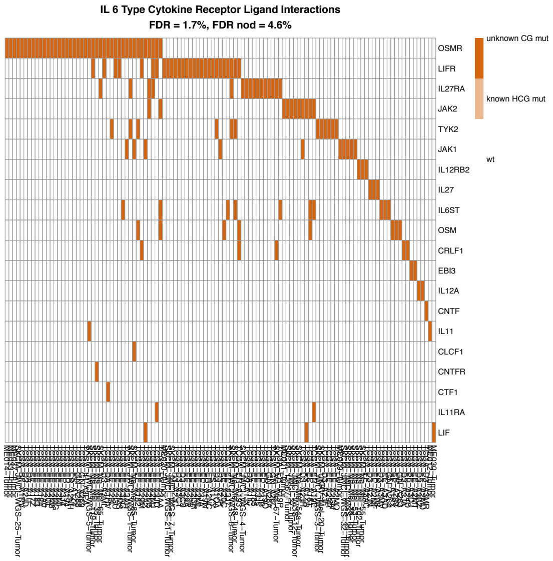
IL 6 Type Cytokine Receptor Ligand Interactions
FDR = 1.7%, FDR nod = 4.6%
OSMR
LIFR
IL27RA
JAK2
TYK2
JAK1
IL12RB2
IL27
IL6ST
OSM
CRLF1
EBI3
IL12A
CNTF
IL11
CLCF1
CNTFR
CTF1
IL11RA
LIF
unknown CG mut
known HCG mut
wt

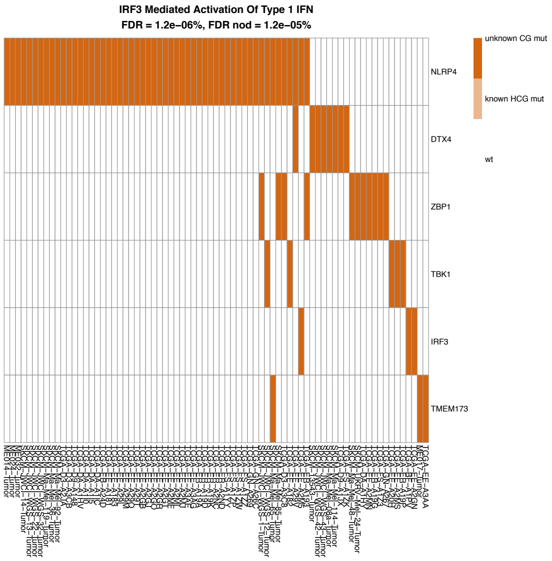
IRF3 Mediated Activation Of Type 1 IFN
FDR = 1.2e−06%, FDR nod = 1.2e−05%
NLRP4
DTX4
ZBP1
TBK1
IRF3
TMEM173
unknown CG mut
known HCG mut
wt
